# A Comprehensive Overview of the Physical Health of the Adolescent Brain Cognitive Development Study (ABCD) Cohort at Baseline

**DOI:** 10.1101/2021.06.30.450555

**Authors:** Clare E. Palmer, Chandni Sheth, Andrew T. Marshall, Shana Adise, Fiona C. Baker, Linda Chang, Duncan B. Clark, Rada K. Dagher, Gayathri J. Dowling, Marybel R. Gonzalez, Frank Haist, Megan M. Herting, Rebekah S. Huber, Terry L. Jernigan, Kimberly LeBlanc, Karen Lee, Krista M. Lisdahl, Gretchen Neigh, Megan W. Patterson, Perry Renshaw, Kyung E. Rhee, Susan Tapert, Wesley K. Thompson, Kristina Uban, Elizabeth R. Sowell, Deborah Yurgelun-Todd

## Abstract

Physical health in childhood is crucial for neurobiological as well as overall development, and can shape long-term outcomes into adulthood. The landmark, longitudinal Adolescent Brain Cognitive Development Study^SM^ (ABCD study®), was designed to investigate brain development and health in almost 12,000 youth who were recruited when they were 9-10 years old and will be followed through adolescence and early adulthood. The overall goal of this paper is to provide descriptive analyses of physical health measures in the ABCD study at baseline, including but not limited to sleep, physical activity and sports involvement, and body mass index, and how these measures vary across demographic groups. This paper outlines how the physical health of the ABCD sample corresponds with that of the US population and highlights important avenues for health disparity research. This manuscript will provide important information for ABCD users and help guide analyses investigating physical health as it pertains to adolescent and young adult development.

## Introduction

It has been increasingly recognized in neurodevelopmental research, policy, and clinical practice communities that early and middle childhood years provide the physical, cognitive, and social-emotional foundation for lifelong health and well-being [1]. Experiences in middle childhood (6-12 years) have been shown to be critically important for a child’s physical development as well as their cognitive, social, and emotional growth and development. Research on a number of adult health and medical conditions points to pre-disease pathways that have their beginnings in early and middle childhood [1]. There are a number of factors including sleep, physical activity, and head injury, that are associated with outcomes in later life. Our understanding of the magnitude of these effects and how these factors interact with one another has been constrained in part by the limited number of large normative samples in this age group with comprehensive assessment of physical health measures. Here we present descriptive analyses of the physical health measures in almost 12,000 youth ages 9-10. We also highlight how this sample corresponds to normative data available for the US population. We propose that these data will aid in the identification of disparities in physical health early in life which in turn may have significant impact on the development of future interventions [2, 3].

During middle childhood there are multiple changes that occur across domains including physical and mental health and cognition [4]. Furthermore, exercise regimes and attitudes towards physical fitness, tobacco use, alcohol use, dietary habits, and coping with stress are important lifestyle characteristics that emerge during middle childhood (National Research Council Panel to Review the Status of Basic Research on School-Age 1984). Physical health in middle childhood is thought to be determined by interactions between a child’s biological function, socioeconomic environment, and the evolution of their lifestyle behaviors [1]. School-age children are generally considered a healthy population and their physical well-being is generally assumed [5]. However, this developmental period may be the most sensitive period for the development of many of the functional patterns that significantly influence physical health status in later life. To date, there have been few comprehensive large-scale studies that have reported physical health data from a cross national middle childhood sample.

The Adolescent Brain Cognitive Development Study^SM^ (ABCD study®) incorporates a broad range of measures assessing predictors and outcomes related to physical health in children, and provides the necessary variability and statistical power to disentangle the extent to which sex and sociodemographic factors contribute to the shared covariance across different physical health measures. The physical health battery at baseline measures a wide-ranging array of constructs relevant to child development and includes both caregiver- and youth-reported assessments [6]. Early developmental physical health measures include prematurity, birth weight, early developmental milestones, medical problems during birth and pregnancy, and prenatal substance exposure. Current physical health measures assess sleep, physical activity and sports involvement, body mass index (BMI), medical problems, traumatic brain injury (TBI), and pubertal development from both self-report of pubertal maturation and salivary biomarkers of pubertal hormones (described in detail in [7]).

Importantly, health disparities across different sociodemographic groups can lead to divergent outcomes in childhood and adolescent physical health and these health disparities have not always been considered in analytic approaches and research study reports. Socioeconomic disadvantage and contextual factors associated with race or ethnicity can mediate the effects of several physical health measures on developmental outcomes due to, for example, lack of access to healthcare and systemic biases [8]. Here, we aim to describe how both good and poor physical health is distributed across sociodemographic groups in order to identify what disparities may be present and may be ameliorated. However, extensive analyses understanding the factors contributing to these disparities are beyond the scope of this paper.

The ABCD Study provides a unique, unprecedented opportunity to explore physical health measures in a large, diverse, typically developing sample, addressing limitations of previous studies that often include smaller, less representative samples. Here, we aim to provide a comprehensive overview of the physical health of the ABCD Study cohort at baseline and how physical health varied across different sociodemographic groups at the baseline time point (i.e., when children were 9- to 10- years-old). In addition, the manuscript will compare physical health measures with current clinical guidelines and norms when appropriate, potentially facilitating clinical recommendations and informing national standards of physical health in this age group. Overall, this manuscript will provide important information for ABCD users and help guide analyses investigating the potential impact of physical health on adolescent and young adult growth and development.

## Methods

### Sample

The ABCD Study is a large-scale, 10-year longitudinal study involving 21 data collection sites across the United States (ABCDStudy.org). The ABCD sample was largely recruited through public, private, and charter elementary schools. ABCD adopted a population neuroscience approach to recruitment [9] by employing epidemiologically informed procedures to ensure demographic variation in its sample that would mirror the variation in the US population of 9- and 10-year-olds [10]. The consortium enrolled 11,880 children aged 9-10 years and data from these subjects came from the ABCD 2.0.1 data release (https://data-archive.nimh.nih.gov/abcd), which included baseline data (i.e., cross-sectional). Centralized institutional review board (IRB) approval was obtained from the University of California San Diego IRB. Study sites obtained approval from their local IRBs. Caregivers provided written consent and children provided written assent. All ethical regulations were complied with during data collection and analysis. Potential participants were excluded for the following reasons: child not fluent in English, MRI contraindication (e.g., irremovable ferromagnetic implants or dental appliances, claustrophobia, major neurological disorder, gestational age less than 28 weeks or birth weight less than 1,200 grams (g), birth complications that resulted in hospitalization for more than one month, uncorrected vision, or current diagnosis of schizophrenia, autism spectrum disorder (moderate, severe), mental retardation/intellectual disability, or alcohol/substance use disorder. The ABCD sample included: non-twin siblings: 1600; Twins: 2100 (1050 pairs); 30 triplets (10 sets); and 8150 singletons. Greater details on recruitment strategies and other demographic characteristics of the ABCD sample have been published previously [10].

### Physical Health Measures

Physical health measures were comprised of questionnaires evaluating past and current physical health. In addition, some objective measures including height, weight, waist circumference were collected at the time of the baseline visit.

#### Developmental history measures

The Developmental History Questionnaire (DHQ), originally developed by the Adolescent Component of the National Comorbidity Survey [11–13] was completed by the caregiver at baseline to obtain information on birth weight, gestational age, early developmental milestones, medical problems during birth and pregnancy, and prenatal substance exposure.

##### Birth weight and prematurity

Caregivers were asked whether their child was born prematurely and, if yes, how many weeks premature. Birth weight (pounds (lbs), ounces) was converted to kilograms (kg) and categorized based on the Centers for Disease Control guidelines as extremely low birth weight (ELBW, < 1,000 g), very low birth weight (VLBW, 1000 to < 1,500 g), low birth weight (LBW, 1500 to < 2,500 g), average/normal birth weight (NBW, 2500 to < 4,000 g), and high birth weight (HBW, > 4,000 g). One exclusionary criterion for study participation was a birth weight of <1200 g; however, there are a few participants who report this low birth weight. Nevertheless, given the low number of participants in the ELBW category, we combined the ELBW and VLBW categories for analysis. Generally, an infant is defined as premature if birth occurred before 37 weeks of gestation. Consistent with the current literature, we further categorized participants into extremely preterm (EPI, if born less than 28 weeks), very preterm (VPI, 28 to <32 weeks), moderate preterm (MPI, 32 to <34 weeks), and late preterm (LPI, 34 to 37 weeks) [14]. One child born before 35 weeks and weighing 14 lbs was removed, likely due to measurement error.

##### Early developmental milestones

Caregivers were asked about the age at which their child began to roll over (delayed if after 6 months), sit without assistance (delayed if after 9 months), walk without assistance (delayed if after 18 months), and say his/her first word (delayed if after 12 months) [15]. Outliers due to measurement error were excluded from analysis by setting an upper threshold of 48 months for rolling over, sitting and walking, and 60 months for first word. The questionnaire also assessed caregiver concern regarding motor and speech delays, with caregivers asked to compare their child’s development to that of other children (earlier, average, later).

##### Medical problems during birth and pregnancy

Caregivers were asked about any complications during birth and pregnancy using dropdown lists. Medical problems during birth were: blue at birth, slow heartbeat, did not breathe at first, convulsions, jaundice needing treatment, required oxygen, required blood transfusion, and Rh incompatibility. Medical problems during pregnancy were severe nausea and vomiting extending past the 6^th^ month accompanied by weight loss, heavy bleeding requiring bed rest or special treatment, (pre)eclampsia/toxemia, severe gall bladder attack, persistent proteinuria, rubella during first 3 months of pregnancy, severe anemia, urinary tract infections, pregnancy related diabetes, pregnancy related high blood pressure, problems with the placenta, accident or injury, or any other conditions. A summary variable was calculated for each of these as a Total Problems score based on a sum of the number of complications endorsed (separately for birth and pregnancy). A categorical variable was created to summarize those who had no problems, one problem, or more than one problem.

##### Prenatal substance exposure

Caregivers were asked about the biological mother’s substance use (i.e., tobacco, alcohol, marijuana, cocaine/crack, heroin/morphine, oxycontin, and any other drugs) before the mother found out about the pregnancy (but could have been pregnant) and once the mother knew about the pregnancy. If drug use was endorsed, follow-up questions were asked about the frequency and quantity of use. In addition to the above drugs of abuse, caffeine use from conception until delivery was also measured. Here, we measured endorsement of use of each substance before and after pregnancy by combining the two variables asking about pre and post pregnancy recognition of substance exposure into a single variable with three categories: 1) pre-recognition no use + post-recognition no use; 2) pre-recognition use + post-recognition no use; 3) pre-recognition use + post-recognition use. There were 59 subjects who endorsed no use before pregnancy recognition but use after recognition; these subjects were excluded from analysis given the small number in this category. This graded exposure variable dependent on timing of pregnancy recognition was computed for alcohol, tobacco, cannabis (aka marijuana), and other substance exposure.

#### Current measures

Caregivers and youth completed several other questionnaires about the child’s current physical health.

##### Sleep

The Sleep Disturbance Scale for Children (SDSC) was used to assess sleep duration and sleep disturbance symptoms at the baseline visit [16]. The SDSC (26 items) assesses frequency of disorders of initiating and maintaining sleep, sleep breathing disorders, disorders of arousal, sleep-wake transition disorders, disorders of excessive somnolence, and sleep hyperhidrosis in the past 6 months. We used the overall sleep-wake disturbance score, which was the sum of all items, with higher scores reflecting a greater clinical severity of sleep disturbance. We excluded 8 subjects who scored greater than 87, which was the maximum score attained in the original study [16]. A cut-off score of 39 is recommended as a threshold for identifying children with disturbed sleep [16]. The individual item from the SDSC ‘How many hours of sleep does your child get on most nights?’ was used as the measure for typical total sleep duration. Possible responses were: 1) 9-11 hours, 2) 8-9 hours, 3) 7-8 hours, 4) 5-7 hours, and 5) Less than 5 hours. As very few participants endorsed fewer than 5 hours of sleep per night, the smallest two categories were combined to form a category of “less than 7 hours”.

##### Physical activity and sports activity

Three items from the Youth Risk Behavior Survey (YRBS) served as a measure of physical activity. The YRBS questionnaire was modified from the Youth Risk Behavior Survey [6, 17]. Youth were asked the number of days in the past week that they exercised for at least 60 minutes per day and the number of days in the past week that they engaged in exercises to strengthen or tone their muscles. The questionnaire also asks about how many days per week the youth has physical education (PE) class in school. The Sports and Activities Involvement Questionnaire (SAI-Q), modeled after the assessment developed for the Vermont Health and Behavioral Questionnaire (VHBQ) and the Dutch Health Behavioral Questionnaire (DHBQ) [18], measured lifetime and past year involvement in 23 different sports, activities such as music and dance, and other hobbies. A summary score of time spent participating in sports (excluding all items associated with activities or hobbies that do not require physical activity) was calculated for each participant. For each sport, caregivers were asked to report: 1) time spent (mins) per session (tspent); 2) number of days per week when participating (perwk); 3) number of months per year (nmonth); 4) number of years participated (at baseline visit); and, 5) whether the child participated in the last 12 months. For each sport endorsed in the past 12 months, the mean participation hours per week was calculated using the following formula: past year mean hours per week per sport = (tspent*perwk*nmonth)/52)/60. This value was summed across all sports endorsed to create a total average time spent participating in sports over the past year for each participant.

##### BMI/Weight Status

Anthropometric measurements of height and weight were taken as the average of up to 3 separate measures using professional grade equipment (e.g., physician weight beam scale with height rod). Body Mass Index (BMI) (kg/m^2^) was calculated according to convention and converted to age- and sex-specific percentiles using the CDC 2000 Growth Chart SAS (SAS Institute, Inc, Cary, NC)[19]. The CDC age- and sex-adjusted percentiles were used to classify participants as underweight (i.e., <5^th^ %ile) healthy weight (≥5^th^ %ile to < 85^th^ %ile), overweight (≥ 85^th^ %ile to <95^th^ %ile), obese (≥95^th^ %ile) [20, 21]. Subjects with potential measurement error who had biologically implausible BMIs (e.g., extremely small BMI (n=23) and extremely large BMI (n=5)) were excluded. These extreme values were identified using cut offs of <−4 and >8 of the modified BMI z-scores, which express an individual’s BMI relative to the median BMI at that age and sex [22]. These scores were calculated using SAS code from the CDC: https://www.cdc.gov/nccdphp/dnpao/growthcharts/resources/sas.htm#reference. Statistical analyses were conducted using the above noted weight status classifications.

##### Medical Problems (lifetime)

A caregiver-report medical history questionnaire about the youth was derived from the Missouri Assessment of Genetics Interview for Children (MAGIC) Health Services Utilization Questionnaire [23]. At baseline, the questionnaire covered both past year and lifetime conditions including the following: asthma, allergies, bronchitis, leukemia, cerebral palsy, diabetes, epilepsy, hearing loss, kidney disease, lead poisoning, muscular dystrophy, multiple sclerosis, vision problems, heart problems, sickle cell anemia, headache, operation, and other illnesses. A Total Problems summary score was created by summing endorsed conditions for each participant. A categorical variable was created based on whether participants endorsed none, one, or more than one problem. Participants endorsing more than 6 medical problems were excluded due to questions regarding the validity of the data (n=47).

##### Traumatic Brain Injury or Head injury (lifetime)

Caregivers reported on the youth’s lifetime history of head injury using the Modified Ohio State University TBI Screen-Short Version [24, 25]. This questionnaire asks whether their child had been to the emergency room due to an injury to the head or neck, whether the child had injured their neck in a fall or from being hit by something or from being in a fight or from a gunshot wound). A positive response to an occurrence question was followed up with questions to determine loss of consciousness (LOC), memory loss, and other details about the event (e.g., age at time of injury). A summary variable with the worst injury overall is generated as follows: Improbable TBI (responses to all head injuries are ‘no’); TBI without LOC or memory loss (response to at least one question about head injury is ‘yes’ but all responses to LOC and memory loss are “no”); possible mild TBI (TBI without LOC but with memory loss); mild TBI (TBI with LOC less than 30 minutes); moderate TBI (TBI with LOC between 30 minutes and 24 hours), or severe TBI (TBI with LOC greater than 24 hours).

#### Statistical Analysis

Chi-squared tests were used to describe the distribution of these physical health measures across different demographic factors using one individual per family to control for family relatedness. For categorical measures, these tests determined whether the proportion of subjects at each level of the categorical variable depended on each demographic factor. For continuous measures, we used clinical guidelines as a threshold to binarize these measures (more details below). We then determined whether a subject was more or less likely to be above or below these guidelines as a function of these demographic factors. For all measures, demographic factors of sex at birth, household income, highest parental education, self-declared race (White/Black/Asian/Other) and ethnicity (Hispanic: Yes/No) were analyzed as variables of interest. For the current, non-developmental (retrospective) measures (weight status, sleep, physical activity, sports and activity involvement, total medical problems and TBI), additional predictors of age and pubertal development were assessed. Age in months was divided into quartiles for chi-squared analyses. Pubertal development was measured using the caregiver report Pubertal Development Scale (PDS) [26]. Baseline information on pubertal development in the ABCD study has been previously reported in detail [7]. Due to very few numbers in the post-puberty category, this was combined with the late-puberty category to generate 4 levels of pubertal development (pre, early, mid, late/post). In instances where p-values could not be estimated from the chi-squared distribution, Monte Carlo simulations were used.

Prior to completing statistical analysis, siblings were randomly removed from the sample to keep one participant per family. Contingency tables showing the proportion of subjects across levels of the dependent variables stratified by each demographic factor can be found for the whole sample in Supplementary Tables 1-20 and for the independent sample (used for statistical analysis) in Supplementary Tables 21-40. All contingency tables additionally show the proportion of subjects at each level of the dependent variable using propensity-weights based on nationally-representative controls from the American Community Survey (ACS) as calculated in [27] and applied in the ABCD Study Data Exploration Analysis Portal (DEAP)(https://deap.nimhda.org). Results of the chi-squared analyses and post-hoc pairwise comparisons can be found in Supplementary Tables 41-60. All post-hoc pairwise p-values have been corrected using the Bonferroni method. For all main effects, an alpha level of 0.00045 was used to determine significance controlling for a false positive rate of 5% across 110 independent tests. Distributions of each variable of interest stratified by the sociodemographic factors can be found in Supplementary Figures 1-20.

## Results

### Developmental Measures

#### Birth weight & Prematurity

Figure 1 illustrates the distribution of males and females in the baseline sample based on prematurity and birthweight. Most participants were born at full term of pregnancy with 86.4% of female infants and 86.2% of male infants being born at term; 9.8% of both male and female infants, late preterm; 2.5% of female infants and 2.8% of male infants, moderate preterm; 0.9% of both male and female infants, very preterm; and 0.3% of both male and female infants, extremely preterm. In terms of birthweight, most participants had normal birthweight with 75.9% of female infants and 75.2% of male infants having normal birthweight; 6.4% of female infants and 10.6% of male infants, high birthweight; 16.2% of female infants and 13.3% of male infants, low birthweight; 1.5% of female infants and 0.9% of male infants, very low birthweight; and only 0.1% of female infants and 0.0% of male infants, extremely low birthweight.

**Figure 1.**
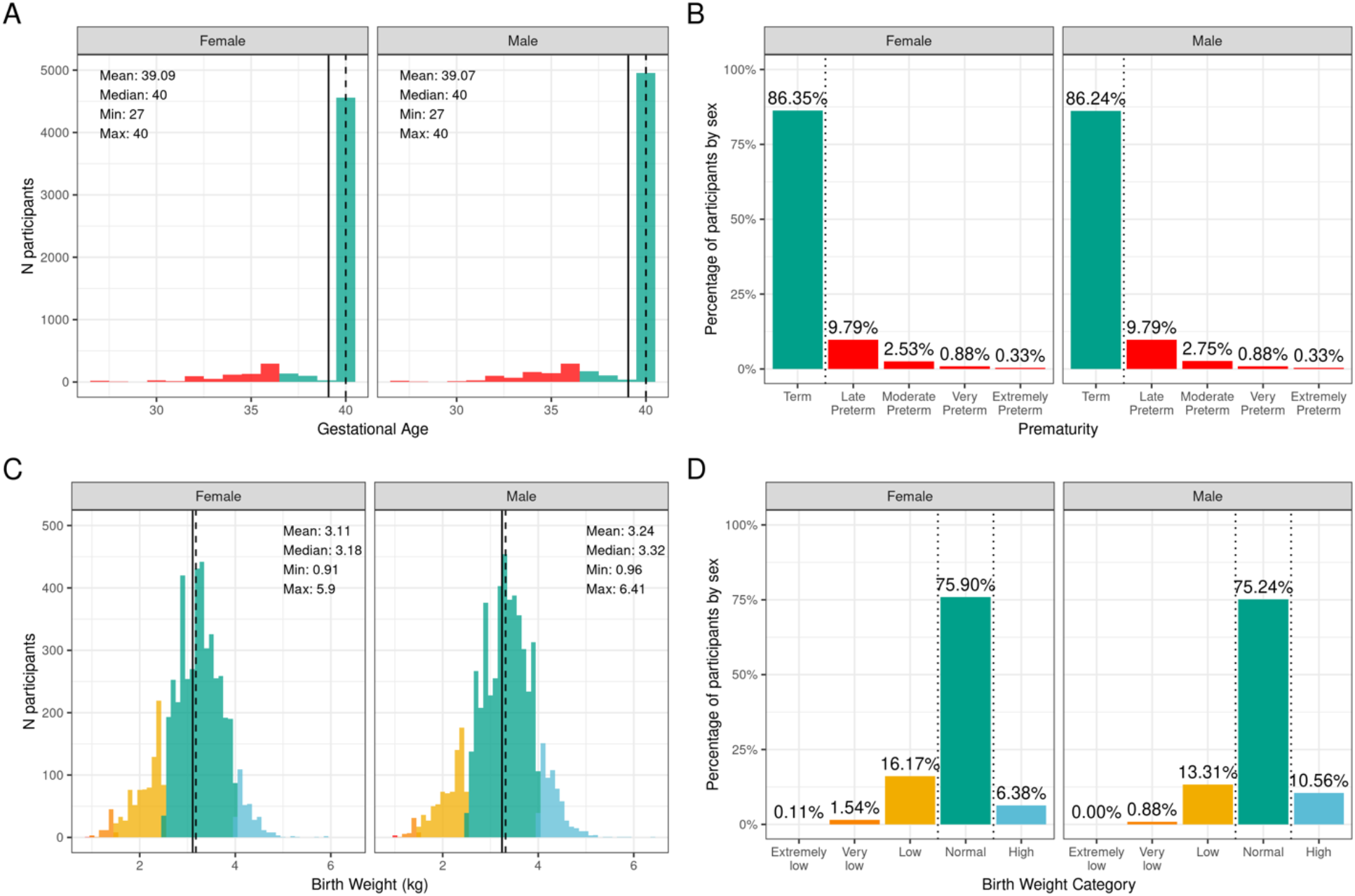
Distribution of gestational age and birth weight in the ABCD sample. A) Distribution of gestational ages in the sample in weeks stratified by sex assigned at birth. Participants born at gestational age <37 weeks are considered premature (red) and ≥37 weeks are considered born at term (green). B) Percentage of participants stratified by sex within different prematurity categories. C) Distribution of birth weight in kilograms (kg) across the sample color coded by birth weight category: high (blue), normal (green), low (yellow), very low (orange) and extremely low (red). D) Percentage of participants stratified by sex within each birth weight category.

Supplementary Figures 1 and 2 show the probability distribution of prematurity categories and birthweight (kg), respectively, for all levels of the sociodemographic factors (race, ethnicity, household income, and parent educational status). Supplementary Table 21 shows the distribution of participants and weighted frequencies across each of the four categories of birthweight (very low, low, normal, and high) stratified by demographics (sex, race, Hispanic status, household income level, and education level). As seen in Supplementary Table 41, the distributions of birth weight differed significantly by sex at birth (χ^2^(3,8536)=59.9, p=6.14e-13), such that females were more likely to be in the low birthweight category and males were more likely to be in the high birthweight category. There were also significant associations with Hispanic status (χ^2^(3,8536)=22.3,p=8.88e-05). Supplementary Table 22 shows the distribution and weighted frequencies of participants across each of the five categories regarding their maturity at birth (Term, Late Preterm, Moderate Preterm, Very Preterm and Extremely Preterm), stratified by demographics (sex, race, Hispanic status, household income level, and education level). The distributions of prematurity status were not different between boys and girls. The dependence on household income (χ^2^(12, 8783)=37.1, p=5e-04), and the parents’ education level (χ^2^(16, 8783)=53.1, p=5e-04) just fell short of corrected statistical significance (Supplementary Table 42).

#### Developmental Milestones

Figure 2 illustrates the distribution and percentage of males and females in the baseline sample meeting standard guidelines for developmental milestones (by age in months). Most participants met developmental milestones on time with 98.8% of both males and females rolling over by 6 months of age; 96.5% of females and 95.0% of males sitting without assistance by 9 months of age; 93.0% of females and 88.2% of males saying her/his first word by 12 months of age; and 97.5% of females and 96.1% of males walking without assistance by 18 months of age.

**Figure 2.**
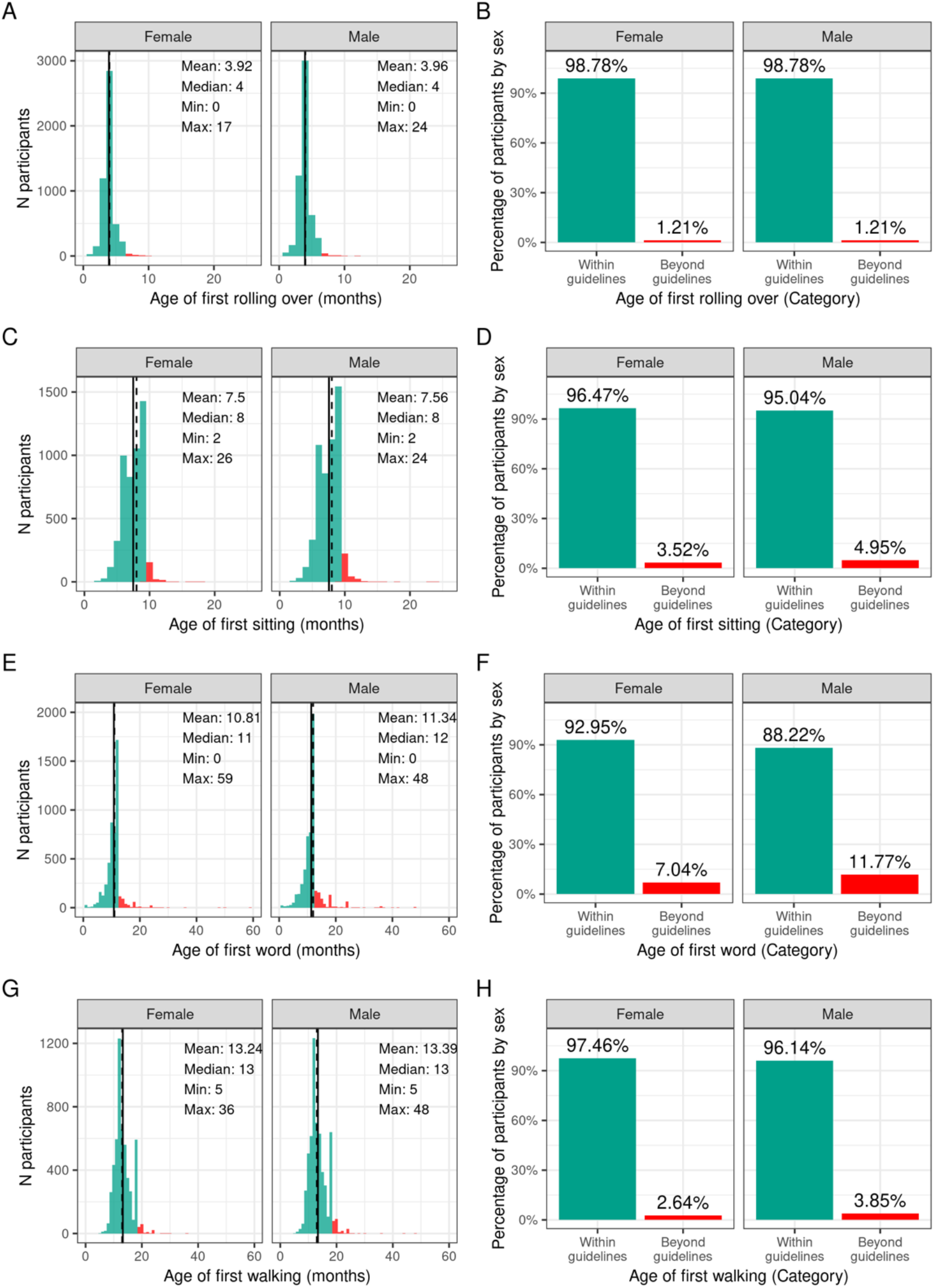
Distribution of age at reaching developmental milestones in the ABCD sample. Distribution of ages in months when each participant reached each developmental milestone stratified by sex assigned at birth, and percentage of those within the recommended developmental guidelines (green) and beyond the guidelines (red) for when each participant first (A,B) rolled over, (C,D) sat on their own, (E,F) said their first word, (G,H) started walking.

Supplementary Figures 3-6 demonstrate the probability distribution of the developmental milestones (age of rolling over, sitting without assistance, walking without assistance, and speaking their first word) for all levels of the other sociodemographic factors (race, ethnicity, household income, and parent educational status). Supplementary Tables 23-26 report the weighted frequencies and percentages stratified by sex of baseline participants meeting guidelines for the age of rolling over, sitting without assistance, walking without assistance, and speaking his/her first word for all levels of the sociodemographic factors. Sex was significantly associated with age of first word spoken (χ^2^(1, 8902)=50.4, *p*=1.26e-12), such that females were more likely to meet guidelines. Sociodemographic factors were not significantly associated with motor developmental milestones (rolling over, sitting without assistance, or walking without assistance) (Supplementary Tables 43-46).

#### Medical problems during birth and pregnancy

Figure 3 illustrates the frequency distribution of males and females in the baseline sample based on the mother having medical problems during birth and pregnancy. Most participants were born to mothers who had no problems during pregnancy (58.6% of female infants and 60.6% of male infants); 25.6% of female infants and 24.6% of male infants were born to mothers who had one medical problem during pregnancy, and 15.7% of female infants and 14.7% of male infants were born to mothers who had two or more medical problems during pregnancy. The majority of participants were born to mothers who had no medical problems during birth (77.4% of female infants and 73.9% of male infants), 16.5% of female infants and 18.7% of male infants having one medical problem during birth, and 6.1% of female infants and 7.4% of male infants were born to mothers who had two or more medical problems during birth.

**Figure 3.**
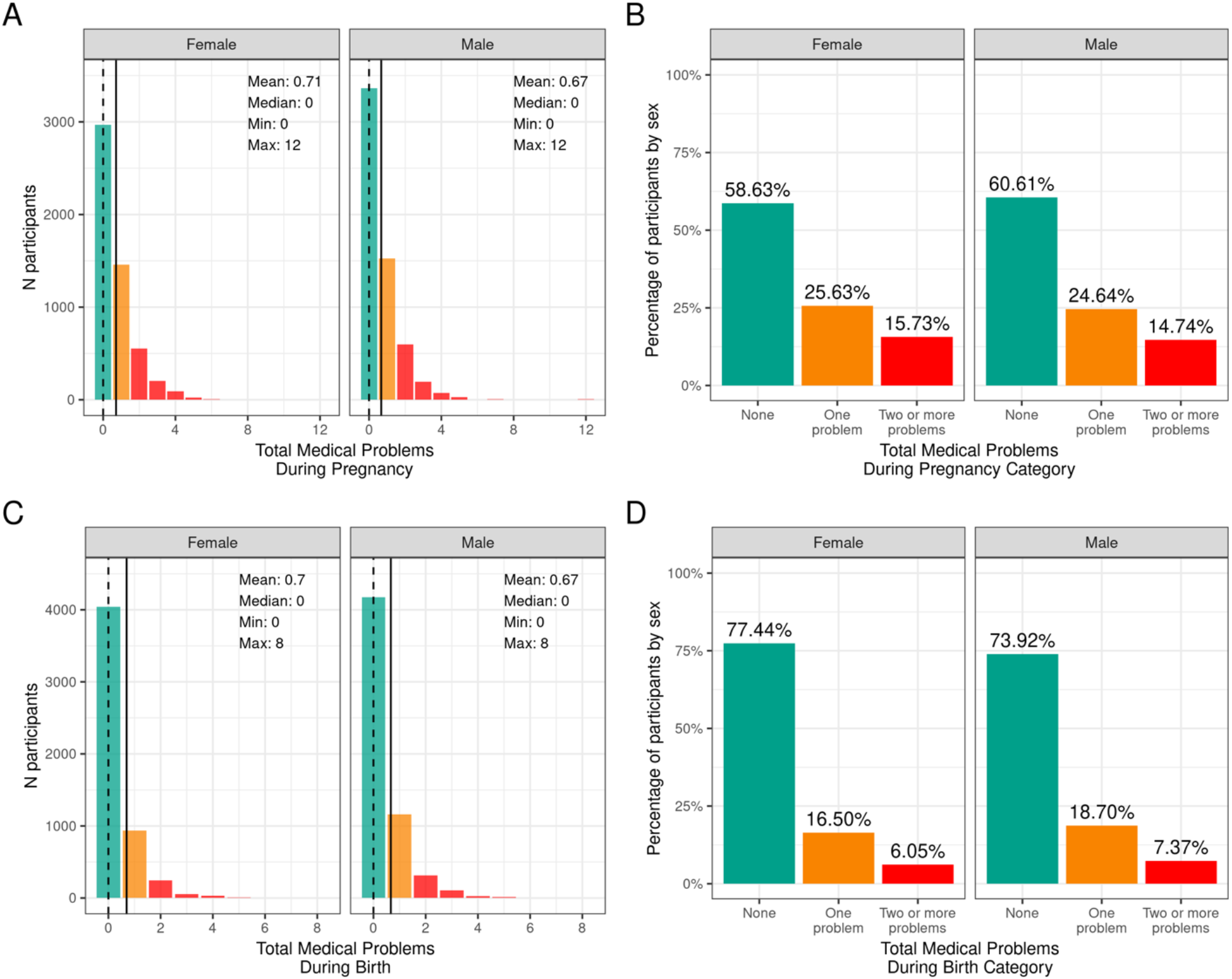
Distribution of participants who experienced medical problems during birth and pregnancy. Continuous distributions of the total number of medical problems experienced by each participant during pregnancy (A) and during birth (C) stratified by sex assigned at birth and color coded by the number of problems experienced: none (green); one problem (orange); two or more problems (red). Percentage of participants (stratified by sex assigned at birth) who fell into these categories for problems experienced during pregnancy (B) and during birth (D).

Supplementary Figures 7 and 8 demonstrate the probability distribution of total medical problems during pregnancy and during childbirth, respectively, for all levels of the other sociodemographic factors (race, ethnicity, household income, and parent educational status). Supplementary Table 27 shows the distribution and weighted frequencies of participants across each of the three categories of problems during pregnancy (no problem, one problem, two or more problems) stratified by demographics (sex, race, Hispanic status, household income level and education level). Chi-squared tests (Supplementary Table 47) showed significant associations between total pregnancy problems and sociodemographic factors of race (χ^2^(6,8902)=68.4, p=8.55e- 13), ethnicity (χ^2^(2,8902)=22.7, p=1.17e-05), household income (χ^2^(4,8902)=115, p=7.64e-24) and parental education (χ^2^(8,8902)=155, p=1.46e-29). Black and Other/Mixed Race infants were less likely to be born to mothers who had no pregnancy problems. Similarly, Hispanic infants were less likely to be born to mothers who had no pregnancy problems. In addition, infants in low income households [<50K] were less likely to be born to mothers who had no pregnancy problems. Similarly, infants whose mothers had a HS diploma/GED or some college education were more likely to be born to mothers who had two or more problems during pregnancy, and infants whose mothers had a postgraduate degree were more likely to be born to mothers who had no pregnancy problems.

Supplementary Table 28 shows the distribution and weighted frequencies of participants across each of the three categories of problems during birth (no problem, one problem, two or more problems) stratified by demographics (sex, race, Hispanic status, household income level and education level). For total birth problems (Supplementary Table 48), chi-squared tests showed a significant association with parental education (χ^2^(8,8902)=37.9, p=7.77e-06). No other sociodemographic factors were significantly associated with total birth problems.

#### Prenatal Substance Exposure

Figure 4 shows the percentage of caregivers who endorsed maternal use of substances of abuse during pregnancy for pre- and postpregnancy recognition, plotted for females and males separately. Differences in maternal reports of substance use during pregnancy did not differ between female versus male children (all *p*-values > 0.30). Among the substances used prior to pregnancy recognition and when a woman was likely pregnant, alcohol use was the most endorsed substance (~23%), followed by tobacco (~8%), then cannabis (aka marijuana) (~3.5%); all other substances with abuse potential, ~1%. Endorsement of substance use declined considerably after pregnancy recognition, with tobacco use having the highest endorsement (~5%), followed by alcohol (~2.5%), then cannabis (aka marijuana (~2%)), and then all other substances with abuse potential (~0.6%).

**Figure 4.**
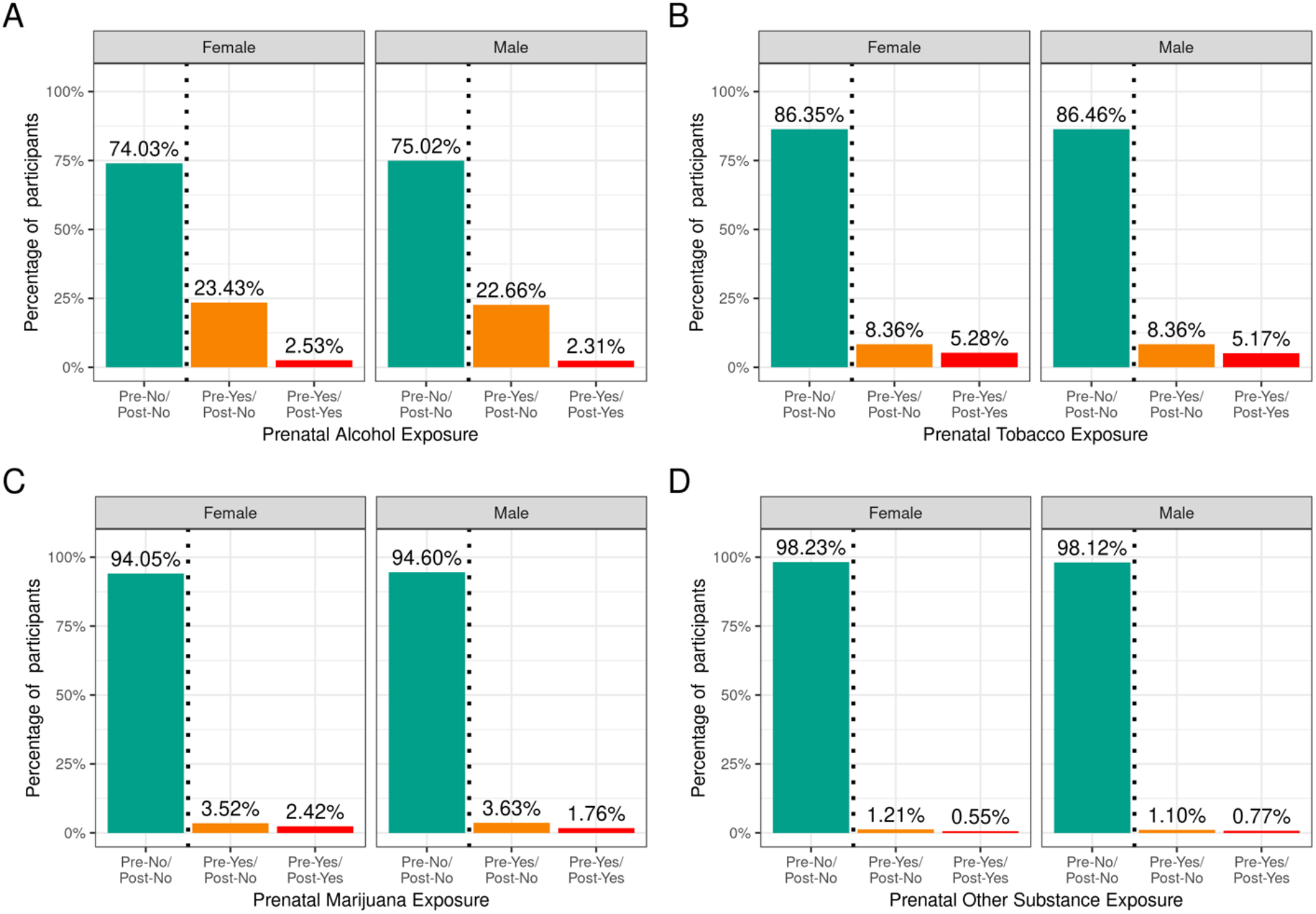
Distribution of participants who experienced prenatal substance exposure. Percentage of participants potentially exposed to alcohol (A), tobacco (B), marijuana (C) or other substances (D) prenatally stratified by sex assigned at birth and grouped by reported use pre and post pregnancy recognition: no use pre- or post-pregnancy recognition (green); use pre-, but not post-pregnancy recognition (orange); use pre- and post-pregnancy recognition (red).

Supplementary Figures 9-12 demonstrate the probability distribution of prenatal alcohol, tobacco, marijuana, and other substance exposure, respectively, for all levels of the other sociodemographic factors. Supplementary Tables 29-32 show the distribution and weighted frequencies of participants across each level of endorsement of prenatal substance exposure stratified by demographics (sex, race, Hispanic status, household income level and education level). The distributions for prenatal alcohol differed significantly by ethnicity (χ^2^(2,8308)=35.8, p=1.66e-08) and household income (χ^2^(4,8308)=196, p=2.6e-41) (see Supplementary Tables 49-52). For tobacco endorsement, distribution significantly differed by household income (χ^2^(4,8641)=456, p=2.51e-97), parental education (χ^2^(8,8641)=636, p=4.5e-132), and ethnicity (χ^2^(2,8641)=16, p=0.00033. The distribution for prenatal marijuana and other substances with potential for abuse significantly differed by household income (marijuana: (χ^2^(4,8599)=226, p=7.55e-48; other substances: (χ^2^(4,8885)=31.9, p=1.97e-06)). The general pattern of prenatal substance use is such that the majority endorsed no substance use during pregnancy, then a significantly reduced number endorsed use pre- and not post- pregnancy recognition, and finally, a significantly smaller percentage endorsing continued use post-pregnancy recognition. A very small percentage of mothers endorsed no use pre- but use post- pregnancy recognition; this sub-group was removed from analyses.

### Current Measures

#### Sleep

Figure 5 shows the distribution of the sample separately for male and female children according to categories of hours of sleep per night and total sleep-wake disturbance scores based on caregiver report. The majority reported between 9-11 hours of sleep per night, which aligns with the recommended sleep duration for children aged 6-12 years old [28]. However, ~12% of participants had between 7-8 hours, and just over 3% had less than 7 hours of sleep per night. There was a wide range in total sleep-wake disturbance scores, with 30% of the sample scoring greater than the cutoff of 39 reflecting possible sleep disturbance [16]. Supplementary Figures 13 and 14 demonstrate the probability distribution of average hours of sleep per night and sleep disturbance score, respectively, for all levels of the other sociodemographic factors.

**Figure 5.**
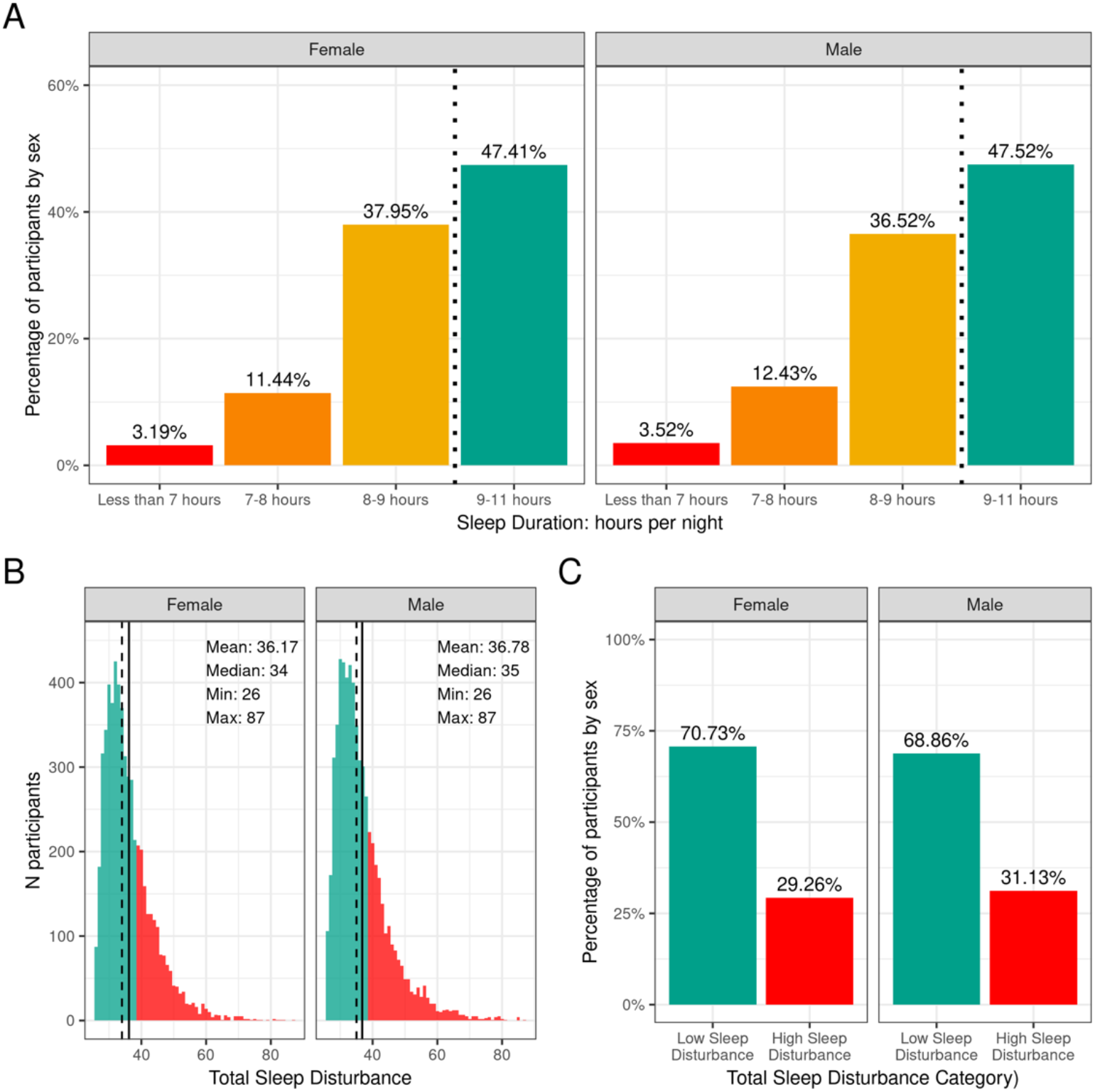
Distribution of participants sleep duration and disturbances. A) Percentage of participants stratified by sex assigned at birth who indicated their average sleep duration to be less than 7 hours (red), 7-8 hours (orange), 8-9 hours (yellow) or 9-11 hours (green). B) Continuous distribution of participant’s scores on the Sleep Disturbance Scale for Children (SDSC) stratified by sex assigned at birth. Participants scoring <39 were deemed to have low sleep-wake disturbance (green) and those scoring >39 wered deemed to have high sleep-wake disturbance (red). C) Percentage of participants (stratified by sex assigned at birth) who experienced low (green) or high sleep-wake disturbance (red).

Supplementary Tables 33-34 show the distribution and weighted frequencies of participants across each level of sleep duration and sleep disturbance stratified by demographics (age, sex, race, Hispanic status, household income level and education level) and pubertal status. There were significant associations between sleep duration and age (χ^2^(9,8593)=72.8,p=4.36-12), such that children of older age were less likely to have 9-11 hours of sleep and those of younger age were more likely to have 9-11 hours (Supplementary Table 53). Sleep duration did not differ according to sex. The distribution was significantly dependent on ethnicity (χ^2^(3,8593)=130,p=4.84e-28), household income (χ^2^(6,8593)=834,p=6.98e-177), and parental education (χ^2^(12,8593)=891,p=5.37e-183). Children who identified as Hispanic were less likely to have more than 9 hours of sleep. Also, children in families of higher income and higher education were more likely to have longer sleep duration, whereas children in families of lower income and lower education were more likely to have <7 hours of sleep.

Age and sex were not associated with having high versus low sleep-wake disturbance (Supplementary Table 54). There was an association of sleep disturbance and race (χ^2^(3,8593)=21.1, p=0.0001), household income (χ^2^(2,8593)=73.8, p=9.3e-17), and parental education (χ^2^(4,8593)=52.2, p=1.3e-10). Children in families with higher incomes and more education were less likely to have high sleep-wake disturbance, whereas children in families with lower incomes were more likely to have high sleep-wake disturbance. In addition, pubertal development was associated with sleep disturbance (χ^2^(3,8593)=3.5,p=2.5e-07), with prepubertal children being more likely to have low sleep-wake disturbance.

#### Physical Activity and Sports Involvement

Figure 6 shows the number of participants engaging in vigorous physical activity for 60 minutes and strengthening exercises as a function of number of days per week stratified by sex. The CDC recommends at least 60 minutes of moderate to vigorous intensity physical activity daily for 5-17 year old children. Relatively few children met these guidelines (17.8% of boys and 15% of girls). The CDC also recommends including muscle-strengthening activities, such as sit-ups or push-ups, at least 3 days per week for 5–17-year-olds; 33.3 % of boys and 28.9 % of girls met these guidelines. Supplementary Figures 15 and 16 illustrate the probability distribution of number of days of 60 minutes of physical activity and number of days engaged in strengthening exercises (past week) as a function of the other sociodemographic factors and pubertal status.

**Figure 6.**
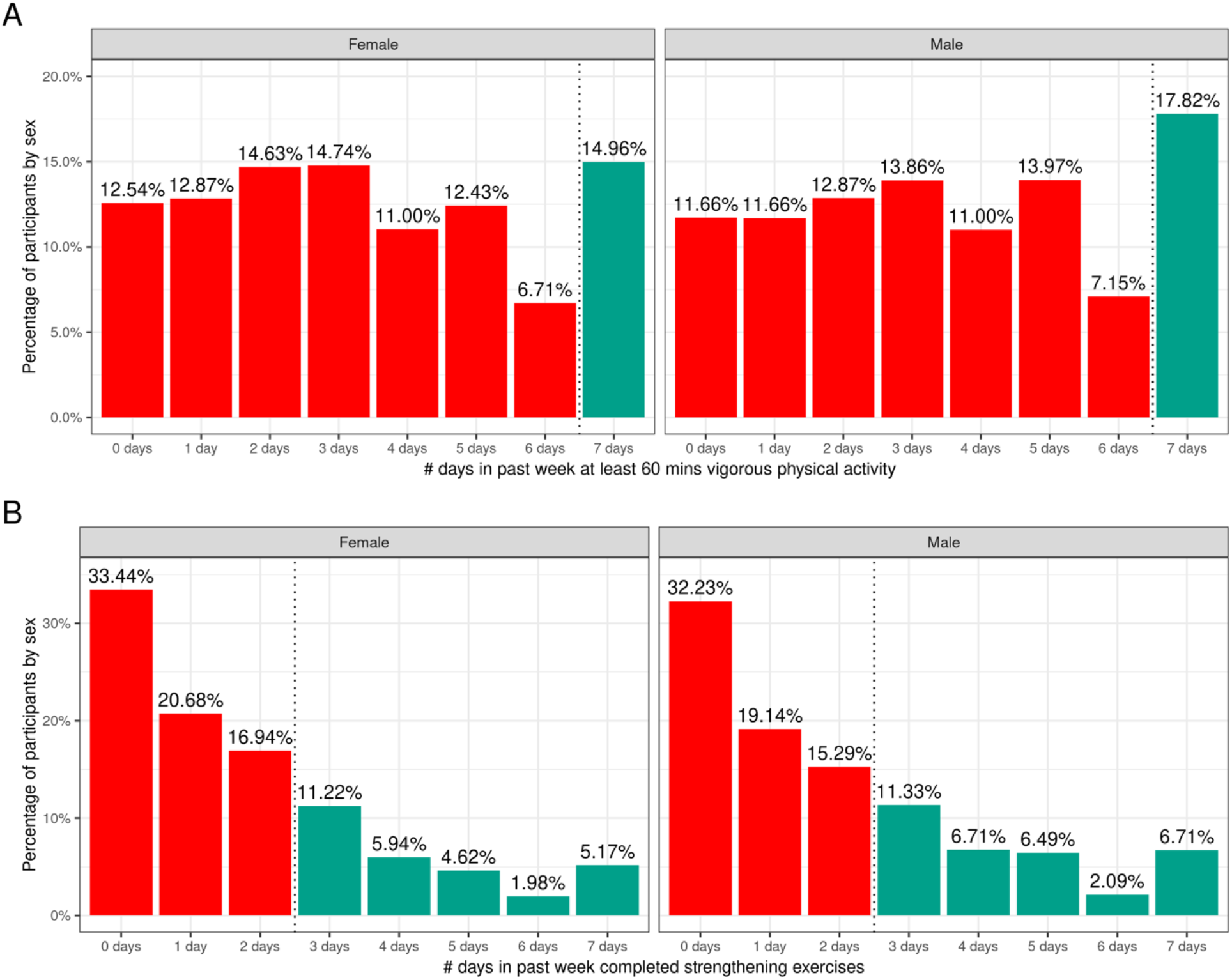
Distribution of time spent participating in physical activity across the ABCD sample. A) Distribution of number of days in the past week participants (stratified by sex assigned at birth) spent participating in at least 60 minutes of vigorous physical activity. B) Distribution of number of days in the past week participants (stratified by sex assigned at birth) completed strengthening exercises. Participants that met CDC guidelines for recommended duration of physical activity (green) and that did not meet guidelines (red).

Supplementary Tables 35 and 36 show the distribution of participants and weighted frequencies who were above and below the CDC guidelines for the number of days the child engaged in vigorous physical activity (PA1) and did strengthening exercises (PA2) as a function of sociodemographic factors. For PA1, the distributions significantly differed by sex (χ^2^(1, 8593)=14.3, p=0.000159) and ethnicity (χ^2^(1,8593)=13.6, p=0.00022) (Supplementary Tables 55 and 56). The association with pubertal development showed a trend towards significance (χ^2^(3,8593)=17.7, p=0.0005) such that those in pre-pubertal stages were more likely to meet guidelines and those in the middle pubertal stage were less likely to meet guidelines. Male participants were significantly more likely to be above the CDC guidelines, and female participants were significantly more likely to be below the CDC guidelines.

For PA2, the distributions significantly differed by sex (χ^2^(1,8593)=19.7, p=9.06e-06), race (χ^2^(3,8593)=75.2, p=3.27e-16), household income (χ^2^(2,8593)=31.3, p=1.6e-07) and parent educational attainment (χ^2^(4,8593)=36, p=3e-06) (Supplementary Tables 55 and 56). Male participants were significantly more likely to be above the CDC guidelines, and female participants were significantly more likely to be below the CDC guidelines. Participants from households with higher income and parental education were less likely to meet CDC guidelines specifically for strengthening exercises, and Black participants were significantly more likely to be within CDC guidelines for strengthening exercises.

Figure 7 shows the distribution of average hours per week spent participating in sports over the past 12 months stratified by sex. Despite the heavy rightward skew of the sports involvement summary score, 73.48% of girls and 77.44% of boys reported engaging in some form of sports participation in the past year with a high degree of variability among the sample. This is a similar proportion of the sample to those reporting participating in some form of physical activity (>0 days for PA1 and PA2), with boys similarly reporting more sports participation compared to girls (χ^2^(1,8593)=14, p=0.00018). The time spent participating in sports was relatively low as many sports are seasonal, so the average time spent participating over the entire year is greatly reduced. It is, however, important to note that participants could only endorse participation in the sports listed in the SAI-Q; although the list is extensive, this measure may underestimate sports participation.

**Figure 7.**
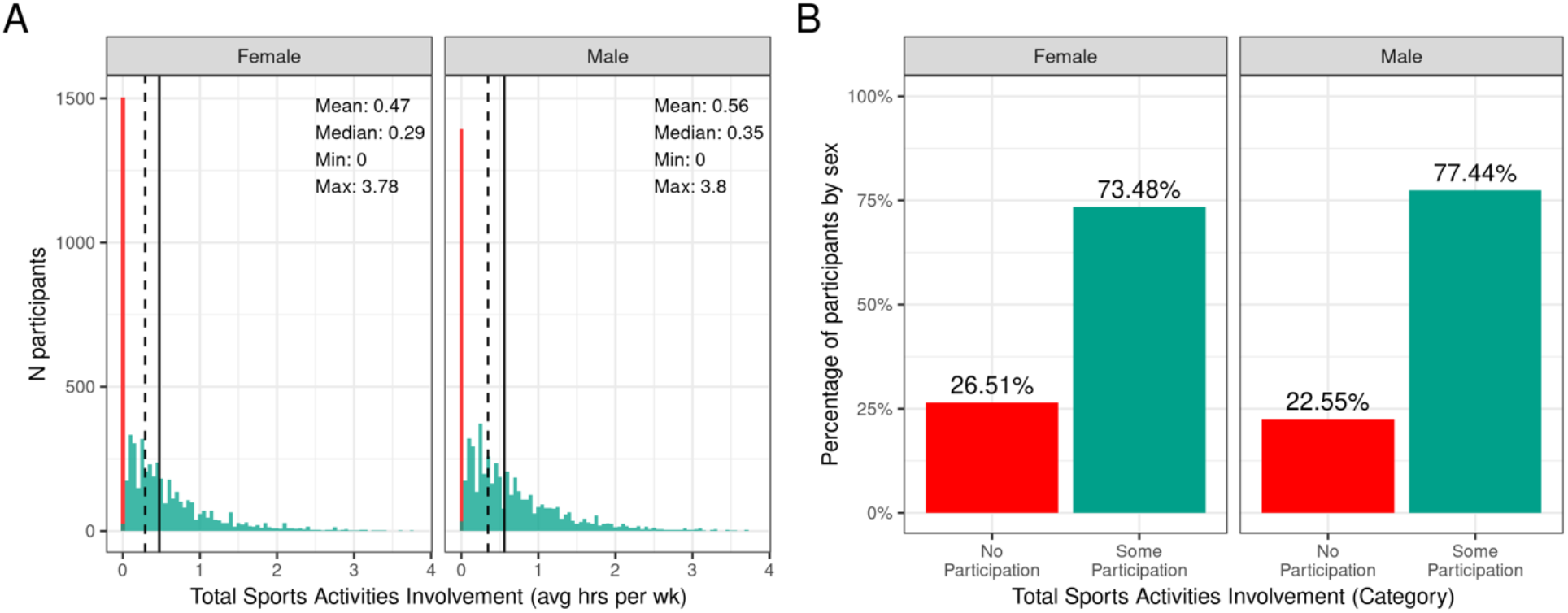
Distribution of time spent participating in sports across the ABCD sample. A) Continuous distribution of the average hours per week each participant (stratified by sex assigned at birth) participated in sports (total across all sports endorsed) in the past year as indicated by the Sports Activites Involvement (SAI) questionnaire. B) Percentage of participants (stratified by sex assigned at birth) who indicated no participation in sports (red) and some participation in sports (green).

Supplementary Figure 17 illustrates the probability distribution of sports participation for all levels of the sociodemographic factors and pubertal status. Supplementary Table 37 show the distribution of participants and weighted frequencies who endorsed some and no participation as a function of sociodemographic factors. Chi-squared tests (Supplementary Table 57) also showed large associations between sports involvement and sociodemographic factors of race (χ^2^(3,8593)=281, p=1.06e-60), ethnicity (χ^2^(1,8593)=81.8, p=1.49e-19), household income (χ^2^(2,8593)=850, p=2.11e-185), and parental education (χ^2^(4,8593)=870, p=4.37e-187), in that families with a higher household income and greater parental educational attainment were more likely to endorse participation. Participants further along in pubertal development (midstage) were less likely to endorse participation (χ^2^(3,8593)=850, p=2.11e-185). This may be driven by females showing greater pubertal development at this age.

#### Body Mass Index (BMI) and Weight Status

The distribution of males’ and females BMI percentiles is shown in Figure 8. Across the whole sample, 3.4% of males and 4.1%of females had underweight (BMI< 5^th^ %ile). Most of the participants (64%) had a healthy-weight (BMI < 85^th^ %ile and ≥5^th^ %ile). Overweight was present in 15% of the sample while an additional 17.4% of males and 16.1% of females had obesity. Supplementary Figure 18 shows the probability distribution of BMI percentiles for all levels of the sociodemographic factors and pubertal status. Supplementary Table 38 shows the distribution of participants and weighted frequencies across weight categories as a function of sociodemographic factors.

**Figure 8.**
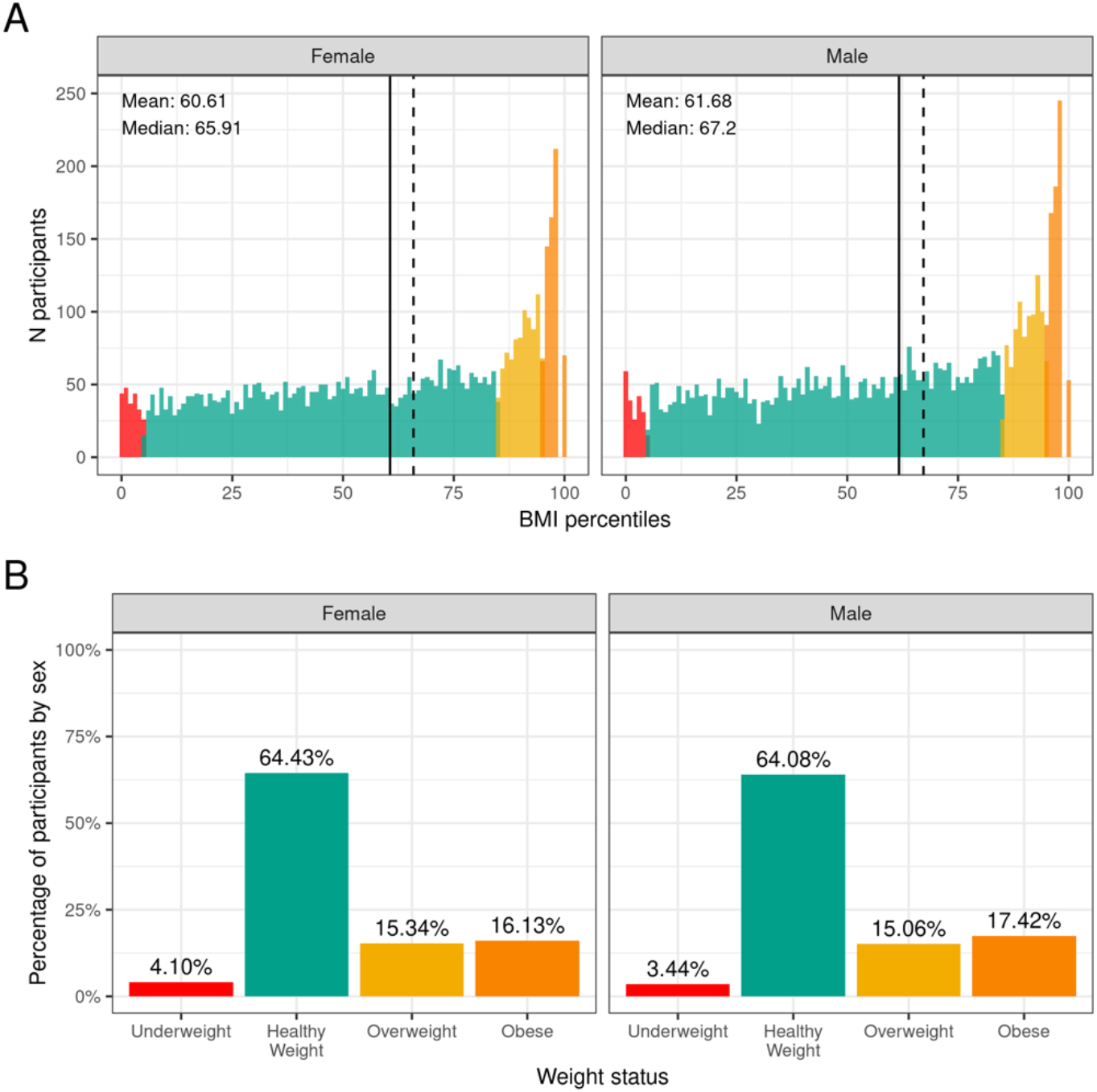
Distribution of BMI and weight status in the ABCD sample. A) Continuous distribution of BMI percentiles stratified by sex assigned at birth color coded based on weight status: underweight (red), healthy weight (green), overweight (yellow) and obesity (orange). B) Percentage of participants (stratified by sex assigned at birth) within each weight status.

Since CDC percentiles are adjusted for age and sex, weight status distributions were not significantly different across sex (χ^2^(3,8568)=5.54, p=0.136) or age (χ^2^(9,8568)=5.27, p= 0.81) (Supplementary Table 58). Conversely, distributions were significantly different across different racial groups (χ^2^(9,8568)=289, p=4.5e-57), ethnicity (χ^2^(3,8568)=202, p=0.1.2e-43), and household income (χ^2^(6,8568)=417, p=5e-87). The association with parental education (χ^2^(8568)=529, p=5e-04) and pubertal status (χ^2^(8568)=383, p=5e-04) showed a trend towards significance. Pre-pubertal children were less likely to have overweight/obese, whereas mid to late/post puberty corresponded with a greater likelihood of having overweight/obesity.

#### Medical Problems (Lifetime)

Figure 9 illustrates the frequency distribution of males and females in the baseline sample for caregiver-reported history of lifetime medical problems for which their child saw a doctor. Across the sample, 32.4% of females and 29.4% of males had no medical problems, 30.7% of females and 29.4% of males had to see a doctor for one medical problem, and 36.8% of females and 41.2% of males had to see a doctor for more than one medical problem according to caregiver report. The most common causes for seeing a doctor for both males and females included allergies, asthma, vision problems, and operations. Participants that reported more than 6 medical problems in their lifetime were excluded (n=47).

**Figure 9.**
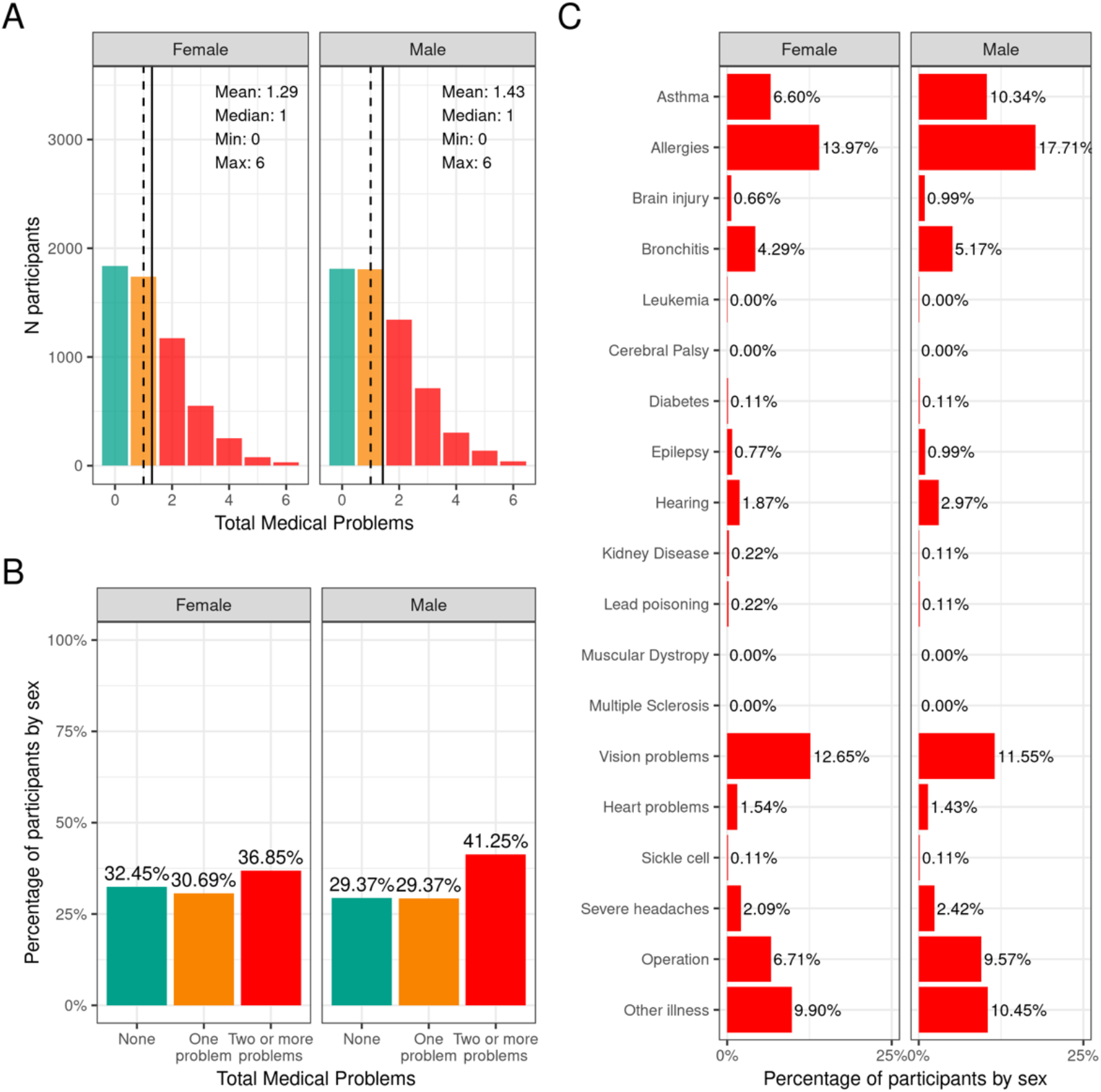
Distribution of participants who experience lifetime medical problems. A) Continuous distribution of the total number of medical problems experienced by each participant within their lifetime stratified by sex assigned at birth and color coded by the number of problems experienced: none (green); one problem (orange); two or more problems (red). B) Percentage of participants (stratified by sex assigned at birth) who fell into these categories for problems experienced during their lifetime. C) Percentage of participants endorsing each medical problem.

Supplementary Figure 19 shows the probability distribution of participants who endorsed none, one, two or more problems for all levels of the sociodemographic factors and pubertal status. Supplementary Table 39 shows the distribution of participants and weighted frequencies who endorsed none, one, and two or more problems as a function of sociodemographic factors. Chi-squared tests (Supplementary Table 59) showed associations between number of medical problems endorsed and sociodemographic factors of ethnicity (χ^2^(2,8561)=17.2, p=0.00018), household income (χ^2^(4,8561)=27.7, p=1.46e-05), and parental education (χ^2^(8,8561)=67.9, p=1.3e-11). Participants from lower income households and with parents from lower educational level were more likely to endorse no medical problems.

#### Traumatic Brain Injury (lifetime)

Figure 10 demonstrates the number of participants in each TBI diagnostic category at baseline stratified by sex. Most participants (95.5% of boys (n=5909) and 96.1% (n=5502) of girls) had improbable TBI (no head injury or head injury without loss of consciousness or alteration in memory). 3.2% of the caregivers of male participants (n=199) and 2.2% of the caregiver of female participants (n=123) reported a head injury consistent with a possible mild TBI (head injury with alteration in memory but no loss of consciousness). Finally, a small proportion of the caregivers (1.2% of boys (n=75) and 0.9% of girls (n=52)) reported a head injury consistent with a mild TBI (head injury with a loss of consciousness of less than 30 minutes). There were less than 10 caregivers who reported head injury consistent with a diagnosis of moderate or severe TBI. Supplementary Figure 20 demonstrates the probability distribution of worst lifetime head injury for all levels of the other sociodemographic factors and pubertal status.

**Figure 10.**
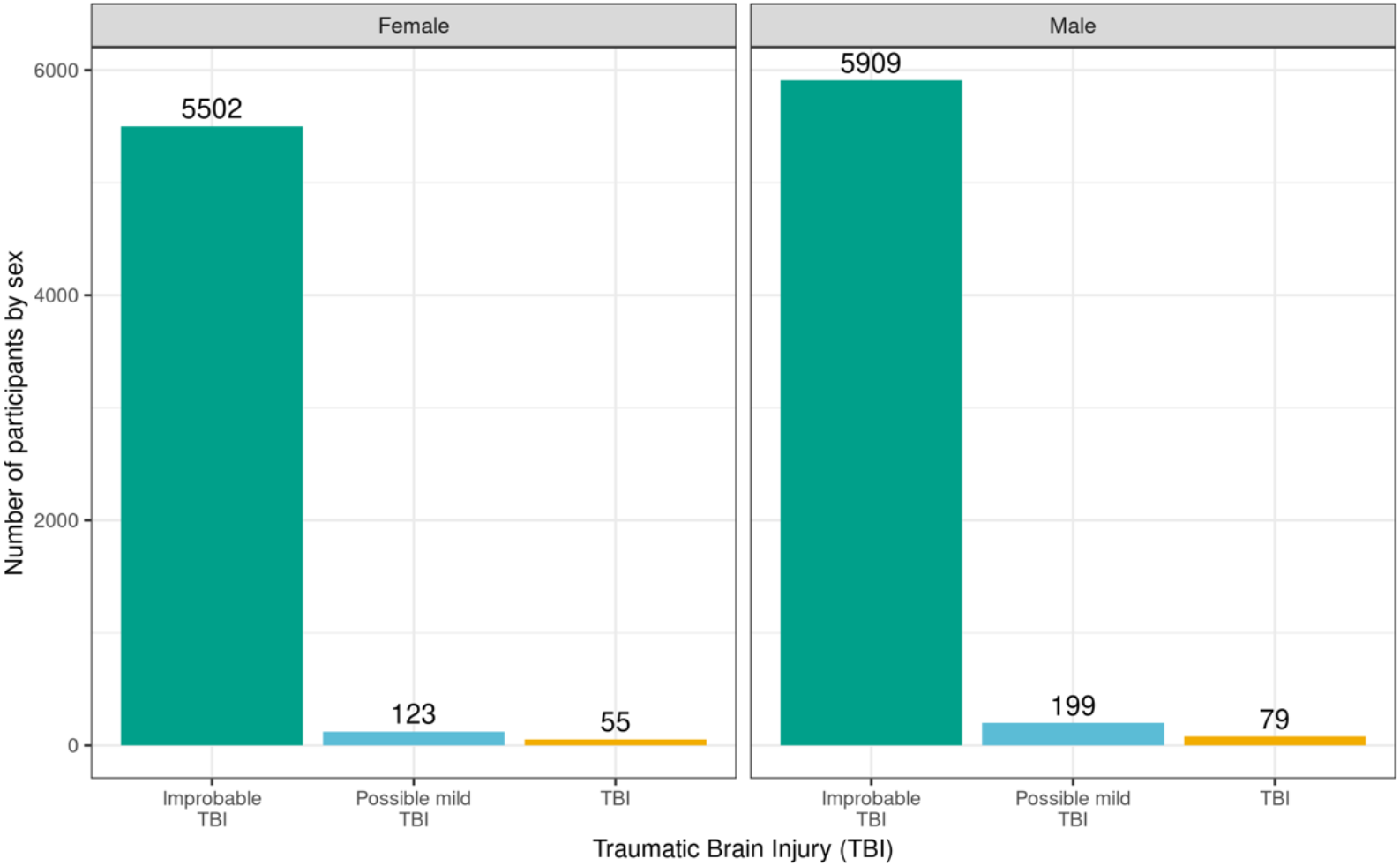
Distribution of participants who experienced a traumatic brain injury (TBI). Percentage of participants (stratified by sex assigned at birth) who experienced improbable TBI (green), possible mild TBI (blue), or experienced a TBI (collapsed across mild, moderate and severe TBI; yellow).

Supplementary Table 40 shows the distribution of participants and weighted frequencies across TBI categories as a function of sociodemographic factors. The proportion of subjects for different TBI diagnosis was not significantly associated with any of the sociodemographic factors (Supplementary Table 60).

## Discussion

We have provided a comprehensive overview of the physical health measures collected in the ABCD Study and the distributions of these measures at baseline (9- to 10-years-old), as well as how these varied across different sociodemographic factors. In addition, we have highlighted where participants did not meet standard guidelines for this developmental stage based on available population data and CDC recommendations, which has allowed us to examine how the ABCD Study sample compares to current national standards.

In this comprehensive study, we identified health disparities across many of the physical health domains, including medical problems during pregnancy and birth, weight status, physical activity, and sleep, which require further clarification and contextualized analyses. The presence of known lifetime youth medical problems was more prevalent in higher income and educated families, but this result, based on a self-report measure, is likely biased by access to healthcare, knowledge of diagnoses, and structural racism/biases within healthcare. Therefore, it is likely that medical problems are under-reported in low-income families in ABCD, highlighting the need for objective markers of physical health, some of which will become available with later releases of the ABCD data. The interpretation of BMI data can be challenging as weight status defined by BMI percentile for age and sex is also recognized as a screening tool, and is not a precise measure for metabolic health since it does not always correlate with percent body fat. Differences seen across race, ethnicity, and parent education are commonly noted in the literature and were also identified in the ABCD cohort [29]. Whether these differences relate to greater metabolic problems in these demographic groups remain to be seen in future ABCD measurements. Finally, sleep disturbances were present in a large proportion of youth, in line with previous observations. Sleep duration and disturbance differed with age, puberty, and across all sociodemographic groups, underscoring the importance of sleep health for development and health disparities research.

In many domains the distribution of physical health characteristics in the ABCD sample were similar to published national normative data [30–32]. Birth weight was an exception: given the high proportion of twins in the ABCD sample, there was a greater proportion of youth born with low birth weight. The ABCD sample also has a lower proportion of youth who did not meet guidelines for developmental milestones; however, this is unsurprising given the inclusion requirements to participate in the study. With respect to physical activity, the majority of the cohort were insufficiently active according to recommended CDC guidelines. This differed by assigned sex at birth, with females less likely to meet guidelines than males in line with previous findings. Below we discuss each of the domains measured in more detail.

### Developmental Measures

#### Birth weight and prematurity

According to data reported by the CDC in 2018, 8.3% of infants were born with a low birthweight and 1.3% of infants were born with a very low birthweight. Results from the ABCD study show that approximately 14.8% of the ACS-weighted sample was born with a low birthweight, which is higher than the national standard. This likely reflects the large twin cohort within the ABCD sample, which must be taken into account when analyzing associations with birthweight and other factors. Rates of very low birthweight and extremely low birthweight should be interpreted with caution since a birthweight of less than 1200g was an exclusionary criterion for the ABCD Study. The sex difference in birthweight, with males being more likely to be in the high birthweight category and females more likely to be in the low birthweight category, has been reported previously [33, 34]. The results from the chi-square analyses showed that participants in the low-birthweight category were less likely to identify as White and those in the high-birthweight category were less likely to identify as Black. This disparity has been identified previously [35] and recent CDC data have shown that the low birthweight rate was more than twice as high for non-Hispanic black infants as for non-Hispanic white infants (11.36% compared with 5.21% in 2016). Further, participants who identified as Hispanic in the ABCD sample were less likely to be in the low-birthweight category [36, 37].

In 2018, the CDC reported that ~10% of births in the United States were preterm (percentage of all births delivered at less than 37 completed weeks of gestation). The current study shows similar rates, with ~12% reporting preterm births. Being born before 28 weeks of gestation (i.e., more than 12 weeks premature; extreme preterm birth) was an exclusionary criterion for ABCD, so the rates of extreme preterm birth reported here may be an underestimate. The proportion of participants in each category based on prematurity depended on household income, such that participants from lower income households (<$50K) were less likely to be in the late preterm category and more likely to be in the extreme preterm category. These findings should be interpreted with caution since the percentage of the sample in the extreme preterm category was very small. Prior studies have shown that preterm birth rates were significantly higher among the women in the lowest (versus the highest) family income group [38] may reflect limited access to prenatal healthcare in the US among poorer individuals, effects of increased stress of poverty and/or structural biases.

#### Developmental Milestones

The majority of ABCD participants met CDC guidelines for the age of achieving developmental milestones. Children who do not meet standard guidelines for developmental milestones are considered to have developmental delay. The percentage of ABCD participants that did not meet the standard guidelines (1.21-11.7%) was lower than national estimates of developmental delays (13-15%) [31, 39]. One reason for this difference may be that children with developmental delays may not have met eligibility criteria for the study, thus excluding from enrollment. Further, retrospective report by caregivers may be subject to recall bias, contributing to the smaller percentage of children in the study that exceeded standard guidelines.

Females in the study were more likely to meet standard guidelines for first word speech milestones. Sex differences in language acquisition have been consistently reported, with females outpacing males as early as 6 months of age on tasks related to sensory discrimination of speech sounds [40]. Recent studies have proposed neurobiological mechanisms to explain sex differences found in speech and language development. Specifically, sex hormones such as testosterone have shown to influence language function and lateralization during the first month of life [41], whereas sex differences have been reported in the development of neural and temporal processes involved in language acquisition [40].

#### Medical problems during birth and pregnancy

The proportion of participants reporting one or more medical problems during pregnancy (41.3% of female infants; 39.3% of male infants) or birth (22.6% of female babies; 26.1% of male infants) were higher than reported general trends of pregnancy and birth complications among US mothers. A recent report on 1.8 million pregnancies between 2014 and 2018 among commercially insured women (ages 18 to 44) showed that pregnancy complications increased from 16.8% in 2014 to 19.6% in 2018 and that birth complications increased from 14.8 to 16.9% during the same time period (Blue Cross Blue Shield, 2020). However, this study was biased by those with commercial insurance highlighting the need for more representative samples. In our study, infants of minority racial and ethnic backgrounds and households of lower socioeconomic status were less likely to be born to mothers with no medical problems during pregnancy, which is in line with research examining racial and ethnic disparities in pregnancy risks and complications in the US [42]. Along the same lines, studies show low socioeconomic status being associated with higher risk of pregnancy complications such as preeclampsia, diabetes, and preterm delivery [43, 44]. However, unexpectedly, birth complications in our sample were positively associated with parental education, such that caregivers of participants from families with lower parental education were more likely to report *no* medical problems at birth. This may be confounded by mother’s age at giving birth, employment status, occupational exposures, and whether the mother was informed that they had any medical problems.

#### Prenatal substance exposure

Alcohol was the most endorsed substance for potential of unintended exposure early on in pregnancy. Endorsement of prenatal substance use declined after pregnancy recognition for all substances with tobacco then becoming the most endorsed substance (~5%). The distribution of maternal prenatal substance use during pregnancy is similar to patterns of use in women who were pregnant in a discrete time period of 2006-2009 [45]. Marijuana was widely legalized throughout the US after 2009, but patterns of prenatal marijuana use in this cohort are similar to preceding data. Retrospective endorsement of prenatal substance use 9 to 10 years earlier has clear limitations due to recall, particularly when there are multiple siblings, and possibility of revealing stigmatizing information. The percentages of caregivers endorsing a specific substance used in pregnancy significantly differed as a function of sociodemographic factors, suggesting that controlling for these factors or matching samples on these factors may be needed in future analyses on teratogenic effects.

Most human studies on the impact of prenatal exposure to substances of abuse on brain and cognitive development either rely on recruitment from clinically identified populations or target offspring of mothers who endorse substance use during pregnancy prior to study enrollment. Thus, the quantity, frequency, and duration of maternal substance use may be higher in those studies than what might be observed in a community sample not specifically targeted for substance use during pregnancy. The ABCD Study is in a unique position to address some limitations, as recruitment was not dependent on maternal endorsement or denial of substance use during pregnancy.

### Current Measures

#### Sleep

The recommended sleep duration for school age children is 9-11 hours [28, 46]. In the ABCD Study, about 85% showed adequate [9-11 hours] or near adequate [8-9 hours] sleep duration while a concerning proportion (~15%) reported inadequate sleep [i.e., 7-8 hours or less], which is somewhat lower than what has been reported previously [47]. Utilizing the recommended overall sleep disturbance symptom score from SDSC [i.e., 39], about 30% showed sufficient symptoms to warrant additional evaluation, similar to findings from the development of the sleep disturbance scale (26% of the control sample had scores consistent with possible sleep disturbance) [16]. While seemingly high, the authors commented that there is a high prevalence of undiagnosed sleep disorders in children (30%) [16]. Here, we report the total score, which could reflect a number of possible sleep problems, including sleep breathing disturbances, night tremors, or insomnia. Follow-up analyses are needed to determine which factors in the scale are most common in this sample.

As expected, and even with the narrow age window examined, older children and those with more advanced pubertal stages were less likely to have sufficient sleep. Pubertal advancement was also associated with more sleep disorder symptoms. Sleep duration declines as children progress across adolescence, due to a combination of biological and psychosocial changes [48–50] however, for the young age group studied here, with already lower than recommended sleep duration, it will be important to track whether their sleep duration shortens even further as they age. Sociodemographic variables were associated with sleep duration and sleep disturbance, which supports prior research [47, 51, 52]. As reported for a large sample of Canadian school-age children [53], we found no sex difference in sleep duration.

While the baseline ABCD sleep data have already been utilized to examine several important issues with mental health and brain development [54–58], it is important to note that these data are limited by caregiver-report, which may not accurately reflect the children’s sleep, the absence of objective sleep indicators, and a lack of differentiation between weekdays and weekends for sleep duration. Future assessments of sleep in the ABCD cohort at follow-up years will address some of these limitations.

#### Physical Activity & Sport Involvement

At baseline, only 17.8% of boys and 15% of girls met CDC guidelines by engaging in daily moderate to vigorous physical activity for 60 minutes. The National Survey of Children’s Health (NSCH) in 2016 showed that less than one-quarter (24%) of children 6 to 17 years of age participated in 60 minutes of physical activity every day. Furthermore, 33.3% of boys and 28.9% of girls endorsed engaging in muscle-strengthening activities, such as sit-ups, push-ups and weight-lifting at least 3 days in the week. In 2017, 51.1% of high school students participated in muscle strengthening exercises (e.g., push-ups, sit-ups, weight lifting) on 3 or more days during the previous week according to CDC data. Although the lower percentage of participants meeting CDC guidelines could be attributed to the younger and narrower age range of 9- to 10-year-old children recruited by ABCD, it is clear that the majority of the cohort were insufficiently active according to CDC guidelines.

On average, young girls are less physically active than boys. According to CDC data, boys (63.7%) engage in muscle-strengthening activities more regularly than girls (42.7%), and approximately 35% of high-school boys versus 18% of girls report at least 60 minutes of daily physical activity [59]. These sex differences are visible in the current study findings wherein a lower percentage of girls met CDC guidelines for moderate to vigorous physical activity as well as for muscle-strengthening exercises. Previous research points to a number of potential factors underpinning this disparity: a) girls participate less in organized sports; b) girls may receive less social support to engage in physical activity; or c) earlier pubertal maturation in girls [60]. In our sample, boys reported greater participation in organized sport, and sports involvement was additionally negatively associated with pubertal development, which supports the potential influence of these factors on physical activity levels in our sample.

Sports involvement and, to a lesser extent, vigorous, cardiac physical activity was additionally associated with higher income and parental education, which reflects the high cost associated with sports participation. This highlights an important need to increase engagement in sport and accessibility of recreational facilities for low-income communities. However, Black participants and those from households with lower income were more likely to participate in strengthening exercises, such as weight-lifting, and fall within CDC guidelines for this type of physical activity. This suggests there may be differences in the types of activity that children participate in in different communities.

#### Body Mass Index & Weight Status

The prevalence of obesity in the ABCD cohort (16.1% females; 17.7% males) closely resembles the national prevalence for obesity for all youth (18.5%), but in particular, for this age group, as reported by the CDC (16.3% for females; 20.4% for males) [30, 61]. The relationship between weight status and race, ethnicity, and household income are also in line with previously established findings of youth in the United States [30, 62]. As in previous research, we found that Hispanic and Black youth are more likely to be in the overweight and obese categories [30]. However, it should be noted that BMI is a screening tool, not a diagnostic tool, for metabolic disease risk. The best measurement of adiposity is obtained via DEXA scans, but this was not feasible in the ABCD study. ABCD did collect measures of waist circumference, which may be a better predictor of adiposity [63]. Yet, waist circumference is also a screening tool and is prone to measurement error because of the presence of abdominal fat folds. Furthermore, no cut-offs exist for clinical interpretation of disease risk in children. Despite these limitations, we presented our results by utilizing the CDC growth curves to classify children into four commonly used weight categories (i.e., underweight, healthy weight, overweight, obese) to allow for direct comparison to national estimates.

It is also important to recognize that BMI does not take into account body shape, composition [64], or genetic differences in fat metabolism [65]. There are known differences in fat metabolism and body composition by race. For example, although Asians appear to be leaner, they have, on average, greater adiposity and less muscle mass [66–68]. This may explain why Asians appear to develop type 2 diabetes at a lower BMI than Whites [69]. Furthermore, in the current study we found that Black participants were more likely than White participants to be within CDC guidelines for strengthening (but not vigorous) activities, which would differentially increase muscle mass and BMI, potentially contributing to the racial differences in BMI we observed. Importantly, regardless of racial-ethnic status, it is clear that good nutrition, access to health care, and good social and general living conditions are integral contributing factors for optimal growth and development. More research is required to contextualize racial-ethnic differences in BMI found in the ABCD sample.

Similar to previous literature [70], we also found participants with overweight/obesity were more likely to present with advanced pubertal development (although this did not reach corrected significance levels). This may be related to the effect of excess adiposity on the production of sex steroids and hormones involved in the hypothalamic-pituitary axis [71]. Given the majority of participants were pre-pubertal at baseline, we expect this relationship to become stronger as the children get older. However, it is important to acknowledge that PDS, used as a measure of physical maturation in the ABCD Study, does not evaluate pubertal stage directly (as opposed to Tanner stage), and a description of the measured construct is better reported as “perceived pubertal stage” [72].

#### Medical Problems (Lifetime)

The results from the current study show that approximately 67% of the ABCD sample had to visit a doctor or healthcare professional due to one or more medical problems or diagnoses in their lifetime. Asthma, allergies, bronchitis, and vision problems were most commonly endorsed. Although the middle childhood population is generally considered healthy, certain health-promoting or health-risk behaviors may take shape during this time and influence outcomes in adolescence and adulthood. Analyses of the National 2016-2017 Survey of Children’s Health data in 6-11 year old children revealed that allergies were the most prevalent physical condition among children (21%), followed by asthma (9.5%), and frequent headaches or migraines (2.6%) [5], showing close agreement with the rates reported in the ABCD sample. However, we found that youth from families with higher household income and parental education were more likely to endorse having medical problems. This highlights a potential bias, as knowledge of a medical problem may rely on access to healthcare and structural racism or biases within medicine and healthcare infrastructures may lead to under diagnosis among certain groups. Therefore, it is likely that the presence of medical problems in lower income youth will be underestimated in the current sample.

#### Traumatic Brain Injury (lifetime)

The prevalence rate for head injury in the current study was approximately 3.9% for girls and 4.5% for boys. These rates are in agreement with prior large-scale community survey studies demonstrating a range of 2-20% [32, 73]. Community surveys tend to screen for multiple health conditions in very large, representative samples, but such screening may artificially result in higher or less reliable head injury prevalence estimates by not assessing injury-specific information in detail [74]. Though boys had a higher prevalence rate, consistent with previous literature [75], the association with sex did not survive correction for multiple comparisons. A number of factors have been identified that may contribute to the sex difference in epidemiology of TBI such as higher incidence of general injury among younger males, variance between males and females in traditional societal roles and activities, and differences in risk-taking behaviors.

### Associations between Race, Sex, and Physical Health Measures in ABCD

Understanding why broad racial groupings or assigned sex at birth relate to physical health measures is outside the scope of analyses in this paper. The broad categories used to capture race in this study are not exhaustive and often conflated with nationality. It is also important to acknowledge that youth who were not proficient/fluent in English were not included in the ABCD study (though caregivers with English or Spanish proficiency were included), therefore, some of the results may have been confounded by this exclusionary criterion. Furthermore, there is an association between race and environmental factors many stemming from systemic racism in the United States, that contribute to health disparities across minoritized youth. Our analyses do not control for this wide variety of influential factors; therefore, we cannot draw any conclusions regarding potential mechanisms underlying these associations. Moreover, although sex differences in physical health may be in part due to biological pathways associated with sex chromosomes, sex is highly conflated with gender identity and there are many societal differences in the experiences of those who identify as male and female that can shape their physical and mental health. Given the high correlation between assigned sex at birth and gender identity and/or expression, we cannot dissociate the contributions of these different factors in the current analysis; however, the ABCD Study has collected data about gender identity to advance our understanding of how assigned sex at birth and gender identity impact individuals’ physical and mental health in future studies. Finally, there is a high degree of variability in response to experiential factors within each sex, race, and ethnic group such that the outcome of an individual cannot be solely predicted by group trends. The current analytic plan was designed to describe variability in the distribution of physical health characteristics across sociodemographic factors at baseline in the ABCD sample. These results highlight areas for future health disparity research, but do not aim to disentangle the complex underpinnings contributing to any disparities. Elucidating the relationship between physical health outcomes and the many contextual factors that are associated with race and sex is a very important and notably understudied area that requires extensive and exclusive attention in future studies designed to specifically address this line of inquiry. Future research should use the plethora of cultural and environmental measures collected as part of the ABCD Study to delve into the complex factors that may underlie sociodemographic disparities in physical health amongst this cohort [76].

## Conclusions and Future Directions

In the current paper, we have summarized physical health descriptive outcomes in the ABCD cohort at baseline and contrasted these with current clinical guidelines and published normative data. Given that the ABCD Study is following youth ages 9 to 10 into adulthood, an important future direction will be to conduct longitudinal analyses. Following the ABCD cohort through adolescence, a period of significant risk, will be crucial in identifying what trajectories of risk factors significantly predict transition to negative physical health outcomes. Further, the physical health category has added new measures examining nutrition and pain at the follow-up visits expanding avenues for research on physical health. The ABCD Study is also collecting deciduous (“baby”) teeth to store for future analysis of prenatal and early infant exposure to environmental toxins. Beginning at the year 2 follow-up visit and every 1-2 years from that point, all youth will be asked for a blood sample, which will be used for genetic/epigenetic analyses as well as lipid, cholesterol, iron, a CBC panel, and glucose measurements. As can be seen with the data summarized in this manuscript, the ABCD Study is well-poised to investigate not only the factors influencing physical health trajectories but also the association between physical health and other domains such as mental health, neurocognition, and substance use.

## Acknowledgements

Data used in the preparation of this article were obtained from the Adolescent Brain Cognitive Development (ABCD) Study (https://abcdstudy.org), held in the NIMH Data Archive (NDA). This is a multisite, longitudinal study designed to recruit more than 10,000 children age 9 –10 and follow them over 10 years into early adulthood. A listing of participating sites and a complete listing of the study investigators can be found at https://abcdstudy.org/principal-investigators.html. ABCD consortium investigators designed and implemented the study and/or provided data but did not necessarily participate in analysis or writing of this report. T. Dr. Gayathri Dowling was substantially involved in all of the cited grants. All other Federal representatives contributed to the interpretation of the data and participated in the preparation, review and approval of the manuscript, consistent with their roles on the ABCD Federal Partners Group. The views and opinions expressed in this manuscript are those of the authors only and do not necessarily represent the views, official policy or position of the ABCD consortium investigators or the U.S. Department of Health and Human Services or any of its affiliated institutions or agencies. In putting together this manuscript, all authors considered their positionality and proximity to the different sociodemographic groups analyzed and took time to determine whether there were any systemic biases in the measures used or the inferences made.

## Data Availability Statement

The ABCD data repository grows and changes over time. The ABCD data used in this report came from NIMH Data Archive Digital Object Identifier (DOI) 10.15154/1522647. DOIs can be found at https://dx.doi.org/10.15154/1522647

