## Supplementary Figures for "A Comprehensive Overview of the Physical Health of the Adolescent Brain Cognitive Development Study (ABCD) Cohort at Baseline"

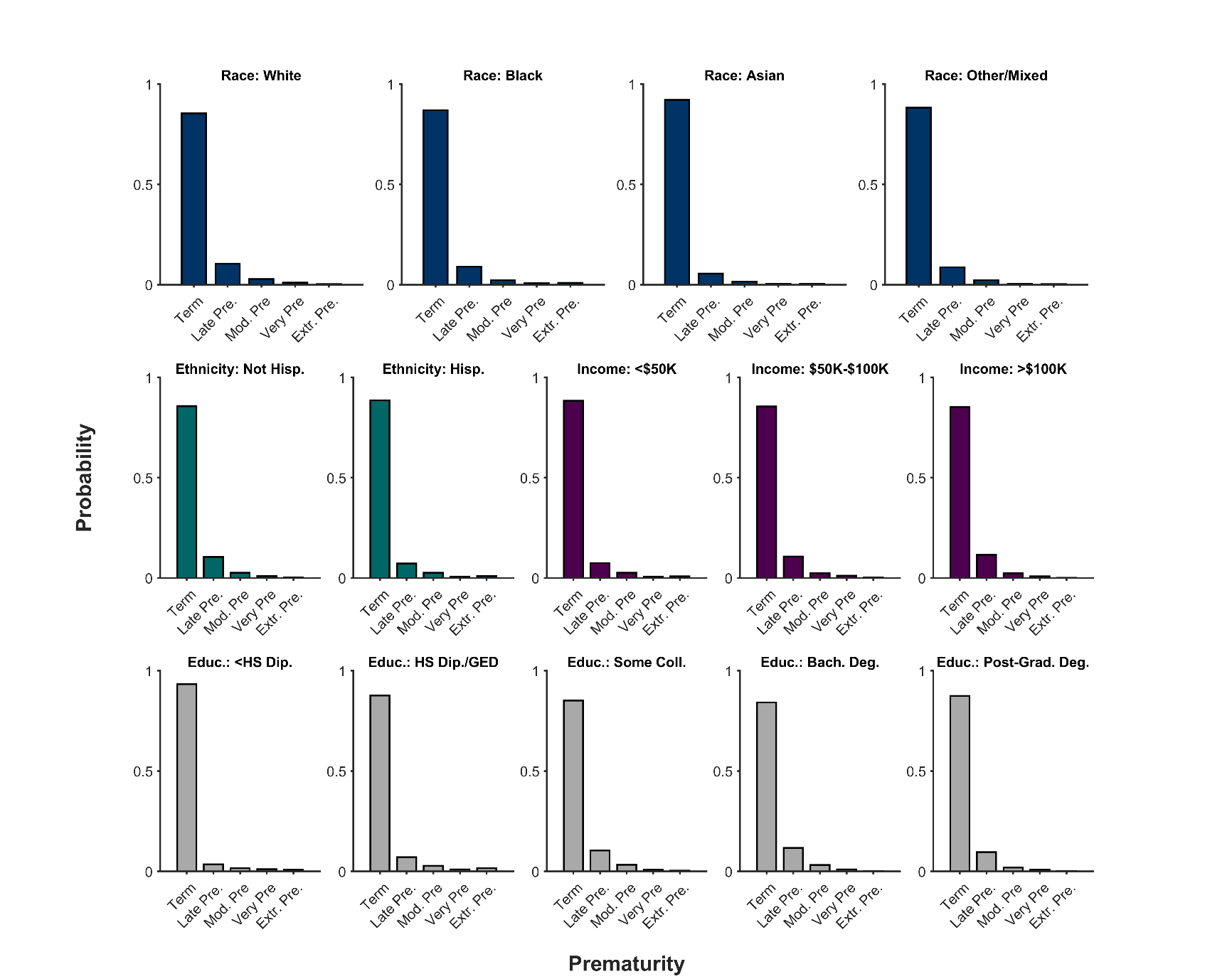

**Supplementary Figure 1**. Probability distribution (frequency histogram) of (pre)maturity status of youth participants’ birth as function of race, ethnicity, household income, and highest parental education level. Data are categorized as births being to term (Term), late pre-term (Late Pre.), moderate pre-term (Mod. Pre.), very pre-term (Very Pre.), and extremely pre-term (Extr. Pre). Hisp. = Hispanic, Educ. = Highest parental education level, HS Dip. = High school diploma, GED = General Educational Development, Coll. = college, Bach. Deg. = Bachelor’s degree.

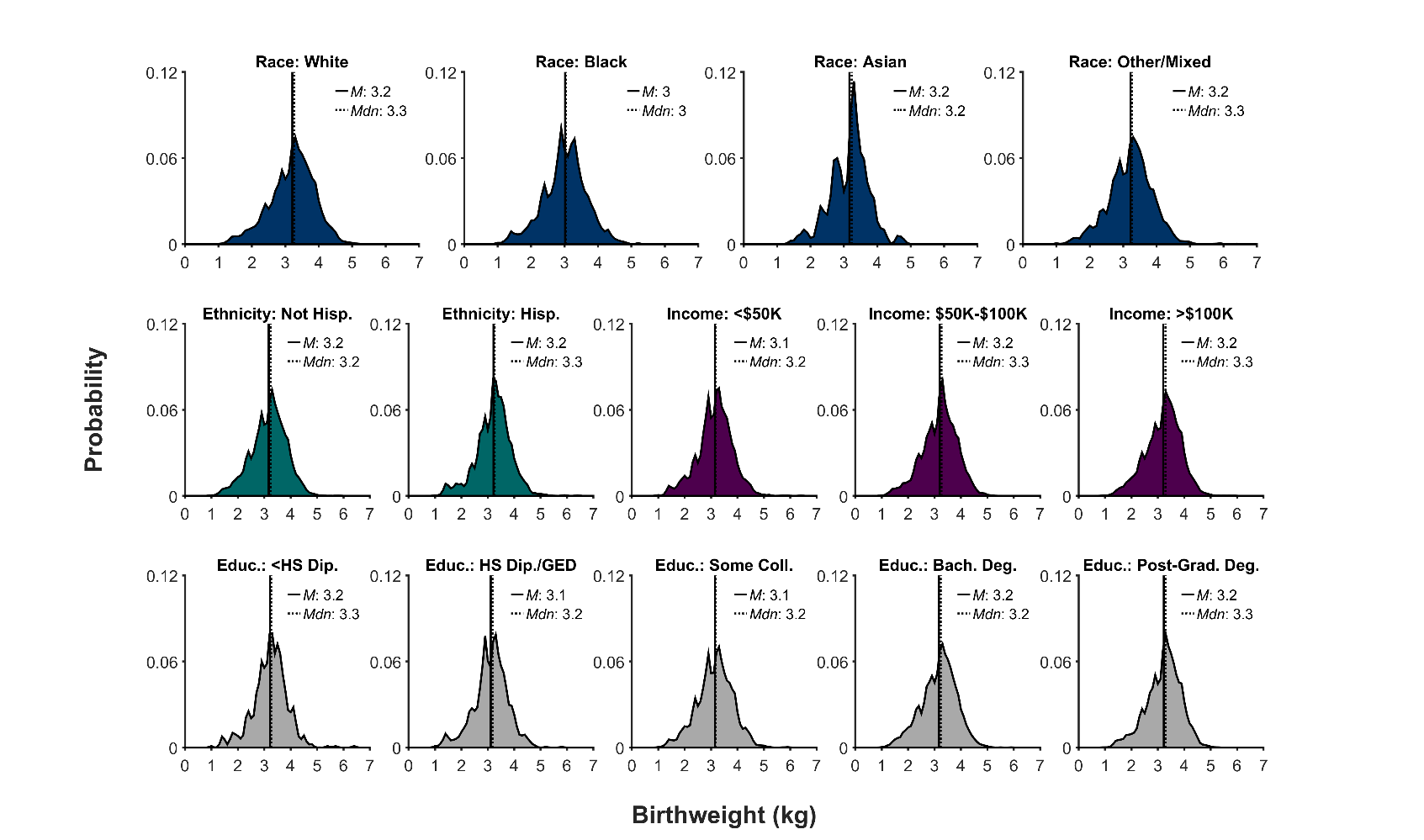

**Supplementary Figure 2**. Probability distribution (frequency histogram) of youth participants’ birthweights (in kilograms, kg) as function of race, ethnicity, household income, and highest parental education level. The solid line represents the mean (*M*) of the corresponding distribution; the dotted line, the median (*Mdn*). Hisp. = Hispanic, Educ. = Highest parental education level, HS Dip. = High school diploma, GED = General Educational Development, Coll. = college, Bach. Deg. = Bachelor’s degree.

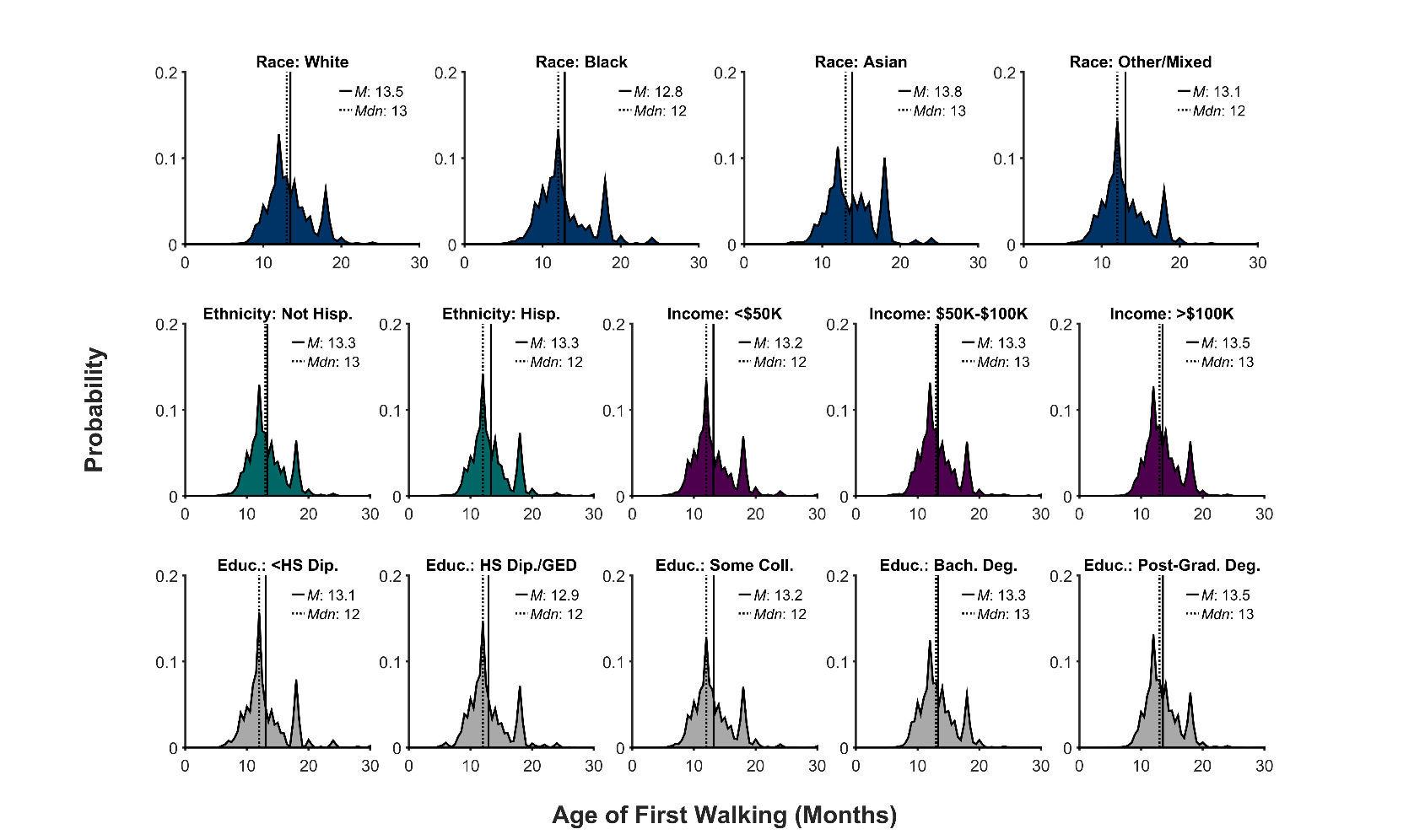

**Supplementary Figure 3**. Probability distribution (frequency histogram) of the age (in months) at which youth participants started walking as a function of race, ethnicity, household income, and highest parental education level. The solid line represents the mean (*M*) of the corresponding distribution; the dotted line, the median (*Mdn*). Hisp. = Hispanic, Educ. = Highest parental education level, HS Dip. = High school diploma, GED = General Educational Development, Coll. = college, Bach. Deg. = Bachelor’s degree.

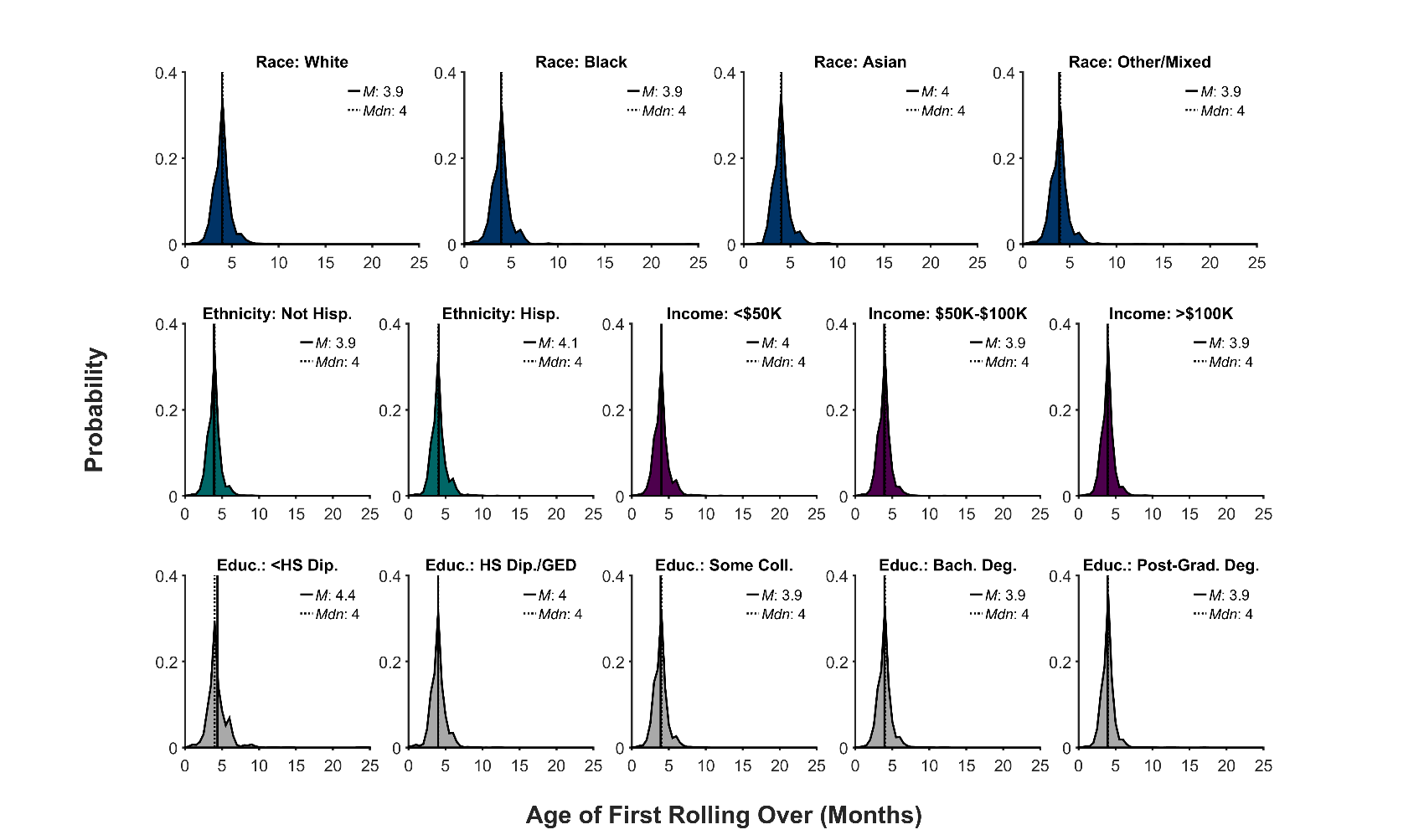

**Supplementary Figure 4**. Probability distribution (frequency histogram) of the age (in months) at which youth participants first rolled over as a function of race, ethnicity, household income, and highest parental education level. The solid line represents the mean (*M*) of the corresponding distribution; the dotted line, the median (*Mdn*). Hisp. = Hispanic, Educ. = Highest parental education level, HS Dip. = High school diploma, GED = General Educational Development, Coll. = college, Bach. Deg. = Bachelor’s degree.

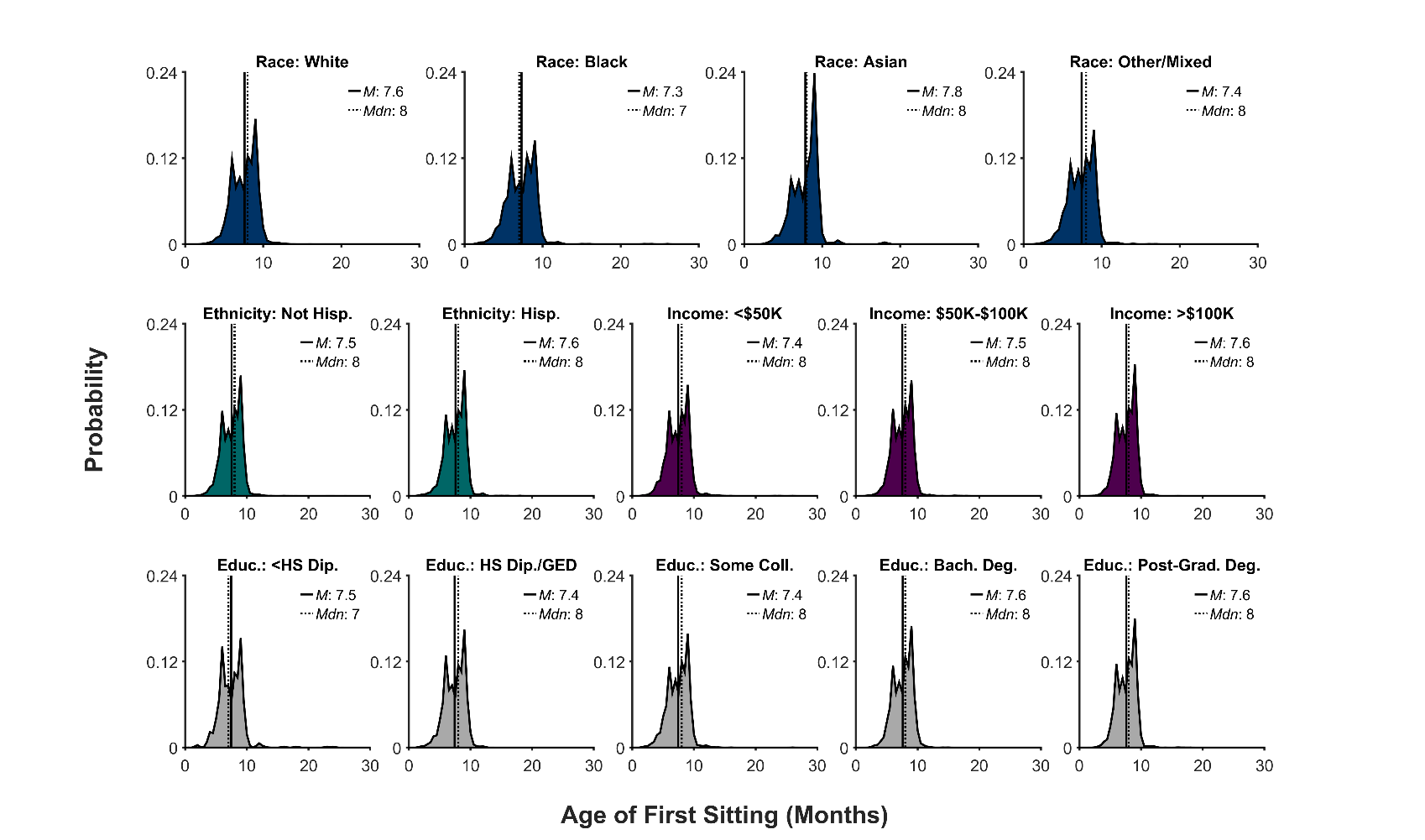

**Supplementary Figure 5**. Probability distribution (frequency histogram) of the age (in months) at which youth participants began sitting as a function of race, ethnicity, household income, and highest parental education level. The solid line represents the mean (*M*) of the corresponding distribution; the dotted line, the median (*Mdn*). Hisp. = Hispanic, Educ. = Highest parental education level, HS Dip. = High school diploma, GED = General Educational Development, Coll. = college, Bach. Deg. = Bachelor’s degree.

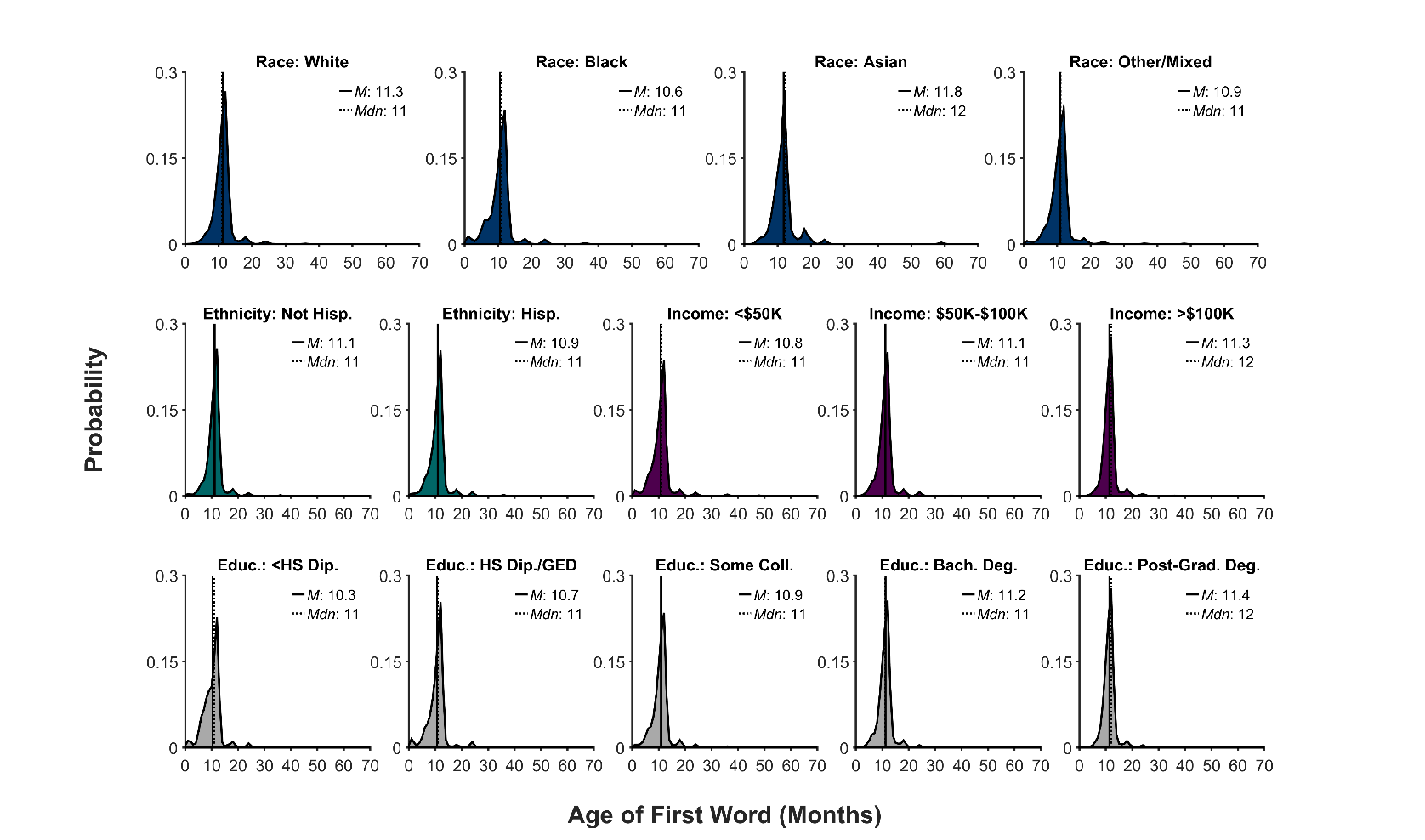

**Supplementary Figure 6**. Probability distribution (frequency histogram) of the age (in months) when youth participants said their first spoken word as function of race, ethnicity, household income, and highest parental education level. The solid line represents the mean (*M*) of the corresponding distribution; the dotted line, the median (*Mdn*). Hisp. = Hispanic, Educ. = Highest parental education level, HS Dip. = High school diploma, GED = General Educational Development, Coll. = college, Bach. Deg. = Bachelor’s degree.

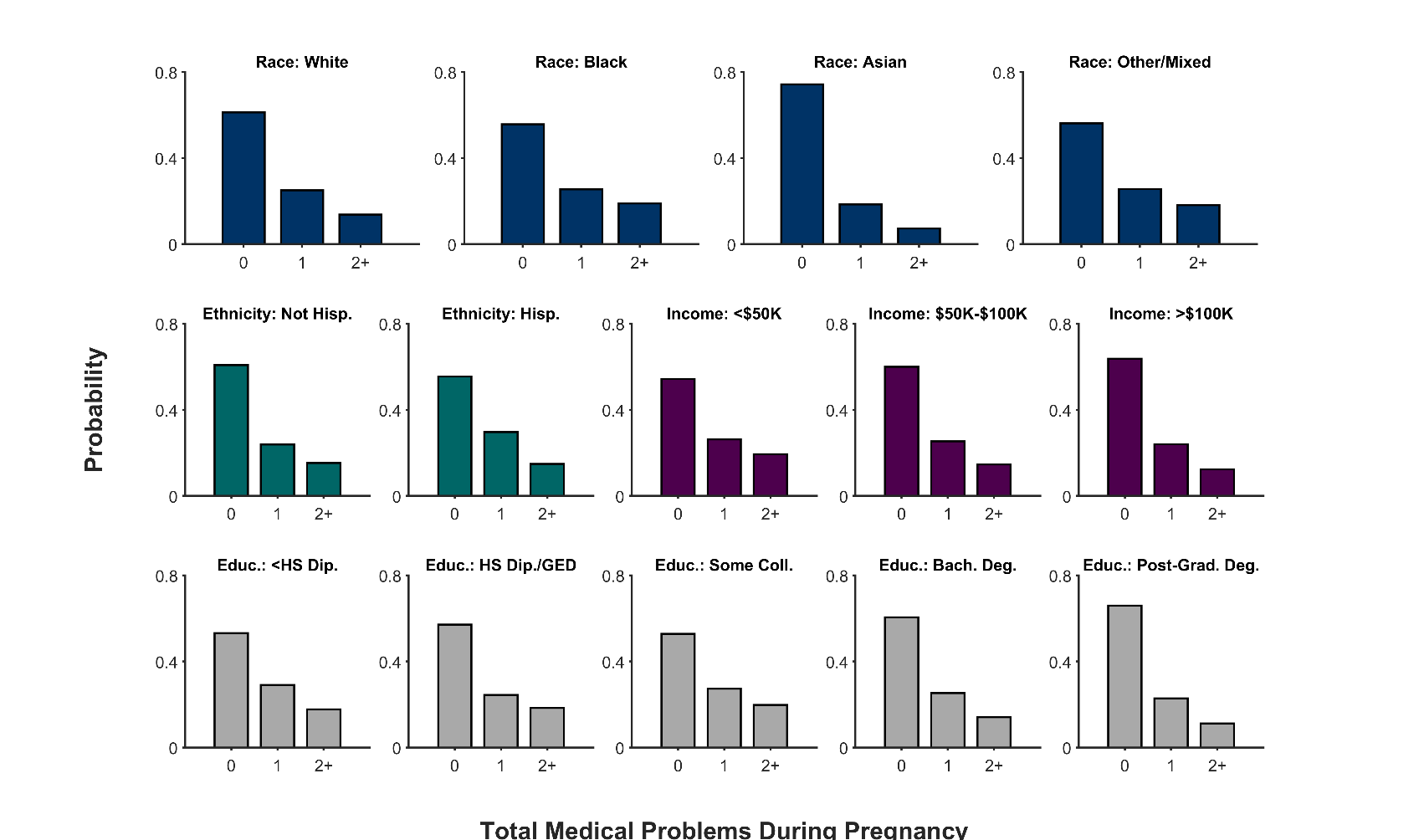

**Supplementary Figure 7**. Probability distribution (frequency histogram) of the total number of medical problems during pregnancy as function of race, ethnicity, household income, and highest parental education level. Hisp. = Hispanic, Educ. = Highest parental education level, HS Dip. = High school diploma, GED = General Educational Development, Coll. = college, Bach. Deg. = Bachelor’s degree.

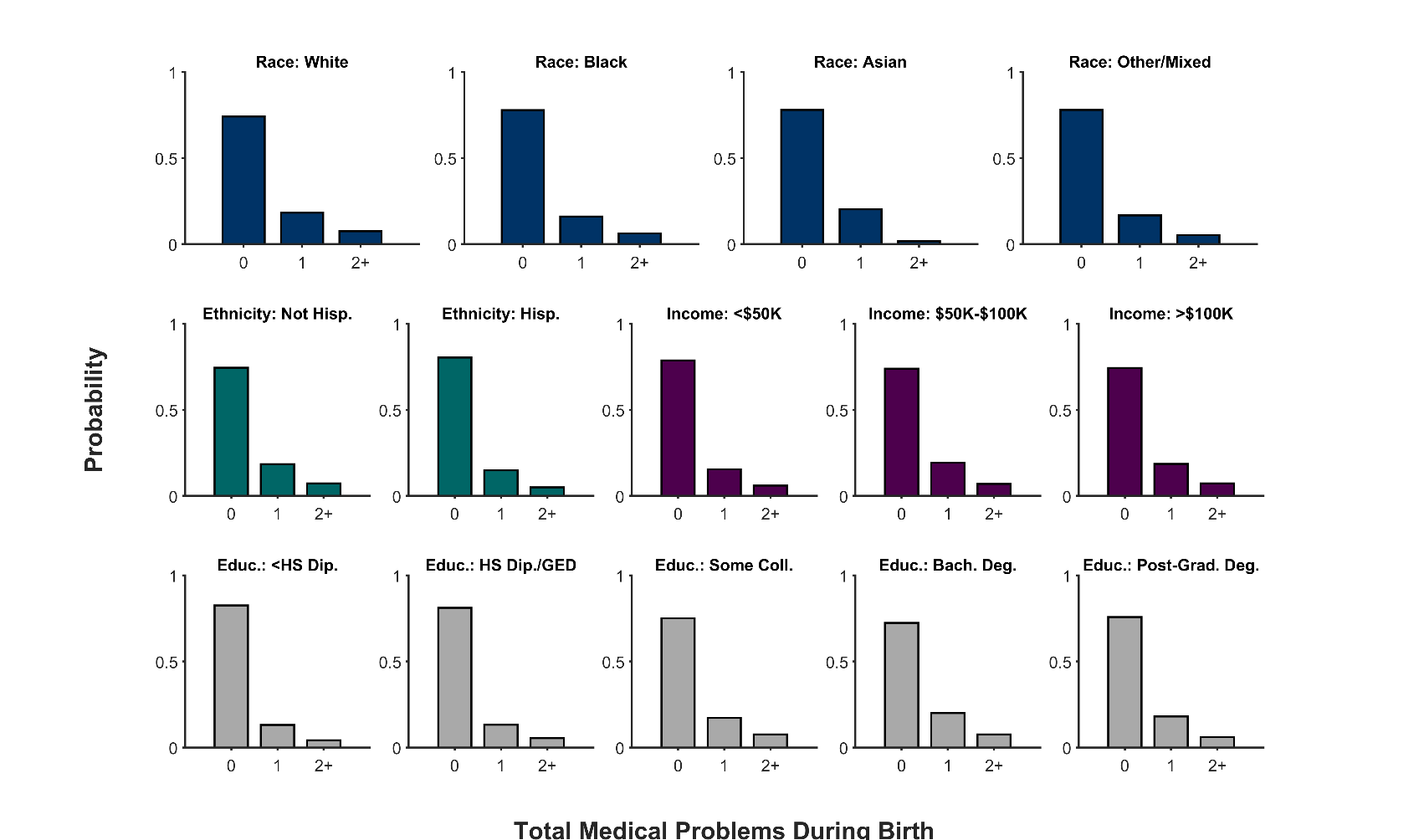

**Supplementary Figure 8**. Probability distribution (frequency histogram) of the total number of medical problems during youth participants’ birth as function of race, ethnicity, and household income, highest parental education level. Hisp. = Hispanic, Educ. = Highest parental education level, HS Dip. = High school diploma, GED = General Educational Development, Coll. = college, Bach. Deg. = Bachelor’s degree.

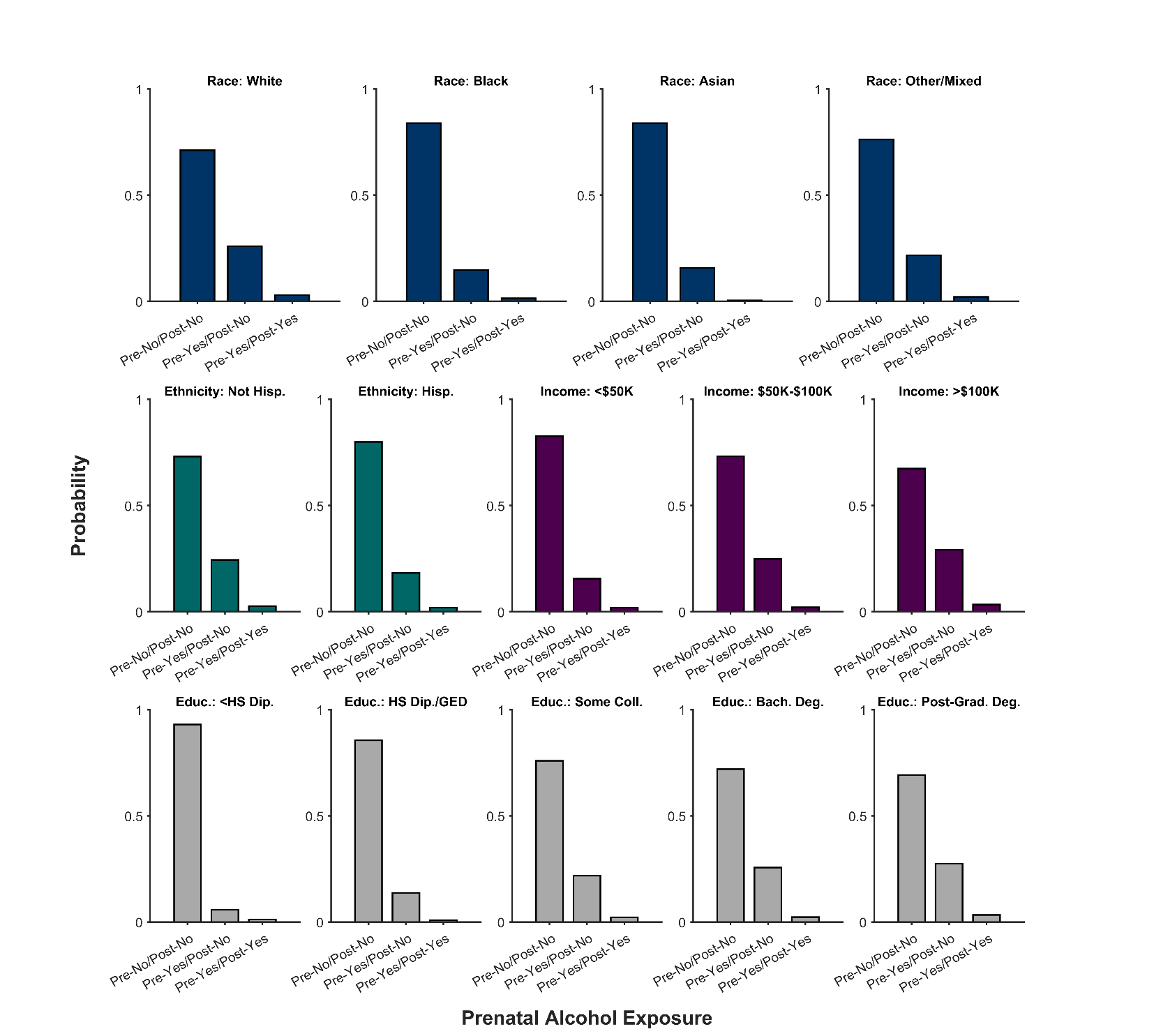

**Supplementary Figure 9**. Probability distribution (frequency histogram) of prenatal alcohol exposure status (No = no prenatal alcohol exposure, Yes = prenatal alcohol exposure) of each youth participant as function of race, ethnicity, household income, and highest parental education level. Data are categorized based on whether alcohol was consumed before (pre) or after (post) knowledge of the pregnancy. Hisp. = Hispanic, Educ. = Highest parental education level, HS Dip. = High school diploma, GED = General Educational Development, Coll. = college, Bach. Deg. = Bachelor’s degree.

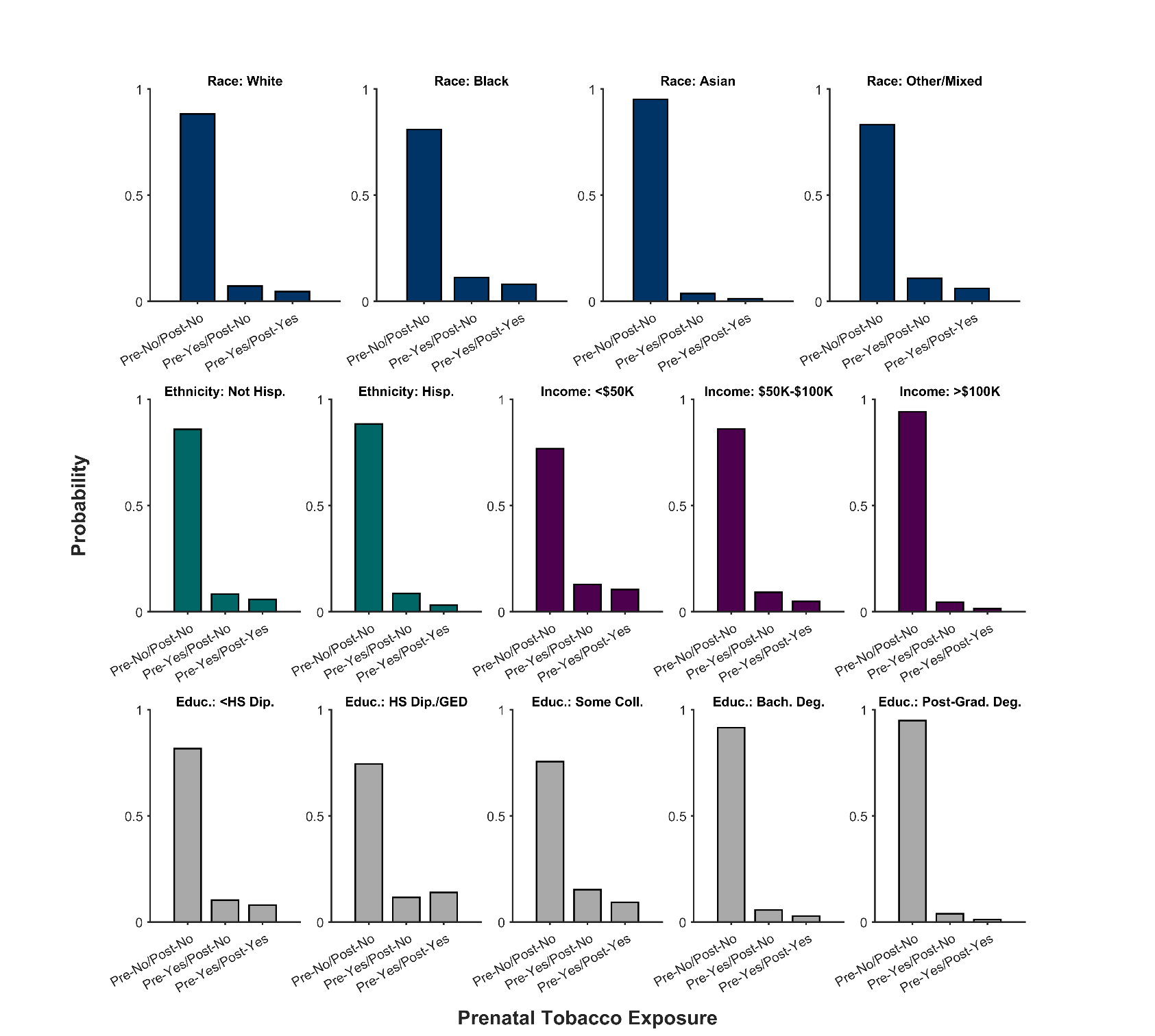

**Supplementary Figure 10**. Probability distribution (frequency histogram) of prenatal tobacco exposure status (No = no prenatal tobacco exposure, Yes = prenatal tobacco exposure) of each youth participant as function of race, ethnicity, household income, and highest parental education level. Data are categorized based on whether tobacco was used before (pre) or after (post) knowledge of the pregnancy. Hisp. = Hispanic, Educ. = Highest parental education level, HS Dip. = High school diploma, GED = General Educational Development, Coll. = college, Bach. Deg. = Bachelor’s degree.

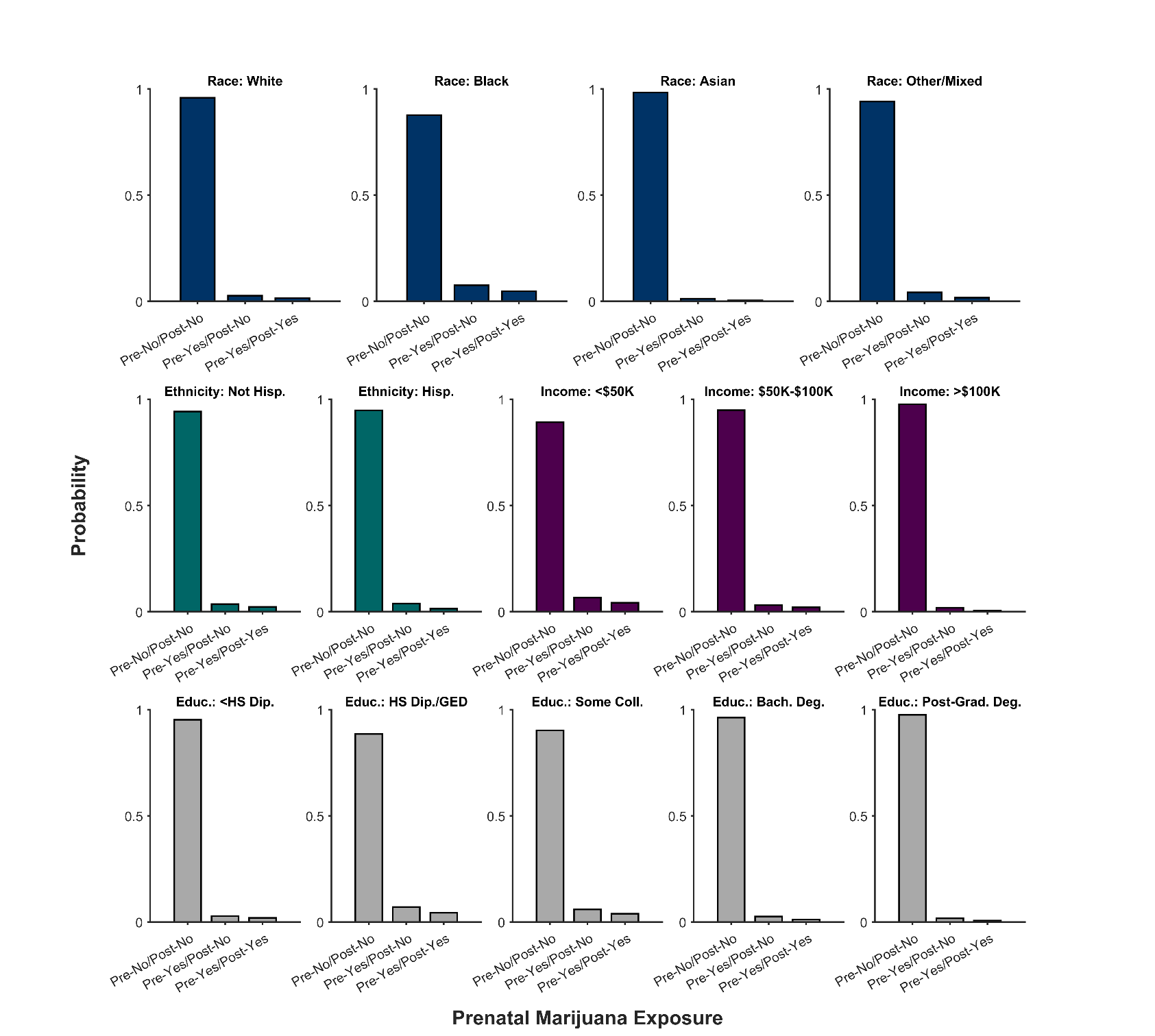

**Supplementary Figure 11**. Probability distribution (frequency histogram) of prenatal marijuana exposure status (No = no prenatal marijuana exposure, Yes = prenatal marijuana exposure) of each youth participant as function of race, ethnicity, household income, and highest parental education level. Data are categorized based on whether marijuana was used before (pre) or after (post) knowledge of the pregnancy. Hisp. = Hispanic, Educ. = Highest parental education level, HS Dip. = High school diploma, GED = General Educational Development, Coll. = college, Bach. Deg. = Bachelor’s degree.

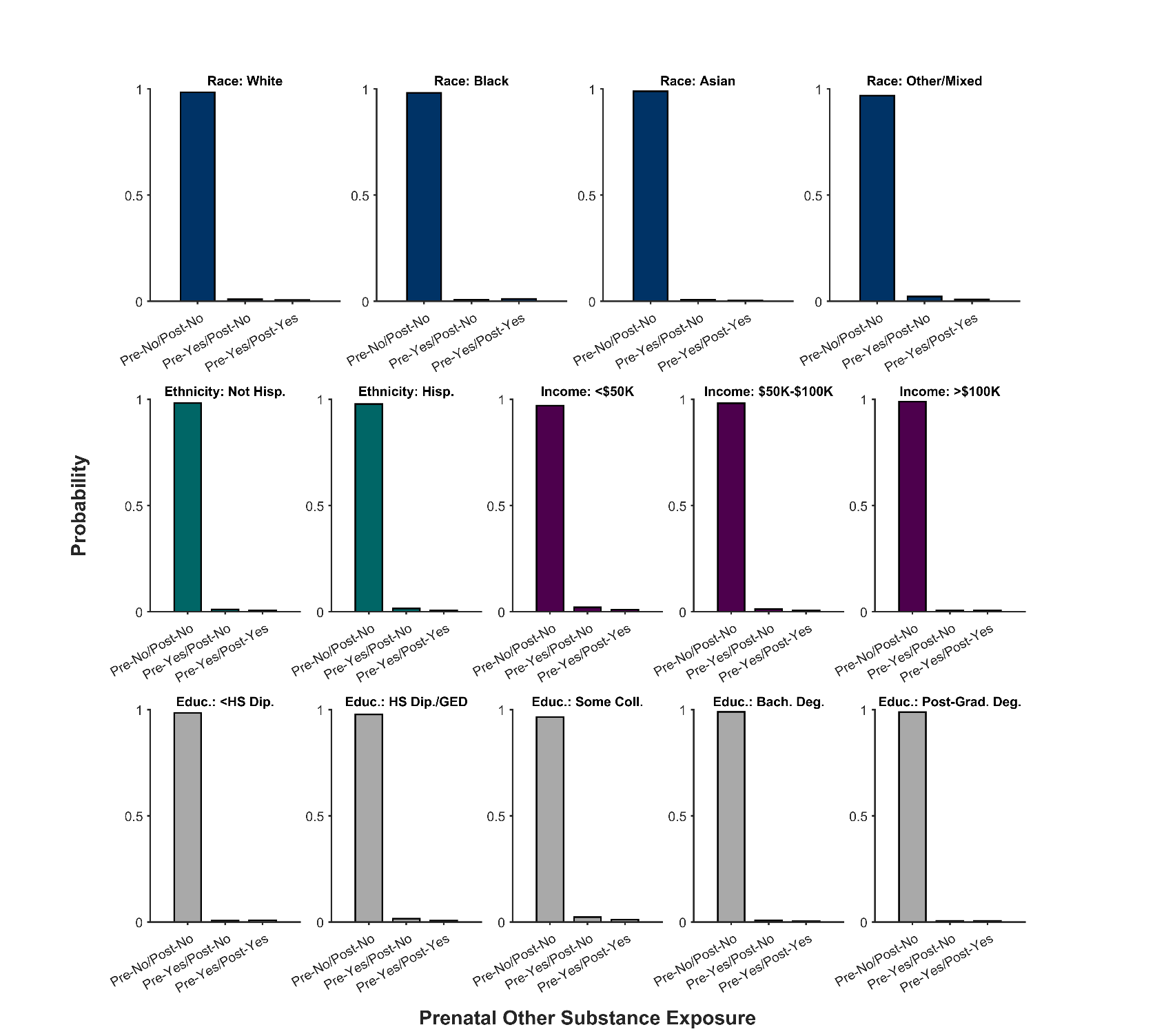

**Supplementary Figure 12**. Probability distribution (frequency histogram) of prenatal exposure status of other substances (No = no exposure, Yes = exposure) of each youth participant as function of race, ethnicity, household income, and highest parental education level. Data are categorized based on whether other substances were used before (pre) or after (post) knowledge of the pregnancy. Hisp. = Hispanic, Educ. = Highest parental education level, HS Dip. = High school diploma, GED = General Educational Development, Coll. = college, Bach. Deg. = Bachelor’s degree.

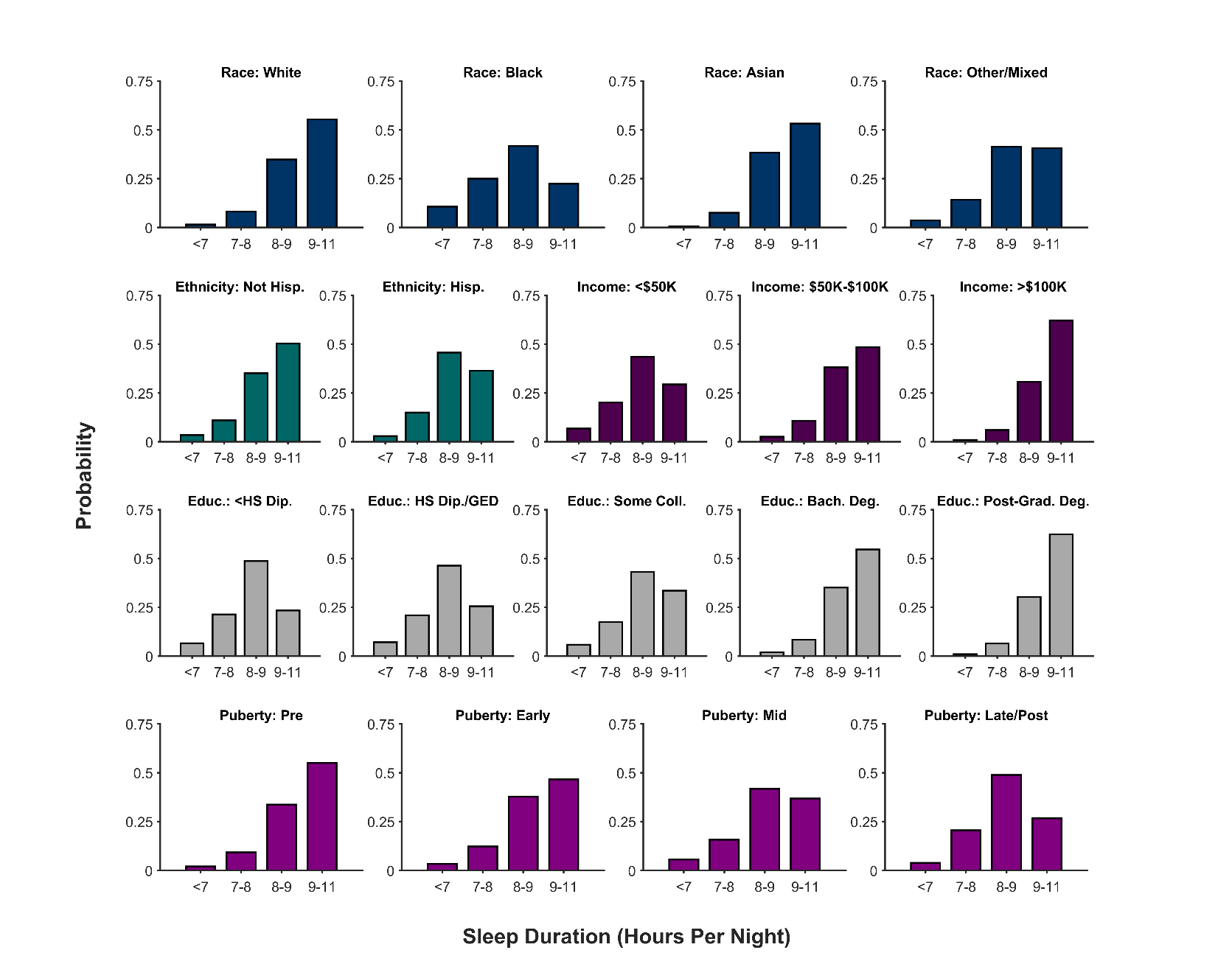

**Supplementary Figure 13**. Probability distribution (frequency histogram) of youth participants’ sleep duration (i.e., hours per night) as a function of race, ethnicity, household income, highest parental education level, and puberty status. Hisp. = Hispanic, Educ. = Highest parental education level, HS Dip. = High school diploma, GED = General Educational Development, Coll. = college, Bach. Deg. = Bachelor’s degree.

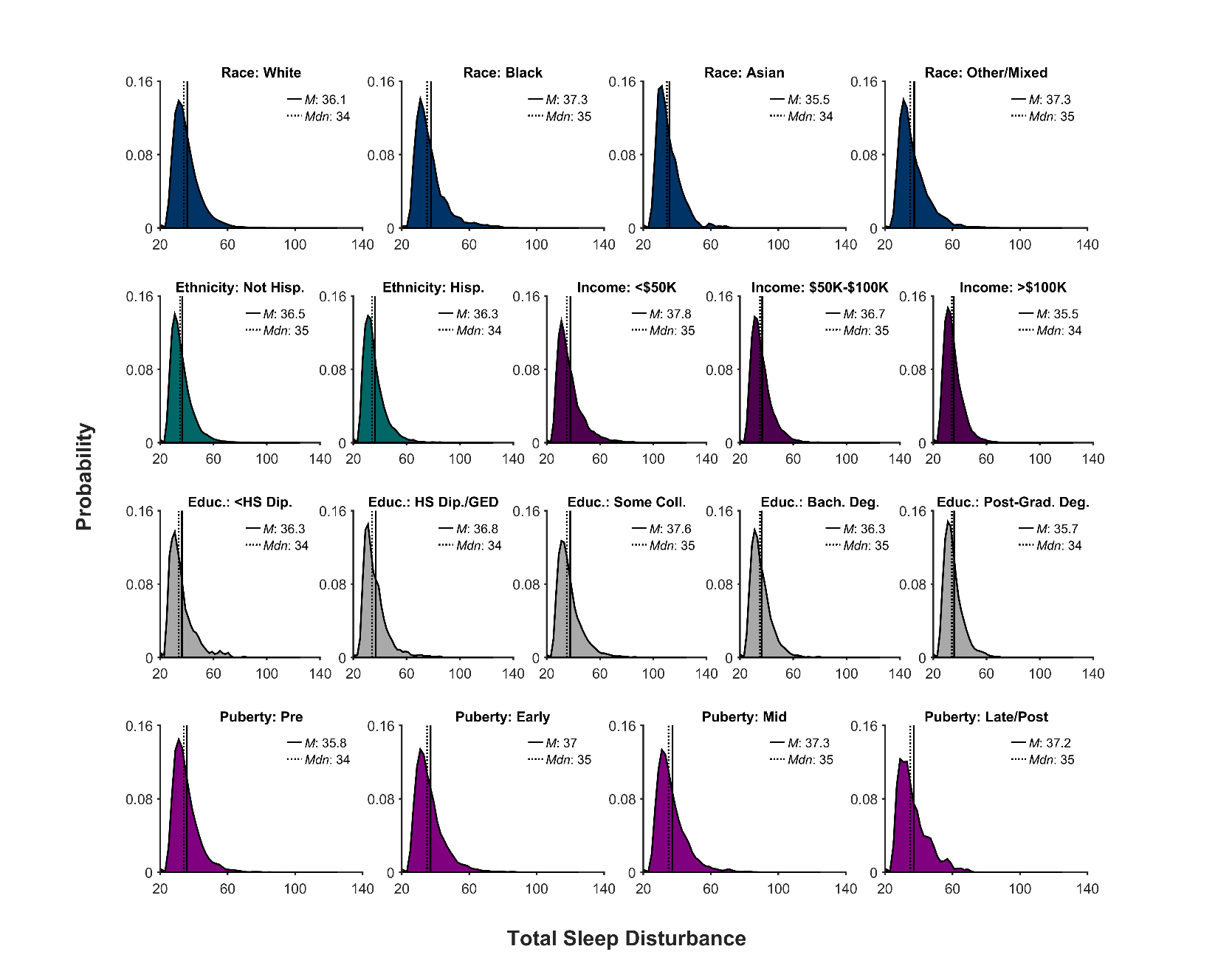

**Supplementary Figure 14**. Probability distribution (frequency histogram) of youth participants’ total sleep disturbances as a function of race, ethnicity, household income, highest parental education level, and puberty status. The solid line represents the mean (*M*) of the corresponding distribution; the dotted line, the median (*Mdn*). Hisp. = Hispanic, Educ. = Highest parental education level, HS Dip. = High school diploma, GED = General Educational Development, Coll. = college, Bach. Deg. = Bachelor’s degree.

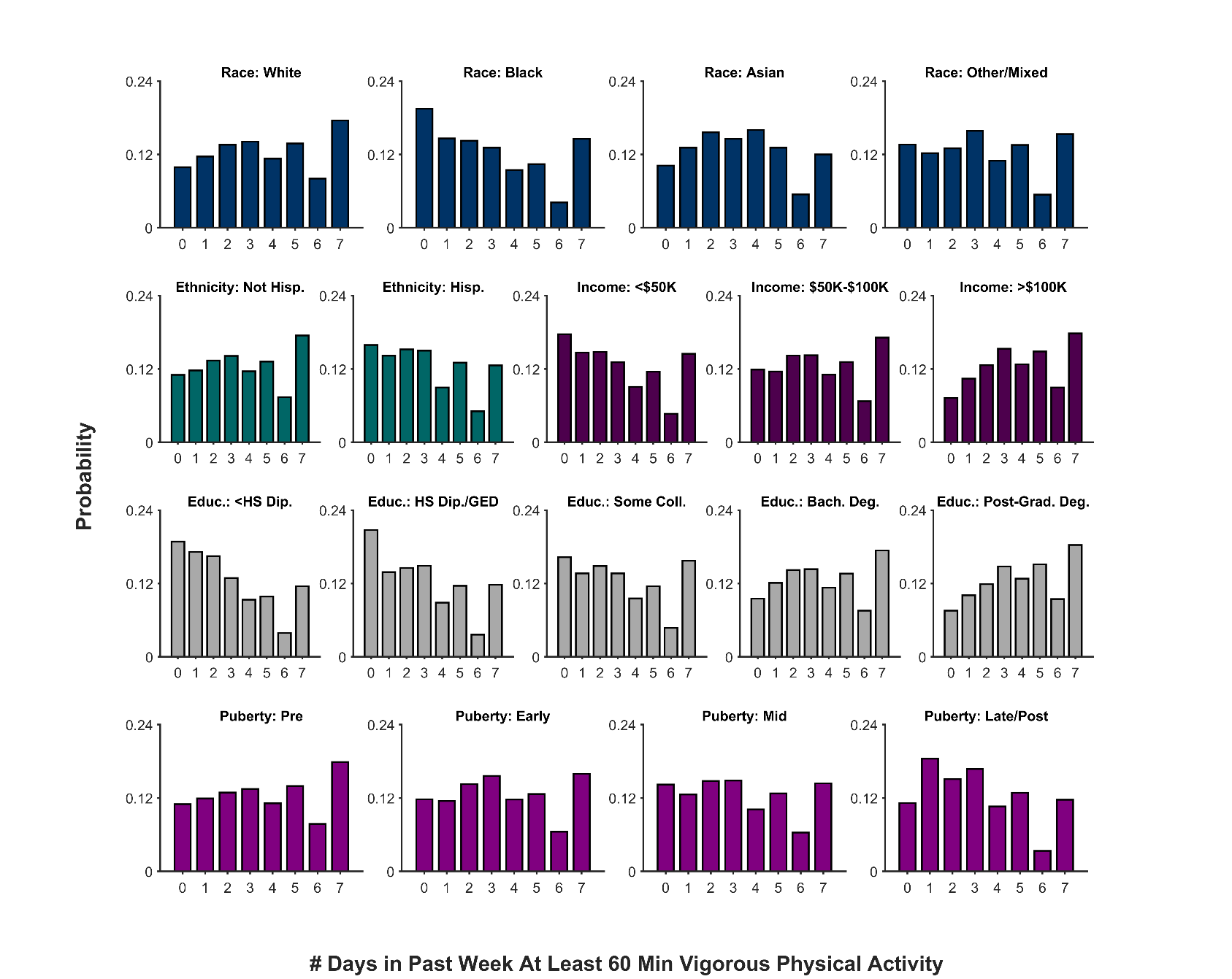

**Supplementary Figure 15**. Probability distribution (frequency histogram) of the number of days in the past week that youth participants engaged in at least 60 min of vigorous physical activity as function of race, ethnicity, household income, highest parental education level, and puberty status. Hisp. = Hispanic, Educ. = Highest parental education level, HS Dip. = High school diploma, GED = General Educational Development, Coll. = college, Bach. Deg. = Bachelor’s degree.

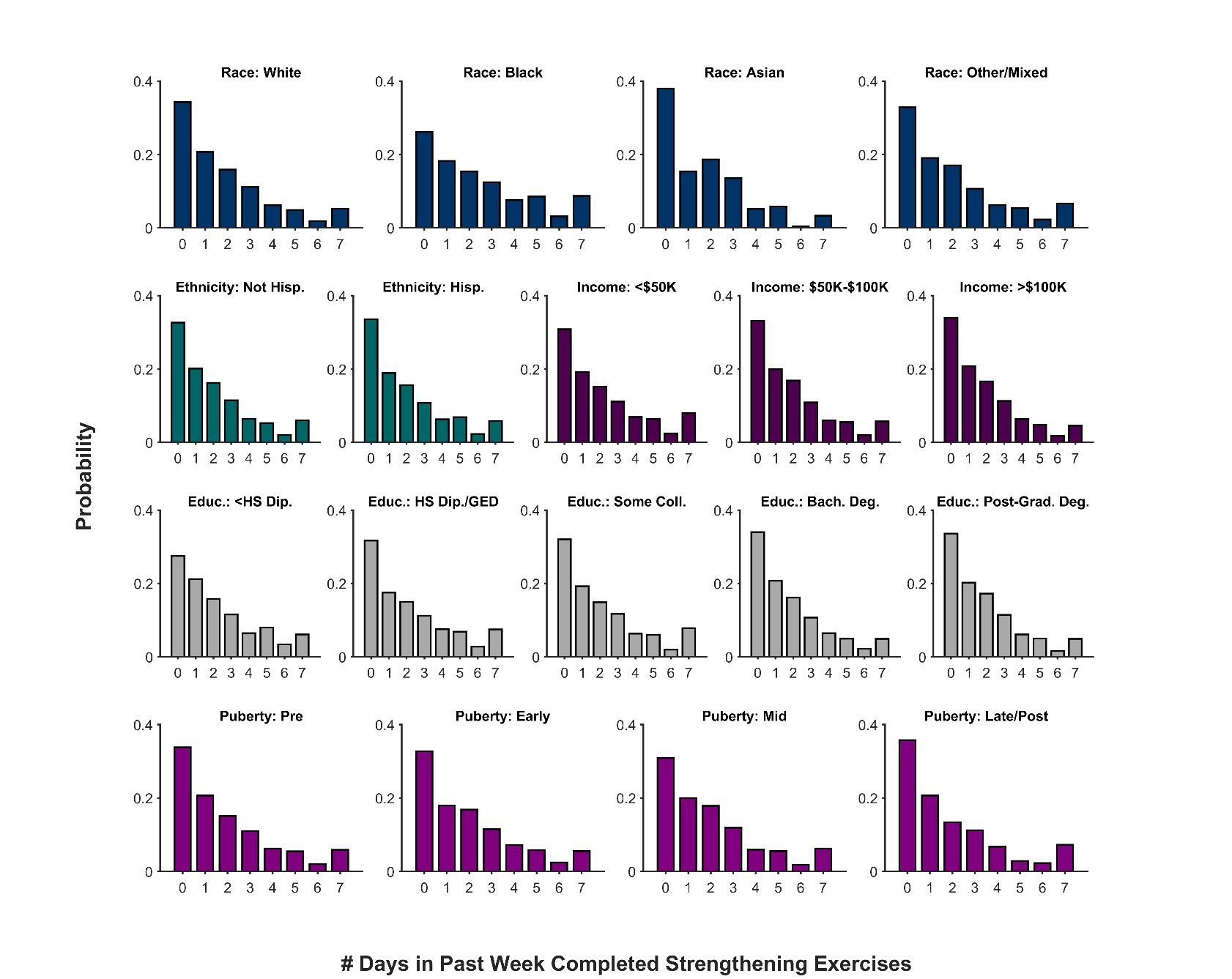

**Supplementary Figure 16**. Probability distribution (frequency histogram) of the number of days in the past week that youth participants completed strengthening exercises as function of race, ethnicity, household income, highest parental education level, and puberty status. Hisp. = Hispanic, Educ. = Highest parental education level, HS Dip. = High school diploma, GED = General Educational Development, Coll. = college, Bach. Deg. = Bachelor’s degree.

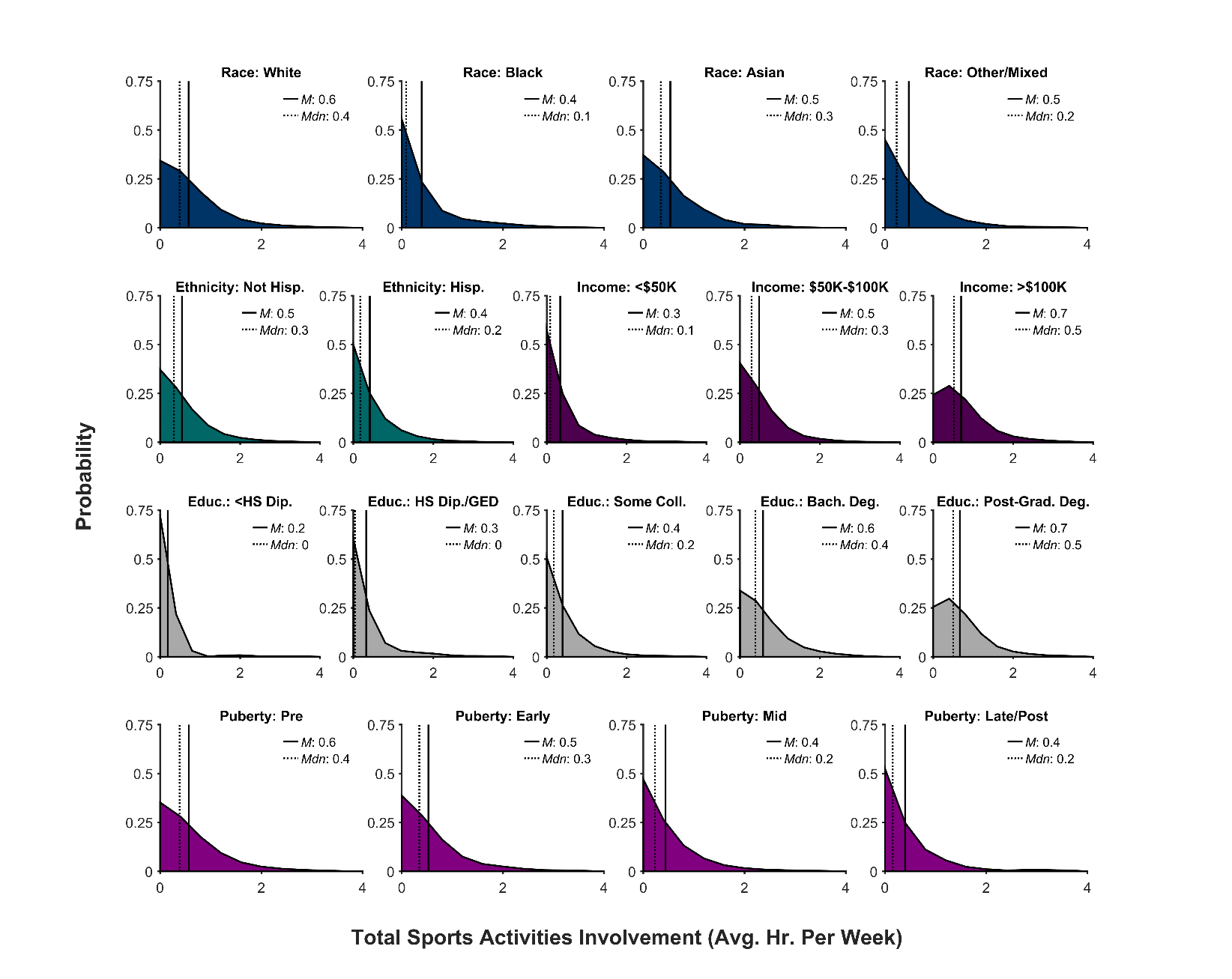

**Supplementary Figure 17**. Probability distribution (frequency histogram) of youth participants’ total sports activities involvement as a function of race, ethnicity, household income, highest parental education level, and puberty status. Total sports activities involvement was quantified as average hours per week. The solid line represents the mean (*M*) of the corresponding distribution; the dotted line, the median (*Mdn*). Hisp. = Hispanic, Educ. = Highest parental education level, HS Dip. = High school diploma, GED = General Educational Development, Coll. = college, Bach. Deg. = Bachelor’s degree.

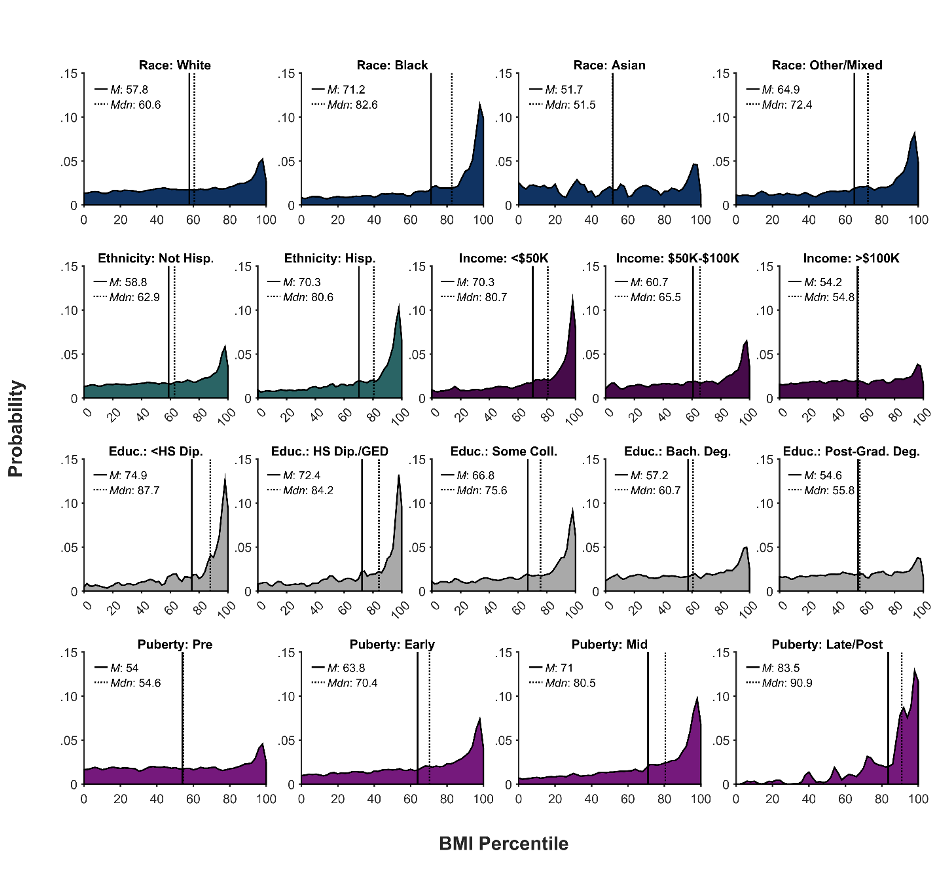

**Supplementary Figure 18**. Probability distribution (frequency histogram) of youth participants’ body mass index (BMI) percentile as a function of race, ethnicity, household income, highest parental education level, and puberty status. The solid line represents the mean (*M*) of the corresponding distribution; the dotted line, the median (*Mdn*). Hisp. = Hispanic, Educ. = Highest parental education level, HS Dip. = High school diploma, GED = General Educational Development, Coll. = college, Bach. Deg. = Bachelor’s degree.

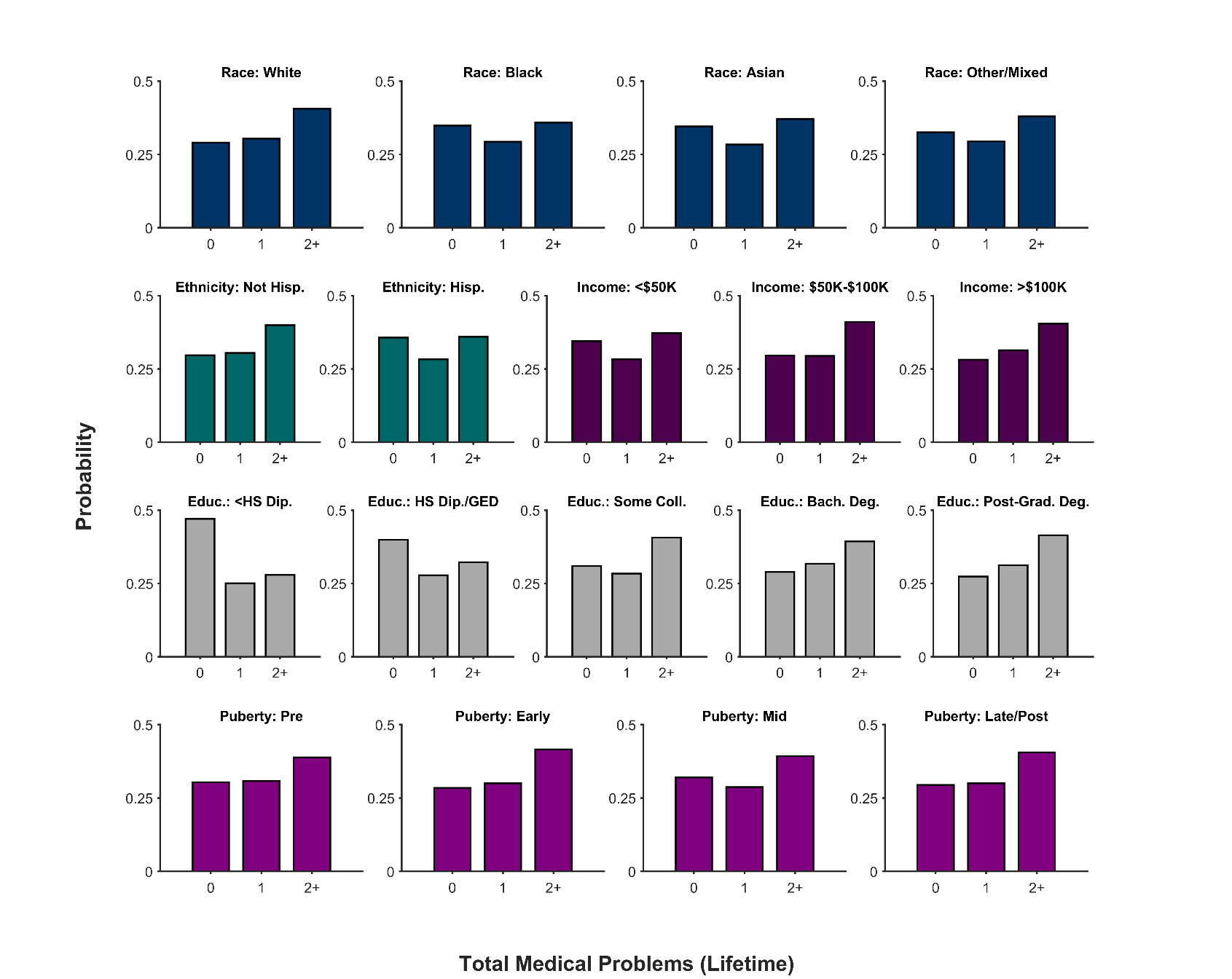

**Supplementary Figure 19**. Probability distribution (frequency histogram) of total lifetime medical problems as a function of race, ethnicity, household income, highest parental education level, and puberty status. Hisp. = Hispanic, Educ. = Highest parental education level, HS Dip. = High school diploma, GED = General Educational Development, Coll. = college, Bach. Deg. = Bachelor’s degree.

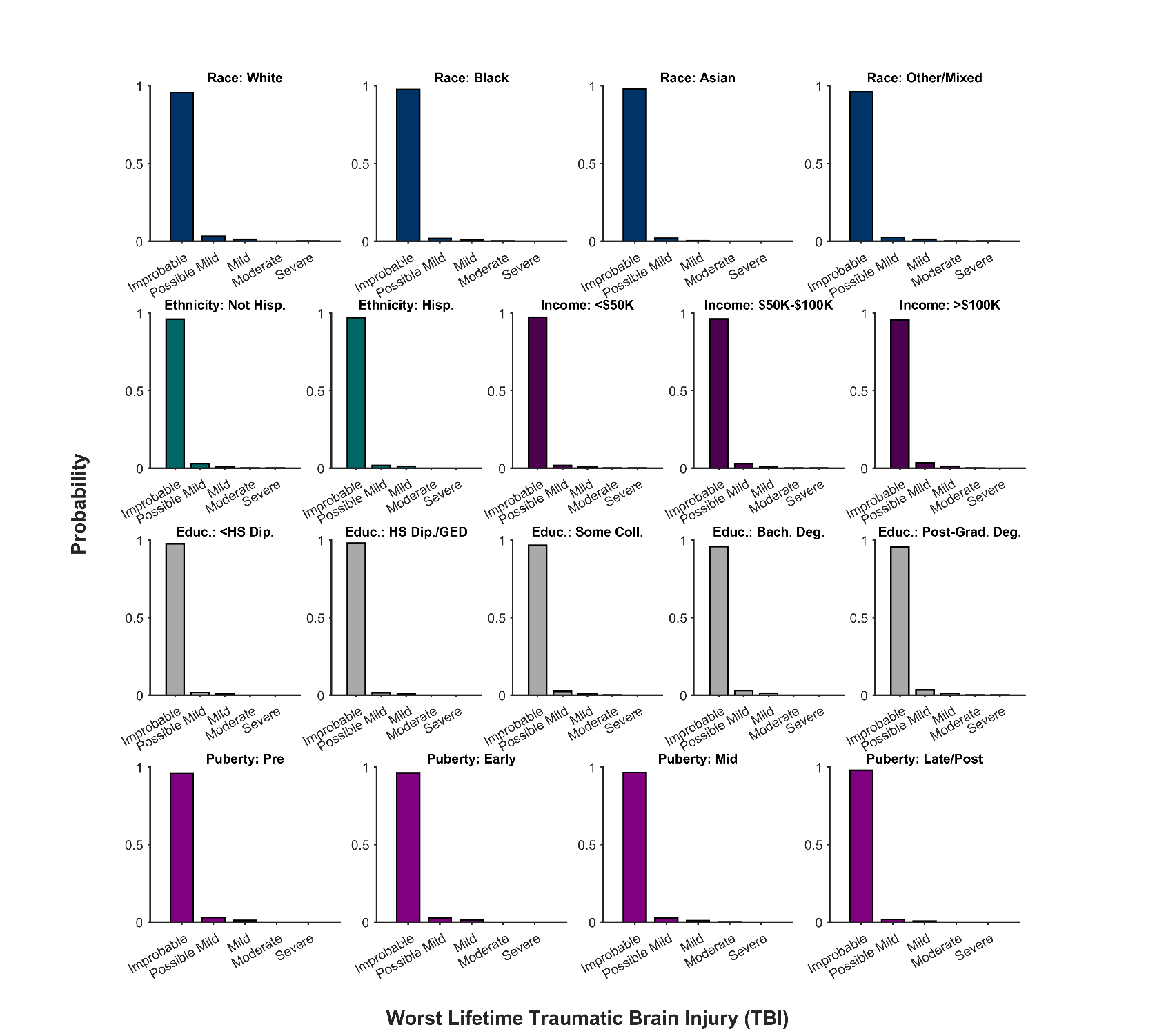

**Supplementary Figure 20**. Probability distribution (frequency histogram) of youth participants’ worse lifetime traumatic brain injury (improbable TBI, possible mild TBI, mild TBI, moderate TBI, or severe TBI) as function of race, ethnicity, household income, highest parental education level, and puberty status. Hisp. = Hispanic, Educ. = Highest parental education level, HS Dip. = High school diploma, GED = General Educational Development, Coll. = college, Bach. Deg. = Bachelor’s degree.
