## Supplementary Tables 1-20 for "A Comprehensive Overview of the Physical Health of the Adolescent Brain Cognitive Development Study (ABCD) Cohort at Baseline"

*Supplementary Table 1. Contingency table for variable of interest: Birth weight. Full sample.*

| **Demographics** | **n** | **Very low** | **Low** | **Normal** | **High** |
| --- | --- | --- | --- | --- | --- |
| **sex_at_birth** |  |  |  |  |  |
| F | 4884 | 67 ( 1.4) | 787 (16.1) | 3717 (76.1) | 313 ( 6.4) |
| *weighted* |  | ( 1.6) | (16.3) | (75.8) | ( 6.2) |
| M | 5290 | 43 ( 0.8) | 710 (13.4) | 3981 (75.3) | 556 (10.5) |
| *weighted* |  | ( 0.9) | (13.3) | (75.2) | (10.6) |
| **race.4level** |  |  |  |  |  |
| White | 6811 | 75 ( 1.1) | 959 (14.1) | 5152 (75.6) | 625 ( 9.2) |
| *weighted* |  | ( 1.3) | (14.3) | (75.3) | ( 9.1) |
| Black | 1450 | 25 ( 1.7) | 288 (19.9) | 1061 (73.2) | 76 ( 5.2) |
| *weighted* |  | ( 1.7) | (20.4) | (72.8) | ( 5.1) |
| Asian | 208 | 0 ( 0.0) | 27 (13.0) | 170 (81.7) | 11 ( 5.3) |
| *weighted* |  | ( 0.0) | (13.2) | (80.9) | ( 5.8) |
| Other/Mixed | 1705 | 10 ( 0.6) | 223 (13.1) | 1315 (77.1) | 157 ( 9.2) |
| *weighted* |  | ( 0.7) | (12.2) | (77.9) | ( 9.2) |
| **hisp** |  |  |  |  |  |
| No | 8264 | 81 ( 1.0) | 1284 (15.5) | 6182 (74.8) | 717 ( 8.7) |
| *weighted* |  | ( 1.1) | (15.8) | (74.5) | ( 8.6) |
| Yes | 1910 | 29 ( 1.5) | 213 (11.2) | 1516 (79.4) | 152 ( 8.0) |
| *weighted* |  | ( 1.6) | (11.2) | (79.2) | ( 8.0) |
| **household.income** |  |  |  |  |  |
| [<50K] | 2912 | 42 ( 1.4) | 426 (14.6) | 2228 (76.5) | 216 ( 7.4) |
| *weighted* |  | ( 1.5) | (14.3) | (76.5) | ( 7.7) |
| [>=50K & <100K] | 2897 | 35 ( 1.2) | 406 (14.0) | 2188 (75.5) | 268 ( 9.3) |
| *weighted* |  | ( 1.4) | (14.4) | (75.2) | ( 9.1) |
| [>=100K] | 4365 | 33 ( 0.8) | 665 (15.2) | 3282 (75.2) | 385 ( 8.8) |
| *weighted* |  | ( 0.8) | (15.8) | (74.6) | ( 8.8) |
| **high.educ** |  |  |  |  |  |
| < HS Diploma | 370 | <10 | 48 (13.0) | 286 (77.3) | 30 ( 8.1) |
| *weighted* |  |  | (11.5) | (79.7) | ( 7.5) |
| HS Diploma/GED | 835 | 12 ( 1.4) | 116 (13.9) | 652 (78.1) | 55 ( 6.6) |
| *weighted* |  | ( 1.7) | (14.2) | (77.5) | ( 6.7) |
| Some College | 2624 | 35 ( 1.3) | 435 (16.6) | 1931 (73.6) | 223 ( 8.5) |
| *weighted* |  | ( 1.4) | (16.5) | (73.8) | ( 8.3) |
| Bachelor | 2706 | 23 ( 0.8) | 429 (15.9) | 2025 (74.8) | 229 ( 8.5) |
| *weighted* |  | ( 1.0) | (15.8) | (74.2) | ( 8.9) |
| Post Graduate Degree | 3639 | 34 ( 0.9) | 469 (12.9) | 2804 (77.1) | 332 ( 9.1) |
| *weighted* |  | ( 1.1) | (12.9) | (77.1) | ( 8.9) |

*Supplementary Table 2. Contingency table for variable of interest: Prematurity. Full sample.*

| **Demographics** | **n** | **Term** | **Late Preterm** | **Moderate Preterm** | **Very Preterm** | **Extremely Preterm** |
| --- | --- | --- | --- | --- | --- | --- |
| **sex_at_birth** |  |  |  |  |  |  |
| F | 5010 | 4321 (86.2) | 507 (10.1) | 121 ( 2.4) | 45 ( 0.9) | 16 ( 0.3) |
| *weighted* |  | (86.1) | ( 9.8) | ( 2.7) | ( 1.0) | ( 0.4) |
| M | 5452 | 4690 (86.0) | 552 (10.1) | 141 ( 2.6) | 48 ( 0.9) | 21 ( 0.4) |
| *weighted* |  | (85.9) | (10.0) | ( 2.5) | ( 1.0) | ( 0.5) |
| **race.4level** |  |  |  |  |  |  |
| White | 6946 | 5929 (85.4) | 740 (10.7) | 183 ( 2.6) | 78 ( 1.1) | 16 ( 0.2) |
| *weighted* |  | (85.0) | (10.7) | ( 2.8) | ( 1.3) | ( 0.3) |
| Black | 1528 | 1323 (86.6) | 150 ( 9.8) | 34 ( 2.2) | <10 | 14 ( 0.9) |
| *weighted* |  | (86.6) | ( 9.8) | ( 2.2) |  | ( 0.9) |
| Asian | 215 | 201 (93.5) | 11 ( 5.1) | <10 | <10 | <10 |
| *weighted* |  | (93.6) | ( 4.7) |  |  |  |
| Other/Mixed | 1773 | 1558 (87.9) | 158 ( 8.9) | 43 ( 2.4) | <10 | <10 |
| *weighted* |  | (88.9) | ( 7.7) | ( 2.6) |  |  |
| **hisp** |  |  |  |  |  |  |
| No | 8486 | 7271 (85.7) | 908 (10.7) | 208 ( 2.5) | 78 ( 0.9) | 21 ( 0.2) |
| *weighted* |  | (85.4) | (10.7) | ( 2.6) | ( 1.1) | ( 0.3) |
| Yes | 1976 | 1740 (88.1) | 151 ( 7.6) | 54 ( 2.7) | 15 ( 0.8) | 16 ( 0.8) |
| *weighted* |  | (88.1) | ( 7.4) | ( 2.8) | ( 0.7) | ( 1.0) |
| **household.income** |  |  |  |  |  |  |
| [<50K] | 3035 | 2674 (88.1) | 230 ( 7.6) | 81 ( 2.7) | 24 ( 0.8) | 26 ( 0.9) |
| *weighted* |  | (87.8) | ( 7.6) | ( 3.0) | ( 0.8) | ( 0.8) |
| [>=50K & <100K] | 2982 | 2551 (85.5) | 320 (10.7) | 72 ( 2.4) | 33 ( 1.1) | <10 |
| *weighted* |  | (85.0) | (11.0) | ( 2.3) | ( 1.5) |  |
| [>=100K] | 4445 | 3786 (85.2) | 509 (11.5) | 109 ( 2.5) | 36 ( 0.8) | <10 |
| *weighted* |  | (84.8) | (11.7) | ( 2.6) | ( 0.8) |  |
| **high.educ** |  |  |  |  |  |  |
| < HS Diploma | 395 | 364 (92.2) | 17 ( 4.3) | <10 | <10 | <10 |
| *weighted* |  | (92.1) | ( 4.0) | <10 |  |  |
| HS Diploma/GED | 872 | 767 (88.0) | 63 ( 7.2) | 19 ( 2.2) | 10 ( 1.1) | 13 ( 1.5) |
| *weighted* |  | (87.2) | ( 7.5) | ( 2.7) | ( 1.3) | ( 1.3) |
| Some College | 2694 | 2275 (84.4) | 297 (11.0) | 91 ( 3.4) | 20 ( 0.7) | 11 ( 0.4) |
| *weighted* |  | (84.5) | (10.9) | ( 3.2) | ( 0.9) | ( 0.4) |
| Bachelor | 2770 | 2331 (84.2) | 332 (12.0) | 78 ( 2.8) | 25 ( 0.9) | <10 |
| *weighted* |  | (84.1) | (11.8) | ( 2.9) | ( 1.1) |  |
| Post Graduate Degree | 3731 | 3274 (87.8) | 350 ( 9.4) | 67 ( 1.8) | 33 ( 0.9) | <10 |
| *weighted* |  | (87.8) | ( 9.1) | ( 1.8) | ( 1.0) |  |

*Supplementary Table 3. Contingency table for variable of interest: Age at first rolling over (months). Full sample.*

| **Demographics** | **n** | **Within guidelines** | **Beyond guidelines** |
| --- | --- | --- | --- |
| **sex_at_birth** |  |  |  |
| F | 5085 | 5026 (98.8) | 59 ( 1.2) |
| *weighted* |  | (98.7) | ( 1.3) |
| M | 5515 | 5449 (98.8) | 66 ( 1.2) |
| *weighted* |  | (98.7) | ( 1.3) |
| **race.4level** |  |  |  |
| White | 6998 | 6910 (98.7) | 88 ( 1.3) |
| *weighted* |  | (98.6) | ( 1.4) |
| Black | 1570 | 1553 (98.9) | 17 ( 1.1) |
| *weighted* |  | (98.9) | ( 1.1) |
| Asian | 235 | 233 (99.1) | <10 |
| *weighted* |  | (99.2) |  |
| Other/Mixed | 1797 | 1779 (99.0) | 18 ( 1.0) |
| *weighted* |  | (98.9) | ( 1.1) |
| **hisp** |  |  |  |
| No | 8593 | 8501 (98.9) | 92 ( 1.1) |
| *weighted* |  | (98.8) | ( 1.2) |
| Yes | 2007 | 1974 (98.4) | 33 ( 1.6) |
| *weighted* |  | (98.2) | ( 1.8) |
| **household.income** |  |  |  |
| [<50K] | 3088 | 3038 (98.4) | 50 ( 1.6) |
| *weighted* |  | (98.3) | ( 1.7) |
| [>=50K & <100K] | 3014 | 2982 (98.9) | 32 ( 1.1) |
| *weighted* |  | (99.0) | ( 1.0) |
| [>=100K] | 4498 | 4455 (99.0) | 43 ( 1.0) |
| *weighted* |  | (98.9) | ( 1.1) |
| **high.educ** |  |  |  |
| < HS Diploma | 402 | 391 (97.3) | 11 ( 2.7) |
| *weighted* |  | (97.2) | ( 2.8) |
| HS Diploma/GED | 891 | 878 (98.5) | 13 ( 1.5) |
| *weighted* |  | (98.6) | ( 1.4) |
| Some College | 2728 | 2693 (98.7) | 35 ( 1.3) |
| *weighted* |  | (98.6) | ( 1.4) |
| Bachelor | 2798 | 2763 (98.7) | 35 ( 1.3) |
| *weighted* |  | (98.8) | ( 1.2) |
| Post Graduate Degree | 3781 | 3750 (99.2) | 31 ( 0.8) |
| *weighted* |  | (99.0) | ( 1.0) |

*Supplementary Table 4. Contingency table for variable of interest: Age at first sitting (months). Full sample.*

| **Demographics** | **n** | **Within guidelines** | **Beyond guidelines** |
| --- | --- | --- | --- |
| **sex_at_birth** |  |  |  |
| F | 5085 | 4908 (96.5) | 177 ( 3.5) |
| *weighted* |  | (96.2) | ( 3.8) |
| M | 5515 | 5242 (95.0) | 273 ( 5.0) |
| *weighted* |  | (94.5) | ( 5.5) |
| **race.4level** |  |  |  |
| White | 6998 | 6668 (95.3) | 330 ( 4.7) |
| *weighted* |  | (94.7) | ( 5.3) |
| Black | 1570 | 1512 (96.3) | 58 ( 3.7) |
| *weighted* |  | (96.5) | ( 3.5) |
| Asian | 235 | 227 (96.6) | <10 |
| *weighted* |  | (96.4) |  |
| Other/Mixed | 1797 | 1743 (97.0) | 54 ( 3.0) |
| *weighted* |  | (97.2) | ( 2.8) |
| **hisp** |  |  |  |
| No | 8593 | 8236 (95.8) | 357 ( 4.2) |
| *weighted* |  | (95.3) | ( 4.7) |
| Yes | 2007 | 1914 (95.4) | 93 ( 4.6) |
| *weighted* |  | (95.4) | ( 4.6) |
| **household.income** |  |  |  |
| [<50K] | 3088 | 2937 (95.1) | 151 ( 4.9) |
| *weighted* |  | (94.6) | ( 5.4) |
| [>=50K & <100K] | 3014 | 2888 (95.8) | 126 ( 4.2) |
| *weighted* |  | (95.5) | ( 4.5) |
| [>=100K] | 4498 | 4325 (96.2) | 173 ( 3.8) |
| *weighted* |  | (96.0) | ( 4.0) |
| **high.educ** |  |  |  |
| < HS Diploma | 402 | 382 (95.0) | 20 ( 5.0) |
| *weighted* |  | (94.4) | ( 5.6) |
| HS Diploma/GED | 891 | 853 (95.7) | 38 ( 4.3) |
| *weighted* |  | (96.0) | ( 4.0) |
| Some College | 2728 | 2605 (95.5) | 123 ( 4.5) |
| *weighted* |  | (95.0) | ( 5.0) |
| Bachelor | 2798 | 2670 (95.4) | 128 ( 4.6) |
| *weighted* |  | (94.9) | ( 5.1) |
| Post Graduate Degree | 3781 | 3640 (96.3) | 141 ( 3.7) |
| *weighted* |  | (95.9) | ( 4.1) |

*Supplementary Table 5. Contingency table for variable of interest: Age at first walking (months). Full sample.*

| **Demographics** | **n** | **Within guidelines** | **Beyond guidelines** |
| --- | --- | --- | --- |
| **sex_at_birth** |  |  |  |
| F | 5085 | 4958 (97.5) | 127 ( 2.5) |
| *weighted* |  | (97.3) | ( 2.7) |
| M | 5515 | 5302 (96.1) | 213 ( 3.9) |
| *weighted* |  | (96.0) | ( 4.0) |
| **race.4level** |  |  |  |
| White | 6998 | 6766 (96.7) | 232 ( 3.3) |
| *weighted* |  | (96.4) | ( 3.6) |
| Black | 1570 | 1515 (96.5) | 55 ( 3.5) |
| *weighted* |  | (97.0) | ( 3.0) |
| Asian | 235 | 228 (97.0) | <10 |
| *weighted* |  | (96.6) |  |
| Other/Mixed | 1797 | 1751 (97.4) | 46 ( 2.6) |
| *weighted* |  | (97.5) | ( 2.5) |
| **hisp** |  |  |  |
| No | 8593 | 8330 (96.9) | 263 ( 3.1) |
| *weighted* |  | (96.8) | ( 3.2) |
| Yes | 2007 | 1930 (96.2) | 77 ( 3.8) |
| *weighted* |  | (96.0) | ( 4.0) |
| **household.income** |  |  |  |
| [<50K] | 3088 | 2968 (96.1) | 120 ( 3.9) |
| *weighted* |  | (96.0) | ( 4.0) |
| [>=50K & <100K] | 3014 | 2928 (97.1) | 86 ( 2.9) |
| *weighted* |  | (97.0) | ( 3.0) |
| [>=100K] | 4498 | 4364 (97.0) | 134 ( 3.0) |
| *weighted* |  | (97.1) | ( 2.9) |
| **high.educ** |  |  |  |
| < HS Diploma | 402 | 383 (95.3) | 19 ( 4.7) |
| *weighted* |  | (95.1) | ( 4.9) |
| HS Diploma/GED | 891 | 864 (97.0) | 27 ( 3.0) |
| *weighted* |  | (97.4) | ( 2.6) |
| Some College | 2728 | 2631 (96.4) | 97 ( 3.6) |
| *weighted* |  | (96.1) | ( 3.9) |
| Bachelor | 2798 | 2714 (97.0) | 84 ( 3.0) |
| *weighted* |  | (96.9) | ( 3.1) |
| Post Graduate Degree | 3781 | 3668 (97.0) | 113 ( 3.0) |
| *weighted* |  | (97.0) | ( 3.0) |

*Supplementary Table 6. Contingency table for variable of interest: Age at first word (months). Full sample.*

| **Demographics** | **n** | **Within guidelines** | **Beyond guidelines** |
| --- | --- | --- | --- |
| **sex_at_birth** |  |  |  |
| F | 5085 | 4717 (92.8) | 368 ( 7.2) |
| *weighted* |  | (92.6) | ( 7.4) |
| M | 5515 | 4846 (87.9) | 669 (12.1) |
| *weighted* |  | (87.5) | (12.5) |
| **race.4level** |  |  |  |
| White | 6998 | 6273 (89.6) | 725 (10.4) |
| *weighted* |  | (89.4) | (10.6) |
| Black | 1570 | 1445 (92.0) | 125 ( 8.0) |
| *weighted* |  | (92.6) | ( 7.4) |
| Asian | 235 | 200 (85.1) | 35 (14.9) |
| *weighted* |  | (84.4) | (15.6) |
| Other/Mixed | 1797 | 1645 (91.5) | 152 ( 8.5) |
| *weighted* |  | (92.1) | ( 7.9) |
| **hisp** |  |  |  |
| No | 8593 | 7742 (90.1) | 851 ( 9.9) |
| *weighted* |  | (89.8) | (10.2) |
| Yes | 2007 | 1821 (90.7) | 186 ( 9.3) |
| *weighted* |  | (90.8) | ( 9.2) |
| **household.income** |  |  |  |
| [<50K] | 3088 | 2823 (91.4) | 265 ( 8.6) |
| *weighted* |  | (90.9) | ( 9.1) |
| [>=50K & <100K] | 3014 | 2708 (89.8) | 306 (10.2) |
| *weighted* |  | (89.7) | (10.3) |
| [>=100K] | 4498 | 4032 (89.6) | 466 (10.4) |
| *weighted* |  | (89.2) | (10.8) |
| **high.educ** |  |  |  |
| < HS Diploma | 402 | 376 (93.5) | 26 ( 6.5) |
| *weighted* |  | (93.7) | ( 6.3) |
| HS Diploma/GED | 891 | 818 (91.8) | 73 ( 8.2) |
| *weighted* |  | (91.9) | ( 8.1) |
| Some College | 2728 | 2472 (90.6) | 256 ( 9.4) |
| *weighted* |  | (90.2) | ( 9.8) |
| Bachelor | 2798 | 2520 (90.1) | 278 ( 9.9) |
| *weighted* |  | (89.6) | (10.4) |
| Post Graduate Degree | 3781 | 3377 (89.3) | 404 (10.7) |
| *weighted* |  | (89.0) | (11.0) |

*Supplementary Table 7. Contingency table for variable of interest: Total pregnancy problems. Full sample.*

| **Demographics** | **n** | **None** | **One problem** | **Two or more problems** |
| --- | --- | --- | --- | --- |
| **sex_at_birth** |  |  |  |  |
| F | 5085 | 2996 (58.9) | 1312 (25.8) | 777 (15.3) |
| *weighted* |  | (57.6) | (26.1) | (16.3) |
| M | 5515 | 3363 (61.0) | 1337 (24.2) | 815 (14.8) |
| *weighted* |  | (60.7) | (24.3) | (15.0) |
| **race.4level** |  |  |  |  |
| White | 6998 | 4286 (61.2) | 1745 (24.9) | 967 (13.8) |
| *weighted* |  | (59.9) | (25.2) | (14.9) |
| Black | 1570 | 869 (55.4) | 407 (25.9) | 294 (18.7) |
| *weighted* |  | (53.5) | (27.4) | (19.1) |
| Asian | 235 | 175 (74.5) | 47 (20.0) | 13 ( 5.5) |
| *weighted* |  | (76.6) | (18.0) | ( 5.4) |
| Other/Mixed | 1797 | 1029 (57.3) | 450 (25.0) | 318 (17.7) |
| *weighted* |  | (56.7) | (24.8) | (18.6) |
| **hisp** |  |  |  |  |
| No | 8593 | 5232 (60.9) | 2065 (24.0) | 1296 (15.1) |
| *weighted* |  | (60.3) | (24.1) | (15.5) |
| Yes | 2007 | 1127 (56.2) | 584 (29.1) | 296 (14.7) |
| *weighted* |  | (55.2) | (28.8) | (16.0) |
| **household.income** |  |  |  |  |
| [<50K] | 3088 | 1677 (54.3) | 811 (26.3) | 600 (19.4) |
| *weighted* |  | (54.8) | (26.1) | (19.1) |
| [>=50K & <100K] | 3014 | 1817 (60.3) | 759 (25.2) | 438 (14.5) |
| *weighted* |  | (60.2) | (25.4) | (14.5) |
| [>=100K] | 4498 | 2865 (63.7) | 1079 (24.0) | 554 (12.3) |
| *weighted* |  | (63.7) | (23.8) | (12.4) |
| **high.educ** |  |  |  |  |
| < HS Diploma | 402 | 218 (54.2) | 112 (27.9) | 72 (17.9) |
| *weighted* |  | (55.0) | (27.4) | (17.6) |
| HS Diploma/GED | 891 | 501 (56.2) | 220 (24.7) | 170 (19.1) |
| *weighted* |  | (55.8) | (24.4) | (19.7) |
| Some College | 2728 | 1445 (53.0) | 738 (27.1) | 545 (20.0) |
| *weighted* |  | (53.1) | (26.6) | (20.2) |
| Bachelor | 2798 | 1704 (60.9) | 703 (25.1) | 391 (14.0) |
| *weighted* |  | (60.5) | (25.5) | (14.0) |
| Post Graduate Degree | 3781 | 2491 (65.9) | 876 (23.2) | 414 (10.9) |
| *weighted* |  | (65.8) | (23.4) | (10.8) |

*Supplementary Table 8. Contingency table for variable of interest: Total birth problems. Full sample.*

| **Demographics** | **n** | **None** | **One problem** | **Two or more problems** |
| --- | --- | --- | --- | --- |
| **sex_at_birth** |  |  |  |  |
| F | 5085 | 3925 (77.2) | 849 (16.7) | 311 ( 6.1) |
| *weighted* |  | (77.6) | (16.2) | ( 6.2) |
| M | 5515 | 4065 (73.7) | 1044 (18.9) | 406 ( 7.4) |
| *weighted* |  | (74.2) | (18.7) | ( 7.1) |
| **race.4level** |  |  |  |  |
| White | 6998 | 5186 (74.1) | 1285 (18.4) | 527 ( 7.5) |
| *weighted* |  | (74.5) | (18.1) | ( 7.5) |
| Black | 1570 | 1225 (78.0) | 249 (15.9) | 96 ( 6.1) |
| *weighted* |  | (77.6) | (16.4) | ( 6.0) |
| Asian | 235 | 184 (78.3) | 47 (20.0) | <10 |
| *weighted* |  | (80.9) | (17.5) |  |
| Other/Mixed | 1797 | 1395 (77.6) | 312 (17.4) | 90 ( 5.0) |
| *weighted* |  | (79.8) | (15.4) | ( 4.7) |
| **hisp** |  |  |  |  |
| No | 8593 | 6397 (74.4) | 1583 (18.4) | 613 ( 7.1) |
| *weighted* |  | (74.7) | (18.1) | ( 7.2) |
| Yes | 2007 | 1593 (79.4) | 310 (15.4) | 104 ( 5.2) |
| *weighted* |  | (80.0) | (15.2) | ( 4.8) |
| **household.income** |  |  |  |  |
| [<50K] | 3088 | 2424 (78.5) | 478 (15.5) | 186 ( 6.0) |
| *weighted* |  | (78.5) | (15.4) | ( 6.1) |
| [>=50K & <100K] | 3014 | 2223 (73.8) | 582 (19.3) | 209 ( 6.9) |
| *weighted* |  | (73.7) | (19.3) | ( 7.0) |
| [>=100K] | 4498 | 3343 (74.3) | 833 (18.5) | 322 ( 7.2) |
| *weighted* |  | (74.7) | (18.1) | ( 7.1) |
| **high.educ** |  |  |  |  |
| < HS Diploma | 402 | 330 (82.1) | 52 (12.9) | 20 ( 5.0) |
| *weighted* |  | (81.0) | (14.5) | ( 4.5) |
| HS Diploma/GED | 891 | 718 (80.6) | 120 (13.5) | 53 ( 5.9) |
| *weighted* |  | (80.0) | (14.4) | ( 5.6) |
| Some College | 2728 | 2037 (74.7) | 481 (17.6) | 210 ( 7.7) |
| *weighted* |  | (75.3) | (17.0) | ( 7.7) |
| Bachelor | 2798 | 2026 (72.4) | 560 (20.0) | 212 ( 7.6) |
| *weighted* |  | (72.7) | (19.5) | ( 7.7) |
| Post Graduate Degree | 3781 | 2879 (76.1) | 680 (18.0) | 222 ( 5.9) |
| *weighted* |  | (76.8) | (17.7) | ( 5.5) |

*Supplementary Table 9. Contingency table for variable of interest: Prenatal alcohol exposure. Full sample.*

| **Demographics** | **n** | **Pre-No/Post-No** | **Pre-Yes/Post-No** | **Pre-Yes/Post-Yes** |
| --- | --- | --- | --- | --- |
| **sex_at_birth** |  |  |  |  |
| F | 4739 | 3455 (72.9) | 1156 (24.4) | 128 ( 2.7) |
| *weighted* |  | (74.1) | (23.4) | ( 2.5) |
| M | 5164 | 3811 (73.8) | 1227 (23.8) | 126 ( 2.4) |
| *weighted* |  | (74.9) | (22.7) | ( 2.4) |
| **race.4level** |  |  |  |  |
| White | 6582 | 4644 (70.6) | 1744 (26.5) | 194 ( 2.9) |
| *weighted* |  | (71.5) | (25.7) | ( 2.8) |
| Black | 1456 | 1203 (82.6) | 229 (15.7) | 24 ( 1.6) |
| *weighted* |  | (83.4) | (15.0) | ( 1.6) |
| Asian | 204 | 168 (82.4) | 35 (17.2) | <10 |
| *weighted* |  | (84.6) | (15.1) |  |
| Other/Mixed | 1661 | 1251 (75.3) | 375 (22.6) | 35 ( 2.1) |
| *weighted* |  | (78.1) | (19.7) | ( 2.2) |
| **hisp** |  |  |  |  |
| No | 8003 | 5784 (72.3) | 2004 (25.0) | 215 ( 2.7) |
| *weighted* |  | (72.9) | (24.5) | ( 2.6) |
| Yes | 1900 | 1482 (78.0) | 379 (19.9) | 39 ( 2.1) |
| *weighted* |  | (79.8) | (18.2) | ( 2.0) |
| **household.income** |  |  |  |  |
| [<50K] | 2924 | 2404 (82.2) | 467 (16.0) | 53 ( 1.8) |
| *weighted* |  | (81.6) | (16.6) | ( 1.8) |
| [>=50K & <100K] | 2799 | 2043 (73.0) | 697 (24.9) | 59 ( 2.1) |
| *weighted* |  | (72.3) | (25.4) | ( 2.3) |
| [>=100K] | 4180 | 2819 (67.4) | 1219 (29.2) | 142 ( 3.4) |
| *weighted* |  | (67.6) | (28.9) | ( 3.5) |
| **high.educ** |  |  |  |  |
| < HS Diploma | 386 | 355 (92.0) | 26 ( 6.7) | <10 |
| *weighted* |  | (92.4) | ( 6.4) |  |
| HS Diploma/GED | 850 | 721 (84.8) | 121 (14.2) | <10 |
| *weighted* |  | (83.7) | (15.4) |  |
| Some College | 2561 | 1922 (75.0) | 581 (22.7) | 58 ( 2.3) |
| *weighted* |  | (75.1) | (22.5) | ( 2.4) |
| Bachelor | 2618 | 1866 (71.3) | 688 (26.3) | 64 ( 2.4) |
| *weighted* |  | (72.3) | (25.2) | ( 2.4) |
| Post Graduate Degree | 3488 | 2402 (68.9) | 967 (27.7) | 119 ( 3.4) |
| *weighted* |  | (69.7) | (27.1) | ( 3.2) |

*Supplementary Table 10. Contingency table for variable of interest: Prenatal tobacco exposure. Full sample.*

| **Demographics** | **n** | **Pre-No/Post-No** | **Pre-Yes/Post-No** | **Pre-Yes/Post-Yes** |
| --- | --- | --- | --- | --- |
| **sex_at_birth** |  |  |  |  |
| F | 4920 | 4279 (87.0) | 394 ( 8.0) | 247 ( 5.0) |
| *weighted* |  | (84.1) | ( 9.2) | ( 6.7) |
| M | 5377 | 4646 (86.4) | 451 ( 8.4) | 280 ( 5.2) |
| *weighted* |  | (83.6) | ( 9.9) | ( 6.4) |
| **race.4level** |  |  |  |  |
| White | 6887 | 6096 (88.5) | 488 ( 7.1) | 303 ( 4.4) |
| *weighted* |  | (84.7) | ( 8.9) | ( 6.3) |
| Black | 1481 | 1202 (81.2) | 161 (10.9) | 118 ( 8.0) |
| *weighted* |  | (80.0) | (11.4) | ( 8.6) |
| Asian | 207 | 195 (94.2) | <10 | <10 |
| *weighted* |  | (94.3) |  |  |
| Other/Mixed | 1722 | 1432 (83.2) | 187 (10.9) | 103 ( 6.0) |
| *weighted* |  | (80.6) | (12.7) | ( 6.7) |
| **hisp** |  |  |  |  |
| No | 8347 | 7210 (86.4) | 677 ( 8.1) | 460 ( 5.5) |
| *weighted* |  | (83.1) | ( 9.5) | ( 7.4) |
| Yes | 1950 | 1715 (87.9) | 168 ( 8.6) | 67 ( 3.4) |
| *weighted* |  | (86.4) | ( 9.7) | ( 3.8) |
| **household.income** |  |  |  |  |
| [<50K] | 2994 | 2283 (76.3) | 389 (13.0) | 322 (10.8) |
| *weighted* |  | (74.9) | (13.4) | (11.7) |
| [>=50K & <100K] | 2922 | 2515 (86.1) | 263 ( 9.0) | 144 ( 4.9) |
| *weighted* |  | (84.9) | ( 9.8) | ( 5.3) |
| [>=100K] | 4381 | 4127 (94.2) | 193 ( 4.4) | 61 ( 1.4) |
| *weighted* |  | (94.1) | ( 4.6) | ( 1.3) |
| **high.educ** |  |  |  |  |
| < HS Diploma | 388 | 301 (77.6) | 50 (12.9) | 37 ( 9.5) |
| *weighted* |  | (77.3) | (11.6) | (11.1) |
| HS Diploma/GED | 865 | 640 (74.0) | 101 (11.7) | 124 (14.3) |
| *weighted* |  | (71.4) | (12.5) | (16.0) |
| Some College | 2649 | 2009 (75.8) | 393 (14.8) | 247 ( 9.3) |
| *weighted* |  | (73.5) | (16.0) | (10.5) |
| Bachelor | 2726 | 2491 (91.4) | 158 ( 5.8) | 77 ( 2.8) |
| *weighted* |  | (89.7) | ( 6.8) | ( 3.6) |
| Post Graduate Degree | 3669 | 3484 (95.0) | 143 ( 3.9) | 42 ( 1.1) |
| *weighted* |  | (94.5) | ( 4.2) | ( 1.3) |

*Supplementary Table 11. Contingency table for variable of interest: Prenatal marijuana exposure. Full sample.*

| **Demographics** | **n** | **Pre-No/Post-No** | **Pre-Yes/Post-No** | **Pre-Yes/Post-Yes** |
| --- | --- | --- | --- | --- |
| **sex_at_birth** |  |  |  |  |
| F | 4909 | 4620 (94.1) | 176 ( 3.6) | 113 ( 2.3) |
| *weighted* |  | (93.4) | ( 3.9) | ( 2.8) |
| M | 5341 | 5049 (94.5) | 193 ( 3.6) | 99 ( 1.9) |
| *weighted* |  | (93.8) | ( 4.0) | ( 2.1) |
| **race.4level** |  |  |  |  |
| White | 6858 | 6567 (95.8) | 181 ( 2.6) | 110 ( 1.6) |
| *weighted* |  | (94.8) | ( 3.1) | ( 2.2) |
| Black | 1478 | 1292 (87.4) | 115 ( 7.8) | 71 ( 4.8) |
| *weighted* |  | (87.0) | ( 8.0) | ( 5.0) |
| Asian | 207 | 204 (98.6) | <10 | <10 |
| *weighted* |  | (98.5) |  |  |
| Other/Mixed | 1707 | 1606 (94.1) | 71 ( 4.2) | 30 ( 1.8) |
| *weighted* |  | (93.0) | ( 5.2) | ( 1.8) |
| **hisp** |  |  |  |  |
| No | 8308 | 7837 (94.3) | 289 ( 3.5) | 182 ( 2.2) |
| *weighted* |  | (93.5) | ( 3.8) | ( 2.7) |
| Yes | 1942 | 1832 (94.3) | 80 ( 4.1) | 30 ( 1.5) |
| *weighted* |  | (93.9) | ( 4.5) | ( 1.6) |
| **household.income** |  |  |  |  |
| [<50K] | 2968 | 2637 (88.8) | 202 ( 6.8) | 129 ( 4.3) |
| *weighted* |  | (89.3) | ( 6.3) | ( 4.4) |
| [>=50K & <100K] | 2909 | 2760 (94.9) | 88 ( 3.0) | 61 ( 2.1) |
| *weighted* |  | (94.8) | ( 3.1) | ( 2.0) |
| [>=100K] | 4373 | 4272 (97.7) | 79 ( 1.8) | 22 ( 0.5) |
| *weighted* |  | (97.8) | ( 1.8) | ( 0.4) |
| **high.educ** |  |  |  |  |
| < HS Diploma | 385 | 364 (94.5) | 13 ( 3.4) | <10 |
| *weighted* |  | (94.0) | ( 3.0) |  |
| HS Diploma/GED | 855 | 755 (88.3) | 62 ( 7.3) | 38 ( 4.4) |
| *weighted* |  | (87.8) | ( 7.8) | ( 4.4) |
| Some College | 2633 | 2364 (89.8) | 160 ( 6.1) | 109 ( 4.1) |
| *weighted* |  | (89.9) | ( 5.8) | ( 4.3) |
| Bachelor | 2724 | 2623 (96.3) | 66 ( 2.4) | 35 ( 1.3) |
| *weighted* |  | (95.7) | ( 2.7) | ( 1.6) |
| Post Graduate Degree | 3653 | 3563 (97.5) | 68 ( 1.9) | 22 ( 0.6) |
| *weighted* |  | (97.4) | ( 2.0) | ( 0.6) |

*Supplementary Table 12. Contingency table for variable of interest: Prenatal other substance exposure. Full sample.*

| **Demographics** | **n** | **Pre-No/Post-No** | **Pre-Yes/Post-No** | **Pre-Yes/Post-Yes** |
| --- | --- | --- | --- | --- |
| **sex_at_birth** |  |  |  |  |
| F | 5075 | 4978 (98.1) | 66 ( 1.3) | 31 ( 0.6) |
| *weighted* |  | (97.7) | ( 1.6) | ( 0.6) |
| M | 5508 | 5403 (98.1) | 62 ( 1.1) | 43 ( 0.8) |
| *weighted* |  | (97.9) | ( 1.3) | ( 0.9) |
| **race.4level** |  |  |  |  |
| White | 6987 | 6878 (98.4) | 72 ( 1.0) | 37 ( 0.5) |
| *weighted* |  | (98.0) | ( 1.3) | ( 0.7) |
| Black | 1565 | 1535 (98.1) | 10 ( 0.6) | 20 ( 1.3) |
| *weighted* |  | (98.2) | ( 0.6) | ( 1.2) |
| Asian | 235 | 232 (98.7) | <10 | <10 |
| *weighted* |  | (98.6) |  |  |
| Other/Mixed | 1796 | 1736 (96.7) | 44 ( 2.4) | 16 ( 0.9) |
| *weighted* |  | (96.1) | ( 3.1) | ( 0.8) |
| **hisp** |  |  |  |  |
| No | 8578 | 8427 (98.2) | 93 ( 1.1) | 58 ( 0.7) |
| *weighted* |  | (98.0) | ( 1.3) | ( 0.7) |
| Yes | 2005 | 1954 (97.5) | 35 ( 1.7) | 16 ( 0.8) |
| *weighted* |  | (97.3) | ( 1.8) | ( 0.9) |
| **household.income** |  |  |  |  |
| [<50K] | 3085 | 2992 (97.0) | 63 ( 2.0) | 30 ( 1.0) |
| *weighted* |  | (96.7) | ( 2.3) | ( 1.0) |
| [>=50K & <100K] | 3006 | 2948 (98.1) | 37 ( 1.2) | 21 ( 0.7) |
| *weighted* |  | (98.0) | ( 1.3) | ( 0.7) |
| [>=100K] | 4492 | 4441 (98.9) | 28 ( 0.6) | 23 ( 0.5) |
| *weighted* |  | (99.1) | ( 0.5) | ( 0.4) |
| **high.educ** |  |  |  |  |
| < HS Diploma | 402 | 395 (98.3) | <10 | <10 |
| *weighted* |  | (97.8) |  |  |
| HS Diploma/GED | 890 | 866 (97.3) | 17 ( 1.9) | <10 |
| *weighted* |  | (96.9) | ( 2.4) |  |
| Some College | 2723 | 2629 (96.5) | 64 ( 2.4) | 30 ( 1.1) |
| *weighted* |  | (96.1) | ( 2.6) | ( 1.3) |
| Bachelor | 2790 | 2758 (98.9) | 21 ( 0.8) | 11 ( 0.4) |
| *weighted* |  | (98.8) | ( 0.9) | ( 0.3) |
| Post Graduate Degree | 3778 | 3733 (98.8) | 23 ( 0.6) | 22 ( 0.6) |
| *weighted* |  | (99.0) | ( 0.5) | ( 0.5) |

*Supplementary Table 13. Contingency table for variable of interest: Average hours of sleep per night. Full sample.*

| **Demographics** | **n** | **Less than 7 hours** | **7-8 hours** | **8-9 hours** | **9-11 hours** |
| --- | --- | --- | --- | --- | --- |
| **age_quartiles** |  |  |  |  |  |
| age<=25% | 2698 | 73 ( 2.7) | 291 (10.8) | 868 (32.2) | 1466 (54.3) |
| *weighted* |  | ( 2.9) | (12.1) | (34.5) | (50.4) |
| age>25&<=50% | 2704 | 69 ( 2.6) | 285 (10.5) | 952 (35.2) | 1398 (51.7) |
| *weighted* |  | ( 2.7) | (11.9) | (36.4) | (49.0) |
| age>50&<=75% | 2306 | 81 ( 3.5) | 248 (10.8) | 873 (37.9) | 1104 (47.9) |
| *weighted* |  | ( 3.8) | (12.0) | (39.8) | (44.4) |
| age>75% | 2536 | 99 ( 3.9) | 321 (12.7) | 1015 (40.0) | 1101 (43.4) |
| *weighted* |  | ( 4.4) | (14.1) | (41.9) | (39.5) |
| **sex_at_birth** |  |  |  |  |  |
| F | 4916 | 143 ( 2.9) | 512 (10.4) | 1835 (37.3) | 2426 (49.3) |
| *weighted* |  | ( 3.4) | (11.9) | (39.0) | (45.7) |
| M | 5328 | 179 ( 3.4) | 633 (11.9) | 1873 (35.2) | 2643 (49.6) |
| *weighted* |  | ( 3.6) | (13.1) | (37.4) | (45.9) |
| **race.4level** |  |  |  |  |  |
| White | 6849 | 102 ( 1.5) | 549 ( 8.0) | 2338 (34.1) | 3860 (56.4) |
| *weighted* |  | ( 1.9) | ( 9.8) | (36.6) | (51.7) |
| Black | 1449 | 156 (10.8) | 349 (24.1) | 596 (41.1) | 348 (24.0) |
| *weighted* |  | (11.6) | (24.9) | (41.1) | (22.4) |
| Asian | 221 | 2 ( 0.9) | 14 ( 6.3) | 84 (38.0) | 121 (54.8) |
| *weighted* |  | ( 1.1) | ( 7.1) | (40.4) | (51.4) |
| Other/Mixed | 1725 | 62 ( 3.6) | 233 (13.5) | 690 (40.0) | 740 (42.9) |
| *weighted* |  | ( 4.2) | (16.2) | (42.6) | (36.9) |
| **hisp** |  |  |  |  |  |
| No | 8329 | 265 ( 3.2) | 854 (10.3) | 2854 (34.3) | 4356 (52.3) |
| *weighted* |  | ( 3.4) | (11.3) | (36.0) | (49.3) |
| Yes | 1915 | 57 ( 3.0) | 291 (15.2) | 854 (44.6) | 713 (37.2) |
| *weighted* |  | ( 3.6) | (16.9) | (45.9) | (33.6) |
| **household.income** |  |  |  |  |  |
| [<50K] | 2879 | 200 ( 6.9) | 573 (19.9) | 1234 (42.9) | 872 (30.3) |
| *weighted* |  | ( 6.1) | (19.6) | (43.6) | (30.6) |
| [>=50K & <100K] | 2933 | 78 ( 2.7) | 311 (10.6) | 1114 (38.0) | 1430 (48.8) |
| *weighted* |  | ( 2.8) | (10.8) | (38.6) | (47.7) |
| [>=100K] | 4432 | 44 ( 1.0) | 261 ( 5.9) | 1360 (30.7) | 2767 (62.4) |
| *weighted* |  | ( 0.9) | ( 5.7) | (31.0) | (62.3) |
| **high.educ** |  |  |  |  |  |
| < HS Diploma | 353 | 29 ( 8.2) | 79 (22.4) | 162 (45.9) | 83 (23.5) |
| *weighted* |  | ( 8.3) | (22.3) | (47.4) | (22.0) |
| HS Diploma/GED | 810 | 58 ( 7.2) | 167 (20.6) | 373 (46.0) | 212 (26.2) |
| *weighted* |  | ( 6.4) | (21.2) | (44.9) | (27.6) |
| Some College | 2621 | 146 ( 5.6) | 456 (17.4) | 1122 (42.8) | 897 (34.2) |
| *weighted* |  | ( 5.3) | (17.9) | (43.5) | (33.3) |
| Bachelor | 2745 | 53 ( 1.9) | 215 ( 7.8) | 953 (34.7) | 1524 (55.5) |
| *weighted* |  | ( 2.3) | ( 8.8) | (36.3) | (52.6) |
| Post Graduate Degree | 3715 | 36 ( 1.0) | 228 ( 6.1) | 1098 (29.6) | 2353 (63.3) |
| *weighted* |  | ( 1.1) | ( 6.4) | (31.1) | (61.4) |
| **pubertal_dev_collapsed** |  |  |  |  |  |
| Pre | 5309 | 106 ( 2.0) | 476 ( 9.0) | 1760 (33.2) | 2967 (55.9) |
| *weighted* |  | ( 2.3) | (10.1) | (35.4) | (52.3) |
| Early | 2424 | 80 ( 3.3) | 282 (11.6) | 891 (36.8) | 1171 (48.3) |
| *weighted* |  | ( 3.6) | (12.8) | (38.7) | (45.0) |
| Mid | 2352 | 132 ( 5.6) | 355 (15.1) | 976 (41.5) | 889 (37.8) |
| *weighted* |  | ( 5.8) | (16.6) | (42.1) | (35.5) |
| Late/Post | 159 | <10 | 32 (20.1) | 81 (50.9) | 42 (26.4) |
| *weighted* |  |  | (21.0) | (53.3) | (23.6) |

*Supplementary Table 14. Contingency table for variable of interest: Total sleep disturbance. Full sample.*

| **Demographics** | **n** | **Low Sleep Disturbance** | **High Sleep Disturbance** |
| --- | --- | --- | --- |
| **age_quartiles** |  |  |  |
| age<=25% | 2698 | 1829 (67.8) | 869 (32.2) |
| *weighted* |  | (66.2) | (33.8) |
| age>25&<=50% | 2704 | 1903 (70.4) | 801 (29.6) |
| *weighted* |  | (69.4) | (30.6) |
| age>50&<=75% | 2306 | 1596 (69.2) | 710 (30.8) |
| *weighted* |  | (67.6) | (32.4) |
| age>75% | 2536 | 1787 (70.5) | 749 (29.5) |
| *weighted* |  | (69.1) | (30.9) |
| **sex_at_birth** |  |  |  |
| F | 4916 | 3458 (70.3) | 1458 (29.7) |
| *weighted* |  | (69.1) | (30.9) |
| M | 5328 | 3657 (68.6) | 1671 (31.4) |
| *weighted* |  | (67.1) | (32.9) |
| **race.4level** |  |  |  |
| White | 6849 | 4852 (70.8) | 1997 (29.2) |
| *weighted* |  | (69.0) | (31.0) |
| Black | 1449 | 968 (66.8) | 481 (33.2) |
| *weighted* |  | (66.5) | (33.5) |
| Asian | 221 | 160 (72.4) | 61 (27.6) |
| *weighted* |  | (70.3) | (29.7) |
| Other/Mixed | 1725 | 1135 (65.8) | 590 (34.2) |
| *weighted* |  | (64.1) | (35.9) |
| **hisp** |  |  |  |
| No | 8329 | 5781 (69.4) | 2548 (30.6) |
| *weighted* |  | (68.1) | (31.9) |
| Yes | 1915 | 1334 (69.7) | 581 (30.3) |
| *weighted* |  | (68.1) | (31.9) |
| **household.income** |  |  |  |
| [<50K] | 2879 | 1843 (64.0) | 1036 (36.0) |
| *weighted* |  | (63.5) | (36.5) |
| [>=50K & <100K] | 2933 | 1989 (67.8) | 944 (32.2) |
| *weighted* |  | (67.3) | (32.7) |
| [>=100K] | 4432 | 3283 (74.1) | 1149 (25.9) |
| *weighted* |  | (74.4) | (25.6) |
| **high.educ** |  |  |  |
| < HS Diploma | 353 | 235 (66.6) | 118 (33.4) |
| *weighted* |  | (64.5) | (35.5) |
| HS Diploma/GED | 810 | 531 (65.6) | 279 (34.4) |
| *weighted* |  | (65.0) | (35.0) |
| Some College | 2621 | 1711 (65.3) | 910 (34.7) |
| *weighted* |  | (64.1) | (35.9) |
| Bachelor | 2745 | 1921 (70.0) | 824 (30.0) |
| *weighted* |  | (69.0) | (31.0) |
| Post Graduate Degree | 3715 | 2717 (73.1) | 998 (26.9) |
| *weighted* |  | (72.7) | (27.3) |
| **pubertal_dev_collapsed** |  |  |  |
| Pre | 5309 | 3843 (72.4) | 1466 (27.6) |
| *weighted* |  | (71.3) | (28.7) |
| Early | 2424 | 1621 (66.9) | 803 (33.1) |
| *weighted* |  | (65.9) | (34.1) |
| Mid | 2352 | 1548 (65.8) | 804 (34.2) |
| *weighted* |  | (64.2) | (35.8) |
| Late/Post | 159 | 103 (64.8) | 56 (35.2) |
| *weighted* |  | (62.6) | (37.4) |

*Supplementary Table 15. Contingency table for variable of interest: Physical activity (vigorous). Full sample.*

| **Demographics** | **n** | **Below guidelines** | **Within guidelines** |
| --- | --- | --- | --- |
| **age_quartiles** |  |  |  |
| age<=25% | 2698 | 2249 (83.4) | 449 (16.6) |
| *weighted* |  | (83.8) | (16.2) |
| age>25&<=50% | 2704 | 2255 (83.4) | 449 (16.6) |
| *weighted* |  | (83.7) | (16.3) |
| age>50&<=75% | 2306 | 1916 (83.1) | 390 (16.9) |
| *weighted* |  | (83.4) | (16.6) |
| age>75% | 2536 | 2087 (82.3) | 449 (17.7) |
| *weighted* |  | (82.2) | (17.8) |
| **sex_at_birth** |  |  |  |
| F | 4916 | 4156 (84.5) | 760 (15.5) |
| *weighted* |  | (84.6) | (15.4) |
| M | 5328 | 4351 (81.7) | 977 (18.3) |
| *weighted* |  | (82.0) | (18.0) |
| **race.4level** |  |  |  |
| White | 6849 | 5630 (82.2) | 1219 (17.8) |
| *weighted* |  | (82.5) | (17.5) |
| Black | 1449 | 1229 (84.8) | 220 (15.2) |
| *weighted* |  | (85.1) | (14.9) |
| Asian | 221 | 189 (85.5) | 32 (14.5) |
| *weighted* |  | (84.2) | (15.8) |
| Other/Mixed | 1725 | 1459 (84.6) | 266 (15.4) |
| *weighted* |  | (85.0) | (15.0) |
| **hisp** |  |  |  |
| No | 8329 | 6850 (82.2) | 1479 (17.8) |
| *weighted* |  | (82.3) | (17.7) |
| Yes | 1915 | 1657 (86.5) | 258 (13.5) |
| *weighted* |  | (86.5) | (13.5) |
| **household.income** |  |  |  |
| [<50K] | 2879 | 2449 (85.1) | 430 (14.9) |
| *weighted* |  | (84.6) | (15.4) |
| [>=50K & <100K] | 2933 | 2421 (82.5) | 512 (17.5) |
| *weighted* |  | (82.5) | (17.5) |
| [>=100K] | 4432 | 3637 (82.1) | 795 (17.9) |
| *weighted* |  | (82.4) | (17.6) |
| **high.educ** |  |  |  |
| < HS Diploma | 353 | 309 (87.5) | 44 (12.5) |
| *weighted* |  | (88.3) | (11.7) |
| HS Diploma/GED | 810 | 704 (86.9) | 106 (13.1) |
| *weighted* |  | (86.7) | (13.3) |
| Some College | 2621 | 2199 (83.9) | 422 (16.1) |
| *weighted* |  | (83.5) | (16.5) |
| Bachelor | 2745 | 2265 (82.5) | 480 (17.5) |
| *weighted* |  | (82.4) | (17.6) |
| Post Graduate Degree | 3715 | 3030 (81.6) | 685 (18.4) |
| *weighted* |  | (81.9) | (18.1) |
| **pubertal_dev_collapsed** |  |  |  |
| Pre | 5309 | 4341 (81.8) | 968 (18.2) |
| *weighted* |  | (81.8) | (18.2) |
| Early | 2424 | 2019 (83.3) | 405 (16.7) |
| *weighted* |  | (83.8) | (16.2) |
| Mid | 2352 | 2006 (85.3) | 346 (14.7) |
| *weighted* |  | (85.2) | (14.8) |
| Late/Post | 159 | 141 (88.7) | 18 (11.3) |
| *weighted* |  | (89.8) | (10.2) |

*Supplementary Table 16. Contingency table for variable of interest: Physical activity (strengthening). Full sample.*

| **Demographics** | **n** | **Below guidelines** | **Within guidelines** |
| --- | --- | --- | --- |
| **age_quartiles** |  |  |  |
| age<=25% | 2698 | 1909 (70.8) | 789 (29.2) |
| *weighted* |  | (70.9) | (29.1) |
| age>25&<=50% | 2704 | 1892 (70.0) | 812 (30.0) |
| *weighted* |  | (69.8) | (30.2) |
| age>50&<=75% | 2306 | 1565 (67.9) | 741 (32.1) |
| *weighted* |  | (68.0) | (32.0) |
| age>75% | 2536 | 1706 (67.3) | 830 (32.7) |
| *weighted* |  | (67.2) | (32.8) |
| **sex_at_birth** |  |  |  |
| F | 4916 | 3502 (71.2) | 1414 (28.8) |
| *weighted* |  | (71.0) | (29.0) |
| M | 5328 | 3570 (67.0) | 1758 (33.0) |
| *weighted* |  | (67.0) | (33.0) |
| **race.4level** |  |  |  |
| White | 6849 | 4862 (71.0) | 1987 (29.0) |
| *weighted* |  | (70.8) | (29.2) |
| Black | 1449 | 857 (59.1) | 592 (40.9) |
| *weighted* |  | (59.2) | (40.8) |
| Asian | 221 | 156 (70.6) | 65 (29.4) |
| *weighted* |  | (70.2) | (29.8) |
| Other/Mixed | 1725 | 1197 (69.4) | 528 (30.6) |
| *weighted* |  | (68.8) | (31.2) |
| **hisp** |  |  |  |
| No | 8329 | 5764 (69.2) | 2565 (30.8) |
| *weighted* |  | (69.6) | (30.4) |
| Yes | 1915 | 1308 (68.3) | 607 (31.7) |
| *weighted* |  | (67.0) | (33.0) |
| **household.income** |  |  |  |
| [<50K] | 2879 | 1878 (65.2) | 1001 (34.8) |
| *weighted* |  | (66.3) | (33.7) |
| [>=50K & <100K] | 2933 | 2047 (69.8) | 886 (30.2) |
| *weighted* |  | (70.2) | (29.8) |
| [>=100K] | 4432 | 3147 (71.0) | 1285 (29.0) |
| *weighted* |  | (71.0) | (29.0) |
| **high.educ** |  |  |  |
| < HS Diploma | 353 | 232 (65.7) | 121 (34.3) |
| *weighted* |  | (66.0) | (34.0) |
| HS Diploma/GED | 810 | 514 (63.5) | 296 (36.5) |
| *weighted* |  | (64.1) | (35.9) |
| Some College | 2621 | 1743 (66.5) | 878 (33.5) |
| *weighted* |  | (66.5) | (33.5) |
| Bachelor | 2745 | 1960 (71.4) | 785 (28.6) |
| *weighted* |  | (71.9) | (28.1) |
| Post Graduate Degree | 3715 | 2623 (70.6) | 1092 (29.4) |
| *weighted* |  | (70.9) | (29.1) |
| **pubertal_dev_collapsed** |  |  |  |
| Pre | 5309 | 3705 (69.8) | 1604 (30.2) |
| *weighted* |  | (69.8) | (30.2) |
| Early | 2424 | 1646 (67.9) | 778 (32.1) |
| *weighted* |  | (68.4) | (31.6) |
| Mid | 2352 | 1608 (68.4) | 744 (31.6) |
| *weighted* |  | (67.8) | (32.2) |
| Late/Post | 159 | 113 (71.1) | 46 (28.9) |
| *weighted* |  | (71.5) | (28.5) |

*Supplementary Table 17. Contingency table for variable of interest: Sports/activities involvement. Full sample.*

| **Demographics** | **n** | **No Participation** | **Some Participation** |
| --- | --- | --- | --- |
| **age_quartiles** |  |  |  |
| age<=25% | 2698 | 665 (24.6) | 2033 (75.4) |
| *weighted* |  | (28.5) | (71.5) |
| age>25&<=50% | 2704 | 597 (22.1) | 2107 (77.9) |
| *weighted* |  | (25.2) | (74.8) |
| age>50&<=75% | 2306 | 499 (21.6) | 1807 (78.4) |
| *weighted* |  | (25.4) | (74.6) |
| age>75% | 2536 | 504 (19.9) | 2032 (80.1) |
| *weighted* |  | (23.0) | (77.0) |
| **sex_at_birth** |  |  |  |
| F | 4916 | 1179 (24.0) | 3737 (76.0) |
| *weighted* |  | (27.7) | (72.3) |
| M | 5328 | 1086 (20.4) | 4242 (79.6) |
| *weighted* |  | (23.5) | (76.5) |
| **race.4level** |  |  |  |
| White | 6849 | 1194 (17.4) | 5655 (82.6) |
| *weighted* |  | (21.3) | (78.7) |
| Black | 1449 | 574 (39.6) | 875 (60.4) |
| *weighted* |  | (43.1) | (56.9) |
| Asian | 221 | 36 (16.3) | 185 (83.7) |
| *weighted* |  | (18.1) | (81.9) |
| Other/Mixed | 1725 | 461 (26.7) | 1264 (73.3) |
| *weighted* |  | (32.5) | (67.5) |
| **hisp** |  |  |  |
| No | 8329 | 1690 (20.3) | 6639 (79.7) |
| *weighted* |  | (23.1) | (76.9) |
| Yes | 1915 | 575 (30.0) | 1340 (70.0) |
| *weighted* |  | (34.2) | (65.8) |
| **household.income** |  |  |  |
| [<50K] | 2879 | 1190 (41.3) | 1689 (58.7) |
| *weighted* |  | (40.8) | (59.2) |
| [>=50K & <100K] | 2933 | 670 (22.8) | 2263 (77.2) |
| *weighted* |  | (23.4) | (76.6) |
| [>=100K] | 4432 | 405 ( 9.1) | 4027 (90.9) |
| *weighted* |  | ( 9.3) | (90.7) |
| **high.educ** |  |  |  |
| < HS Diploma | 353 | 183 (51.8) | 170 (48.2) |
| *weighted* |  | (53.3) | (46.7) |
| HS Diploma/GED | 810 | 370 (45.7) | 440 (54.3) |
| *weighted* |  | (45.8) | (54.2) |
| Some College | 2621 | 908 (34.6) | 1713 (65.4) |
| *weighted* |  | (35.8) | (64.2) |
| Bachelor | 2745 | 463 (16.9) | 2282 (83.1) |
| *weighted* |  | (18.9) | (81.1) |
| Post Graduate Degree | 3715 | 341 ( 9.2) | 3374 (90.8) |
| *weighted* |  | (10.9) | (89.1) |
| **pubertal_dev_collapsed** |  |  |  |
| Pre | 5309 | 1000 (18.8) | 4309 (81.2) |
| *weighted* |  | (22.1) | (77.9) |
| Early | 2424 | 513 (21.2) | 1911 (78.8) |
| *weighted* |  | (24.6) | (75.4) |
| Mid | 2352 | 702 (29.8) | 1650 (70.2) |
| *weighted* |  | (32.7) | (67.3) |
| Late/Post | 159 | 50 (31.4) | 109 (68.6) |
| *weighted* |  | (31.4) | (68.6) |

*Supplementary Table 18. Contingency table for variable of interest: Weight status (from BMI). Full sample.*

| **Demographics** | **n** | **Underweight** | **Healthy Weight** | **Overweight** | **Obese** |
| --- | --- | --- | --- | --- | --- |
| **age_quartiles** |  |  |  |  |  |
| age<=25% | 2691 | 92 ( 3.4) | 1769 (65.7) | 376 (14.0) | 454 (16.9) |
| *weighted* |  | ( 3.2) | (63.8) | (14.3) | (18.7) |
| age>25&<=50% | 2696 | 104 ( 3.9) | 1769 (65.6) | 403 (14.9) | 420 (15.6) |
| *weighted* |  | ( 3.7) | (64.0) | (15.7) | (16.6) |
| age>50&<=75% | 2296 | 93 ( 4.1) | 1499 (65.3) | 335 (14.6) | 369 (16.1) |
| *weighted* |  | ( 4.0) | (63.5) | (15.2) | (17.4) |
| age>75% | 2531 | 105 ( 4.1) | 1669 (65.9) | 379 (15.0) | 378 (14.9) |
| *weighted* |  | ( 3.9) | (63.9) | (15.8) | (16.4) |
| **sex_at_birth** |  |  |  |  |  |
| F | 4903 | 207 ( 4.2) | 3241 (66.1) | 710 (14.5) | 745 (15.2) |
| *weighted* |  | ( 4.0) | (64.5) | (15.4) | (16.2) |
| M | 5311 | 187 ( 3.5) | 3465 (65.2) | 783 (14.7) | 876 (16.5) |
| *weighted* |  | ( 3.4) | (63.2) | (15.1) | (18.3) |
| **race.4level** |  |  |  |  |  |
| White | 6833 | 286 ( 4.2) | 4778 (69.9) | 936 (13.7) | 833 (12.2) |
| *weighted* |  | ( 4.0) | (67.4) | (14.5) | (14.2) |
| Black | 1440 | 35 ( 2.4) | 728 (50.6) | 251 (17.4) | 426 (29.6) |
| *weighted* |  | ( 2.3) | (50.2) | (17.6) | (29.9) |
| Asian | 220 | 14 ( 6.4) | 157 (71.4) | 26 (11.8) | 23 (10.5) |
| *weighted* |  | ( 5.5) | (69.8) | (13.5) | (11.2) |
| Other/Mixed | 1721 | 59 ( 3.4) | 1043 (60.6) | 280 (16.3) | 339 (19.7) |
| *weighted* |  | ( 3.1) | (57.0) | (17.3) | (22.7) |
| **hisp** |  |  |  |  |  |
| No | 8302 | 344 ( 4.1) | 5690 (68.5) | 1121 (13.5) | 1147 (13.8) |
| *weighted* |  | ( 4.0) | (67.1) | (14.0) | (14.9) |
| Yes | 1912 | 50 ( 2.6) | 1016 (53.1) | 372 (19.5) | 474 (24.8) |
| *weighted* |  | ( 2.6) | (52.1) | (19.7) | (25.6) |
| **household.income** |  |  |  |  |  |
| [<50K] | 2869 | 71 ( 2.5) | 1536 (53.5) | 506 (17.6) | 756 (26.4) |
| *weighted* |  | ( 2.6) | (54.5) | (17.7) | (25.2) |
| [>=50K & <100K] | 2925 | 121 ( 4.1) | 1886 (64.5) | 450 (15.4) | 468 (16.0) |
| *weighted* |  | ( 4.2) | (64.7) | (15.2) | (15.9) |
| [>=100K] | 4420 | 202 ( 4.6) | 3284 (74.3) | 537 (12.1) | 397 ( 9.0) |
| *weighted* |  | ( 4.5) | (74.2) | (12.2) | ( 9.0) |
| **high.educ** |  |  |  |  |  |
| < HS Diploma | 352 | <10 | 159 (45.2) | 74 (21.0) | 111 (31.5) |
| *weighted* |  |  | (44.8) | (22.3) | (30.4) |
| HS Diploma/GED | 810 | 21 ( 2.6) | 402 (49.6) | 124 (15.3) | 263 (32.5) |
| *weighted* |  | ( 2.8) | (48.9) | (15.2) | (33.1) |
| Some College | 2609 | 78 ( 3.0) | 1481 (56.8) | 455 (17.4) | 595 (22.8) |
| *weighted* |  | ( 2.9) | (57.3) | (17.4) | (22.4) |
| Bachelor | 2737 | 115 ( 4.2) | 1916 (70.0) | 388 (14.2) | 318 (11.6) |
| *weighted* |  | ( 4.2) | (68.3) | (14.9) | (12.6) |
| Post Graduate Degree | 3706 | 172 ( 4.6) | 2748 (74.2) | 452 (12.2) | 334 ( 9.0) |
| *weighted* |  | ( 4.5) | (73.7) | (12.4) | ( 9.4) |
| **pubertal_dev_collapsed** |  |  |  |  |  |
| Pre | 5296 | 270 ( 5.1) | 3832 (72.4) | 615 (11.6) | 579 (10.9) |
| *weighted* |  | ( 5.0) | (70.5) | (12.1) | (12.4) |
| Early | 2416 | 81 ( 3.4) | 1536 (63.6) | 387 (16.0) | 412 (17.1) |
| *weighted* |  | ( 3.3) | (62.7) | (16.0) | (18.1) |
| Mid | 2344 | 43 ( 1.8) | 1282 (54.7) | 441 (18.8) | 578 (24.7) |
| *weighted* |  | ( 1.8) | (53.7) | (19.2) | (25.3) |
| Late/Post | 158 | 0 ( 0.0) | 56 (35.4) | 50 (31.6) | 52 (32.9) |
| *weighted* |  | ( 0.0) | (36.3) | (34.4) | (29.2) |

*Supplementary Table 19. Contingency table for variable of interest: Total medical problems. Full sample.*

| **Demographics** | **n** | **None** | **One problem** | **Two or more problems** |
| --- | --- | --- | --- | --- |
| **age_quartiles** |  |  |  |  |
| age<=25% | 2687 | 814 (30.3) | 831 (30.9) | 1042 (38.8) |
| *weighted* |  | (29.5) | (30.5) | (40.0) |
| age>25&<=50% | 2697 | 825 (30.6) | 792 (29.4) | 1080 (40.0) |
| *weighted* |  | (30.6) | (28.6) | (40.8) |
| age>50&<=75% | 2292 | 668 (29.1) | 726 (31.7) | 898 (39.2) |
| *weighted* |  | (28.2) | (30.7) | (41.1) |
| age>75% | 2530 | 736 (29.1) | 722 (28.5) | 1072 (42.4) |
| *weighted* |  | (29.0) | (27.5) | (43.5) |
| **sex_at_birth** |  |  |  |  |
| F | 4900 | 1533 (31.3) | 1499 (30.6) | 1868 (38.1) |
| *weighted* |  | (30.4) | (29.8) | (39.8) |
| M | 5306 | 1510 (28.5) | 1572 (29.6) | 2224 (41.9) |
| *weighted* |  | (28.3) | (28.9) | (42.9) |
| **race.4level** |  |  |  |  |
| White | 6824 | 1944 (28.5) | 2081 (30.5) | 2799 (41.0) |
| *weighted* |  | (27.7) | (29.6) | (42.7) |
| Black | 1443 | 481 (33.3) | 419 (29.0) | 543 (37.6) |
| *weighted* |  | (33.8) | (28.7) | (37.5) |
| Asian | 221 | 73 (33.0) | 61 (27.6) | 87 (39.4) |
| *weighted* |  | (32.8) | (27.2) | (40.0) |
| Other/Mixed | 1718 | 545 (31.7) | 510 (29.7) | 663 (38.6) |
| *weighted* |  | (32.3) | (28.9) | (38.7) |
| **hisp** |  |  |  |  |
| No | 8300 | 2405 (29.0) | 2527 (30.4) | 3368 (40.6) |
| *weighted* |  | (28.3) | (29.5) | (42.2) |
| Yes | 1906 | 638 (33.5) | 544 (28.5) | 724 (38.0) |
| *weighted* |  | (32.9) | (28.7) | (38.5) |
| **household.income** |  |  |  |  |
| [<50K] | 2859 | 957 (33.5) | 811 (28.4) | 1091 (38.2) |
| *weighted* |  | (32.0) | (27.8) | (40.2) |
| [>=50K & <100K] | 2926 | 851 (29.1) | 865 (29.6) | 1210 (41.4) |
| *weighted* |  | (28.2) | (29.0) | (42.8) |
| [>=100K] | 4421 | 1235 (27.9) | 1395 (31.6) | 1791 (40.5) |
| *weighted* |  | (27.2) | (31.5) | (41.3) |
| **high.educ** |  |  |  |  |
| < HS Diploma | 350 | 156 (44.6) | 90 (25.7) | 104 (29.7) |
| *weighted* |  | (44.1) | (26.1) | (29.9) |
| HS Diploma/GED | 807 | 312 (38.7) | 219 (27.1) | 276 (34.2) |
| *weighted* |  | (36.1) | (27.9) | (35.9) |
| Some College | 2609 | 797 (30.5) | 737 (28.2) | 1075 (41.2) |
| *weighted* |  | (29.3) | (27.7) | (43.0) |
| Bachelor | 2740 | 776 (28.3) | 870 (31.8) | 1094 (39.9) |
| *weighted* |  | (27.8) | (30.9) | (41.4) |
| Post Graduate Degree | 3700 | 1002 (27.1) | 1155 (31.2) | 1543 (41.7) |
| *weighted* |  | (26.5) | (30.5) | (43.1) |
| **pubertal_dev_collapsed** |  |  |  |  |
| Pre | 5289 | 1596 (30.2) | 1633 (30.9) | 2060 (38.9) |
| *weighted* |  | (29.9) | (30.4) | (39.7) |
| Early | 2415 | 668 (27.7) | 726 (30.1) | 1021 (42.3) |
| *weighted* |  | (27.0) | (28.8) | (44.2) |
| Mid | 2343 | 731 (31.2) | 667 (28.5) | 945 (40.3) |
| *weighted* |  | (30.4) | (27.9) | (41.7) |
| Late/Post | 159 | 48 (30.2) | 45 (28.3) | 66 (41.5) |
| *weighted* |  | (30.1) | (25.6) | (44.4) |

*Supplementary Table 20. Contingency table for variable of interest: Traumatic Brain Injury (TBI). Full sample.*

| **Demographics** | **n** | **Improbable TBI** | **Possible mild TBI** | **TBI** |
| --- | --- | --- | --- | --- |
| **age_quartiles** |  |  |  |  |
| age<=25% | 2698 | 2611 (96.8) | 54 ( 2.0) | 33 ( 1.2) |
| *weighted* |  | (96.8) | ( 2.0) | ( 1.2) |
| age>25&<=50% | 2704 | 2606 (96.4) | 71 ( 2.6) | 27 ( 1.0) |
| *weighted* |  | (96.6) | ( 2.4) | ( 1.0) |
| age>50&<=75% | 2306 | 2214 (96.0) | 70 ( 3.0) | 22 ( 1.0) |
| *weighted* |  | (96.4) | ( 2.7) | ( 0.9) |
| age>75% | 2536 | 2418 (95.3) | 82 ( 3.2) | 36 ( 1.4) |
| *weighted* |  | (95.7) | ( 3.0) | ( 1.3) |
| **sex_at_birth** |  |  |  |  |
| F | 4916 | 4762 (96.9) | 107 ( 2.2) | 47 ( 1.0) |
| *weighted* |  | (97.0) | ( 2.0) | ( 1.0) |
| M | 5328 | 5087 (95.5) | 170 ( 3.2) | 71 ( 1.3) |
| *weighted* |  | (95.7) | ( 3.0) | ( 1.2) |
| **race.4level** |  |  |  |  |
| White | 6849 | 6564 (95.8) | 204 ( 3.0) | 81 ( 1.2) |
| *weighted* |  | (96.0) | ( 2.8) | ( 1.2) |
| Black | 1449 | 1410 (97.3) | 27 ( 1.9) | 12 ( 0.8) |
| *weighted* |  | (97.6) | ( 1.7) | ( 0.7) |
| Asian | 221 | 215 (97.3) | <10 | <10 |
| *weighted* |  | (97.8) |  |  |
| Other/Mixed | 1725 | 1660 (96.2) | 41 ( 2.4) | 24 ( 1.4) |
| *weighted* |  | (96.8) | ( 2.0) | ( 1.2) |
| **hisp** |  |  |  |  |
| No | 8329 | 7993 (96.0) | 243 ( 2.9) | 93 ( 1.1) |
| *weighted* |  | (96.1) | ( 2.8) | ( 1.1) |
| Yes | 1915 | 1856 (96.9) | 34 ( 1.8) | 25 ( 1.3) |
| *weighted* |  | (97.2) | ( 1.7) | ( 1.2) |
| **household.income** |  |  |  |  |
| [<50K] | 2879 | 2797 (97.2) | 54 ( 1.9) | 28 ( 1.0) |
| *weighted* |  | (97.1) | ( 1.8) | ( 1.1) |
| [>=50K & <100K] | 2933 | 2820 (96.1) | 83 ( 2.8) | 30 ( 1.0) |
| *weighted* |  | (96.1) | ( 2.8) | ( 1.0) |
| [>=100K] | 4432 | 4232 (95.5) | 140 ( 3.2) | 60 ( 1.4) |
| *weighted* |  | (95.7) | ( 3.1) | ( 1.3) |
| **high.educ** |  |  |  |  |
| < HS Diploma | 353 | 342 (96.9) | <10 | <10 |
| *weighted* |  | (96.4) |  |  |
| HS Diploma/GED | 810 | 791 (97.7) | 14 ( 1.7) | <10 |
| *weighted* |  | (97.7) | ( 1.8) |  |
| Some College | 2621 | 2541 (96.9) | 54 ( 2.1) | 26 ( 1.0) |
| *weighted* |  | (96.9) | ( 2.0) | ( 1.1) |
| Bachelor | 2745 | 2629 (95.8) | 79 ( 2.9) | 37 ( 1.3) |
| *weighted* |  | (96.2) | ( 2.6) | ( 1.2) |
| Post Graduate Degree | 3715 | 3546 (95.5) | 123 ( 3.3) | 46 ( 1.2) |
| *weighted* |  | (95.6) | ( 3.3) | ( 1.2) |
| **pubertal_dev_collapsed** |  |  |  |  |
| Pre | 5309 | 5101 (96.1) | 149 ( 2.8) | 59 ( 1.1) |
| *weighted* |  | (96.3) | ( 2.7) | ( 1.0) |
| Early | 2424 | 2326 (96.0) | 63 ( 2.6) | 35 ( 1.4) |
| *weighted* |  | (96.2) | ( 2.3) | ( 1.5) |
| Mid | 2352 | 2267 (96.4) | 62 ( 2.6) | 23 ( 1.0) |
| *weighted* |  | (96.7) | ( 2.4) | ( 0.9) |
| Late/Post | 159 | 155 (97.5) | <10 | <10 |
| *weighted* |  | (97.4) |  |  |
