## Supplemental Tables 21-40 for "A Comprehensive Overview of the Physical Health of the Adolescent Brain Cognitive Development Study (ABCD) Cohort at Baseline"

*Supplementary Table 21. Contingency table for variable of interest: Birth weight. Independent sample used for statistics.*

| **Demographics** | **n** | **Very low** | **Low** | **Normal** | **High** |
| --- | --- | --- | --- | --- | --- |
| **sex_at_birth** |  |  |  |  |  |
| F | 4089 | 45 ( 1.1) | 507 (12.4) | 3249 (79.5) | 288 ( 7.0) |
| *weighted* |  | ( 1.3) | (12.8) | (79.1) | ( 6.7) |
| M | 4447 | 27 ( 0.6) | 449 (10.1) | 3465 (77.9) | 506 (11.4) |
| *weighted* |  | ( 0.7) | (10.0) | (77.8) | (11.5) |
| **race.4level** |  |  |  |  |  |
| White | 5659 | 47 ( 0.8) | 590 (10.4) | 4451 (78.7) | 571 (10.1) |
| *weighted* |  | ( 1.0) | (10.8) | (78.3) | ( 9.9) |
| Black | 1228 | 18 ( 1.5) | 194 (15.8) | 944 (76.9) | 72 ( 5.9) |
| *weighted* |  | ( 1.5) | (16.4) | (76.6) | ( 5.5) |
| Asian | 190 | 0 ( 0.0) | 24 (12.6) | 155 (81.6) | 11 ( 5.8) |
| *weighted* |  | ( 0.0) | (12.7) | (81.0) | ( 6.4) |
| Other/Mixed | 1459 | 7 ( 0.5) | 148 (10.1) | 1164 (79.8) | 140 ( 9.6) |
| *weighted* |  | ( 0.6) | ( 9.4) | (80.3) | ( 9.7) |
| **hisp** |  |  |  |  |  |
| No | 6855 | 49 ( 0.7) | 806 (11.8) | 5344 (78.0) | 656 ( 9.6) |
| *weighted* |  | ( 0.8) | (12.1) | (77.6) | ( 9.5) |
| Yes | 1681 | 23 ( 1.4) | 150 ( 8.9) | 1370 (81.5) | 138 ( 8.2) |
| *weighted* |  | ( 1.4) | ( 9.1) | (81.3) | ( 8.2) |
| **household.income** |  |  |  |  |  |
| [<50K] | 2519 | 29 ( 1.2) | 310 (12.3) | 1983 (78.7) | 197 ( 7.8) |
| *weighted* |  | ( 1.2) | (12.0) | (78.8) | ( 8.1) |
| [>=50K & <100K] | 2425 | 25 ( 1.0) | 248 (10.2) | 1906 (78.6) | 246 (10.1) |
| *weighted* |  | ( 1.2) | (10.5) | (78.3) | (10.0) |
| [>=100K] | 3592 | 18 ( 0.5) | 398 (11.1) | 2825 (78.6) | 351 ( 9.8) |
| *weighted* |  | ( 0.5) | (11.6) | (78.2) | ( 9.8) |
| **high.educ** |  |  |  |  |  |
| < HS Diploma | 318 | <10 | 34 (10.7) | 252 (79.2) | 28 ( 8.8) |
| *weighted* |  |  | ( 8.6) | (82.1) | ( 8.1) |
| HS Diploma/GED | 729 | <10 | 85 (11.7) | 586 (80.4) | 51 ( 7.0) |
| *weighted* |  |  | (11.7) | (80.3) | ( 6.9) |
| Some College | 2199 | 26 ( 1.2) | 278 (12.6) | 1689 (76.8) | 206 ( 9.4) |
| *weighted* |  | ( 1.3) | (12.6) | (76.9) | ( 9.2) |
| Bachelor | 2219 | 12 ( 0.5) | 262 (11.8) | 1738 (78.3) | 207 ( 9.3) |
| *weighted* |  | ( 0.6) | (12.3) | (77.3) | ( 9.8) |
| Post Graduate Degree | 3071 | 23 ( 0.7) | 297 ( 9.7) | 2449 (79.7) | 302 ( 9.8) |
| *weighted* |  | ( 0.9) | ( 9.8) | (79.8) | ( 9.5) |

*Supplementary Table 22. Contingency table for variable of interest: Prematurity. Independent sample used for statistics.*

| **Demographics** | **n** | **Term** | **Late Preterm** | **Moderate Preterm** | **Very Preterm** | **Extremely Preterm** |
| --- | --- | --- | --- | --- | --- | --- |
| **sex_at_birth** |  |  |  |  |  |  |
| F | 4194 | 3749 (89.4) | 330 ( 7.9) | 75 ( 1.8) | 29 ( 0.7) | 11 ( 0.3) |
| *weighted* |  | (89.2) | ( 7.6) | ( 2.0) | ( 0.8) | ( 0.3) |
| M | 4589 | 4094 (89.2) | 361 ( 7.9) | 90 ( 2.0) | 29 ( 0.6) | 15 ( 0.3) |
| *weighted* |  | (89.1) | ( 7.8) | ( 1.9) | ( 0.7) | ( 0.4) |
| **race.4level** |  |  |  |  |  |  |
| White | 5779 | 5140 (88.9) | 469 ( 8.1) | 110 ( 1.9) | 49 ( 0.8) | 11 ( 0.2) |
| *weighted* |  | (88.6) | ( 8.2) | ( 2.0) | ( 1.0) | ( 0.3) |
| Black | 1291 | 1145 (88.7) | 104 ( 8.1) | 27 ( 2.1) | <10 | 10 ( 0.8) |
| *weighted* |  | (88.6) | ( 8.0) | ( 2.2) |  | ( 0.8) |
| Asian | 197 | 184 (93.4) | 10 ( 5.1) | <10 | <10 | <10 |
| *weighted* |  | (93.4) | ( 4.7) |  |  |  |
| Other/Mixed | 1516 | 1374 (90.6) | 108 ( 7.1) | 26 ( 1.7) | <10 | <10 |
| *weighted* |  | (91.4) | ( 6.2) | ( 1.8) |  |  |
| **hisp** |  |  |  |  |  |  |
| No | 7039 | 6266 (89.0) | 581 ( 8.3) | 127 ( 1.8) | 49 ( 0.7) | 16 ( 0.2) |
| *weighted* |  | (88.8) | ( 8.2) | ( 1.9) | ( 0.9) | ( 0.2) |
| Yes | 1744 | 1577 (90.4) | 110 ( 6.3) | 38 ( 2.2) | <10 | 10 ( 0.6) |
| *weighted* |  | (90.4) | ( 6.2) | ( 2.3) |  | ( 0.7) |
| **household.income** |  |  |  |  |  |  |
| [<50K] | 2623 | 2366 (90.2) | 163 ( 6.2) | 59 ( 2.2) | 17 ( 0.6) | 18 ( 0.7) |
| *weighted* |  | (90.1) | ( 6.2) | ( 2.4) | ( 0.6) | ( 0.6) |
| [>=50K & <100K] | 2500 | 2225 (89.0) | 207 ( 8.3) | 44 ( 1.8) | 20 ( 0.8) | <10 |
| *weighted* |  | (88.6) | ( 8.5) | ( 1.6) | ( 1.1) |  |
| [>=100K] | 3660 | 3252 (88.9) | 321 ( 8.8) | 62 ( 1.7) | 21 ( 0.6) | <10 |
| *weighted* |  | (88.6) | ( 9.0) | ( 1.7) | ( 0.6) |  |
| **high.educ** |  |  |  |  |  |  |
| < HS Diploma | 341 | 320 (93.8) | 12 ( 3.5) | <10 | <10 | <10 |
| *weighted* |  | (94.4) | ( 3.1) |  |  |  |
| HS Diploma/GED | 757 | 681 (90.0) | 47 ( 6.2) | 15 ( 2.0) | <10 | <10 |
| *weighted* |  | (89.4) | ( 6.4) | ( 2.3) |  |  |
| Some College | 2257 | 1983 (87.9) | 196 ( 8.7) | 57 ( 2.5) | 13 ( 0.6) | <10 |
| *weighted* |  | (88.1) | ( 8.4) | ( 2.4) | ( 0.7) |  |
| Bachelor | 2276 | 2004 (88.0) | 205 ( 9.0) | 48 ( 2.1) | 16 ( 0.7) | <10 |
| *weighted* |  | (87.8) | ( 9.0) | ( 2.2) | ( 0.9) |  |
| Post Graduate Degree | 3152 | 2855 (90.6) | 231 ( 7.3) | 41 ( 1.3) | 20 ( 0.6) | <10 |
| *weighted* |  | (90.5) | ( 7.2) | ( 1.4) | ( 0.7) |  |

*Supplementary Table 23. Contingency table for variable of interest: Age at first rolling over (months). Independent sample used for statistics.*

| **Demographics** | **n** | **Within guidelines** | **Beyond guidelines** |
| --- | --- | --- | --- |
| **sex_at_birth** |  |  |  |
| F | 4256 | 4210 (98.9) | 46 ( 1.1) |
| *weighted* |  | (98.8) | ( 1.2) |
| M | 4646 | 4594 (98.9) | 52 ( 1.1) |
| *weighted* |  | (98.8) | ( 1.2) |
| **race.4level** |  |  |  |
| White | 5824 | 5758 (98.9) | 66 ( 1.1) |
| *weighted* |  | (98.7) | ( 1.3) |
| Black | 1325 | 1310 (98.9) | 15 ( 1.1) |
| *weighted* |  | (98.9) | ( 1.1) |
| Asian | 215 | 213 (99.1) | <10 |
| *weighted* |  | (99.1) |  |
| Other/Mixed | 1538 | 1523 (99.0) | 15 ( 1.0) |
| *weighted* |  | (99.1) | ( 0.9) |
| **hisp** |  |  |  |
| No | 7130 | 7056 (99.0) | 74 ( 1.0) |
| *weighted* |  | (98.9) | ( 1.1) |
| Yes | 1772 | 1748 (98.6) | 24 ( 1.4) |
| *weighted* |  | (98.6) | ( 1.4) |
| **household.income** |  |  |  |
| [<50K] | 2667 | 2626 (98.5) | 41 ( 1.5) |
| *weighted* |  | (98.4) | ( 1.6) |
| [>=50K & <100K] | 2529 | 2505 (99.1) | 24 ( 0.9) |
| *weighted* |  | (99.1) | ( 0.9) |
| [>=100K] | 3706 | 3673 (99.1) | 33 ( 0.9) |
| *weighted* |  | (99.1) | ( 0.9) |
| **high.educ** |  |  |  |
| < HS Diploma | 347 | 341 (98.3) | <10 |
| *weighted* |  | (98.5) |  |
| HS Diploma/GED | 773 | 763 (98.7) | 10 ( 1.3) |
| *weighted* |  | (98.6) | ( 1.4) |
| Some College | 2289 | 2261 (98.8) | 28 ( 1.2) |
| *weighted* |  | (98.6) | ( 1.4) |
| Bachelor | 2298 | 2271 (98.8) | 27 ( 1.2) |
| *weighted* |  | (98.9) | ( 1.1) |
| Post Graduate Degree | 3195 | 3168 (99.2) | 27 ( 0.8) |
| *weighted* |  | (99.0) | ( 1.0) |

*Supplementary Table 24. Contingency table for variable of interest: Age at first sitting (months). Independent sample used for statistics.*

| **Demographics** | **n** | **Within guidelines** | **Beyond guidelines** |
| --- | --- | --- | --- |
| **sex_at_birth** |  |  |  |
| F | 4256 | 4118 (96.8) | 138 ( 3.2) |
| *weighted* |  | (96.5) | ( 3.5) |
| M | 4646 | 4437 (95.5) | 209 ( 4.5) |
| *weighted* |  | (94.9) | ( 5.1) |
| **race.4level** |  |  |  |
| White | 5824 | 5572 (95.7) | 252 ( 4.3) |
| *weighted* |  | (95.0) | ( 5.0) |
| Black | 1325 | 1280 (96.6) | 45 ( 3.4) |
| *weighted* |  | (96.7) | ( 3.3) |
| Asian | 215 | 208 (96.7) | <10 |
| *weighted* |  | (96.4) |  |
| Other/Mixed | 1538 | 1495 (97.2) | 43 ( 2.8) |
| *weighted* |  | (97.4) | ( 2.6) |
| **hisp** |  |  |  |
| No | 7130 | 6859 (96.2) | 271 ( 3.8) |
| *weighted* |  | (95.7) | ( 4.3) |
| Yes | 1772 | 1696 (95.7) | 76 ( 4.3) |
| *weighted* |  | (95.7) | ( 4.3) |
| **household.income** |  |  |  |
| [<50K] | 2667 | 2546 (95.5) | 121 ( 4.5) |
| *weighted* |  | (95.0) | ( 5.0) |
| [>=50K & <100K] | 2529 | 2429 (96.0) | 100 ( 4.0) |
| *weighted* |  | (95.8) | ( 4.2) |
| [>=100K] | 3706 | 3580 (96.6) | 126 ( 3.4) |
| *weighted* |  | (96.5) | ( 3.5) |
| **high.educ** |  |  |  |
| < HS Diploma | 347 | 329 (94.8) | 18 ( 5.2) |
| *weighted* |  | (94.2) | ( 5.8) |
| HS Diploma/GED | 773 | 741 (95.9) | 32 ( 4.1) |
| *weighted* |  | (96.1) | ( 3.9) |
| Some College | 2289 | 2196 (95.9) | 93 ( 4.1) |
| *weighted* |  | (95.5) | ( 4.5) |
| Bachelor | 2298 | 2209 (96.1) | 89 ( 3.9) |
| *weighted* |  | (95.6) | ( 4.4) |
| Post Graduate Degree | 3195 | 3080 (96.4) | 115 ( 3.6) |
| *weighted* |  | (96.0) | ( 4.0) |

*Supplementary Table 25. Contingency table for variable of interest: Age at first walking (months). Independent sample used for statistics.*

| **Demographics** | **n** | **Within guidelines** | **Beyond guidelines** |
| --- | --- | --- | --- |
| **sex_at_birth** |  |  |  |
| F | 4256 | 4154 (97.6) | 102 ( 2.4) |
| *weighted* |  | (97.4) | ( 2.6) |
| M | 4646 | 4476 (96.3) | 170 ( 3.7) |
| *weighted* |  | (96.1) | ( 3.9) |
| **race.4level** |  |  |  |
| White | 5824 | 5638 (96.8) | 186 ( 3.2) |
| *weighted* |  | (96.5) | ( 3.5) |
| Black | 1325 | 1284 (96.9) | 41 ( 3.1) |
| *weighted* |  | (97.2) | ( 2.8) |
| Asian | 215 | 208 (96.7) | <10 |
| *weighted* |  | (96.3) |  |
| Other/Mixed | 1538 | 1500 (97.5) | 38 ( 2.5) |
| *weighted* |  | (97.4) | ( 2.6) |
| **hisp** |  |  |  |
| No | 7130 | 6918 (97.0) | 212 ( 3.0) |
| *weighted* |  | (96.8) | ( 3.2) |
| Yes | 1772 | 1712 (96.6) | 60 ( 3.4) |
| *weighted* |  | (96.4) | ( 3.6) |
| **household.income** |  |  |  |
| [<50K] | 2667 | 2566 (96.2) | 101 ( 3.8) |
| *weighted* |  | (96.0) | ( 4.0) |
| [>=50K & <100K] | 2529 | 2457 (97.2) | 72 ( 2.8) |
| *weighted* |  | (97.1) | ( 2.9) |
| [>=100K] | 3706 | 3607 (97.3) | 99 ( 2.7) |
| *weighted* |  | (97.3) | ( 2.7) |
| **high.educ** |  |  |  |
| < HS Diploma | 347 | 332 (95.7) | 15 ( 4.3) |
| *weighted* |  | (95.5) | ( 4.5) |
| HS Diploma/GED | 773 | 748 (96.8) | 25 ( 3.2) |
| *weighted* |  | (97.2) | ( 2.8) |
| Some College | 2289 | 2210 (96.5) | 79 ( 3.5) |
| *weighted* |  | (96.1) | ( 3.9) |
| Bachelor | 2298 | 2235 (97.3) | 63 ( 2.7) |
| *weighted* |  | (97.0) | ( 3.0) |
| Post Graduate Degree | 3195 | 3105 (97.2) | 90 ( 2.8) |
| *weighted* |  | (97.1) | ( 2.9) |

*Supplementary Table 26. Contingency table for variable of interest: Age at first word (months). Independent sample used for statistics.*

| **Demographics** | **n** | **Within guidelines** | **Beyond guidelines** |
| --- | --- | --- | --- |
| **sex_at_birth** |  |  |  |
| F | 4256 | 3951 (92.8) | 305 ( 7.2) |
| *weighted* |  | (92.7) | ( 7.3) |
| M | 4646 | 4107 (88.4) | 539 (11.6) |
| *weighted* |  | (88.1) | (11.9) |
| **race.4level** |  |  |  |
| White | 5824 | 5245 (90.1) | 579 ( 9.9) |
| *weighted* |  | (89.9) | (10.1) |
| Black | 1325 | 1219 (92.0) | 106 ( 8.0) |
| *weighted* |  | (92.4) | ( 7.6) |
| Asian | 215 | 186 (86.5) | 29 (13.5) |
| *weighted* |  | (85.7) | (14.3) |
| Other/Mixed | 1538 | 1408 (91.5) | 130 ( 8.5) |
| *weighted* |  | (92.1) | ( 7.9) |
| **hisp** |  |  |  |
| No | 7130 | 6446 (90.4) | 684 ( 9.6) |
| *weighted* |  | (90.2) | ( 9.8) |
| Yes | 1772 | 1612 (91.0) | 160 ( 9.0) |
| *weighted* |  | (91.0) | ( 9.0) |
| **household.income** |  |  |  |
| [<50K] | 2667 | 2441 (91.5) | 226 ( 8.5) |
| *weighted* |  | (91.1) | ( 8.9) |
| [>=50K & <100K] | 2529 | 2283 (90.3) | 246 ( 9.7) |
| *weighted* |  | (90.3) | ( 9.7) |
| [>=100K] | 3706 | 3334 (90.0) | 372 (10.0) |
| *weighted* |  | (89.5) | (10.5) |
| **high.educ** |  |  |  |
| < HS Diploma | 347 | 326 (93.9) | 21 ( 6.1) |
| *weighted* |  | (94.2) | ( 5.8) |
| HS Diploma/GED | 773 | 710 (91.8) | 63 ( 8.2) |
| *weighted* |  | (91.7) | ( 8.3) |
| Some College | 2289 | 2077 (90.7) | 212 ( 9.3) |
| *weighted* |  | (90.4) | ( 9.6) |
| Bachelor | 2298 | 2074 (90.3) | 224 ( 9.7) |
| *weighted* |  | (89.9) | (10.1) |
| Post Graduate Degree | 3195 | 2871 (89.9) | 324 (10.1) |
| *weighted* |  | (89.6) | (10.4) |

*Supplementary Table 27. Contingency table for variable of interest: Total pregnancy problems. Independent sample used for statistics.*

| **Demographics** | **n** | **None** | **One problem** | **Two or more problems** |
| --- | --- | --- | --- | --- |
| **sex_at_birth** |  |  |  |  |
| F | 4256 | 2555 (60.0) | 1092 (25.7) | 609 (14.3) |
| *weighted* |  | (58.7) | (26.0) | (15.3) |
| M | 4646 | 2893 (62.3) | 1106 (23.8) | 647 (13.9) |
| *weighted* |  | (61.7) | (24.2) | (14.1) |
| **race.4level** |  |  |  |  |
| White | 5824 | 3663 (62.9) | 1427 (24.5) | 734 (12.6) |
| *weighted* |  | (61.4) | (25.0) | (13.6) |
| Black | 1325 | 735 (55.5) | 341 (25.7) | 249 (18.8) |
| *weighted* |  | (53.4) | (27.3) | (19.3) |
| Asian | 215 | 158 (73.5) | 44 (20.5) | 13 ( 6.0) |
| *weighted* |  | (75.3) | (18.8) | ( 5.9) |
| Other/Mixed | 1538 | 892 (58.0) | 386 (25.1) | 260 (16.9) |
| *weighted* |  | (56.8) | (25.0) | (18.1) |
| **hisp** |  |  |  |  |
| No | 7130 | 4445 (62.3) | 1689 (23.7) | 996 (14.0) |
| *weighted* |  | (61.7) | (24.0) | (14.3) |
| Yes | 1772 | 1003 (56.6) | 509 (28.7) | 260 (14.7) |
| *weighted* |  | (55.4) | (28.6) | (16.0) |
| **household.income** |  |  |  |  |
| [<50K] | 2667 | 1451 (54.4) | 711 (26.7) | 505 (18.9) |
| *weighted* |  | (55.1) | (26.6) | (18.3) |
| [>=50K & <100K] | 2529 | 1551 (61.3) | 627 (24.8) | 351 (13.9) |
| *weighted* |  | (61.3) | (25.1) | (13.6) |
| [>=100K] | 3706 | 2446 (66.0) | 860 (23.2) | 400 (10.8) |
| *weighted* |  | (66.1) | (23.0) | (10.9) |
| **high.educ** |  |  |  |  |
| < HS Diploma | 347 | 188 (54.2) | 98 (28.2) | 61 (17.6) |
| *weighted* |  | (54.9) | (27.9) | (17.2) |
| HS Diploma/GED | 773 | 434 (56.1) | 191 (24.7) | 148 (19.1) |
| *weighted* |  | (55.8) | (24.5) | (19.7) |
| Some College | 2289 | 1236 (54.0) | 618 (27.0) | 435 (19.0) |
| *weighted* |  | (54.2) | (26.8) | (19.0) |
| Bachelor | 2298 | 1440 (62.7) | 562 (24.5) | 296 (12.9) |
| *weighted* |  | (61.8) | (25.1) | (13.0) |
| Post Graduate Degree | 3195 | 2150 (67.3) | 729 (22.8) | 316 ( 9.9) |
| *weighted* |  | (67.2) | (23.1) | ( 9.7) |

*Supplementary Table 28. Contingency table for variable of interest: Total birth problems. Independent sample used for statistics.*

| **Demographics** | **n** | **None** | **One problem** | **Two or more problems** |
| --- | --- | --- | --- | --- |
| **sex_at_birth** |  |  |  |  |
| F | 4256 | 3318 (78.0) | 711 (16.7) | 227 ( 5.3) |
| *weighted* |  | (78.4) | (16.2) | ( 5.4) |
| M | 4646 | 3485 (75.0) | 862 (18.6) | 299 ( 6.4) |
| *weighted* |  | (75.3) | (18.3) | ( 6.3) |
| **race.4level** |  |  |  |  |
| White | 5824 | 4401 (75.6) | 1042 (17.9) | 381 ( 6.5) |
| *weighted* |  | (75.8) | (17.7) | ( 6.5) |
| Black | 1325 | 1041 (78.6) | 213 (16.1) | 71 ( 5.4) |
| *weighted* |  | (77.9) | (16.6) | ( 5.4) |
| Asian | 215 | 168 (78.1) | 43 (20.0) | <10 |
| *weighted* |  | (80.9) | (17.4) |  |
| Other/Mixed | 1538 | 1193 (77.6) | 275 (17.9) | 70 ( 4.6) |
| *weighted* |  | (79.5) | (16.1) | ( 4.4) |
| **hisp** |  |  |  |  |
| No | 7130 | 5394 (75.7) | 1294 (18.1) | 442 ( 6.2) |
| *weighted* |  | (75.8) | (17.8) | ( 6.3) |
| Yes | 1772 | 1409 (79.5) | 279 (15.7) | 84 ( 4.7) |
| *weighted* |  | (80.1) | (15.6) | ( 4.3) |
| **household.income** |  |  |  |  |
| [<50K] | 2667 | 2113 (79.2) | 412 (15.4) | 142 ( 5.3) |
| *weighted* |  | (79.1) | (15.5) | ( 5.4) |
| [>=50K & <100K] | 2529 | 1880 (74.3) | 495 (19.6) | 154 ( 6.1) |
| *weighted* |  | (74.5) | (19.3) | ( 6.1) |
| [>=100K] | 3706 | 2810 (75.8) | 666 (18.0) | 230 ( 6.2) |
| *weighted* |  | (76.3) | (17.6) | ( 6.1) |
| **high.educ** |  |  |  |  |
| < HS Diploma | 347 | 283 (81.6) | 49 (14.1) | 15 ( 4.3) |
| *weighted* |  | (80.1) | (15.9) | ( 4.1) |
| HS Diploma/GED | 773 | 629 (81.4) | 102 (13.2) | 42 ( 5.4) |
| *weighted* |  | (80.7) | (14.3) | ( 5.0) |
| Some College | 2289 | 1734 (75.8) | 401 (17.5) | 154 ( 6.7) |
| *weighted* |  | (76.3) | (16.9) | ( 6.8) |
| Bachelor | 2298 | 1680 (73.1) | 465 (20.2) | 153 ( 6.7) |
| *weighted* |  | (73.5) | (19.7) | ( 6.8) |
| Post Graduate Degree | 3195 | 2477 (77.5) | 556 (17.4) | 162 ( 5.1) |
| *weighted* |  | (78.3) | (17.0) | ( 4.8) |

*Supplementary Table 29. Contingency table for variable of interest: Prenatal alcohol exposure. Independent sample used for statistics.*

| **Demographics** | **n** | **Pre-No/Post-No** | **Pre-Yes/Post-No** | **Pre-Yes/Post-Yes** |
| --- | --- | --- | --- | --- |
| **sex_at_birth** |  |  |  |  |
| F | 3965 | 2847 (71.8) | 1003 (25.3) | 115 ( 2.9) |
| *weighted* |  | (73.3) | (24.0) | ( 2.6) |
| M | 4343 | 3173 (73.1) | 1057 (24.3) | 113 ( 2.6) |
| *weighted* |  | (74.5) | (23.0) | ( 2.5) |
| **race.4level** |  |  |  |  |
| White | 5469 | 3788 (69.3) | 1504 (27.5) | 177 ( 3.2) |
| *weighted* |  | (70.7) | (26.3) | ( 3.0) |
| Black | 1229 | 1008 (82.0) | 199 (16.2) | 22 ( 1.8) |
| *weighted* |  | (82.8) | (15.5) | ( 1.7) |
| Asian | 187 | 154 (82.4) | 33 (17.6) | 0 ( 0.0) |
| *weighted* |  | (84.3) | (15.7) | ( 0.0) |
| Other/Mixed | 1423 | 1070 (75.2) | 324 (22.8) | 29 ( 2.0) |
| *weighted* |  | (78.4) | (19.6) | ( 2.0) |
| **hisp** |  |  |  |  |
| No | 6635 | 4710 (71.0) | 1733 (26.1) | 192 ( 2.9) |
| *weighted* |  | (71.9) | (25.3) | ( 2.8) |
| Yes | 1673 | 1310 (78.3) | 327 (19.5) | 36 ( 2.2) |
| *weighted* |  | (80.3) | (17.7) | ( 2.0) |
| **household.income** |  |  |  |  |
| [<50K] | 2530 | 2074 (82.0) | 410 (16.2) | 46 ( 1.8) |
| *weighted* |  | (81.7) | (16.6) | ( 1.7) |
| [>=50K & <100K] | 2350 | 1692 (72.0) | 604 (25.7) | 54 ( 2.3) |
| *weighted* |  | (71.4) | (26.1) | ( 2.5) |
| [>=100K] | 3428 | 2254 (65.8) | 1046 (30.5) | 128 ( 3.7) |
| *weighted* |  | (66.0) | (30.2) | ( 3.8) |
| **high.educ** |  |  |  |  |
| < HS Diploma | 334 | 306 (91.6) | 24 ( 7.2) | <10 |
| *weighted* |  | (92.4) | ( 6.6) |  |
| HS Diploma/GED | 738 | 624 (84.6) | 107 (14.5) | <10 |
| *weighted* |  | (83.4) | (15.8) |  |
| Some College | 2148 | 1607 (74.8) | 490 (22.8) | 51 ( 2.4) |
| *weighted* |  | (75.2) | (22.2) | ( 2.5) |
| Bachelor | 2147 | 1499 (69.8) | 587 (27.3) | 61 ( 2.8) |
| *weighted* |  | (71.2) | (26.0) | ( 2.8) |
| Post Graduate Degree | 2941 | 1984 (67.5) | 852 (29.0) | 105 ( 3.6) |
| *weighted* |  | (68.5) | (28.2) | ( 3.3) |

*Supplementary Table 30. Contingency table for variable of interest: Prenatal tobacco exposure. Independent sample used for statistics.*

| **Demographics** | **n** | **Pre-No/Post-No** | **Pre-Yes/Post-No** | **Pre-Yes/Post-Yes** |
| --- | --- | --- | --- | --- |
| **sex_at_birth** |  |  |  |  |
| F | 4117 | 3566 (86.6) | 337 ( 8.2) | 214 ( 5.2) |
| *weighted* |  | (83.7) | ( 9.4) | ( 6.9) |
| M | 4524 | 3904 (86.3) | 388 ( 8.6) | 232 ( 5.1) |
| *weighted* |  | (83.7) | ( 9.9) | ( 6.4) |
| **race.4level** |  |  |  |  |
| White | 5726 | 5059 (88.4) | 409 ( 7.1) | 258 ( 4.5) |
| *weighted* |  | (84.6) | ( 8.8) | ( 6.5) |
| Black | 1250 | 1007 (80.6) | 143 (11.4) | 100 ( 8.0) |
| *weighted* |  | (79.2) | (12.1) | ( 8.7) |
| Asian | 191 | 179 (93.7) | <10 | <10 |
| *weighted* |  | (93.8) |  |  |
| Other/Mixed | 1474 | 1225 (83.1) | 164 (11.1) | 85 ( 5.8) |
| *weighted* |  | (80.7) | (13.0) | ( 6.3) |
| **hisp** |  |  |  |  |
| No | 6919 | 5953 (86.0) | 576 ( 8.3) | 390 ( 5.6) |
| *weighted* |  | (82.8) | ( 9.6) | ( 7.6) |
| Yes | 1722 | 1517 (88.1) | 149 ( 8.7) | 56 ( 3.3) |
| *weighted* |  | (86.6) | ( 9.8) | ( 3.6) |
| **household.income** |  |  |  |  |
| [<50K] | 2591 | 1967 (75.9) | 345 (13.3) | 279 (10.8) |
| *weighted* |  | (74.9) | (13.4) | (11.7) |
| [>=50K & <100K] | 2448 | 2113 (86.3) | 221 ( 9.0) | 114 ( 4.7) |
| *weighted* |  | (85.1) | ( 9.8) | ( 5.1) |
| [>=100K] | 3602 | 3390 (94.1) | 159 ( 4.4) | 53 ( 1.5) |
| *weighted* |  | (94.0) | ( 4.6) | ( 1.4) |
| **high.educ** |  |  |  |  |
| < HS Diploma | 335 | 261 (77.9) | 44 (13.1) | 30 ( 9.0) |
| *weighted* |  | (77.9) | (11.6) | (10.5) |
| HS Diploma/GED | 754 | 555 (73.6) | 91 (12.1) | 108 (14.3) |
| *weighted* |  | (71.0) | (12.6) | (16.3) |
| Some College | 2223 | 1679 (75.5) | 337 (15.2) | 207 ( 9.3) |
| *weighted* |  | (73.2) | (16.2) | (10.6) |
| Bachelor | 2233 | 2037 (91.2) | 131 ( 5.9) | 65 ( 2.9) |
| *weighted* |  | (89.7) | ( 6.8) | ( 3.5) |
| Post Graduate Degree | 3096 | 2938 (94.9) | 122 ( 3.9) | 36 ( 1.2) |
| *weighted* |  | (94.4) | ( 4.3) | ( 1.4) |

*Supplementary Table 31. Contingency table for variable of interest: Prenatal marijuana exposure. Independent sample used for statistics.*

| **Demographics** | **n** | **Pre-No/Post-No** | **Pre-Yes/Post-No** | **Pre-Yes/Post-Yes** |
| --- | --- | --- | --- | --- |
| **sex_at_birth** |  |  |  |  |
| F | 4102 | 3855 (94.0) | 149 ( 3.6) | 98 ( 2.4) |
| *weighted* |  | (93.2) | ( 4.0) | ( 2.9) |
| M | 4497 | 4235 (94.2) | 174 ( 3.9) | 88 ( 2.0) |
| *weighted* |  | (93.5) | ( 4.3) | ( 2.2) |
| **race.4level** |  |  |  |  |
| White | 5699 | 5442 (95.5) | 158 ( 2.8) | 99 ( 1.7) |
| *weighted* |  | (94.5) | ( 3.3) | ( 2.3) |
| Black | 1246 | 1085 (87.1) | 101 ( 8.1) | 60 ( 4.8) |
| *weighted* |  | (86.7) | ( 8.4) | ( 5.0) |
| Asian | 191 | 188 (98.4) | <10 | <10 |
| *weighted* |  | (98.4) |  |  |
| Other/Mixed | 1463 | 1375 (94.0) | 62 ( 4.2) | 26 ( 1.8) |
| *weighted* |  | (93.0) | ( 5.3) | ( 1.8) |
| **hisp** |  |  |  |  |
| No | 6883 | 6473 (94.0) | 251 ( 3.6) | 159 ( 2.3) |
| *weighted* |  | (93.2) | ( 4.0) | ( 2.8) |
| Yes | 1716 | 1617 (94.2) | 72 ( 4.2) | 27 ( 1.6) |
| *weighted* |  | (93.8) | ( 4.6) | ( 1.7) |
| **household.income** |  |  |  |  |
| [<50K] | 2568 | 2275 (88.6) | 182 ( 7.1) | 111 ( 4.3) |
| *weighted* |  | (89.1) | ( 6.6) | ( 4.3) |
| [>=50K & <100K] | 2437 | 2307 (94.7) | 74 ( 3.0) | 56 ( 2.3) |
| *weighted* |  | (94.6) | ( 3.2) | ( 2.3) |
| [>=100K] | 3594 | 3508 (97.6) | 67 ( 1.9) | 19 ( 0.5) |
| *weighted* |  | (97.7) | ( 1.9) | ( 0.4) |
| **high.educ** |  |  |  |  |
| < HS Diploma | 332 | 313 (94.3) | 12 ( 3.6) | <10 |
| *weighted* |  | (93.6) | ( 3.2) |  |
| HS Diploma/GED | 744 | 654 (87.9) | 56 ( 7.5) | 34 ( 4.6) |
| *weighted* |  | (87.4) | ( 8.2) | ( 4.4) |
| Some College | 2210 | 1974 (89.3) | 143 ( 6.5) | 93 ( 4.2) |
| *weighted* |  | (89.5) | ( 6.2) | ( 4.3) |
| Bachelor | 2231 | 2144 (96.1) | 55 ( 2.5) | 32 ( 1.4) |
| *weighted* |  | (95.5) | ( 2.8) | ( 1.7) |
| Post Graduate Degree | 3082 | 3005 (97.5) | 57 ( 1.8) | 20 ( 0.6) |
| *weighted* |  | (97.3) | ( 2.0) | ( 0.7) |

*Supplementary Table 32. Contingency table for variable of interest: Prenatal other substance exposure. Independent sample used for statistics.*

| **Demographics** | **n** | **Pre-No/Post-No** | **Pre-Yes/Post-No** | **Pre-Yes/Post-Yes** |
| --- | --- | --- | --- | --- |
| **sex_at_birth** |  |  |  |  |
| F | 4246 | 4162 (98.0) | 57 ( 1.3) | 27 ( 0.6) |
| *weighted* |  | (97.7) | ( 1.7) | ( 0.7) |
| M | 4639 | 4547 (98.0) | 55 ( 1.2) | 37 ( 0.8) |
| *weighted* |  | (97.8) | ( 1.3) | ( 0.9) |
| **race.4level** |  |  |  |  |
| White | 5813 | 5716 (98.3) | 64 ( 1.1) | 33 ( 0.6) |
| *weighted* |  | (97.9) | ( 1.4) | ( 0.7) |
| Black | 1320 | 1294 (98.0) | <10 | 18 ( 1.4) |
| *weighted* |  | (98.1) |  | ( 1.3) |
| Asian | 215 | 213 (99.1) | <10 | <10 |
| *weighted* |  | (98.8) |  |  |
| Other/Mixed | 1537 | 1486 (96.7) | 38 ( 2.5) | 13 ( 0.8) |
| *weighted* |  | (96.4) | ( 2.9) | ( 0.6) |
| **hisp** |  |  |  |  |
| No | 7115 | 6984 (98.2) | 80 ( 1.1) | 51 ( 0.7) |
| *weighted* |  | (97.9) | ( 1.4) | ( 0.7) |
| Yes | 1770 | 1725 (97.5) | 32 ( 1.8) | 13 ( 0.7) |
| *weighted* |  | (97.4) | ( 1.9) | ( 0.8) |
| **household.income** |  |  |  |  |
| [<50K] | 2664 | 2581 (96.9) | 56 ( 2.1) | 27 ( 1.0) |
| *weighted* |  | (96.7) | ( 2.3) | ( 1.0) |
| [>=50K & <100K] | 2521 | 2471 (98.0) | 32 ( 1.3) | 18 ( 0.7) |
| *weighted* |  | (97.9) | ( 1.3) | ( 0.7) |
| [>=100K] | 3700 | 3657 (98.8) | 24 ( 0.6) | 19 ( 0.5) |
| *weighted* |  | (99.1) | ( 0.5) | ( 0.4) |
| **high.educ** |  |  |  |  |
| < HS Diploma | 347 | 341 (98.3) | <10 | <10 |
| *weighted* |  | (97.9) |  |  |
| HS Diploma/GED | 772 | 750 (97.2) | 15 ( 1.9) | <10 |
| *weighted* |  | (96.8) | ( 2.4) |  |
| Some College | 2284 | 2203 (96.5) | 55 ( 2.4) | 26 ( 1.1) |
| *weighted* |  | (96.1) | ( 2.6) | ( 1.3) |
| Bachelor | 2290 | 2261 (98.7) | 19 ( 0.8) | 10 ( 0.4) |
| *weighted* |  | (98.6) | ( 1.0) | ( 0.3) |
| Post Graduate Degree | 3192 | 3154 (98.8) | 20 ( 0.6) | 18 ( 0.6) |
| *weighted* |  | (99.0) | ( 0.5) | ( 0.5) |

*Supplementary Table 33. Contingency table for variable of interest: Average hours of sleep per night. Independent sample used for statistics.*

| **Demographics** | **n** | **Less than 7 hours** | **7-8 hours** | **8-9 hours** | **9-11 hours** |
| --- | --- | --- | --- | --- | --- |
| **age_quartiles** |  |  |  |  |  |
| age<=25% | 2308 | 66 ( 2.9) | 250 (10.8) | 745 (32.3) | 1247 (54.0) |
| *weighted* |  | ( 3.1) | (12.0) | (34.5) | (50.4) |
| age>25&<=50% | 2286 | 56 ( 2.4) | 241 (10.5) | 803 (35.1) | 1186 (51.9) |
| *weighted* |  | ( 2.5) | (12.0) | (36.2) | (49.3) |
| age>50&<=75% | 1960 | 69 ( 3.5) | 213 (10.9) | 751 (38.3) | 927 (47.3) |
| *weighted* |  | ( 3.8) | (12.3) | (40.4) | (43.4) |
| age>75% | 2039 | 83 ( 4.1) | 271 (13.3) | 813 (39.9) | 872 (42.8) |
| *weighted* |  | ( 4.5) | (14.5) | (41.8) | (39.1) |
| **sex_at_birth** |  |  |  |  |  |
| F | 4105 | 118 ( 2.9) | 440 (10.7) | 1516 (36.9) | 2031 (49.5) |
| *weighted* |  | ( 3.3) | (12.1) | (38.7) | (45.9) |
| M | 4488 | 156 ( 3.5) | 535 (11.9) | 1596 (35.6) | 2201 (49.0) |
| *weighted* |  | ( 3.7) | (13.3) | (37.6) | (45.4) |
| **race.4level** |  |  |  |  |  |
| White | 5705 | 82 ( 1.4) | 466 ( 8.2) | 1940 (34.0) | 3217 (56.4) |
| *weighted* |  | ( 1.8) | (10.0) | (36.5) | (51.7) |
| Black | 1217 | 133 (10.9) | 303 (24.9) | 501 (41.2) | 280 (23.0) |
| *weighted* |  | (11.9) | (25.9) | (40.9) | (21.3) |
| Asian | 201 | 2 ( 1.0) | 13 ( 6.5) | 78 (38.8) | 108 (53.7) |
| *weighted* |  | ( 1.2) | ( 7.0) | (41.4) | (50.5) |
| Other/Mixed | 1470 | 57 ( 3.9) | 193 (13.1) | 593 (40.3) | 627 (42.7) |
| *weighted* |  | ( 4.5) | (15.6) | (42.9) | (37.0) |
| **hisp** |  |  |  |  |  |
| No | 6901 | 221 ( 3.2) | 721 (10.4) | 2355 (34.1) | 3604 (52.2) |
| *weighted* |  | ( 3.4) | (11.5) | (35.7) | (49.3) |
| Yes | 1692 | 53 ( 3.1) | 254 (15.0) | 757 (44.7) | 628 (37.1) |
| *weighted* |  | ( 3.8) | (16.6) | (46.1) | (33.6) |
| **household.income** |  |  |  |  |  |
| [<50K] | 2480 | 177 ( 7.1) | 500 (20.2) | 1062 (42.8) | 741 (29.9) |
| *weighted* |  | ( 6.3) | (19.9) | (43.7) | (30.1) |
| [>=50K & <100K] | 2460 | 62 ( 2.5) | 261 (10.6) | 932 (37.9) | 1205 (49.0) |
| *weighted* |  | ( 2.6) | (10.6) | (38.5) | (48.3) |
| [>=100K] | 3653 | 35 ( 1.0) | 214 ( 5.9) | 1118 (30.6) | 2286 (62.6) |
| *weighted* |  | ( 0.9) | ( 5.7) | (30.7) | (62.6) |
| **high.educ** |  |  |  |  |  |
| < HS Diploma | 304 | 22 ( 7.2) | 69 (22.7) | 140 (46.1) | 73 (24.0) |
| *weighted* |  | ( 7.4) | (22.2) | (47.9) | (22.5) |
| HS Diploma/GED | 700 | 52 ( 7.4) | 148 (21.1) | 321 (45.9) | 179 (25.6) |
| *weighted* |  | ( 6.7) | (21.7) | (44.9) | (26.7) |
| Some College | 2196 | 127 ( 5.8) | 386 (17.6) | 939 (42.8) | 744 (33.9) |
| *weighted* |  | ( 5.5) | (18.2) | (43.6) | (32.7) |
| Bachelor | 2251 | 45 ( 2.0) | 184 ( 8.2) | 778 (34.6) | 1244 (55.3) |
| *weighted* |  | ( 2.3) | ( 9.1) | (36.1) | (52.4) |
| Post Graduate Degree | 3142 | 28 ( 0.9) | 188 ( 6.0) | 934 (29.7) | 1992 (63.4) |
| *weighted* |  | ( 1.0) | ( 6.2) | (31.0) | (61.8) |
| **pubertal_dev_collapsed** |  |  |  |  |  |
| Pre | 4409 | 93 ( 2.1) | 394 ( 8.9) | 1474 (33.4) | 2448 (55.5) |
| *weighted* |  | ( 2.5) | (10.0) | (35.6) | (51.9) |
| Early | 2058 | 66 ( 3.2) | 243 (11.8) | 755 (36.7) | 994 (48.3) |
| *weighted* |  | ( 3.3) | (12.9) | (38.8) | (45.0) |
| Mid | 2000 | 111 ( 5.5) | 315 (15.8) | 817 (40.8) | 757 (37.9) |
| *weighted* |  | ( 5.7) | (17.2) | (41.4) | (35.6) |
| Late/Post | 126 | 4 ( 3.2) | 23 (18.3) | 66 (52.4) | 33 (26.2) |
| *weighted* |  | ( 2.6) | (19.1) | (54.0) | (24.3) |

*Supplementary Table 34. Contingency table for variable of interest: Total sleep disturbance. Independent sample used for statistics.*

| **Demographics** | **n** | **Low Sleep Disturbance** | **High Sleep Disturbance** |
| --- | --- | --- | --- |
| **age_quartiles** |  |  |  |
| age<=25% | 2308 | 1540 (66.7) | 768 (33.3) |
| *weighted* |  | (65.0) | (35.0) |
| age>25&<=50% | 2286 | 1585 (69.3) | 701 (30.7) |
| *weighted* |  | (68.4) | (31.6) |
| age>50&<=75% | 1960 | 1337 (68.2) | 623 (31.8) |
| *weighted* |  | (66.5) | (33.5) |
| age>75% | 2039 | 1401 (68.7) | 638 (31.3) |
| *weighted* |  | (67.6) | (32.4) |
| **sex_at_birth** |  |  |  |
| F | 4105 | 2840 (69.2) | 1265 (30.8) |
| *weighted* |  | (68.0) | (32.0) |
| M | 4488 | 3023 (67.4) | 1465 (32.6) |
| *weighted* |  | (65.9) | (34.1) |
| **race.4level** |  |  |  |
| White | 5705 | 3975 (69.7) | 1730 (30.3) |
| *weighted* |  | (67.9) | (32.1) |
| Black | 1217 | 794 (65.2) | 423 (34.8) |
| *weighted* |  | (64.7) | (35.3) |
| Asian | 201 | 145 (72.1) | 56 (27.9) |
| *weighted* |  | (70.2) | (29.8) |
| Other/Mixed | 1470 | 949 (64.6) | 521 (35.4) |
| *weighted* |  | (62.9) | (37.1) |
| **hisp** |  |  |  |
| No | 6901 | 4700 (68.1) | 2201 (31.9) |
| *weighted* |  | (66.9) | (33.1) |
| Yes | 1692 | 1163 (68.7) | 529 (31.3) |
| *weighted* |  | (66.9) | (33.1) |
| **household.income** |  |  |  |
| [<50K] | 2480 | 1558 (62.8) | 922 (37.2) |
| *weighted* |  | (62.2) | (37.8) |
| [>=50K & <100K] | 2460 | 1640 (66.7) | 820 (33.3) |
| *weighted* |  | (66.4) | (33.6) |
| [>=100K] | 3653 | 2665 (73.0) | 988 (27.0) |
| *weighted* |  | (73.3) | (26.7) |
| **high.educ** |  |  |  |
| < HS Diploma | 304 | 198 (65.1) | 106 (34.9) |
| *weighted* |  | (63.3) | (36.7) |
| HS Diploma/GED | 700 | 450 (64.3) | 250 (35.7) |
| *weighted* |  | (63.5) | (36.5) |
| Some College | 2196 | 1393 (63.4) | 803 (36.6) |
| *weighted* |  | (62.2) | (37.8) |
| Bachelor | 2251 | 1557 (69.2) | 694 (30.8) |
| *weighted* |  | (68.4) | (31.6) |
| Post Graduate Degree | 3142 | 2265 (72.1) | 877 (27.9) |
| *weighted* |  | (71.8) | (28.2) |
| **pubertal_dev_collapsed** |  |  |  |
| Pre | 4409 | 3132 (71.0) | 1277 (29.0) |
| *weighted* |  | (70.0) | (30.0) |
| Early | 2058 | 1352 (65.7) | 706 (34.3) |
| *weighted* |  | (64.8) | (35.2) |
| Mid | 2000 | 1300 (65.0) | 700 (35.0) |
| *weighted* |  | (63.4) | (36.6) |
| Late/Post | 126 | 79 (62.7) | 47 (37.3) |
| *weighted* |  | (60.6) | (39.4) |

*Supplementary Table 35. Contingency table for variable of interest: Physical activity (vigorous). Independent sample used for statistics.*

| **Demographics** | **n** | **Below guidelines** | **Within guidelines** |
| --- | --- | --- | --- |
| **age_quartiles** |  |  |  |
| age<=25% | 2308 | 1917 (83.1) | 391 (16.9) |
| *weighted* |  | (83.6) | (16.4) |
| age>25&<=50% | 2286 | 1913 (83.7) | 373 (16.3) |
| *weighted* |  | (84.3) | (15.7) |
| age>50&<=75% | 1960 | 1632 (83.3) | 328 (16.7) |
| *weighted* |  | (83.5) | (16.5) |
| age>75% | 2039 | 1676 (82.2) | 363 (17.8) |
| *weighted* |  | (81.9) | (18.1) |
| **sex_at_birth** |  |  |  |
| F | 4105 | 3476 (84.7) | 629 (15.3) |
| *weighted* |  | (84.7) | (15.3) |
| M | 4488 | 3662 (81.6) | 826 (18.4) |
| *weighted* |  | (82.0) | (18.0) |
| **race.4level** |  |  |  |
| White | 5705 | 4710 (82.6) | 995 (17.4) |
| *weighted* |  | (82.8) | (17.2) |
| Black | 1217 | 1027 (84.4) | 190 (15.6) |
| *weighted* |  | (84.6) | (15.4) |
| Asian | 201 | 173 (86.1) | 28 (13.9) |
| *weighted* |  | (85.1) | (14.9) |
| Other/Mixed | 1470 | 1228 (83.5) | 242 (16.5) |
| *weighted* |  | (84.0) | (16.0) |
| **hisp** |  |  |  |
| No | 6901 | 5681 (82.3) | 1220 (17.7) |
| *weighted* |  | (82.4) | (17.6) |
| Yes | 1692 | 1457 (86.1) | 235 (13.9) |
| *weighted* |  | (86.3) | (13.7) |
| **household.income** |  |  |  |
| [<50K] | 2480 | 2102 (84.8) | 378 (15.2) |
| *weighted* |  | (84.4) | (15.6) |
| [>=50K & <100K] | 2460 | 2027 (82.4) | 433 (17.6) |
| *weighted* |  | (82.4) | (17.6) |
| [>=100K] | 3653 | 3009 (82.4) | 644 (17.6) |
| *weighted* |  | (83.0) | (17.0) |
| **high.educ** |  |  |  |
| < HS Diploma | 304 | 266 (87.5) | 38 (12.5) |
| *weighted* |  | (88.3) | (11.7) |
| HS Diploma/GED | 700 | 609 (87.0) | 91 (13.0) |
| *weighted* |  | (86.9) | (13.1) |
| Some College | 2196 | 1828 (83.2) | 368 (16.8) |
| *weighted* |  | (82.8) | (17.2) |
| Bachelor | 2251 | 1861 (82.7) | 390 (17.3) |
| *weighted* |  | (82.8) | (17.2) |
| Post Graduate Degree | 3142 | 2574 (81.9) | 568 (18.1) |
| *weighted* |  | (82.4) | (17.6) |
| **pubertal_dev_collapsed** |  |  |  |
| Pre | 4409 | 3596 (81.6) | 813 (18.4) |
| *weighted* |  | (81.6) | (18.4) |
| Early | 2058 | 1723 (83.7) | 335 (16.3) |
| *weighted* |  | (84.2) | (15.8) |
| Mid | 2000 | 1708 (85.4) | 292 (14.6) |
| *weighted* |  | (85.2) | (14.8) |
| Late/Post | 126 | 111 (88.1) | 15 (11.9) |
| *weighted* |  | (89.9) | (10.1) |

*Supplementary Table 36. Contingency table for variable of interest: Physical activity (strengthening). Independent sample used for statistics.*

| **Demographics** | **n** | **Below guidelines** | **Within guidelines** |
| --- | --- | --- | --- |
| **age_quartiles** |  |  |  |
| age<=25% | 2308 | 1635 (70.8) | 673 (29.2) |
| *weighted* |  | (71.1) | (28.9) |
| age>25&<=50% | 2286 | 1608 (70.3) | 678 (29.7) |
| *weighted* |  | (69.9) | (30.1) |
| age>50&<=75% | 1960 | 1325 (67.6) | 635 (32.4) |
| *weighted* |  | (67.7) | (32.3) |
| age>75% | 2039 | 1374 (67.4) | 665 (32.6) |
| *weighted* |  | (67.1) | (32.9) |
| **sex_at_birth** |  |  |  |
| F | 4105 | 2934 (71.5) | 1171 (28.5) |
| *weighted* |  | (70.9) | (29.1) |
| M | 4488 | 3008 (67.0) | 1480 (33.0) |
| *weighted* |  | (67.2) | (32.8) |
| **race.4level** |  |  |  |
| White | 5705 | 4074 (71.4) | 1631 (28.6) |
| *weighted* |  | (71.1) | (28.9) |
| Black | 1217 | 716 (58.8) | 501 (41.2) |
| *weighted* |  | (58.7) | (41.3) |
| Asian | 201 | 144 (71.6) | 57 (28.4) |
| *weighted* |  | (72.1) | (27.9) |
| Other/Mixed | 1470 | 1008 (68.6) | 462 (31.4) |
| *weighted* |  | (67.5) | (32.5) |
| **hisp** |  |  |  |
| No | 6901 | 4786 (69.4) | 2115 (30.6) |
| *weighted* |  | (69.7) | (30.3) |
| Yes | 1692 | 1156 (68.3) | 536 (31.7) |
| *weighted* |  | (66.9) | (33.1) |
| **household.income** |  |  |  |
| [<50K] | 2480 | 1610 (64.9) | 870 (35.1) |
| *weighted* |  | (65.8) | (34.2) |
| [>=50K & <100K] | 2460 | 1718 (69.8) | 742 (30.2) |
| *weighted* |  | (70.6) | (29.4) |
| [>=100K] | 3653 | 2614 (71.6) | 1039 (28.4) |
| *weighted* |  | (71.4) | (28.6) |
| **high.educ** |  |  |  |
| < HS Diploma | 304 | 196 (64.5) | 108 (35.5) |
| *weighted* |  | (64.7) | (35.3) |
| HS Diploma/GED | 700 | 439 (62.7) | 261 (37.3) |
| *weighted* |  | (63.2) | (36.8) |
| Some College | 2196 | 1461 (66.5) | 735 (33.5) |
| *weighted* |  | (66.4) | (33.6) |
| Bachelor | 2251 | 1615 (71.7) | 636 (28.3) |
| *weighted* |  | (72.3) | (27.7) |
| Post Graduate Degree | 3142 | 2231 (71.0) | 911 (29.0) |
| *weighted* |  | (71.3) | (28.7) |
| **pubertal_dev_collapsed** |  |  |  |
| Pre | 4409 | 3075 (69.7) | 1334 (30.3) |
| *weighted* |  | (69.8) | (30.2) |
| Early | 2058 | 1402 (68.1) | 656 (31.9) |
| *weighted* |  | (68.5) | (31.5) |
| Mid | 2000 | 1378 (68.9) | 622 (31.1) |
| *weighted* |  | (67.9) | (32.1) |
| Late/Post | 126 | 87 (69.0) | 39 (31.0) |
| *weighted* |  | (70.1) | (29.9) |

*Supplementary Table 37. Contingency table for variable of interest: Sports/activities involvement. Independent sample used for statistics.*

| **Demographics** | **n** | **No Participation** | **Some Participation** |
| --- | --- | --- | --- |
| **age_quartiles** |  |  |  |
| age<=25% | 2308 | 570 (24.7) | 1738 (75.3) |
| *weighted* |  | (28.6) | (71.4) |
| age>25&<=50% | 2286 | 505 (22.1) | 1781 (77.9) |
| *weighted* |  | (25.2) | (74.8) |
| age>50&<=75% | 1960 | 428 (21.8) | 1532 (78.2) |
| *weighted* |  | (25.7) | (74.3) |
| age>75% | 2039 | 415 (20.4) | 1624 (79.6) |
| *weighted* |  | (23.2) | (76.8) |
| **sex_at_birth** |  |  |  |
| F | 4105 | 989 (24.1) | 3116 (75.9) |
| *weighted* |  | (27.7) | (72.3) |
| M | 4488 | 929 (20.7) | 3559 (79.3) |
| *weighted* |  | (23.8) | (76.2) |
| **race.4level** |  |  |  |
| White | 5705 | 1010 (17.7) | 4695 (82.3) |
| *weighted* |  | (21.5) | (78.5) |
| Black | 1217 | 470 (38.6) | 747 (61.4) |
| *weighted* |  | (42.1) | (57.9) |
| Asian | 201 | 35 (17.4) | 166 (82.6) |
| *weighted* |  | (19.4) | (80.6) |
| Other/Mixed | 1470 | 403 (27.4) | 1067 (72.6) |
| *weighted* |  | (33.2) | (66.8) |
| **hisp** |  |  |  |
| No | 6901 | 1401 (20.3) | 5500 (79.7) |
| *weighted* |  | (23.0) | (77.0) |
| Yes | 1692 | 517 (30.6) | 1175 (69.4) |
| *weighted* |  | (34.6) | (65.4) |
| **household.income** |  |  |  |
| [<50K] | 2480 | 1017 (41.0) | 1463 (59.0) |
| *weighted* |  | (40.7) | (59.3) |
| [>=50K & <100K] | 2460 | 557 (22.6) | 1903 (77.4) |
| *weighted* |  | (23.0) | (77.0) |
| [>=100K] | 3653 | 344 ( 9.4) | 3309 (90.6) |
| *weighted* |  | ( 9.6) | (90.4) |
| **high.educ** |  |  |  |
| < HS Diploma | 304 | 155 (51.0) | 149 (49.0) |
| *weighted* |  | (52.9) | (47.1) |
| HS Diploma/GED | 700 | 318 (45.4) | 382 (54.6) |
| *weighted* |  | (45.7) | (54.3) |
| Some College | 2196 | 756 (34.4) | 1440 (65.6) |
| *weighted* |  | (35.9) | (64.1) |
| Bachelor | 2251 | 385 (17.1) | 1866 (82.9) |
| *weighted* |  | (18.9) | (81.1) |
| Post Graduate Degree | 3142 | 304 ( 9.7) | 2838 (90.3) |
| *weighted* |  | (11.3) | (88.7) |
| **pubertal_dev_collapsed** |  |  |  |
| Pre | 4409 | 849 (19.3) | 3560 (80.7) |
| *weighted* |  | (22.7) | (77.3) |
| Early | 2058 | 437 (21.2) | 1621 (78.8) |
| *weighted* |  | (24.4) | (75.6) |
| Mid | 2000 | 597 (29.8) | 1403 (70.2) |
| *weighted* |  | (32.8) | (67.2) |
| Late/Post | 126 | 35 (27.8) | 91 (72.2) |
| *weighted* |  | (27.5) | (72.5) |

*Supplementary Table 38. Contingency table for variable of interest: Weight status (for BMI). Independent sample used for statistics.*

| **Demographics** | **n** | **Underweight** | **Healthy Weight** | **Overweight** | **Obese** |
| --- | --- | --- | --- | --- | --- |
| **age_quartiles** |  |  |  |  |  |
| age<=25% | 2301 | 80 ( 3.5) | 1511 (65.7) | 320 (13.9) | 390 (16.9) |
| *weighted* |  | ( 3.3) | (63.6) | (14.3) | (18.9) |
| age>25&<=50% | 2280 | 89 ( 3.9) | 1491 (65.4) | 328 (14.4) | 372 (16.3) |
| *weighted* |  | ( 3.7) | (63.9) | (15.2) | (17.2) |
| age>50&<=75% | 1951 | 72 ( 3.7) | 1253 (64.2) | 294 (15.1) | 332 (17.0) |
| *weighted* |  | ( 3.6) | (62.5) | (15.6) | (18.3) |
| age>75% | 2036 | 85 ( 4.2) | 1329 (65.3) | 307 (15.1) | 315 (15.5) |
| *weighted* |  | ( 3.9) | (63.4) | (15.6) | (17.0) |
| **sex_at_birth** |  |  |  |  |  |
| F | 4096 | 168 ( 4.1) | 2697 (65.8) | 590 (14.4) | 641 (15.6) |
| *weighted* |  | ( 3.8) | (64.4) | (15.1) | (16.7) |
| M | 4472 | 158 ( 3.5) | 2887 (64.6) | 659 (14.7) | 768 (17.2) |
| *weighted* |  | ( 3.5) | (62.4) | (15.2) | (19.0) |
| **race.4level** |  |  |  |  |  |
| White | 5690 | 235 ( 4.1) | 3946 (69.3) | 785 (13.8) | 724 (12.7) |
| *weighted* |  | ( 3.9) | (66.8) | (14.5) | (14.7) |
| Black | 1211 | 31 ( 2.6) | 613 (50.6) | 200 (16.5) | 367 (30.3) |
| *weighted* |  | ( 2.5) | (50.1) | (16.6) | (30.8) |
| Asian | 200 | 12 ( 6.0) | 144 (72.0) | 24 (12.0) | 20 (10.0) |
| *weighted* |  | ( 5.0) | (70.7) | (13.8) | (10.5) |
| Other/Mixed | 1467 | 48 ( 3.3) | 881 (60.1) | 240 (16.4) | 298 (20.3) |
| *weighted* |  | ( 2.9) | (56.6) | (17.2) | (23.3) |
| **hisp** |  |  |  |  |  |
| No | 6879 | 278 ( 4.0) | 4705 (68.4) | 918 (13.3) | 978 (14.2) |
| *weighted* |  | ( 3.9) | (67.1) | (13.7) | (15.3) |
| Yes | 1689 | 48 ( 2.8) | 879 (52.0) | 331 (19.6) | 431 (25.5) |
| *weighted* |  | ( 2.8) | (51.2) | (19.8) | (26.2) |
| **household.income** |  |  |  |  |  |
| [<50K] | 2471 | 63 ( 2.5) | 1315 (53.2) | 427 (17.3) | 666 (27.0) |
| *weighted* |  | ( 2.6) | (54.0) | (17.6) | (25.8) |
| [>=50K & <100K] | 2455 | 99 ( 4.0) | 1578 (64.3) | 373 (15.2) | 405 (16.5) |
| *weighted* |  | ( 4.0) | (64.7) | (14.8) | (16.4) |
| [>=100K] | 3642 | 164 ( 4.5) | 2691 (73.9) | 449 (12.3) | 338 ( 9.3) |
| *weighted* |  | ( 4.6) | (73.8) | (12.4) | ( 9.3) |
| **high.educ** |  |  |  |  |  |
| < HS Diploma | 303 | 7 ( 2.3) | 133 (43.9) | 62 (20.5) | 101 (33.3) |
| *weighted* |  | ( 2.3) | (43.4) | (22.0) | (32.2) |
| HS Diploma/GED | 700 | 19 ( 2.7) | 344 (49.1) | 102 (14.6) | 235 (33.6) |
| *weighted* |  | ( 2.9) | (48.6) | (14.4) | (34.1) |
| Some College | 2185 | 68 ( 3.1) | 1224 (56.0) | 383 (17.5) | 510 (23.3) |
| *weighted* |  | ( 3.0) | (56.4) | (17.5) | (23.1) |
| Bachelor | 2245 | 90 ( 4.0) | 1563 (69.6) | 318 (14.2) | 274 (12.2) |
| *weighted* |  | ( 3.9) | (68.0) | (14.9) | (13.2) |
| Post Graduate Degree | 3135 | 142 ( 4.5) | 2320 (74.0) | 384 (12.2) | 289 ( 9.2) |
| *weighted* |  | ( 4.5) | (73.7) | (12.4) | ( 9.4) |
| **pubertal_dev_collapsed** |  |  |  |  |  |
| Pre | 4396 | 219 ( 5.0) | 3153 (71.7) | 513 (11.7) | 511 (11.6) |
| *weighted* |  | ( 4.9) | (69.8) | (12.2) | (13.1) |
| Early | 2051 | 69 ( 3.4) | 1309 (63.8) | 320 (15.6) | 353 (17.2) |
| *weighted* |  | ( 3.3) | (63.0) | (15.7) | (18.0) |
| Mid | 1996 | 38 ( 1.9) | 1079 (54.1) | 375 (18.8) | 504 (25.3) |
| *weighted* |  | ( 1.8) | (53.3) | (18.9) | (26.0) |
| Late/Post | 125 | 0 ( 0.0) | 43 (34.4) | 41 (32.8) | 41 (32.8) |
| *weighted* |  | ( 0.0) | (35.1) | (35.2) | (29.7) |

*Supplementary Table 39. Contingency table for variable of interest: Total medical problems. Independent sample used for statistics.*

| **Demographics** | **n** | **None** | **One problem** | **Two or more problems** |
| --- | --- | --- | --- | --- |
| **age_quartiles** |  |  |  |  |
| age<=25% | 2300 | 667 (29.0) | 735 (32.0) | 898 (39.0) |
| *weighted* |  | (28.3) | (31.3) | (40.4) |
| age>25&<=50% | 2279 | 664 (29.1) | 683 (30.0) | 932 (40.9) |
| *weighted* |  | (29.5) | (28.9) | (41.7) |
| age>50&<=75% | 1948 | 573 (29.4) | 609 (31.3) | 766 (39.3) |
| *weighted* |  | (28.3) | (30.5) | (41.2) |
| age>75% | 2034 | 579 (28.5) | 575 (28.3) | 880 (43.3) |
| *weighted* |  | (28.4) | (27.5) | (44.1) |
| **sex_at_birth** |  |  |  |  |
| F | 4091 | 1236 (30.2) | 1255 (30.7) | 1600 (39.1) |
| *weighted* |  | (29.6) | (29.8) | (40.6) |
| M | 4470 | 1247 (27.9) | 1347 (30.1) | 1876 (42.0) |
| *weighted* |  | (27.7) | (29.3) | (43.0) |
| **race.4level** |  |  |  |  |
| White | 5685 | 1579 (27.8) | 1756 (30.9) | 2350 (41.3) |
| *weighted* |  | (27.2) | (29.9) | (42.9) |
| Black | 1211 | 384 (31.7) | 360 (29.7) | 467 (38.6) |
| *weighted* |  | (32.3) | (29.3) | (38.4) |
| Asian | 201 | 68 (33.8) | 56 (27.9) | 77 (38.3) |
| *weighted* |  | (33.7) | (27.4) | (38.9) |
| Other/Mixed | 1464 | 452 (30.9) | 430 (29.4) | 582 (39.8) |
| *weighted* |  | (30.9) | (28.5) | (40.6) |
| **hisp** |  |  |  |  |
| No | 6876 | 1925 (28.0) | 2120 (30.8) | 2831 (41.2) |
| *weighted* |  | (27.5) | (29.7) | (42.8) |
| Yes | 1685 | 558 (33.1) | 482 (28.6) | 645 (38.3) |
| *weighted* |  | (32.3) | (28.9) | (38.8) |
| **household.income** |  |  |  |  |
| [<50K] | 2462 | 809 (32.9) | 692 (28.1) | 961 (39.0) |
| *weighted* |  | (31.6) | (27.5) | (40.9) |
| [>=50K & <100K] | 2454 | 689 (28.1) | 745 (30.4) | 1020 (41.6) |
| *weighted* |  | (27.1) | (29.9) | (43.0) |
| [>=100K] | 3645 | 985 (27.0) | 1165 (32.0) | 1495 (41.0) |
| *weighted* |  | (26.3) | (31.8) | (41.9) |
| **high.educ** |  |  |  |  |
| < HS Diploma | 303 | 129 (42.6) | 78 (25.7) | 96 (31.7) |
| *weighted* |  | (42.1) | (26.9) | (31.0) |
| HS Diploma/GED | 697 | 260 (37.3) | 188 (27.0) | 249 (35.7) |
| *weighted* |  | (35.0) | (27.8) | (37.1) |
| Some College | 2184 | 650 (29.8) | 627 (28.7) | 907 (41.5) |
| *weighted* |  | (28.6) | (27.8) | (43.6) |
| Bachelor | 2247 | 620 (27.6) | 718 (32.0) | 909 (40.5) |
| *weighted* |  | (27.0) | (31.0) | (42.0) |
| Post Graduate Degree | 3130 | 824 (26.3) | 991 (31.7) | 1315 (42.0) |
| *weighted* |  | (26.0) | (31.0) | (43.1) |
| **pubertal_dev_collapsed** |  |  |  |  |
| Pre | 4394 | 1300 (29.6) | 1377 (31.3) | 1717 (39.1) |
| *weighted* |  | (29.3) | (30.7) | (40.0) |
| Early | 2050 | 541 (26.4) | 623 (30.4) | 886 (43.2) |
| *weighted* |  | (26.2) | (29.2) | (44.6) |
| Mid | 1991 | 609 (30.6) | 567 (28.5) | 815 (40.9) |
| *weighted* |  | (29.9) | (28.0) | (42.1) |
| Late/Post | 126 | 33 (26.2) | 35 (27.8) | 58 (46.0) |
| *weighted* |  | (25.7) | (24.8) | (49.6) |

*Supplementary Table 40. Contingency table for variable of interest: Traumatic Brain Injury (TBI). Independent sample used for statistics.*

| **Demographics** | **n** | **Improbable TBI** | **Possible mild TBI** | **TBI** |
| --- | --- | --- | --- | --- |
| **age_quartiles** |  |  |  |  |
| age<=25% | 2308 | 2234 (96.8) | 45 ( 1.9) | 29 ( 1.3) |
| *weighted* |  | (96.7) | ( 1.9) | ( 1.3) |
| age>25&<=50% | 2286 | 2199 (96.2) | 61 ( 2.7) | 26 ( 1.1) |
| *weighted* |  | (96.4) | ( 2.5) | ( 1.2) |
| age>50&<=75% | 1960 | 1882 (96.0) | 58 ( 3.0) | 20 ( 1.0) |
| *weighted* |  | (96.4) | ( 2.6) | ( 1.0) |
| age>75% | 2039 | 1942 (95.2) | 65 ( 3.2) | 32 ( 1.6) |
| *weighted* |  | (95.6) | ( 3.0) | ( 1.4) |
| **sex_at_birth** |  |  |  |  |
| F | 4105 | 3974 (96.8) | 87 ( 2.1) | 44 ( 1.1) |
| *weighted* |  | (97.0) | ( 1.9) | ( 1.1) |
| M | 4488 | 4283 (95.4) | 142 ( 3.2) | 63 ( 1.4) |
| *weighted* |  | (95.6) | ( 3.0) | ( 1.3) |
| **race.4level** |  |  |  |  |
| White | 5705 | 5463 (95.8) | 166 ( 2.9) | 76 ( 1.3) |
| *weighted* |  | (95.9) | ( 2.7) | ( 1.3) |
| Black | 1217 | 1184 (97.3) | 22 ( 1.8) | 11 ( 0.9) |
| *weighted* |  | (97.6) | ( 1.6) | ( 0.8) |
| Asian | 201 | 195 (97.0) | <10 | <10 |
| *weighted* |  | (97.6) |  |  |
| Other/Mixed | 1470 | 1415 (96.3) | 36 ( 2.4) | 19 ( 1.3) |
| *weighted* |  | (96.6) | ( 2.1) | ( 1.2) |
| **hisp** |  |  |  |  |
| No | 6901 | 6620 (95.9) | 199 ( 2.9) | 82 ( 1.2) |
| *weighted* |  | (96.1) | ( 2.7) | ( 1.2) |
| Yes | 1692 | 1637 (96.7) | 30 ( 1.8) | 25 ( 1.5) |
| *weighted* |  | (97.0) | ( 1.7) | ( 1.3) |
| **household.income** |  |  |  |  |
| [<50K] | 2480 | 2410 (97.2) | 46 ( 1.9) | 24 ( 1.0) |
| *weighted* |  | (97.1) | ( 1.8) | ( 1.1) |
| [>=50K & <100K] | 2460 | 2368 (96.3) | 64 ( 2.6) | 28 ( 1.1) |
| *weighted* |  | (96.2) | ( 2.7) | ( 1.2) |
| [>=100K] | 3653 | 3479 (95.2) | 119 ( 3.3) | 55 ( 1.5) |
| *weighted* |  | (95.4) | ( 3.2) | ( 1.4) |
| **high.educ** |  |  |  |  |
| < HS Diploma | 304 | 294 (96.7) | <10 | <10 |
| *weighted* |  | (96.4) |  |  |
| HS Diploma/GED | 700 | 682 (97.4) | 13 ( 1.9) | <10 |
| *weighted* |  | (97.5) | ( 1.8) |  |
| Some College | 2196 | 2127 (96.9) | 46 ( 2.1) | 23 ( 1.0) |
| *weighted* |  | (96.8) | ( 2.1) | ( 1.1) |
| Bachelor | 2251 | 2151 (95.6) | 66 ( 2.9) | 34 ( 1.5) |
| *weighted* |  | (96.0) | ( 2.6) | ( 1.4) |
| Post Graduate Degree | 3142 | 3003 (95.6) | 98 ( 3.1) | 41 ( 1.3) |
| *weighted* |  | (95.7) | ( 3.0) | ( 1.3) |
| **pubertal_dev_collapsed** |  |  |  |  |
| Pre | 4409 | 4232 (96.0) | 124 ( 2.8) | 53 ( 1.2) |
| *weighted* |  | (96.2) | ( 2.7) | ( 1.1) |
| Early | 2058 | 1975 (96.0) | 52 ( 2.5) | 31 ( 1.5) |
| *weighted* |  | (96.0) | ( 2.3) | ( 1.7) |
| Mid | 2000 | 1928 (96.4) | 50 ( 2.5) | 22 ( 1.1) |
| *weighted* |  | (96.7) | ( 2.3) | ( 1.0) |
| Late/Post | 126 | 122 (96.8) | <10 | <10 |
| *weighted* |  | (96.7) |  |  |
