## Supplemental Tables 41-60 for "A Comprehensive Overview of the Physical Health of the Adolescent Brain Cognitive Development Study (ABCD) Cohort at Baseline"

*Supplementary Table 41. DV: Birth weight. Chi-squared test and pairwise post-hoc comparisons for associations with the sociodemographic factors.*

| **Demographic** | **X.squared** | **DF** | **p.value** |
| --- | --- | --- | --- |
| **sex_at_birth** | 59.9 | 3 | 6.14e-13 |
| **race.4level** | 60.5 |  | 5e-04 |
| **hisp** | 21.4 | 3 | 8.88e-05 |
| **household.income** | 22.3 | 6 | 0.00106 |
| **high.educ** | 24.9 |  | 0.0175 |
| **Measure** | **sex_at_birth** | **StdResidual** | **P-value** |
| **Very low** | F | 2.49 | 1.02e-01 |
| **Low** | F | 3.37 | 6.02e-03 |
| **Normal** | F | 1.73 | 6.63e-01 |
| **High** | F | -6.89 | 4.50e-11 |
| **Very low** | M | -2.49 | 1.02e-01 |
| **Low** | M | -3.37 | 6.02e-03 |
| **Normal** | M | -1.73 | 6.63e-01 |
| **High** | M | 6.89 | 4.50e-11 |
| **Measure** | **race.4level** | **StdResidual** | **P-value** |
| **Very low** | White | -0.18350 | 1.00e+00 |
| **Low** | White | -3.17923 | 2.36e-02 |
| **Normal** | White | -0.00517 | 1.00e+00 |
| **High** | White | 3.51689 | 6.99e-03 |
| **Very low** | Black | 2.57712 | 1.59e-01 |
| **Low** | Black | 5.52238 | 5.35e-07 |
| **Normal** | Black | -1.64725 | 1.00e+00 |
| **High** | Black | -4.48358 | 1.17e-04 |
| **Very low** | Asian | -1.28571 | 1.00e+00 |
| **Low** | Asian | 0.63297 | 1.00e+00 |
| **Normal** | Asian | 0.99474 | 1.00e+00 |
| **High** | Asian | -1.68568 | 1.00e+00 |
| **Very low** | Other/Mixed | -1.66833 | 1.00e+00 |
| **Low** | Other/Mixed | -1.40429 | 1.00e+00 |
| **Normal** | Other/Mixed | 1.15236 | 1.00e+00 |
| **High** | Other/Mixed | 0.42437 | 1.00e+00 |
| **Measure** | **hisp** | **StdResidual** | **P-value** |
| **Very low** | No | -2.63 | 0.06930 |
| **Low** | No | 3.30 | 0.00767 |
| **Normal** | No | -3.18 | 0.01200 |
| **High** | No | 1.72 | 0.68200 |
| **Very low** | Yes | 2.63 | 0.06930 |
| **Low** | Yes | -3.30 | 0.00767 |
| **Normal** | Yes | 3.18 | 0.01200 |
| **High** | Yes | -1.72 | 0.68200 |
| **Measure** | **household.income** | **StdResidual** | **P-value** |
| **Very low** | [<50K] | 2.0117 | 0.5310 |
| **Low** | [<50K] | 2.0981 | 0.4310 |
| **Normal** | [<50K] | 0.0972 | 1.0000 |
| **High** | [<50K] | -3.0485 | 0.0276 |
| **Very low** | [>=50K & <100K] | 1.1929 | 1.0000 |
| **Low** | [>=50K & <100K] | -1.7954 | 0.8710 |
| **Normal** | [>=50K & <100K] | -0.0812 | 1.0000 |
| **High** | [>=50K & <100K] | 1.6883 | 1.0000 |
| **Very low** | [>=100K] | -2.9482 | 0.0384 |
| **Low** | [>=100K] | -0.2983 | 1.0000 |
| **Normal** | [>=100K] | -0.0156 | 1.0000 |
| **High** | [>=100K] | 1.2741 | 1.0000 |
| **Measure** | **high.educ** | **StdResidual** | **P-value** |
| **Very low** | < HS Diploma | 0.8235 | 1.0000 |
| **Low** | < HS Diploma | -0.2926 | 1.0000 |
| **Normal** | < HS Diploma | 0.2618 | 1.0000 |
| **High** | < HS Diploma | -0.3108 | 1.0000 |
| **Very low** | HS Diploma/GED | 0.3604 | 1.0000 |
| **Low** | HS Diploma/GED | 0.4120 | 1.0000 |
| **Normal** | HS Diploma/GED | 1.1913 | 1.0000 |
| **High** | HS Diploma/GED | -2.2413 | 0.5000 |
| **Very low** | Some College | 2.0166 | 0.8750 |
| **Low** | Some College | 2.4894 | 0.2560 |
| **Normal** | Some College | -2.4539 | 0.2830 |
| **High** | Some College | 0.1239 | 1.0000 |
| **Very low** | Bachelor | -1.8125 | 1.0000 |
| **Low** | Bachelor | 1.0548 | 1.0000 |
| **Normal** | Bachelor | -0.4431 | 1.0000 |
| **High** | Bachelor | 0.0504 | 1.0000 |
| **Very low** | Post Graduate Degree | -0.7160 | 1.0000 |
| **Low** | Post Graduate Degree | -3.3568 | 0.0158 |
| **Normal** | Post Graduate Degree | 1.8439 | 1.0000 |
| **High** | Post Graduate Degree | 1.2689 | 1.0000 |

*Supplementary Table 42. DV: Prematurity. Chi-squared test and pairwise post-hoc comparisons for associations with the sociodemographic factors.*

| **Demographic** | **X.squared** | **DF** | **p.value** |
| --- | --- | --- | --- |
| **sex_at_birth** | 0.783 | 4 | 0.941 |
| **race.4level** | 27.6 |  | 0.011 |
| **hisp** | 14.4 |  | 0.006 |
| **household.income** | 37.1 |  | 5e-04 |
| **high.educ** | 53.1 |  | 5e-04 |
| **Measure** | **sex_at_birth** | **StdResidual** | **P-value** |
| **Term** | F | 0.26691 | 1 |
| **Late Preterm** | F | 0.00304 | 1 |
| **Moderate Preterm** | F | -0.59628 | 1 |
| **Very Preterm** | F | 0.34399 | 1 |
| **Extremely Preterm** | F | -0.55653 | 1 |
| **Term** | M | -0.26691 | 1 |
| **Late Preterm** | M | -0.00304 | 1 |
| **Moderate Preterm** | M | 0.59628 | 1 |
| **Very Preterm** | M | -0.34399 | 1 |
| **Extremely Preterm** | M | 0.55653 | 1 |
| **Measure** | **race.4level** | **StdResidual** | **P-value** |
| **Term** | White | -1.492 | 1.0000 |
| **Late Preterm** | White | 1.198 | 1.0000 |
| **Moderate Preterm** | White | 0.238 | 1.0000 |
| **Very Preterm** | White | 3.010 | 0.0523 |
| **Extremely Preterm** | White | -2.529 | 0.2290 |
| **Term** | Black | -0.763 | 1.0000 |
| **Late Preterm** | Black | 0.272 | 1.0000 |
| **Moderate Preterm** | Black | 0.610 | 1.0000 |
| **Very Preterm** | Black | -1.312 | 1.0000 |
| **Extremely Preterm** | Black | 3.427 | 0.0122 |
| **Term** | Asian | 1.884 | 1.0000 |
| **Late Preterm** | Asian | -1.472 | 1.0000 |
| **Moderate Preterm** | Asian | -0.903 | 1.0000 |
| **Very Preterm** | Asian | -1.157 | 1.0000 |
| **Extremely Preterm** | Asian | 0.553 | 1.0000 |
| **Term** | Other/Mixed | 1.849 | 1.0000 |
| **Late Preterm** | Other/Mixed | -1.182 | 1.0000 |
| **Moderate Preterm** | Other/Mixed | -0.516 | 1.0000 |
| **Very Preterm** | Other/Mixed | -2.096 | 0.7220 |
| **Extremely Preterm** | Other/Mixed | -0.254 | 1.0000 |
| **Measure** | **hisp** | **StdResidual** | **P-value** |
| **Term** | No | -1.700 | 0.8910 |
| **Late Preterm** | No | 2.703 | 0.0687 |
| **Moderate Preterm** | No | -1.032 | 1.0000 |
| **Very Preterm** | No | 0.831 | 1.0000 |
| **Extremely Preterm** | No | -2.382 | 0.1720 |
| **Term** | Yes | 1.700 | 0.8910 |
| **Late Preterm** | Yes | -2.703 | 0.0687 |
| **Moderate Preterm** | Yes | 1.032 | 1.0000 |
| **Very Preterm** | Yes | -0.831 | 1.0000 |
| **Extremely Preterm** | Yes | 2.382 | 0.1720 |
| **Measure** | **household.income** | **StdResidual** | **P-value** |
| **Term** | [<50K] | 1.7894 | 1.000000 |
| **Late Preterm** | [<50K] | -3.7552 | 0.002600 |
| **Moderate Preterm** | [<50K] | 1.6698 | 1.000000 |
| **Very Preterm** | [<50K] | -0.0925 | 1.000000 |
| **Extremely Preterm** | [<50K] | 4.3925 | 0.000168 |
| **Term** | [>=50K & <100K] | -0.5689 | 1.000000 |
| **Late Preterm** | [>=50K & <100K] | 0.9058 | 1.000000 |
| **Moderate Preterm** | [>=50K & <100K] | -0.5165 | 1.000000 |
| **Very Preterm** | [>=50K & <100K] | 1.0192 | 1.000000 |
| **Extremely Preterm** | [>=50K & <100K] | -1.4802 | 1.000000 |
| **Term** | [>=100K] | -1.1404 | 1.000000 |
| **Late Preterm** | [>=100K] | 2.6569 | 0.118000 |
| **Moderate Preterm** | [>=100K] | -1.0773 | 1.000000 |
| **Very Preterm** | [>=100K] | -0.8469 | 1.000000 |
| **Extremely Preterm** | [>=100K] | -2.7228 | 0.097100 |
| **Measure** | **high.educ** | **StdResidual** | **P-value** |
| **Term** | < HS Diploma | 2.769 | 0.14100 |
| **Late Preterm** | < HS Diploma | -3.042 | 0.05870 |
| **Moderate Preterm** | < HS Diploma | -0.979 | 1.00000 |
| **Very Preterm** | < HS Diploma | 0.510 | 1.00000 |
| **Extremely Preterm** | < HS Diploma | 1.007 | 1.00000 |
| **Term** | HS Diploma/GED | 0.617 | 1.00000 |
| **Late Preterm** | HS Diploma/GED | -1.773 | 1.00000 |
| **Moderate Preterm** | HS Diploma/GED | 0.218 | 1.00000 |
| **Very Preterm** | HS Diploma/GED | 0.470 | 1.00000 |
| **Extremely Preterm** | HS Diploma/GED | 4.030 | 0.00139 |
| **Term** | Some College | -2.563 | 0.26000 |
| **Late Preterm** | Some College | 1.672 | 1.00000 |
| **Moderate Preterm** | Some College | 2.626 | 0.21600 |
| **Very Preterm** | Some College | -0.574 | 1.00000 |
| **Extremely Preterm** | Some College | 0.593 | 1.00000 |
| **Term** | Bachelor | -2.238 | 0.63000 |
| **Late Preterm** | Bachelor | 2.346 | 0.47400 |
| **Moderate Preterm** | Bachelor | 0.940 | 1.00000 |
| **Very Preterm** | Bachelor | 0.292 | 1.00000 |
| **Extremely Preterm** | Bachelor | -1.675 | 1.00000 |
| **Term** | Post Graduate Degree | 2.903 | 0.09240 |
| **Late Preterm** | Post Graduate Degree | -1.403 | 1.00000 |
| **Moderate Preterm** | Post Graduate Degree | -2.984 | 0.07100 |
| **Very Preterm** | Post Graduate Degree | -0.224 | 1.00000 |
| **Extremely Preterm** | Post Graduate Degree | -1.773 | 1.00000 |

*Supplementary Table 43. DV: Age at first rolling over (months). Chi-squared test and pairwise post-hoc comparisons for associations with the sociodemographic factors.*

| **Demographic** | **X.squared** | **DF** | **p.value** |
| --- | --- | --- | --- |
| **sex_at_birth** | 0.00516 | 1 | 0.943 |
| **race.4level** | 0.348 |  | 0.952 |
| **hisp** | 1.03 | 1 | 0.31 |
| **household.income** | 6.71 | 2 | 0.0349 |
| **high.educ** | 3.87 |  | 0.409 |
| **Measure** | **sex_at_birth** | **StdResidual** | **P-value** |
| **Within guidelines** | F | 0.174 | 1 |
| **Beyond guidelines** | F | -0.174 | 1 |
| **Within guidelines** | M | -0.174 | 1 |
| **Beyond guidelines** | M | 0.174 | 1 |
| **Measure** | **race.4level** | **StdResidual** | **P-value** |
| **Within guidelines** | White | -0.403 | 1 |
| **Beyond guidelines** | White | 0.403 | 1 |
| **Within guidelines** | Black | -0.118 | 1 |
| **Beyond guidelines** | Black | 0.118 | 1 |
| **Within guidelines** | Asian | 0.243 | 1 |
| **Beyond guidelines** | Asian | -0.243 | 1 |
| **Within guidelines** | Other/Mixed | 0.519 | 1 |
| **Beyond guidelines** | Other/Mixed | -0.519 | 1 |
| **Measure** | **hisp** | **StdResidual** | **P-value** |
| **Within guidelines** | No | 1.14 | 1 |
| **Beyond guidelines** | No | -1.14 | 1 |
| **Within guidelines** | Yes | -1.14 | 1 |
| **Beyond guidelines** | Yes | 1.14 | 1 |
| **Measure** | **household.income** | **StdResidual** | **P-value** |
| **Within guidelines** | [<50K] | -2.581 | 0.0591 |
| **Beyond guidelines** | [<50K] | 2.581 | 0.0591 |
| **Within guidelines** | [>=50K & <100K] | 0.865 | 1.0000 |
| **Beyond guidelines** | [>=50K & <100K] | -0.865 | 1.0000 |
| **Within guidelines** | [>=100K] | 1.607 | 0.6480 |
| **Beyond guidelines** | [>=100K] | -1.607 | 0.6480 |
| **Measure** | **high.educ** | **StdResidual** | **P-value** |
| **Within guidelines** | < HS Diploma | -1.144 | 1.000 |
| **Beyond guidelines** | < HS Diploma | 1.144 | 1.000 |
| **Within guidelines** | HS Diploma/GED | -0.538 | 1.000 |
| **Beyond guidelines** | HS Diploma/GED | 0.538 | 1.000 |
| **Within guidelines** | Some College | -0.651 | 1.000 |
| **Beyond guidelines** | Some College | 0.651 | 1.000 |
| **Within guidelines** | Bachelor | -0.395 | 1.000 |
| **Beyond guidelines** | Bachelor | 0.395 | 1.000 |
| **Within guidelines** | Post Graduate Degree | 1.731 | 0.835 |
| **Beyond guidelines** | Post Graduate Degree | -1.731 | 0.835 |

*Supplementary Table 44. DV: Age at first sitting (months). Chi-squared test and pairwise post-hoc comparisons for associations with the sociodemographic factors.*

| **Demographic** | **X.squared** | **DF** | **p.value** |
| --- | --- | --- | --- |
| **sex_at_birth** | 9.02 | 1 | 0.00267 |
| **race.4level** | 8.97 |  | 0.0275 |
| **hisp** | 0.777 | 1 | 0.378 |
| **household.income** | 5.38 | 2 | 0.0678 |
| **high.educ** | 2.59 | 4 | 0.628 |
| **Measure** | **sex_at_birth** | **StdResidual** | **P-value** |
| **Within guidelines** | F | 3.06 | 0.0089 |
| **Beyond guidelines** | F | -3.06 | 0.0089 |
| **Within guidelines** | M | -3.06 | 0.0089 |
| **Beyond guidelines** | M | 3.06 | 0.0089 |
| **Measure** | **race.4level** | **StdResidual** | **P-value** |
| **Within guidelines** | White | -2.876 | 0.0322 |
| **Beyond guidelines** | White | 2.876 | 0.0322 |
| **Within guidelines** | Black | 1.023 | 1.0000 |
| **Beyond guidelines** | Black | -1.023 | 1.0000 |
| **Within guidelines** | Asian | 0.492 | 1.0000 |
| **Beyond guidelines** | Asian | -0.492 | 1.0000 |
| **Within guidelines** | Other/Mixed | 2.455 | 0.1130 |
| **Beyond guidelines** | Other/Mixed | -2.455 | 0.1130 |
| **Measure** | **hisp** | **StdResidual** | **P-value** |
| **Within guidelines** | No | 0.95 | 1 |
| **Beyond guidelines** | No | -0.95 | 1 |
| **Within guidelines** | Yes | -0.95 | 1 |
| **Beyond guidelines** | Yes | 0.95 | 1 |
| **Measure** | **household.income** | **StdResidual** | **P-value** |
| **Within guidelines** | [<50K] | -2.037 | 0.250 |
| **Beyond guidelines** | [<50K] | 2.037 | 0.250 |
| **Within guidelines** | [>=50K & <100K] | -0.172 | 1.000 |
| **Beyond guidelines** | [>=50K & <100K] | 0.172 | 1.000 |
| **Within guidelines** | [>=100K] | 2.051 | 0.242 |
| **Beyond guidelines** | [>=100K] | -2.051 | 0.242 |
| **Measure** | **high.educ** | **StdResidual** | **P-value** |
| **Within guidelines** | < HS Diploma | -1.2658 | 1 |
| **Beyond guidelines** | < HS Diploma | 1.2658 | 1 |
| **Within guidelines** | HS Diploma/GED | -0.3634 | 1 |
| **Beyond guidelines** | HS Diploma/GED | 0.3634 | 1 |
| **Within guidelines** | Some College | -0.4730 | 1 |
| **Beyond guidelines** | Some College | 0.4730 | 1 |
| **Within guidelines** | Bachelor | 0.0721 | 1 |
| **Beyond guidelines** | Bachelor | -0.0721 | 1 |
| **Within guidelines** | Post Graduate Degree | 1.0892 | 1 |
| **Beyond guidelines** | Post Graduate Degree | -1.0892 | 1 |

*Supplementary Table 45. DV: Age at first walking (months). Chi-squared test and pairwise post-hoc comparisons for associations with the sociodemographic factors.*

| **Demographic** | **X.squared** | **DF** | **p.value** |
| --- | --- | --- | --- |
| **sex_at_birth** | 11.5 | 1 | 0.000685 |
| **race.4level** | 2.19 |  | 0.547 |
| **hisp** | 0.683 | 1 | 0.409 |
| **household.income** | 7.04 | 2 | 0.0297 |
| **high.educ** | 4.55 | 4 | 0.336 |
| **Measure** | **sex_at_birth** | **StdResidual** | **P-value** |
| **Within guidelines** | F | 3.46 | 0.00218 |
| **Beyond guidelines** | F | -3.46 | 0.00218 |
| **Within guidelines** | M | -3.46 | 0.00218 |
| **Beyond guidelines** | M | 3.46 | 0.00218 |
| **Measure** | **race.4level** | **StdResidual** | **P-value** |
| **Within guidelines** | White | -1.0421 | 1 |
| **Beyond guidelines** | White | 1.0421 | 1 |
| **Within guidelines** | Black | -0.0891 | 1 |
| **Beyond guidelines** | Black | 0.0891 | 1 |
| **Within guidelines** | Asian | -0.1728 | 1 |
| **Beyond guidelines** | Asian | 0.1728 | 1 |
| **Within guidelines** | Other/Mixed | 1.4650 | 1 |
| **Beyond guidelines** | Other/Mixed | -1.4650 | 1 |
| **Measure** | **hisp** | **StdResidual** | **P-value** |
| **Within guidelines** | No | 0.903 | 1 |
| **Beyond guidelines** | No | -0.903 | 1 |
| **Within guidelines** | Yes | -0.903 | 1 |
| **Beyond guidelines** | Yes | 0.903 | 1 |
| **Measure** | **household.income** | **StdResidual** | **P-value** |
| **Within guidelines** | [<50K] | -2.62 | 0.0523 |
| **Beyond guidelines** | [<50K] | 2.62 | 0.0523 |
| **Within guidelines** | [>=50K & <100K] | 0.72 | 1.0000 |
| **Beyond guidelines** | [>=50K & <100K] | -0.72 | 1.0000 |
| **Within guidelines** | [>=100K] | 1.78 | 0.4520 |
| **Beyond guidelines** | [>=100K] | -1.78 | 0.4520 |
| **Measure** | **high.educ** | **StdResidual** | **P-value** |
| **Within guidelines** | < HS Diploma | -1.399 | 1 |
| **Beyond guidelines** | < HS Diploma | 1.399 | 1 |
| **Within guidelines** | HS Diploma/GED | -0.302 | 1 |
| **Beyond guidelines** | HS Diploma/GED | 0.302 | 1 |
| **Within guidelines** | Some College | -1.277 | 1 |
| **Beyond guidelines** | Some College | 1.277 | 1 |
| **Within guidelines** | Bachelor | 1.015 | 1 |
| **Beyond guidelines** | Bachelor | -1.015 | 1 |
| **Within guidelines** | Post Graduate Degree | 0.979 | 1 |
| **Beyond guidelines** | Post Graduate Degree | -0.979 | 1 |

*Supplementary Table 46. DV: Age at first word (months). Chi-squared test and pairwise post-hoc comparisons for associations with the sociodemographic factors.*

| **Demographic** | **X.squared** | **DF** | **p.value** |
| --- | --- | --- | --- |
| **sex_at_birth** | 50.4 | 1 | 1.26e-12 |
| **race.4level** | 10.7 | 3 | 0.0132 |
| **hisp** | 0.462 | 1 | 0.497 |
| **household.income** | 4.67 | 2 | 0.0969 |
| **high.educ** | 8.29 | 4 | 0.0815 |
| **Measure** | **sex_at_birth** | **StdResidual** | **P-value** |
| **Within guidelines** | F | 7.14 | 3.87e-12 |
| **Beyond guidelines** | F | -7.14 | 3.87e-12 |
| **Within guidelines** | M | -7.14 | 3.87e-12 |
| **Beyond guidelines** | M | 7.14 | 3.87e-12 |
| **Measure** | **race.4level** | **StdResidual** | **P-value** |
| **Within guidelines** | White | -2.04 | 0.330 |
| **Beyond guidelines** | White | 2.04 | 0.330 |
| **Within guidelines** | Black | 1.99 | 0.369 |
| **Beyond guidelines** | Black | -1.99 | 0.369 |
| **Within guidelines** | Asian | -2.03 | 0.338 |
| **Beyond guidelines** | Asian | 2.03 | 0.338 |
| **Within guidelines** | Other/Mixed | 1.51 | 1.000 |
| **Beyond guidelines** | Other/Mixed | -1.51 | 1.000 |
| **Measure** | **hisp** | **StdResidual** | **P-value** |
| **Within guidelines** | No | -0.725 | 1 |
| **Beyond guidelines** | No | 0.725 | 1 |
| **Within guidelines** | Yes | 0.725 | 1 |
| **Beyond guidelines** | Yes | -0.725 | 1 |
| **Measure** | **household.income** | **StdResidual** | **P-value** |
| **Within guidelines** | [<50K] | 2.121 | 0.203 |
| **Beyond guidelines** | [<50K] | -2.121 | 0.203 |
| **Within guidelines** | [>=50K & <100K] | -0.499 | 1.000 |
| **Beyond guidelines** | [>=50K & <100K] | 0.499 | 1.000 |
| **Within guidelines** | [>=100K] | -1.514 | 0.780 |
| **Beyond guidelines** | [>=100K] | 1.514 | 0.780 |
| **Measure** | **high.educ** | **StdResidual** | **P-value** |
| **Within guidelines** | < HS Diploma | 2.224 | 0.261 |
| **Beyond guidelines** | < HS Diploma | -2.224 | 0.261 |
| **Within guidelines** | HS Diploma/GED | 1.322 | 1.000 |
| **Beyond guidelines** | HS Diploma/GED | -1.322 | 1.000 |
| **Within guidelines** | Some College | 0.416 | 1.000 |
| **Beyond guidelines** | Some College | -0.416 | 1.000 |
| **Within guidelines** | Bachelor | -0.506 | 1.000 |
| **Beyond guidelines** | Bachelor | 0.506 | 1.000 |
| **Within guidelines** | Post Graduate Degree | -1.590 | 1.000 |
| **Beyond guidelines** | Post Graduate Degree | 1.590 | 1.000 |

*Supplementary Table 47. DV: Total pregnancy problems. Chi-squared test and pairwise post-hoc comparisons for associations with the sociodemographic factors.*

| **Demographic** | **X.squared** | **DF** | **p.value** |
| --- | --- | --- | --- |
| **sex_at_birth** | 5.13 | 2 | 0.0768 |
| **race.4level** | 68.4 | 6 | 8.55e-13 |
| **hisp** | 22.7 | 2 | 1.17e-05 |
| **household.income** | 115 | 4 | 7.64e-24 |
| **high.educ** | 155 | 8 | 1.46e-29 |
| **Measure** | **sex_at_birth** | **StdResidual** | **P-value** |
| **None** | F | -2.162 | 0.184 |
| **One problem** | F | 2.025 | 0.257 |
| **Two or more problems** | F | 0.519 | 1.000 |
| **None** | M | 2.162 | 0.184 |
| **One problem** | M | -2.025 | 0.257 |
| **Two or more problems** | M | -0.519 | 1.000 |
| **Measure** | **race.4level** | **StdResidual** | **P-value** |
| **None** | White | 4.515 | 7.60e-05 |
| **One problem** | White | -0.569 | 1.00e+00 |
| **Two or more problems** | White | -5.615 | 2.36e-07 |
| **None** | Black | -4.638 | 4.22e-05 |
| **One problem** | Black | 0.956 | 1.00e+00 |
| **Two or more problems** | Black | 5.308 | 1.33e-06 |
| **None** | Asian | 3.743 | 2.18e-03 |
| **One problem** | Asian | -1.455 | 1.00e+00 |
| **Two or more problems** | Asian | -3.438 | 7.04e-03 |
| **None** | Other/Mixed | -2.834 | 5.52e-02 |
| **One problem** | Other/Mixed | 0.406 | 1.00e+00 |
| **Two or more problems** | Other/Mixed | 3.463 | 6.41e-03 |
| **Measure** | **hisp** | **StdResidual** | **P-value** |
| **None** | No | 4.437 | 5.47e-05 |
| **One problem** | No | -4.400 | 6.50e-05 |
| **Two or more problems** | No | -0.761 | 1.00e+00 |
| **None** | Yes | -4.437 | 5.47e-05 |
| **One problem** | Yes | 4.400 | 6.50e-05 |
| **Two or more problems** | Yes | 0.761 | 1.00e+00 |
| **Measure** | **household.income** | **StdResidual** | **P-value** |
| **None** | [<50K] | -8.603 | 6.96e-17 |
| **One problem** | [<50K] | 2.816 | 4.37e-02 |
| **Two or more problems** | [<50K] | 8.555 | 1.07e-16 |
| **None** | [>=50K & <100K] | 0.157 | 1.00e+00 |
| **One problem** | [>=50K & <100K] | 0.140 | 1.00e+00 |
| **Two or more problems** | [>=50K & <100K] | -0.393 | 1.00e+00 |
| **None** | [>=100K] | 7.851 | 3.71e-14 |
| **One problem** | [>=100K] | -2.745 | 5.45e-02 |
| **Two or more problems** | [>=100K] | -7.590 | 2.88e-13 |
| **Measure** | **high.educ** | **StdResidual** | **P-value** |
| **None** | < HS Diploma | -2.738 | 9.28e-02 |
| **One problem** | < HS Diploma | 1.565 | 1.00e+00 |
| **Two or more problems** | < HS Diploma | 1.894 | 8.73e-01 |
| **None** | HS Diploma/GED | -3.018 | 3.82e-02 |
| **One problem** | HS Diploma/GED | 0.012 | 1.00e+00 |
| **Two or more problems** | HS Diploma/GED | 4.210 | 3.83e-04 |
| **None** | Some College | -8.204 | 3.47e-15 |
| **One problem** | Some College | 2.971 | 4.46e-02 |
| **Two or more problems** | Some College | 7.805 | 8.93e-14 |
| **None** | Bachelor | 1.671 | 1.00e+00 |
| **One problem** | Bachelor | -0.303 | 1.00e+00 |
| **Two or more problems** | Bachelor | -1.964 | 7.43e-01 |
| **None** | Post Graduate Degree | 8.827 | 1.61e-17 |
| **One problem** | Post Graduate Degree | -3.068 | 3.23e-02 |
| **Two or more problems** | Post Graduate Degree | -8.555 | 1.76e-16 |

*Supplementary Table 48. DV: Total birth problems. Chi-squared test and pairwise post-hoc comparisons for associations with the sociodemographic factors.*

| **Demographic** | **X.squared** | **DF** | **p.value** |
| --- | --- | --- | --- |
| **sex_at_birth** | 11.4 | 2 | 0.00337 |
| **race.4level** | 19.9 |  | 0.0055 |
| **hisp** | 12.5 | 2 | 0.00191 |
| **household.income** | 19.4 | 4 | 0.00065 |
| **high.educ** | 37.9 | 8 | 7.77e-06 |
| **Measure** | **sex_at_birth** | **StdResidual** | **P-value** |
| **None** | F | 3.28 | 0.00634 |
| **One problem** | F | -2.28 | 0.13500 |
| **Two or more problems** | F | -2.20 | 0.16600 |
| **None** | M | -3.28 | 0.00634 |
| **One problem** | M | 2.28 | 0.13500 |
| **Two or more problems** | M | 2.20 | 0.16600 |
| **Measure** | **race.4level** | **StdResidual** | **P-value** |
| **None** | White | -2.612 | 0.10800 |
| **One problem** | White | 0.753 | 1.00000 |
| **Two or more problems** | White | 3.485 | 0.00591 |
| **None** | Black | 1.994 | 0.55400 |
| **One problem** | Black | -1.650 | 1.00000 |
| **Two or more problems** | Black | -0.921 | 1.00000 |
| **None** | Asian | 0.601 | 1.00000 |
| **One problem** | Asian | 0.907 | 1.00000 |
| **Two or more problems** | Asian | -2.548 | 0.13000 |
| **None** | Other/Mixed | 1.165 | 1.00000 |
| **One problem** | Other/Mixed | 0.238 | 1.00000 |
| **Two or more problems** | Other/Mixed | -2.482 | 0.15700 |
| **Measure** | **hisp** | **StdResidual** | **P-value** |
| **None** | No | -3.43 | 0.00365 |
| **One problem** | No | 2.37 | 0.10600 |
| **Two or more problems** | No | 2.33 | 0.11900 |
| **None** | Yes | 3.43 | 0.00365 |
| **One problem** | Yes | -2.37 | 0.10600 |
| **Two or more problems** | Yes | -2.33 | 0.11900 |
| **Measure** | **household.income** | **StdResidual** | **P-value** |
| **None** | [<50K] | 4.080 | 0.000406 |
| **One problem** | [<50K] | -3.595 | 0.002920 |
| **Two or more problems** | [<50K] | -1.530 | 1.000000 |
| **None** | [>=50K & <100K] | -2.917 | 0.031800 |
| **One problem** | [>=50K & <100K] | 2.965 | 0.027200 |
| **Two or more problems** | [>=50K & <100K] | 0.455 | 1.000000 |
| **None** | [>=100K] | -1.123 | 1.000000 |
| **One problem** | [>=100K] | 0.628 | 1.000000 |
| **Two or more problems** | [>=100K] | 1.005 | 1.000000 |
| **Measure** | **high.educ** | **StdResidual** | **P-value** |
| **None** | < HS Diploma | 2.299 | 0.323000 |
| **One problem** | < HS Diploma | -1.768 | 1.000000 |
| **Two or more problems** | < HS Diploma | -1.278 | 1.000000 |
| **None** | HS Diploma/GED | 3.393 | 0.010400 |
| **One problem** | HS Diploma/GED | -3.413 | 0.009620 |
| **Two or more problems** | HS Diploma/GED | -0.587 | 1.000000 |
| **None** | Some College | -0.873 | 1.000000 |
| **One problem** | Some College | -0.221 | 1.000000 |
| **Two or more problems** | Some College | 1.928 | 0.807000 |
| **None** | Bachelor | -4.345 | 0.000209 |
| **One problem** | Bachelor | 3.743 | 0.002730 |
| **Two or more problems** | Bachelor | 1.768 | 1.000000 |
| **None** | Post Graduate Degree | 1.840 | 0.987000 |
| **One problem** | Post Graduate Degree | -0.496 | 1.000000 |
| **Two or more problems** | Post Graduate Degree | -2.510 | 0.181000 |

*Supplementary Table 49. DV: Prenatal alcohol exposure. Chi-squared test and pairwise post-hoc comparisons for associations with the sociodemographic factors.*

| **Demographic** | **X.squared** | **DF** | **p.value** |
| --- | --- | --- | --- |
| **sex_at_birth** | 1.89 | 2 | 0.388 |
| **race.4level** | 103 |  | 5e-04 |
| **hisp** | 35.8 | 2 | 1.66e-08 |
| **household.income** | 196 | 4 | 2.6e-41 |
| **high.educ** | 169 |  | 5e-04 |
| **Measure** | **sex_at_birth** | **StdResidual** | **P-value** |
| **Pre-No/Post-No** | F | -1.281 | 1 |
| **Pre-Yes/Post-No** | F | 1.010 | 1 |
| **Pre-Yes/Post-Yes** | F | 0.832 | 1 |
| **Pre-No/Post-No** | M | 1.281 | 1 |
| **Pre-Yes/Post-No** | M | -1.010 | 1 |
| **Pre-Yes/Post-Yes** | M | -0.832 | 1 |
| **Measure** | **race.4level** | **StdResidual** | **P-value** |
| **Pre-No/Post-No** | White | -9.05 | 1.65e-18 |
| **Pre-Yes/Post-No** | White | 7.92 | 2.74e-14 |
| **Pre-Yes/Post-Yes** | White | 3.81 | 1.66e-03 |
| **Pre-No/Post-No** | Black | 8.13 | 5.34e-15 |
| **Pre-Yes/Post-No** | Black | -7.57 | 4.60e-13 |
| **Pre-Yes/Post-Yes** | Black | -2.22 | 3.18e-01 |
| **Pre-No/Post-No** | Asian | 3.06 | 2.63e-02 |
| **Pre-Yes/Post-No** | Asian | -2.29 | 2.65e-01 |
| **Pre-Yes/Post-Yes** | Asian | -2.32 | 2.42e-01 |
| **Pre-No/Post-No** | Other/Mixed | 2.54 | 1.35e-01 |
| **Pre-Yes/Post-No** | Other/Mixed | -1.94 | 6.22e-01 |
| **Pre-Yes/Post-Yes** | Other/Mixed | -1.79 | 8.78e-01 |
| **Measure** | **hisp** | **StdResidual** | **P-value** |
| **Pre-No/Post-No** | No | -5.99 | 1.29e-08 |
| **Pre-Yes/Post-No** | No | 5.56 | 1.58e-07 |
| **Pre-Yes/Post-Yes** | No | 1.66 | 5.82e-01 |
| **Pre-No/Post-No** | Yes | 5.99 | 1.29e-08 |
| **Pre-Yes/Post-No** | Yes | -5.56 | 1.58e-07 |
| **Pre-Yes/Post-Yes** | Yes | -1.66 | 5.82e-01 |
| **Measure** | **household.income** | **StdResidual** | **P-value** |
| **Pre-No/Post-No** | [<50K] | 12.85 | 7.92e-37 |
| **Pre-Yes/Post-No** | [<50K] | -12.00 | 3.29e-32 |
| **Pre-Yes/Post-Yes** | [<50K] | -3.42 | 5.65e-03 |
| **Pre-No/Post-No** | [>=50K & <100K] | -0.59 | 1.00e+00 |
| **Pre-Yes/Post-No** | [>=50K & <100K] | 1.20 | 1.00e+00 |
| **Pre-Yes/Post-Yes** | [>=50K & <100K] | -1.56 | 1.00e+00 |
| **Pre-No/Post-No** | [>=100K] | -11.47 | 1.66e-29 |
| **Pre-Yes/Post-No** | [>=100K] | 10.12 | 4.23e-23 |
| **Pre-Yes/Post-Yes** | [>=100K] | 4.63 | 3.33e-05 |
| **Measure** | **high.educ** | **StdResidual** | **P-value** |
| **Pre-No/Post-No** | < HS Diploma | 8.000 | 1.87e-14 |
| **Pre-Yes/Post-No** | < HS Diploma | -7.607 | 4.20e-13 |
| **Pre-Yes/Post-Yes** | < HS Diploma | -1.766 | 1.00e+00 |
| **Pre-No/Post-No** | HS Diploma/GED | 7.704 | 1.98e-13 |
| **Pre-Yes/Post-No** | HS Diploma/GED | -6.786 | 1.73e-10 |
| **Pre-Yes/Post-Yes** | HS Diploma/GED | -3.128 | 2.64e-02 |
| **Pre-No/Post-No** | Some College | 2.836 | 6.86e-02 |
| **Pre-Yes/Post-No** | Some College | -2.472 | 2.01e-01 |
| **Pre-Yes/Post-Yes** | Some College | -1.219 | 1.00e+00 |
| **Pre-No/Post-No** | Bachelor | -3.182 | 2.19e-02 |
| **Pre-Yes/Post-No** | Bachelor | 3.171 | 2.28e-02 |
| **Pre-Yes/Post-Yes** | Bachelor | 0.319 | 1.00e+00 |
| **Pre-No/Post-No** | Post Graduate Degree | -7.552 | 6.41e-13 |
| **Pre-Yes/Post-No** | Post Graduate Degree | 6.522 | 1.04e-09 |
| **Pre-Yes/Post-Yes** | Post Graduate Degree | 3.411 | 9.71e-03 |

*Supplementary Table 50. DV: Prenatal tobacco exposure. Chi-squared test and pairwise post-hoc comparisons for associations with the sociodemographic factors.*

| **Demographic** | **X.squared** | **DF** | **p.value** |
| --- | --- | --- | --- |
| **sex_at_birth** | 0.439 | 2 | 0.803 |
| **race.4level** | 81.3 |  | 5e-04 |
| **hisp** | 16 | 2 | 0.000328 |
| **household.income** | 456 | 4 | 2.51e-97 |
| **high.educ** | 636 | 8 | 4.52e-132 |
| **Measure** | **sex_at_birth** | **StdResidual** | **P-value** |
| **Pre-No/Post-No** | F | 0.436 | 1 |
| **Pre-Yes/Post-No** | F | -0.655 | 1 |
| **Pre-Yes/Post-Yes** | F | 0.146 | 1 |
| **Pre-No/Post-No** | M | -0.436 | 1 |
| **Pre-Yes/Post-No** | M | 0.655 | 1 |
| **Pre-Yes/Post-Yes** | M | -0.146 | 1 |
| **Measure** | **race.4level** | **StdResidual** | **P-value** |
| **Pre-No/Post-No** | White | 7.24 | 5.24e-12 |
| **Pre-Yes/Post-No** | White | -5.86 | 5.50e-08 |
| **Pre-Yes/Post-Yes** | White | -3.86 | 1.36e-03 |
| **Pre-No/Post-No** | Black | -6.58 | 5.78e-10 |
| **Pre-Yes/Post-No** | Black | 4.21 | 3.13e-04 |
| **Pre-Yes/Post-Yes** | Black | 4.90 | 1.12e-05 |
| **Pre-No/Post-No** | Asian | 2.97 | 3.60e-02 |
| **Pre-Yes/Post-No** | Asian | -1.85 | 7.65e-01 |
| **Pre-Yes/Post-Yes** | Asian | -2.27 | 2.80e-01 |
| **Pre-No/Post-No** | Other/Mixed | -4.12 | 4.64e-04 |
| **Pre-Yes/Post-No** | Other/Mixed | 4.16 | 3.82e-04 |
| **Pre-Yes/Post-Yes** | Other/Mixed | 1.15 | 1.00e+00 |
| **Measure** | **hisp** | **StdResidual** | **P-value** |
| **Pre-No/Post-No** | No | -2.231 | 0.154000 |
| **Pre-Yes/Post-No** | No | -0.439 | 1.000000 |
| **Pre-Yes/Post-Yes** | No | 4.002 | 0.000377 |
| **Pre-No/Post-No** | Yes | 2.231 | 0.154000 |
| **Pre-Yes/Post-No** | Yes | 0.439 | 1.000000 |
| **Pre-Yes/Post-Yes** | Yes | -4.002 | 0.000377 |
| **Measure** | **household.income** | **StdResidual** | **P-value** |
| **Pre-No/Post-No** | [<50K] | -18.718 | 3.17e-77 |
| **Pre-Yes/Post-No** | [<50K] | 10.807 | 2.88e-26 |
| **Pre-Yes/Post-Yes** | [<50K] | 15.416 | 1.16e-52 |
| **Pre-No/Post-No** | [>=50K & <100K] | -0.227 | 1.00e+00 |
| **Pre-Yes/Post-No** | [>=50K & <100K] | 1.344 | 1.00e+00 |
| **Pre-Yes/Post-Yes** | [>=50K & <100K] | -1.333 | 1.00e+00 |
| **Pre-No/Post-No** | [>=100K] | 17.603 | 2.11e-68 |
| **Pre-Yes/Post-No** | [>=100K] | -11.271 | 1.64e-28 |
| **Pre-Yes/Post-Yes** | [>=100K] | -13.108 | 2.67e-38 |
| **Measure** | **high.educ** | **StdResidual** | **P-value** |
| **Pre-No/Post-No** | < HS Diploma | -4.66 | 4.82e-05 |
| **Pre-Yes/Post-No** | < HS Diploma | 3.19 | 2.10e-02 |
| **Pre-Yes/Post-Yes** | < HS Diploma | 3.20 | 2.05e-02 |
| **Pre-No/Post-No** | HS Diploma/GED | -10.78 | 6.22e-26 |
| **Pre-Yes/Post-No** | HS Diploma/GED | 3.81 | 2.05e-03 |
| **Pre-Yes/Post-Yes** | HS Diploma/GED | 11.90 | 1.73e-31 |
| **Pre-No/Post-No** | Some College | -17.45 | 4.83e-67 |
| **Pre-Yes/Post-No** | Some College | 13.36 | 1.59e-39 |
| **Pre-Yes/Post-Yes** | Some College | 10.26 | 1.56e-23 |
| **Pre-No/Post-No** | Bachelor | 7.65 | 2.92e-13 |
| **Pre-Yes/Post-No** | Bachelor | -5.00 | 8.82e-06 |
| **Pre-Yes/Post-Yes** | Bachelor | -5.58 | 3.57e-07 |
| **Pre-No/Post-No** | Post Graduate Degree | 17.14 | 1.03e-64 |
| **Pre-Yes/Post-No** | Post Graduate Degree | -11.15 | 1.10e-27 |
| **Pre-Yes/Post-Yes** | Post Graduate Degree | -12.55 | 5.70e-35 |

*Supplementary Table 51. DV: Prenatal marijuana exposure. Chi-squared test and pairwise post-hoc comparisons for associations with the sociodemographic factors.*

| **Demographic** | **X.squared** | **DF** | **p.value** |
| --- | --- | --- | --- |
| **sex_at_birth** | 2.18 | 2 | 0.336 |
| **race.4level** | 139 |  | 5e-04 |
| **hisp** | 4.55 | 2 | 0.103 |
| **household.income** | 226 | 4 | 7.55e-48 |
| **high.educ** | 225 |  | 5e-04 |
| **Measure** | **sex_at_birth** | **StdResidual** | **P-value** |
| **Pre-No/Post-No** | F | -0.383 | 1 |
| **Pre-Yes/Post-No** | F | -0.577 | 1 |
| **Pre-Yes/Post-Yes** | F | 1.376 | 1 |
| **Pre-No/Post-No** | M | 0.383 | 1 |
| **Pre-Yes/Post-No** | M | 0.577 | 1 |
| **Pre-Yes/Post-Yes** | M | -1.376 | 1 |
| **Measure** | **race.4level** | **StdResidual** | **P-value** |
| **Pre-No/Post-No** | White | 7.77 | 9.75e-14 |
| **Pre-Yes/Post-No** | White | -6.73 | 2.09e-10 |
| **Pre-Yes/Post-Yes** | White | -3.81 | 1.70e-03 |
| **Pre-No/Post-No** | Black | -11.33 | 1.17e-28 |
| **Pre-Yes/Post-No** | Black | 8.73 | 2.99e-17 |
| **Pre-Yes/Post-Yes** | Black | 6.96 | 4.09e-11 |
| **Pre-No/Post-No** | Asian | 2.58 | 1.20e-01 |
| **Pre-Yes/Post-No** | Asian | -1.99 | 5.57e-01 |
| **Pre-Yes/Post-Yes** | Asian | -1.58 | 1.00e+00 |
| **Pre-No/Post-No** | Other/Mixed | -0.17 | 1.00e+00 |
| **Pre-Yes/Post-No** | Other/Mixed | 1.06 | 1.00e+00 |
| **Pre-Yes/Post-Yes** | Other/Mixed | -1.11 | 1.00e+00 |
| **Measure** | **hisp** | **StdResidual** | **P-value** |
| **Pre-No/Post-No** | No | -0.294 | 1.000 |
| **Pre-Yes/Post-No** | No | -1.070 | 1.000 |
| **Pre-Yes/Post-Yes** | No | 1.877 | 0.363 |
| **Pre-No/Post-No** | Yes | 0.294 | 1.000 |
| **Pre-Yes/Post-No** | Yes | 1.070 | 1.000 |
| **Pre-Yes/Post-Yes** | Yes | -1.877 | 0.363 |
| **Measure** | **household.income** | **StdResidual** | **P-value** |
| **Pre-No/Post-No** | [<50K] | -14.078 | 4.66e-44 |
| **Pre-Yes/Post-No** | [<50K] | 10.601 | 2.66e-25 |
| **Pre-Yes/Post-Yes** | [<50K] | 8.982 | 2.39e-18 |
| **Pre-No/Post-No** | [>=50K & <100K] | 1.445 | 1.00e+00 |
| **Pre-Yes/Post-No** | [>=50K & <100K] | -2.207 | 2.46e-01 |
| **Pre-Yes/Post-Yes** | [>=50K & <100K] | 0.541 | 1.00e+00 |
| **Pre-No/Post-No** | [>=100K] | 11.742 | 6.95e-31 |
| **Pre-Yes/Post-No** | [>=100K] | -7.819 | 4.78e-14 |
| **Pre-Yes/Post-Yes** | [>=100K] | -8.828 | 9.56e-18 |
| **Measure** | **high.educ** | **StdResidual** | **P-value** |
| **Pre-No/Post-No** | < HS Diploma | 0.1547 | 1.00e+00 |
| **Pre-Yes/Post-No** | < HS Diploma | -0.1386 | 1.00e+00 |
| **Pre-Yes/Post-Yes** | < HS Diploma | -0.0698 | 1.00e+00 |
| **Pre-No/Post-No** | HS Diploma/GED | -7.4707 | 1.20e-12 |
| **Pre-Yes/Post-No** | HS Diploma/GED | 5.6596 | 2.28e-07 |
| **Pre-Yes/Post-Yes** | HS Diploma/GED | 4.7217 | 3.51e-05 |
| **Pre-No/Post-No** | Some College | -10.9995 | 5.76e-27 |
| **Pre-Yes/Post-No** | Some College | 7.7858 | 1.04e-13 |
| **Pre-Yes/Post-Yes** | Some College | 7.6672 | 2.64e-13 |
| **Pre-No/Post-No** | Bachelor | 4.6976 | 3.95e-05 |
| **Pre-Yes/Post-No** | Bachelor | -3.7268 | 2.91e-03 |
| **Pre-Yes/Post-Yes** | Bachelor | -2.7494 | 8.96e-02 |
| **Pre-No/Post-No** | Post Graduate Degree | 10.0472 | 1.42e-22 |
| **Pre-Yes/Post-No** | Post Graduate Degree | -6.9507 | 5.45e-11 |
| **Pre-Yes/Post-Yes** | Post Graduate Degree | -7.2138 | 8.16e-12 |

*Supplementary Table 52. DV: Prenatal other substance exposure. Chi-squared test and pairwise post-hoc comparisons for associations with the sociodemographic factors.*

| **Demographic** | **X.squared** | **DF** | **p.value** |
| --- | --- | --- | --- |
| **sex_at_birth** | 1.24 | 2 | 0.539 |
| **race.4level** | 35.5 |  | 5e-04 |
| **hisp** | 5.33 | 2 | 0.0696 |
| **household.income** | 31.9 | 4 | 1.97e-06 |
| **high.educ** | 51.3 |  | 5e-04 |
| **Measure** | **sex_at_birth** | **StdResidual** | **P-value** |
| **Pre-No/Post-No** | F | 0.0164 | 1 |
| **Pre-Yes/Post-No** | F | 0.6619 | 1 |
| **Pre-Yes/Post-Yes** | F | -0.9003 | 1 |
| **Pre-No/Post-No** | M | -0.0164 | 1 |
| **Pre-Yes/Post-No** | M | -0.6619 | 1 |
| **Pre-Yes/Post-Yes** | M | 0.9003 | 1 |
| **Measure** | **race.4level** | **StdResidual** | **P-value** |
| **Pre-No/Post-No** | White | 2.9051 | 0.044100 |
| **Pre-Yes/Post-No** | White | -1.8546 | 0.764000 |
| **Pre-Yes/Post-Yes** | White | -2.3402 | 0.231000 |
| **Pre-No/Post-No** | Black | 0.0316 | 1.000000 |
| **Pre-Yes/Post-No** | Black | -2.3099 | 0.251000 |
| **Pre-Yes/Post-Yes** | Black | 2.9954 | 0.032900 |
| **Pre-No/Post-No** | Asian | 1.1192 | 1.000000 |
| **Pre-Yes/Post-No** | Asian | -0.4395 | 1.000000 |
| **Pre-Yes/Post-Yes** | Asian | -1.2644 | 1.000000 |
| **Pre-No/Post-No** | Other/Mixed | -4.1373 | 0.000422 |
| **Pre-Yes/Post-No** | Other/Mixed | 4.6826 | 0.000034 |
| **Pre-Yes/Post-Yes** | Other/Mixed | 0.6397 | 1.000000 |
| **Measure** | **hisp** | **StdResidual** | **P-value** |
| **Pre-No/Post-No** | No | 1.8945 | 0.349 |
| **Pre-Yes/Post-No** | No | -2.3066 | 0.126 |
| **Pre-Yes/Post-Yes** | No | -0.0787 | 1.000 |
| **Pre-No/Post-No** | Yes | -1.8945 | 0.349 |
| **Pre-Yes/Post-No** | Yes | 2.3066 | 0.126 |
| **Pre-Yes/Post-Yes** | Yes | 0.0787 | 1.000 |
| **Measure** | **household.income** | **StdResidual** | **P-value** |
| **Pre-No/Post-No** | [<50K] | -5.0232 | 4.57e-06 |
| **Pre-Yes/Post-No** | [<50K] | 4.6529 | 2.95e-05 |
| **Pre-Yes/Post-Yes** | [<50K] | 2.1386 | 2.92e-01 |
| **Pre-No/Post-No** | [>=50K & <100K] | -0.0105 | 1.00e+00 |
| **Pre-Yes/Post-No** | [>=50K & <100K] | 0.0467 | 1.00e+00 |
| **Pre-Yes/Post-Yes** | [>=50K & <100K] | -0.0443 | 1.00e+00 |
| **Pre-No/Post-No** | [>=100K] | 4.6784 | 2.60e-05 |
| **Pre-Yes/Post-No** | [>=100K] | -4.3673 | 1.13e-04 |
| **Pre-Yes/Post-Yes** | [>=100K] | -1.9472 | 4.64e-01 |
| **Measure** | **high.educ** | **StdResidual** | **P-value** |
| **Pre-No/Post-No** | < HS Diploma | 0.343 | 1.00e+00 |
| **Pre-Yes/Post-No** | < HS Diploma | -0.675 | 1.00e+00 |
| **Pre-Yes/Post-Yes** | < HS Diploma | 0.324 | 1.00e+00 |
| **Pre-No/Post-No** | HS Diploma/GED | -1.813 | 1.00e+00 |
| **Pre-Yes/Post-No** | HS Diploma/GED | 1.779 | 1.00e+00 |
| **Pre-Yes/Post-Yes** | HS Diploma/GED | 0.641 | 1.00e+00 |
| **Pre-No/Post-No** | Some College | -6.230 | 7.02e-09 |
| **Pre-Yes/Post-No** | Some College | 5.703 | 1.77e-07 |
| **Pre-Yes/Post-Yes** | Some College | 2.741 | 9.19e-02 |
| **Pre-No/Post-No** | Bachelor | 2.848 | 6.60e-02 |
| **Pre-Yes/Post-No** | Bachelor | -2.145 | 4.79e-01 |
| **Pre-Yes/Post-Yes** | Bachelor | -1.863 | 9.37e-01 |
| **Pre-No/Post-No** | Post Graduate Degree | 4.004 | 9.36e-04 |
| **Pre-Yes/Post-No** | Post Graduate Degree | -4.011 | 9.07e-04 |
| **Pre-Yes/Post-Yes** | Post Graduate Degree | -1.305 | 1.00e+00 |

*Supplementary Table 53. DV: Average hours of sleep per night. Chi-squared test and pairwise post-hoc comparisons for associations with the sociodemographic factors.*

| **Demographic** | **X.squared** | **DF** | **p.value** |
| --- | --- | --- | --- |
| **age_quartiles** | 72.8 | 9 | 4.36e-12 |
| **sex_at_birth** | 6.35 | 3 | 0.0956 |
| **race.4level** | 811 |  | 5e-04 |
| **hisp** | 130 | 3 | 4.84e-28 |
| **household.income** | 834 | 6 | 6.98e-177 |
| **high.educ** | 891 | 12 | 5.37e-183 |
| **pubertal_dev_collapsed** | 246 |  | 5e-04 |
| **Measure** | **age_quartiles** | **StdResidual** | **P-value** |
| **Less than 7 hours** | age<=25% | -1.052 | 1.00e+00 |
| **7-8 hours** | age<=25% | -0.911 | 1.00e+00 |
| **8-9 hours** | age<=25% | -4.601 | 6.73e-05 |
| **9-11 hours** | age<=25% | 5.371 | 1.25e-06 |
| **Less than 7 hours** | age>25&<=50% | -2.347 | 3.03e-01 |
| **7-8 hours** | age>25&<=50% | -1.415 | 1.00e+00 |
| **8-9 hours** | age>25&<=50% | -1.264 | 1.00e+00 |
| **9-11 hours** | age>25&<=50% | 2.938 | 5.29e-02 |
| **Less than 7 hours** | age>50&<=75% | 0.952 | 1.00e+00 |
| **7-8 hours** | age>50&<=75% | -0.761 | 1.00e+00 |
| **8-9 hours** | age>50&<=75% | 2.203 | 4.42e-01 |
| **9-11 hours** | age>50&<=75% | -1.969 | 7.83e-01 |
| **Less than 7 hours** | age>75% | 2.595 | 1.51e-01 |
| **7-8 hours** | age>75% | 3.170 | 2.44e-02 |
| **8-9 hours** | age>75% | 3.934 | 1.34e-03 |
| **9-11 hours** | age>75% | -6.705 | 3.22e-10 |
| **Measure** | **sex_at_birth** | **StdResidual** | **P-value** |
| **Less than 7 hours** | F | -1.585 | 0.904 |
| **7-8 hours** | F | -1.755 | 0.634 |
| **8-9 hours** | F | 1.319 | 1.000 |
| **9-11 hours** | F | 0.402 | 1.000 |
| **Less than 7 hours** | M | 1.585 | 0.904 |
| **7-8 hours** | M | 1.755 | 0.634 |
| **8-9 hours** | M | -1.319 | 1.000 |
| **9-11 hours** | M | -0.402 | 1.000 |
| **Measure** | **race.4level** | **StdResidual** | **P-value** |
| **Less than 7 hours** | White | -12.987 | 2.33e-37 |
| **7-8 hours** | White | -13.056 | 9.43e-38 |
| **8-9 hours** | White | -5.992 | 3.33e-08 |
| **9-11 hours** | White | 18.606 | 4.56e-76 |
| **Less than 7 hours** | Black | 16.587 | 1.38e-60 |
| **7-8 hours** | Black | 16.088 | 4.97e-57 |
| **8-9 hours** | Black | 3.879 | 1.68e-03 |
| **9-11 hours** | Black | -19.764 | 9.66e-86 |
| **Less than 7 hours** | Asian | -1.791 | 1.00e+00 |
| **7-8 hours** | Asian | -2.207 | 4.37e-01 |
| **8-9 hours** | Asian | 0.773 | 1.00e+00 |
| **9-11 hours** | Asian | 1.286 | 1.00e+00 |
| **Less than 7 hours** | Other/Mixed | 1.651 | 1.00e+00 |
| **7-8 hours** | Other/Mixed | 2.367 | 2.87e-01 |
| **8-9 hours** | Other/Mixed | 3.614 | 4.83e-03 |
| **9-11 hours** | Other/Mixed | -5.556 | 4.41e-07 |
| **Measure** | **hisp** | **StdResidual** | **P-value** |
| **Less than 7 hours** | No | 0.147 | 1.00e+00 |
| **7-8 hours** | No | -5.305 | 9.03e-07 |
| **8-9 hours** | No | -8.141 | 3.14e-15 |
| **9-11 hours** | No | 11.140 | 6.41e-28 |
| **Less than 7 hours** | Yes | -0.147 | 1.00e+00 |
| **7-8 hours** | Yes | 5.305 | 9.03e-07 |
| **8-9 hours** | Yes | 8.141 | 3.14e-15 |
| **9-11 hours** | Yes | -11.140 | 6.41e-28 |
| **Measure** | **household.income** | **StdResidual** | **P-value** |
| **Less than 7 hours** | [<50K] | 13.269 | 4.21e-39 |
| **7-8 hours** | [<50K] | 16.410 | 1.95e-59 |
| **8-9 hours** | [<50K] | 8.117 | 5.75e-15 |
| **9-11 hours** | [<50K] | -22.876 | 0.00e+00 |
| **Less than 7 hours** | [>=50K & <100K] | -2.233 | 3.06e-01 |
| **7-8 hours** | [>=50K & <100K] | -1.364 | 1.00e+00 |
| **8-9 hours** | [>=50K & <100K] | 2.041 | 4.95e-01 |
| **9-11 hours** | [>=50K & <100K] | -0.312 | 1.00e+00 |
| **Less than 7 hours** | [>=100K] | -10.120 | 5.41e-23 |
| **7-8 hours** | [>=100K] | -13.794 | 3.32e-42 |
| **8-9 hours** | [>=100K] | -9.305 | 1.60e-19 |
| **9-11 hours** | [>=100K] | 21.253 | 3.70e-99 |
| **Measure** | **high.educ** | **StdResidual** | **P-value** |
| **Less than 7 hours** | < HS Diploma | 4.09 | 8.62e-04 |
| **7-8 hours** | < HS Diploma | 6.35 | 4.21e-09 |
| **8-9 hours** | < HS Diploma | 3.63 | 5.59e-03 |
| **9-11 hours** | < HS Diploma | -8.96 | 6.43e-18 |
| **Less than 7 hours** | HS Diploma/GED | 6.66 | 5.41e-10 |
| **7-8 hours** | HS Diploma/GED | 8.53 | 3.01e-16 |
| **8-9 hours** | HS Diploma/GED | 5.54 | 6.12e-07 |
| **9-11 hours** | HS Diploma/GED | -13.07 | 9.22e-38 |
| **Less than 7 hours** | Some College | 8.02 | 2.11e-14 |
| **7-8 hours** | Some College | 10.67 | 2.80e-25 |
| **8-9 hours** | Some College | 7.40 | 2.83e-12 |
| **9-11 hours** | Some College | -16.70 | 2.75e-61 |
| **Less than 7 hours** | Bachelor | -3.74 | 3.69e-03 |
| **7-8 hours** | Bachelor | -5.52 | 6.63e-07 |
| **8-9 hours** | Bachelor | -1.90 | 1.00e+00 |
| **9-11 hours** | Bachelor | 6.64 | 6.09e-10 |
| **Less than 7 hours** | Post Graduate Degree | -9.20 | 6.97e-19 |
| **7-8 hours** | Post Graduate Degree | -11.90 | 2.35e-31 |
| **8-9 hours** | Post Graduate Degree | -9.50 | 4.11e-20 |
| **9-11 hours** | Post Graduate Degree | 19.92 | 5.59e-87 |
| **Measure** | **pubertal_dev_collapsed** | **StdResidual** | **P-value** |
| **Less than 7 hours** | Pre | -5.84562 | 8.07e-08 |
| **7-8 hours** | Pre | -7.23132 | 7.65e-12 |
| **8-9 hours** | Pre | -5.51184 | 5.68e-07 |
| **9-11 hours** | Pre | 11.94068 | 1.16e-31 |
| **Less than 7 hours** | Early | 0.05435 | 1.00e+00 |
| **7-8 hours** | Early | 0.75635 | 1.00e+00 |
| **8-9 hours** | Early | 0.50933 | 1.00e+00 |
| **9-11 hours** | Early | -0.98857 | 1.00e+00 |
| **Less than 7 hours** | Mid | 6.86186 | 1.09e-10 |
| **7-8 hours** | Mid | 7.08878 | 2.16e-11 |
| **8-9 hours** | Mid | 4.92313 | 1.36e-05 |
| **9-11 hours** | Mid | -11.64143 | 4.06e-30 |
| **Less than 7 hours** | Late/Post | -0.00904 | 1.00e+00 |
| **7-8 hours** | Late/Post | 2.46285 | 2.21e-01 |
| **8-9 hours** | Late/Post | 3.80343 | 2.28e-03 |
| **9-11 hours** | Late/Post | -5.21568 | 2.93e-06 |

*Supplementary Table 54. DV: Total sleep disturbance. Chi-squared test and pairwise post-hoc comparisons for associations with the sociodemographic factors.*

| **Demographic** | **X.squared** | **DF** | **p.value** |
| --- | --- | --- | --- |
| **age_quartiles** | 3.92 | 3 | 0.27 |
| **sex_at_birth** | 3.22 | 1 | 0.0729 |
| **race.4level** | 21.1 | 3 | 0.000102 |
| **hisp** | 0.22 | 1 | 0.639 |
| **household.income** | 73.8 | 2 | 9.29e-17 |
| **high.educ** | 52.2 | 4 | 1.27e-10 |
| **pubertal_dev_collapsed** | 33.5 | 3 | 2.49e-07 |
| **Measure** | **age_quartiles** | **StdResidual** | **P-value** |
| **Low Sleep Disturbance** | age<=25% | -1.816 | 0.554 |
| **High Sleep Disturbance** | age<=25% | 1.816 | 0.554 |
| **Low Sleep Disturbance** | age>25&<=50% | 1.325 | 1.000 |
| **High Sleep Disturbance** | age>25&<=50% | -1.325 | 1.000 |
| **Low Sleep Disturbance** | age>50&<=75% | -0.017 | 1.000 |
| **High Sleep Disturbance** | age>50&<=75% | 0.017 | 1.000 |
| **Low Sleep Disturbance** | age>75% | 0.533 | 1.000 |
| **High Sleep Disturbance** | age>75% | -0.533 | 1.000 |
| **Measure** | **sex_at_birth** | **StdResidual** | **P-value** |
| **Low Sleep Disturbance** | F | 1.82 | 0.277 |
| **High Sleep Disturbance** | F | -1.82 | 0.277 |
| **Low Sleep Disturbance** | M | -1.82 | 0.277 |
| **High Sleep Disturbance** | M | 1.82 | 0.277 |
| **Measure** | **race.4level** | **StdResidual** | **P-value** |
| **Low Sleep Disturbance** | White | 4.05 | 0.000417 |
| **High Sleep Disturbance** | White | -4.05 | 0.000417 |
| **Low Sleep Disturbance** | Black | -2.42 | 0.125000 |
| **High Sleep Disturbance** | Black | 2.42 | 0.125000 |
| **Low Sleep Disturbance** | Asian | 1.20 | 1.000000 |
| **High Sleep Disturbance** | Asian | -1.20 | 1.000000 |
| **Low Sleep Disturbance** | Other/Mixed | -3.32 | 0.007170 |
| **High Sleep Disturbance** | Other/Mixed | 3.32 | 0.007170 |
| **Measure** | **hisp** | **StdResidual** | **P-value** |
| **Low Sleep Disturbance** | No | -0.498 | 1 |
| **High Sleep Disturbance** | No | 0.498 | 1 |
| **Low Sleep Disturbance** | Yes | 0.498 | 1 |
| **High Sleep Disturbance** | Yes | -0.498 | 1 |
| **Measure** | **household.income** | **StdResidual** | **P-value** |
| **Low Sleep Disturbance** | [<50K] | -6.86 | 4.21e-11 |
| **High Sleep Disturbance** | [<50K] | 6.86 | 4.21e-11 |
| **Low Sleep Disturbance** | [>=50K & <100K] | -1.97 | 2.92e-01 |
| **High Sleep Disturbance** | [>=50K & <100K] | 1.97 | 2.92e-01 |
| **Low Sleep Disturbance** | [>=100K] | 8.09 | 3.65e-15 |
| **High Sleep Disturbance** | [>=100K] | -8.09 | 3.65e-15 |
| **Measure** | **high.educ** | **StdResidual** | **P-value** |
| **Low Sleep Disturbance** | < HS Diploma | -1.18 | 1.00e+00 |
| **High Sleep Disturbance** | < HS Diploma | 1.18 | 1.00e+00 |
| **Low Sleep Disturbance** | HS Diploma/GED | -2.34 | 1.94e-01 |
| **High Sleep Disturbance** | HS Diploma/GED | 2.34 | 1.94e-01 |
| **Low Sleep Disturbance** | Some College | -5.60 | 2.20e-07 |
| **High Sleep Disturbance** | Some College | 5.60 | 2.20e-07 |
| **Low Sleep Disturbance** | Bachelor | 1.11 | 1.00e+00 |
| **High Sleep Disturbance** | Bachelor | -1.11 | 1.00e+00 |
| **Low Sleep Disturbance** | Post Graduate Degree | 5.83 | 5.49e-08 |
| **High Sleep Disturbance** | Post Graduate Degree | -5.83 | 5.49e-08 |
| **Measure** | **pubertal_dev_collapsed** | **StdResidual** | **P-value** |
| **Low Sleep Disturbance** | Pre | 5.74 | 7.75e-08 |
| **High Sleep Disturbance** | Pre | -5.74 | 7.75e-08 |
| **Low Sleep Disturbance** | Early | -2.83 | 3.69e-02 |
| **High Sleep Disturbance** | Early | 2.83 | 3.69e-02 |
| **Low Sleep Disturbance** | Mid | -3.54 | 3.18e-03 |
| **High Sleep Disturbance** | Mid | 3.54 | 3.18e-03 |
| **Low Sleep Disturbance** | Late/Post | -1.34 | 1.00e+00 |
| **High Sleep Disturbance** | Late/Post | 1.34 | 1.00e+00 |

*Supplementary Table 55. DV: Physical activity (vigorous). Chi-squared test and pairwise post-hoc comparisons for associations with the sociodemographic factors.*

| **Demographic** | **X.squared** | **DF** | **p.value** |
| --- | --- | --- | --- |
| **age_quartiles** | 1.77 | 3 | 0.622 |
| **sex_at_birth** | 14.3 | 1 | 0.000159 |
| **race.4level** | 4.08 | 3 | 0.253 |
| **hisp** | 13.6 | 1 | 0.000225 |
| **household.income** | 7.08 | 2 | 0.029 |
| **high.educ** | 15.2 | 4 | 0.00437 |
| **pubertal_dev_collapsed** | 17.7 | 3 | 0.000496 |
| **Measure** | **age_quartiles** | **StdResidual** | **P-value** |
| **Below guidelines** | age<=25% | -0.013 | 1 |
| **Within guidelines** | age<=25% | 0.013 | 1 |
| **Below guidelines** | age>25&<=50% | 0.916 | 1 |
| **Within guidelines** | age>25&<=50% | -0.916 | 1 |
| **Below guidelines** | age>50&<=75% | 0.266 | 1 |
| **Within guidelines** | age>50&<=75% | -0.266 | 1 |
| **Below guidelines** | age>75% | -1.200 | 1 |
| **Within guidelines** | age>75% | 1.200 | 1 |
| **Measure** | **sex_at_birth** | **StdResidual** | **P-value** |
| **Below guidelines** | F | 3.8 | 0.000567 |
| **Within guidelines** | F | -3.8 | 0.000567 |
| **Below guidelines** | M | -3.8 | 0.000567 |
| **Within guidelines** | M | 3.8 | 0.000567 |
| **Measure** | **race.4level** | **StdResidual** | **P-value** |
| **Below guidelines** | White | -1.766 | 0.619 |
| **Within guidelines** | White | 1.766 | 0.619 |
| **Below guidelines** | Black | 1.326 | 1.000 |
| **Within guidelines** | Black | -1.326 | 1.000 |
| **Below guidelines** | Asian | 1.148 | 1.000 |
| **Within guidelines** | Asian | -1.148 | 1.000 |
| **Below guidelines** | Other/Mixed | 0.528 | 1.000 |
| **Within guidelines** | Other/Mixed | -0.528 | 1.000 |
| **Measure** | **hisp** | **StdResidual** | **P-value** |
| **Below guidelines** | No | -3.72 | 0.000782 |
| **Within guidelines** | No | 3.72 | 0.000782 |
| **Below guidelines** | Yes | 3.72 | 0.000782 |
| **Within guidelines** | Yes | -3.72 | 0.000782 |
| **Measure** | **household.income** | **StdResidual** | **P-value** |
| **Below guidelines** | [<50K] | 2.66 | 0.0467 |
| **Within guidelines** | [<50K] | -2.66 | 0.0467 |
| **Below guidelines** | [>=50K & <100K] | -1.05 | 1.0000 |
| **Within guidelines** | [>=50K & <100K] | 1.05 | 1.0000 |
| **Below guidelines** | [>=100K] | -1.48 | 0.8310 |
| **Within guidelines** | [>=100K] | 1.48 | 0.8310 |
| **Measure** | **high.educ** | **StdResidual** | **P-value** |
| **Below guidelines** | < HS Diploma | 2.098 | 0.359 |
| **Within guidelines** | < HS Diploma | -2.098 | 0.359 |
| **Below guidelines** | HS Diploma/GED | 2.895 | 0.038 |
| **Within guidelines** | HS Diploma/GED | -2.895 | 0.038 |
| **Below guidelines** | Some College | 0.253 | 1.000 |
| **Within guidelines** | Some College | -0.253 | 1.000 |
| **Below guidelines** | Bachelor | -0.579 | 1.000 |
| **Within guidelines** | Bachelor | 0.579 | 1.000 |
| **Below guidelines** | Post Graduate Degree | -2.149 | 0.316 |
| **Within guidelines** | Post Graduate Degree | 2.149 | 0.316 |
| **Measure** | **pubertal_dev_collapsed** | **StdResidual** | **P-value** |
| **Below guidelines** | Pre | -3.824 | 0.00105 |
| **Within guidelines** | Pre | 3.824 | 0.00105 |
| **Below guidelines** | Early | 0.908 | 1.00000 |
| **Within guidelines** | Early | -0.908 | 1.00000 |
| **Below guidelines** | Mid | 3.175 | 0.01200 |
| **Within guidelines** | Mid | -3.175 | 0.01200 |
| **Below guidelines** | Late/Post | 1.516 | 1.00000 |
| **Within guidelines** | Late/Post | -1.516 | 1.00000 |

*Supplementary Table 56. DV: Physical activity (strengthening). Chi-squared test and pairwise post-hoc comparisons for associations with the sociodemographic factors.*

| **Demographic** | **X.squared** | **DF** | **p.value** |
| --- | --- | --- | --- |
| **age_quartiles** | 9.79 | 3 | 0.0205 |
| **sex_at_birth** | 19.7 | 1 | 9.06e-06 |
| **race.4level** | 75.2 | 3 | 3.27e-16 |
| **hisp** | 0.629 | 1 | 0.428 |
| **household.income** | 31.3 | 2 | 1.61e-07 |
| **high.educ** | 36 | 4 | 2.96e-07 |
| **pubertal_dev_collapsed** | 1.8 | 3 | 0.614 |
| **Measure** | **age_quartiles** | **StdResidual** | **P-value** |
| **Below guidelines** | age<=25% | 2.06 | 0.318 |
| **Within guidelines** | age<=25% | -2.06 | 0.318 |
| **Below guidelines** | age>25&<=50% | 1.44 | 1.000 |
| **Within guidelines** | age>25&<=50% | -1.44 | 1.000 |
| **Below guidelines** | age>50&<=75% | -1.69 | 0.731 |
| **Within guidelines** | age>50&<=75% | 1.69 | 0.731 |
| **Below guidelines** | age>75% | -1.97 | 0.387 |
| **Within guidelines** | age>75% | 1.97 | 0.387 |
| **Measure** | **sex_at_birth** | **StdResidual** | **P-value** |
| **Below guidelines** | F | 4.46 | 3.25e-05 |
| **Within guidelines** | F | -4.46 | 3.25e-05 |
| **Below guidelines** | M | -4.46 | 3.25e-05 |
| **Within guidelines** | M | 4.46 | 3.25e-05 |
| **Measure** | **race.4level** | **StdResidual** | **P-value** |
| **Below guidelines** | White | 6.380 | 1.42e-09 |
| **Within guidelines** | White | -6.380 | 1.42e-09 |
| **Below guidelines** | Black | -8.410 | 3.28e-16 |
| **Within guidelines** | Black | 8.410 | 3.28e-16 |
| **Below guidelines** | Asian | 0.774 | 1.00e+00 |
| **Within guidelines** | Asian | -0.774 | 1.00e+00 |
| **Below guidelines** | Other/Mixed | -0.527 | 1.00e+00 |
| **Within guidelines** | Other/Mixed | 0.527 | 1.00e+00 |
| **Measure** | **hisp** | **StdResidual** | **P-value** |
| **Below guidelines** | No | 0.823 | 1 |
| **Within guidelines** | No | -0.823 | 1 |
| **Below guidelines** | Yes | -0.823 | 1 |
| **Within guidelines** | Yes | 0.823 | 1 |
| **Measure** | **household.income** | **StdResidual** | **P-value** |
| **Below guidelines** | [<50K] | -5.407 | 3.84e-07 |
| **Within guidelines** | [<50K] | 5.407 | 3.84e-07 |
| **Below guidelines** | [>=50K & <100K] | 0.875 | 1.00e+00 |
| **Within guidelines** | [>=50K & <100K] | -0.875 | 1.00e+00 |
| **Below guidelines** | [>=100K] | 4.156 | 1.94e-04 |
| **Within guidelines** | [>=100K] | -4.156 | 1.94e-04 |
| **Measure** | **high.educ** | **StdResidual** | **P-value** |
| **Below guidelines** | < HS Diploma | -1.80 | 0.7230 |
| **Within guidelines** | < HS Diploma | 1.80 | 0.7230 |
| **Below guidelines** | HS Diploma/GED | -3.85 | 0.0012 |
| **Within guidelines** | HS Diploma/GED | 3.85 | 0.0012 |
| **Below guidelines** | Some College | -3.08 | 0.0207 |
| **Within guidelines** | Some College | 3.08 | 0.0207 |
| **Below guidelines** | Bachelor | 3.10 | 0.0190 |
| **Within guidelines** | Bachelor | -3.10 | 0.0190 |
| **Below guidelines** | Post Graduate Degree | 2.83 | 0.0467 |
| **Within guidelines** | Post Graduate Degree | -2.83 | 0.0467 |
| **Measure** | **pubertal_dev_collapsed** | **StdResidual** | **P-value** |
| **Below guidelines** | Pre | 1.2246 | 1 |
| **Within guidelines** | Pre | -1.2246 | 1 |
| **Below guidelines** | Early | -1.1543 | 1 |
| **Within guidelines** | Early | 1.1543 | 1 |
| **Below guidelines** | Mid | -0.2756 | 1 |
| **Within guidelines** | Mid | 0.2756 | 1 |
| **Below guidelines** | Late/Post | -0.0249 | 1 |
| **Within guidelines** | Late/Post | 0.0249 | 1 |

*Supplementary Table 57. DV: Sports/activities involvement. Chi-squared test and pairwise post-hoc comparisons for associations with the sociodemographic factors.*

| **Demographic** | **X.squared** | **DF** | **p.value** |
| --- | --- | --- | --- |
| **age_quartiles** | 12.4 | 3 | 0.00613 |
| **sex_at_birth** | 14 | 1 | 0.000179 |
| **race.4level** | 281 | 3 | 1.06e-60 |
| **hisp** | 81.8 | 1 | 1.49e-19 |
| **household.income** | 850 | 2 | 2.11e-185 |
| **high.educ** | 870 | 4 | 4.37e-187 |
| **pubertal_dev_collapsed** | 92.8 | 3 | 5.37e-20 |
| **Measure** | **age_quartiles** | **StdResidual** | **P-value** |
| **No Participation** | age<=25% | 3.206 | 0.0108 |
| **Some Participation** | age<=25% | -3.206 | 0.0108 |
| **No Participation** | age>25&<=50% | -0.308 | 1.0000 |
| **Some Participation** | age>25&<=50% | 0.308 | 1.0000 |
| **No Participation** | age>50&<=75% | -0.585 | 1.0000 |
| **Some Participation** | age>50&<=75% | 0.585 | 1.0000 |
| **No Participation** | age>75% | -2.443 | 0.1170 |
| **Some Participation** | age>75% | 2.443 | 0.1170 |
| **Measure** | **sex_at_birth** | **StdResidual** | **P-value** |
| **No Participation** | F | 3.77 | 0.000645 |
| **Some Participation** | F | -3.77 | 0.000645 |
| **No Participation** | M | -3.77 | 0.000645 |
| **Some Participation** | M | 3.77 | 0.000645 |
| **Measure** | **race.4level** | **StdResidual** | **P-value** |
| **No Participation** | White | -14.45 | 2.14e-46 |
| **Some Participation** | White | 14.45 | 2.14e-46 |
| **No Participation** | Black | 14.74 | 2.90e-48 |
| **Some Participation** | Black | -14.74 | 2.90e-48 |
| **No Participation** | Asian | -1.69 | 7.27e-01 |
| **Some Participation** | Asian | 1.69 | 7.27e-01 |
| **No Participation** | Other/Mixed | 5.15 | 2.06e-06 |
| **Some Participation** | Other/Mixed | -5.15 | 2.06e-06 |
| **Measure** | **hisp** | **StdResidual** | **P-value** |
| **No Participation** | No | -9.08 | 4.43e-19 |
| **Some Participation** | No | 9.08 | 4.43e-19 |
| **No Participation** | Yes | 9.08 | 4.43e-19 |
| **Some Participation** | Yes | -9.08 | 4.43e-19 |
| **Measure** | **household.income** | **StdResidual** | **P-value** |
| **No Participation** | [<50K] | 26.498 | 0 |
| **Some Participation** | [<50K] | -26.498 | 0 |
| **No Participation** | [>=50K & <100K] | 0.454 | 1 |
| **Some Participation** | [>=50K & <100K] | -0.454 | 1 |
| **No Participation** | [>=100K] | -24.702 | 0 |
| **Some Participation** | [>=100K] | 24.702 | 0 |
| **Measure** | **high.educ** | **StdResidual** | **P-value** |
| **No Participation** | < HS Diploma | 12.22 | 2.39e-33 |
| **Some Participation** | < HS Diploma | -12.22 | 2.39e-33 |
| **No Participation** | HS Diploma/GED | 15.32 | 5.62e-52 |
| **Some Participation** | HS Diploma/GED | -15.32 | 5.62e-52 |
| **No Participation** | Some College | 15.79 | 3.64e-55 |
| **Some Participation** | Some College | -15.79 | 3.64e-55 |
| **No Participation** | Bachelor | -6.92 | 4.54e-11 |
| **Some Participation** | Bachelor | 6.92 | 4.54e-11 |
| **No Participation** | Post Graduate Degree | -21.37 | 2.00e-100 |
| **Some Participation** | Post Graduate Degree | 21.37 | 2.00e-100 |
| **Measure** | **pubertal_dev_collapsed** | **StdResidual** | **P-value** |
| **No Participation** | Pre | -7.00 | 2.00e-11 |
| **Some Participation** | Pre | 7.00 | 2.00e-11 |
| **No Participation** | Early | -1.36 | 1.00e+00 |
| **Some Participation** | Early | 1.36 | 1.00e+00 |
| **No Participation** | Mid | 9.23 | 2.12e-19 |
| **Some Participation** | Mid | -9.23 | 2.12e-19 |
| **No Participation** | Late/Post | 1.48 | 1.00e+00 |
| **Some Participation** | Late/Post | -1.48 | 1.00e+00 |

*Supplementary Table 58. DV: Weight status (from BMI). Chi-squared test and pairwise post-hoc comparisons for associations with the sociodemographic factors.*

| **Demographic** | **X.squared** | **DF** | **p.value** |
| --- | --- | --- | --- |
| **age_quartiles** | 5.27 | 9 | 0.81 |
| **sex_at_birth** | 5.54 | 3 | 0.136 |
| **race.4level** | 289 | 9 | 4.51e-57 |
| **hisp** | 202 | 3 | 1.25e-43 |
| **household.income** | 417 | 6 | 5.03e-87 |
| **high.educ** | 529 |  | 5e-04 |
| **pubertal_dev_collapsed** | 383 |  | 5e-04 |
| **Measure** | **age_quartiles** | **StdResidual** | **P-value** |
| **Underweight** | age<=25% | -0.962 | 1 |
| **Healthy Weight** | age<=25% | 0.582 | 1 |
| **Overweight** | age<=25% | -1.066 | 1 |
| **Obese** | age<=25% | 0.763 | 1 |
| **Underweight** | age>25&<=50% | 0.287 | 1 |
| **Healthy Weight** | age>25&<=50% | 0.260 | 1 |
| **Overweight** | age>25&<=50% | -0.303 | 1 |
| **Obese** | age>25&<=50% | -0.194 | 1 |
| **Underweight** | age>50&<=75% | -0.301 | 1 |
| **Healthy Weight** | age>50&<=75% | -1.001 | 1 |
| **Overweight** | age>50&<=75% | 0.700 | 1 |
| **Obese** | age>50&<=75% | 0.776 | 1 |
| **Underweight** | age>75% | 0.999 | 1 |
| **Healthy Weight** | age>75% | 0.111 | 1 |
| **Overweight** | age>75% | 0.734 | 1 |
| **Obese** | age>75% | -1.357 | 1 |
| **Measure** | **sex_at_birth** | **StdResidual** | **P-value** |
| **Underweight** | F | 1.374 | 1.000 |
| **Healthy Weight** | F | 1.250 | 1.000 |
| **Overweight** | F | -0.435 | 1.000 |
| **Obese** | F | -1.901 | 0.458 |
| **Underweight** | M | -1.374 | 1.000 |
| **Healthy Weight** | M | -1.250 | 1.000 |
| **Overweight** | M | 0.435 | 1.000 |
| **Obese** | M | 1.901 | 0.458 |
| **Measure** | **race.4level** | **StdResidual** | **P-value** |
| **Underweight** | White | 2.21 | 4.31e-01 |
| **Healthy Weight** | White | 11.41 | 5.90e-29 |
| **Overweight** | White | -2.88 | 6.32e-02 |
| **Obese** | White | -13.06 | 8.42e-38 |
| **Underweight** | Black | -2.44 | 2.32e-01 |
| **Healthy Weight** | Black | -11.47 | 2.92e-29 |
| **Overweight** | Black | 2.06 | 6.27e-01 |
| **Obese** | Black | 14.04 | 1.37e-43 |
| **Underweight** | Asian | 1.64 | 1.00e+00 |
| **Healthy Weight** | Asian | 2.05 | 6.45e-01 |
| **Overweight** | Asian | -1.05 | 1.00e+00 |
| **Obese** | Asian | -2.49 | 2.06e-01 |
| **Underweight** | Other/Mixed | -1.17 | 1.00e+00 |
| **Healthy Weight** | Other/Mixed | -4.52 | 9.90e-05 |
| **Overweight** | Other/Mixed | 2.13 | 5.37e-01 |
| **Obese** | Other/Mixed | 4.39 | 1.81e-04 |
| **Measure** | **hisp** | **StdResidual** | **P-value** |
| **Underweight** | No | 2.31 | 1.68e-01 |
| **Healthy Weight** | No | 12.64 | 1.01e-35 |
| **Overweight** | No | -6.52 | 5.45e-10 |
| **Obese** | No | -11.23 | 2.42e-28 |
| **Underweight** | Yes | -2.31 | 1.68e-01 |
| **Healthy Weight** | Yes | -12.64 | 1.01e-35 |
| **Overweight** | Yes | 6.52 | 5.45e-10 |
| **Obese** | Yes | 11.23 | 2.42e-28 |
| **Measure** | **household.income** | **StdResidual** | **P-value** |
| **Underweight** | [<50K] | -3.8665 | 1.33e-03 |
| **Healthy Weight** | [<50K] | -14.7873 | 2.12e-48 |
| **Overweight** | [<50K] | 4.5137 | 7.65e-05 |
| **Obese** | [<50K] | 16.7042 | 1.47e-61 |
| **Underweight** | [>=50K & <100K] | 0.6983 | 1.00e+00 |
| **Healthy Weight** | [>=50K & <100K] | -1.1029 | 1.00e+00 |
| **Overweight** | [>=50K & <100K] | 1.0240 | 1.00e+00 |
| **Obese** | [>=50K & <100K] | 0.0823 | 1.00e+00 |
| **Underweight** | [>=100K] | 2.9045 | 4.41e-02 |
| **Healthy Weight** | [>=100K] | 14.5596 | 6.09e-47 |
| **Overweight** | [>=100K] | -5.0728 | 4.70e-06 |
| **Obese** | [>=100K] | -15.3827 | 2.56e-52 |
| **Measure** | **high.educ** | **StdResidual** | **P-value** |
| **Underweight** | < HS Diploma | -1.38461 | 1.00e+00 |
| **Healthy Weight** | < HS Diploma | -7.91562 | 4.92e-14 |
| **Overweight** | < HS Diploma | 2.95546 | 6.24e-02 |
| **Obese** | < HS Diploma | 8.07470 | 1.35e-14 |
| **Underweight** | HS Diploma/GED | -1.57385 | 1.00e+00 |
| **Healthy Weight** | HS Diploma/GED | -9.28954 | 3.10e-19 |
| **Overweight** | HS Diploma/GED | -0.00475 | 1.00e+00 |
| **Obese** | HS Diploma/GED | 12.75626 | 5.75e-36 |
| **Underweight** | Some College | -1.96096 | 9.98e-01 |
| **Healthy Weight** | Some College | -10.40619 | 4.65e-24 |
| **Overweight** | Some College | 4.52908 | 1.18e-04 |
| **Obese** | Some College | 10.07514 | 1.42e-22 |
| **Underweight** | Bachelor | 0.58828 | 1.00e+00 |
| **Healthy Weight** | Bachelor | 5.15017 | 5.21e-06 |
| **Overweight** | Bachelor | -0.64503 | 1.00e+00 |
| **Obese** | Bachelor | -6.30886 | 5.62e-09 |
| **Underweight** | Post Graduate Degree | 2.66331 | 1.55e-01 |
| **Healthy Weight** | Post Graduate Degree | 13.03254 | 1.60e-37 |
| **Overweight** | Post Graduate Degree | -4.64006 | 6.97e-05 |
| **Obese** | Post Graduate Degree | -13.70753 | 1.83e-41 |
| **Measure** | **pubertal_dev_collapsed** | **StdResidual** | **P-value** |
| **Underweight** | Pre | 5.85 | 8.09e-08 |
| **Healthy Weight** | Pre | 13.07 | 8.22e-38 |
| **Overweight** | Pre | -7.83 | 7.84e-14 |
| **Obese** | Pre | -12.36 | 7.16e-34 |
| **Underweight** | Early | -1.20 | 1.00e+00 |
| **Healthy Weight** | Early | -1.47 | 1.00e+00 |
| **Overweight** | Early | 1.51 | 1.00e+00 |
| **Obese** | Early | 1.07 | 1.00e+00 |
| **Underweight** | Mid | -5.07 | 6.40e-06 |
| **Healthy Weight** | Mid | -11.90 | 1.88e-31 |
| **Overweight** | Mid | 6.09 | 1.85e-08 |
| **Obese** | Mid | 12.12 | 1.36e-32 |
| **Underweight** | Late/Post | -2.24 | 4.02e-01 |
| **Healthy Weight** | Late/Post | -7.27 | 5.55e-12 |
| **Overweight** | Late/Post | 5.82 | 9.64e-08 |
| **Obese** | Late/Post | 4.97 | 1.08e-05 |

*Supplementary Table 59. DV: Total medical problems. Chi-squared test and pairwise post-hoc comparisons for associations with the sociodemographic factors.*

| **Demographic** | **X.squared** | **DF** | **p.value** |
| --- | --- | --- | --- |
| **age_quartiles** | 11.6 | 6 | 0.0721 |
| **sex_at_birth** | 8.45 | 2 | 0.0146 |
| **race.4level** | 13.5 | 6 | 0.0361 |
| **hisp** | 17.2 | 2 | 0.000181 |
| **household.income** | 27.7 | 4 | 1.46e-05 |
| **high.educ** | 67.9 | 8 | 1.3e-11 |
| **pubertal_dev_collapsed** | 18.3 | 6 | 0.00545 |
| **Measure** | **age_quartiles** | **StdResidual** | **P-value** |
| **None** | age<=25% | -0.00448 | 1.0000 |
| **One problem** | age<=25% | 1.90552 | 0.6810 |
| **Two or more problems** | age<=25% | -1.78058 | 0.9000 |
| **None** | age>25&<=50% | 0.16207 | 1.0000 |
| **One problem** | age>25&<=50% | -0.51417 | 1.0000 |
| **Two or more problems** | age>25&<=50% | 0.33182 | 1.0000 |
| **None** | age>50&<=75% | 0.45502 | 1.0000 |
| **One problem** | age>50&<=75% | 0.94898 | 1.0000 |
| **Two or more problems** | age>50&<=75% | -1.30926 | 1.0000 |
| **None** | age>75% | -0.61186 | 1.0000 |
| **One problem** | age>75% | -2.38542 | 0.2050 |
| **Two or more problems** | age>75% | 2.79957 | 0.0614 |
| **Measure** | **sex_at_birth** | **StdResidual** | **P-value** |
| **None** | F | 2.358 | 0.1100 |
| **One problem** | F | 0.545 | 1.0000 |
| **Two or more problems** | F | -2.690 | 0.0429 |
| **None** | M | -2.358 | 0.1100 |
| **One problem** | M | -0.545 | 1.0000 |
| **Two or more problems** | M | 2.690 | 0.0429 |
| **Measure** | **race.4level** | **StdResidual** | **P-value** |
| **None** | White | -3.523 | 0.00513 |
| **One problem** | White | 1.399 | 1.00000 |
| **Two or more problems** | White | 1.945 | 0.62200 |
| **None** | Black | 2.239 | 0.30200 |
| **One problem** | Black | -0.544 | 1.00000 |
| **Two or more problems** | Black | -1.560 | 1.00000 |
| **None** | Asian | 1.526 | 1.00000 |
| **One problem** | Asian | -0.790 | 1.00000 |
| **Two or more problems** | Asian | -0.670 | 1.00000 |
| **None** | Other/Mixed | 1.732 | 0.99800 |
| **One problem** | Other/Mixed | -0.934 | 1.00000 |
| **Two or more problems** | Other/Mixed | -0.726 | 1.00000 |
| **Measure** | **hisp** | **StdResidual** | **P-value** |
| **None** | No | -4.15 | 0.000199 |
| **One problem** | No | 1.78 | 0.450000 |
| **Two or more problems** | No | 2.17 | 0.181000 |
| **None** | Yes | 4.15 | 0.000199 |
| **One problem** | Yes | -1.78 | 0.450000 |
| **Two or more problems** | Yes | -2.17 | 0.181000 |
| **Measure** | **household.income** | **StdResidual** | **P-value** |
| **None** | [<50K] | 4.9952 | 5.29e-06 |
| **One problem** | [<50K] | -2.9222 | 3.13e-02 |
| **Two or more problems** | [<50K] | -1.8787 | 5.43e-01 |
| **None** | [>=50K & <100K] | -1.1982 | 1.00e+00 |
| **One problem** | [>=50K & <100K] | -0.0447 | 1.00e+00 |
| **Two or more problems** | [>=50K & <100K] | 1.1490 | 1.00e+00 |
| **None** | [>=100K] | -3.4769 | 4.57e-03 |
| **One problem** | [>=100K] | 2.7159 | 5.95e-02 |
| **Two or more problems** | [>=100K] | 0.6690 | 1.00e+00 |
| **Measure** | **high.educ** | **StdResidual** | **P-value** |
| **None** | < HS Diploma | 5.300 | 1.73e-06 |
| **One problem** | < HS Diploma | -1.792 | 1.00e+00 |
| **Two or more problems** | < HS Diploma | -3.219 | 1.93e-02 |
| **None** | HS Diploma/GED | 5.038 | 7.06e-06 |
| **One problem** | HS Diploma/GED | -2.049 | 6.07e-01 |
| **Two or more problems** | HS Diploma/GED | -2.736 | 9.32e-02 |
| **None** | Some College | 0.905 | 1.00e+00 |
| **One problem** | Some College | -1.983 | 7.10e-01 |
| **Two or more problems** | Some College | 1.022 | 1.00e+00 |
| **None** | Bachelor | -1.717 | 1.00e+00 |
| **One problem** | Bachelor | 1.872 | 9.18e-01 |
| **Two or more problems** | Bachelor | -0.167 | 1.00e+00 |
| **None** | Post Graduate Degree | -4.145 | 5.10e-04 |
| **One problem** | Post Graduate Degree | 1.936 | 7.93e-01 |
| **Two or more problems** | Post Graduate Degree | 2.017 | 6.56e-01 |
| **Measure** | **pubertal_dev_collapsed** | **StdResidual** | **P-value** |
| **None** | Pre | 1.21897 | 1.0000 |
| **One problem** | Pre | 1.95113 | 0.6120 |
| **Two or more problems** | Pre | -2.95379 | 0.0377 |
| **None** | Early | -2.99001 | 0.0335 |
| **One problem** | Early | -0.00384 | 1.0000 |
| **Two or more problems** | Early | 2.76644 | 0.0680 |
| **None** | Mid | 1.77800 | 0.9050 |
| **One problem** | Mid | -2.12119 | 0.4070 |
| **Two or more problems** | Mid | 0.34379 | 1.0000 |
| **None** | Late/Post | -0.70106 | 1.0000 |
| **One problem** | Late/Post | -0.64314 | 1.0000 |
| **Two or more problems** | Late/Post | 1.25016 | 1.0000 |

*Supplementary Table 60. DV: Traumatic Brain Injury (TBI). Chi-squared test and pairwise post-hoc comparisons for associations with the sociodemographic factors.*

| **Demographic** | **X.squared** | **DF** | **p.value** |
| --- | --- | --- | --- |
| **age_quartiles** | 10.2 | 6 | 0.118 |
| **sex_at_birth** | 11.1 | 2 | 0.00389 |
| **race.4level** | 7.6 |  | 0.256 |
| **hisp** | 7.3 | 2 | 0.026 |
| **household.income** | 15.3 | 4 | 0.00412 |
| **high.educ** | 12.1 |  | 0.152 |
| **pubertal_dev_collapsed** | 2.52 |  | 0.865 |
| **Measure** | **age_quartiles** | **StdResidual** | **P-value** |
| **Improbable TBI** | age<=25% | 2.0400 | 0.496 |
| **Possible mild TBI** | age<=25% | -2.4946 | 0.151 |
| **TBI** | age<=25% | 0.0572 | 1.000 |
| **Improbable TBI** | age>25&<=50% | 0.3005 | 1.000 |
| **Possible mild TBI** | age>25&<=50% | 0.0120 | 1.000 |
| **TBI** | age>25&<=50% | -0.5427 | 1.000 |
| **Improbable TBI** | age>50&<=75% | -0.1805 | 1.000 |
| **Possible mild TBI** | age>50&<=75% | 0.9205 | 1.000 |
| **TBI** | age>50&<=75% | -1.0215 | 1.000 |
| **Improbable TBI** | age>75% | -2.2595 | 0.286 |
| **Possible mild TBI** | age>75% | 1.6786 | 1.000 |
| **TBI** | age>75% | 1.5116 | 1.000 |
| **Measure** | **sex_at_birth** | **StdResidual** | **P-value** |
| **Improbable TBI** | F | 3.29 | 0.00605 |
| **Possible mild TBI** | F | -3.00 | 0.01600 |
| **TBI** | F | -1.39 | 0.99500 |
| **Improbable TBI** | M | -3.29 | 0.00605 |
| **Possible mild TBI** | M | 3.00 | 0.01600 |
| **TBI** | M | 1.39 | 0.99500 |
| **Measure** | **race.4level** | **StdResidual** | **P-value** |
| **Improbable TBI** | White | -2.230 | 0.309 |
| **Possible mild TBI** | White | 1.980 | 0.572 |
| **TBI** | White | 1.022 | 1.000 |
| **Improbable TBI** | Black | 2.328 | 0.239 |
| **Possible mild TBI** | Black | -2.004 | 0.541 |
| **TBI** | Black | -1.159 | 1.000 |
| **Improbable TBI** | Asian | 0.685 | 1.000 |
| **Possible mild TBI** | Asian | -0.158 | 1.000 |
| **TBI** | Asian | -0.967 | 1.000 |
| **Improbable TBI** | Other/Mixed | 0.366 | 1.000 |
| **Possible mild TBI** | Other/Mixed | -0.565 | 1.000 |
| **TBI** | Other/Mixed | 0.180 | 1.000 |
| **Measure** | **hisp** | **StdResidual** | **P-value** |
| **Improbable TBI** | No | -1.562 | 0.7100 |
| **Possible mild TBI** | No | 2.542 | 0.0662 |
| **TBI** | No | -0.962 | 1.0000 |
| **Improbable TBI** | Yes | 1.562 | 0.7100 |
| **Possible mild TBI** | Yes | -2.542 | 0.0662 |
| **TBI** | Yes | 0.962 | 1.0000 |
| **Measure** | **household.income** | **StdResidual** | **P-value** |
| **Improbable TBI** | [<50K] | 3.313 | 0.00831 |
| **Possible mild TBI** | [<50K] | -2.970 | 0.02680 |
| **TBI** | [<50K] | -1.477 | 1.00000 |
| **Improbable TBI** | [>=50K & <100K] | 0.516 | 1.00000 |
| **Possible mild TBI** | [>=50K & <100K] | -0.231 | 1.00000 |
| **TBI** | [>=50K & <100K] | -0.566 | 1.00000 |
| **Improbable TBI** | [>=100K] | -3.508 | 0.00406 |
| **Possible mild TBI** | [>=100K] | 2.933 | 0.03020 |
| **TBI** | [>=100K] | 1.872 | 0.55100 |
| **Measure** | **high.educ** | **StdResidual** | **P-value** |
| **Improbable TBI** | < HS Diploma | 0.568 | 1.000 |
| **Possible mild TBI** | < HS Diploma | -0.762 | 1.000 |
| **TBI** | < HS Diploma | 0.113 | 1.000 |
| **Improbable TBI** | HS Diploma/GED | 1.907 | 0.849 |
| **Possible mild TBI** | HS Diploma/GED | -1.385 | 1.000 |
| **TBI** | HS Diploma/GED | -1.322 | 1.000 |
| **Improbable TBI** | Some College | 2.152 | 0.471 |
| **Possible mild TBI** | Some College | -1.923 | 0.817 |
| **TBI** | Some College | -0.969 | 1.000 |
| **Improbable TBI** | Bachelor | -1.517 | 1.000 |
| **Possible mild TBI** | Bachelor | 0.916 | 1.000 |
| **TBI** | Bachelor | 1.321 | 1.000 |
| **Improbable TBI** | Post Graduate Degree | -1.865 | 0.932 |
| **Possible mild TBI** | Post Graduate Degree | 1.984 | 0.709 |
| **TBI** | Post Graduate Degree | 0.379 | 1.000 |
| **Measure** | **pubertal_dev_collapsed** | **StdResidual** | **P-value** |
| **Improbable TBI** | Pre | -0.512 | 1 |
| **Possible mild TBI** | Pre | 0.871 | 1 |
| **TBI** | Pre | -0.370 | 1 |
| **Improbable TBI** | Early | -0.330 | 1 |
| **Possible mild TBI** | Early | -0.446 | 1 |
| **TBI** | Early | 1.225 | 1 |
| **Improbable TBI** | Mid | 0.817 | 1 |
| **Possible mild TBI** | Mid | -0.523 | 1 |
| **TBI** | Mid | -0.669 | 1 |
| **Improbable TBI** | Late/Post | 0.429 | 1 |
| **Possible mild TBI** | Late/Post | -0.199 | 1 |
| **TBI** | Late/Post | -0.460 | 1 |
